# Modeling site-and-branch-heterogeneity with GFmix

**DOI:** 10.1101/2025.08.07.669136

**Authors:** Charley G. P. McCarthy, Edward Susko, Andrew J. Roger

**Affiliations:** Institute for Comparative Genomics, Dalhousie University, Halifax, NS B3H 4R2, Canada; Department of Biochemistry and Molecular Biology, Dalhousie University, Halifax, NS B3H 4R2, Canada; Department of Mathematics and Statistics, Dalhousie University, Halifax, NS B3H 4R2, Canada

**Keywords:** Phylogenetics, Mixture models, Compositional heterogeneity, Simulation

## Abstract

Phylogenetic trees are often inferred from protein sequences sampled from diverse taxa across the tree of life. The compositions of these amino acid sequences may be heterogeneous across both sites and branches, particularly if deep phylogenetic divergences are the focus. Under some conditions, failure to model this compositional heterogeneity can lead to phylogenetic artefacts. However, the computational cost of phylogenetic inference with models accounting for compositional heterogeneity can be prohibitive.

The originally proposed site-and-branch-heterogeneous GFmix model accounts for changing relative frequencies of G, A, R, and P (GARP) vs. F, Y, M, I, N, and K (FYMINK) amino acids resulting from extreme variation in G+C content among taxa. This GFmix model modifies a fitted site-heterogeneous profile mixture model in a branch-specific manner using parameters that reflect branch-specific amino acid compositions. This approach has been shown to improve likelihoods and reduce compositional artifacts. However, the original implementation of the model includes constraints which may sacrifice accuracy for computability and is limited to modeling variation in GARP/FYMINK composition.

Here we investigate the properties of the original GFmix model in greater depth and present several improvements to the model. The improved GFmix models permit fewer constraints on branch-specific composition parameters, allow modeling of user-defined compositional heterogeneity, and provide for full maximum-likelihood optimization of parameters. We have also developed new methods for detecting compositional heterogeneity directly from sequence data. Analyses of simulated site-and-branch-heterogeneous data indicates that the improved GFmix models better estimate branch-specific compositions and branch lengths in heterogeneous trees.

We applied the various versions of the GFmix model to a real dataset with known compositional heterogeneity artefacts. We find that the most complex GFmix model with full maximum likelihood parameter optimization consistently supports the correct tree over the artefactual tree with improved likelihoods. All versions of the GFmix model are available from https://www.mathstat.dal.ca/~tsusko/software.html.

## Introduction

Deep phylogenies are often inferred from aligned sequences of multiple proteins retrieved from many species with very different lifestyles and histories from across the tree of life. Increasing the number of sequences and/or species sampled for deeper phylogenies may not always be beneficial (Philippe et al. 2011). Resolving tricky relationships like locating the roots of the animals and land plants subtrees has remained difficult, even with improved data quality and sampling (Feuda et al. 2017; Whelan et al. 2017; Wickett et al. 2014; Puttick et al. 2018). Many of these deep-time phylogenetic problems have remained difficult to resolve in part because most phylogenetic models do not capture the complexity of sequence evolution at this deep evolutionary scale and are therefore prone to systematic errors in tree inference. Substitution processes across sequences may be heterogeneous due to site-specific functional or structural constraints (Halpern and Bruno 1998; Goldstein 2008). Amino acid composition across species may be heterogeneous due to genomic changes arising from loss of DNA repair mechanisms, genome reduction or adaptation to extreme environments (Acosta et al. 2015; Zeldovich et al. 2007; Siglioccolo et al. 2011). Failing to model or mismodeling such heterogeneity can induce long-branch attraction (LBA) artefacts and lead to incorrect tree estimation (Felsenstein 1978; Foster 2004; Brinkmann et al. 2005).

The simplest protein sequence evolution models typically do not capture the heterogeneity of the substitution process over sites and over branches. Amino acid substitutions over the tree at a site are modeled as occurring according to a Markov process with rate matrix *Q*_*ij*_ = *S*_*ij*_*π*_*j*_. The stationary frequencies, *π*_*j*_, are usually estimated from the data, but the exchangeability rates between amino acids *i* and *j, S*_*ij*_, are fixed. Common exchangability matrices such as LG and WAG were estimated based on large empirical data sets. WAG was estimated assuming homogeneous evolutionary rates and substitution processes across all sites (Whelan and Goldman 2001), whereas LG was optimized allowing for differing rates across sites (Le and Gascuel 2008). Heterogeneity of process over sites is typically accommodated *via* mixtures of these base models where some parameter in the base model differs across mixture classes. For instance, heterogeneity of evolutionary rates across sites is typically modeled using discrete rate multipliers derived from the Gamma (Γ) distribution or estimated from data (Yang 1994; Soubrier et al. 2012). Heterogeneous substitution processes across sites have been accommodated in several ways. In some cases, mixtures of empirically-estimated rate matrices are used (Le et al. 2012) or alternatively different site classes are modeled by using a common set of exchangeabilities and a mixture of frequency profiles (Si Quang et al. 2008). The latter profile mixture models allow the *π*_*j*_ to vary over mixture classes and assume that all substitution patterns across sites can be adequately described using a set of equilibrium amino acid frequency profiles. The frequency profiles may be derived from empirical data as in the C-series (Si Quang et al. 2008) and UDM (Schrempf et al. 2020) models, estimated *via* a Dirichlet process as in the Bayesian CAT model (Lartillot and Philippe 2004) or estimated by maximum likelihood (ML) using the composite likelihood method MAMMaL (Susko et al. 2018). Combining site profile models with site-rate heterogeneity models - e.g. LG+C20+Γor CAT-GTR+Γ- is common in large-scale phylogenetics studies and is readily implemented in software such as IQ-TREE and PhyloBayes (Lartillot et al. 2009; Minh et al. 2020; Wong et al. 2025).

Heterogeneity of amino acid composition across species has been modeled using several approaches. In the ML framework, the branch non-homogeneous and non-stationary Correspondence and Likelihood Analysis (COaLA) model uses correspondence analysis to capture the most important features of compositional variation observed in the data while minimizing the number of parameters requiring optimization (Groussin et al. 2013). In the Bayesian framework, the node-discrete compositional heterogeneity (NDCH) model fits multiple amino acid composition vectors across a tree (Foster 2004; Foster et al. 2009) whereas the CAT-BP model varies the CAT model across a tree to reflect branch-specific amino acid composition (Blanquart and Lartillot 2006; Blanquart and Lartillot 2008). Of these three models, only CAT-BP accommodates both across-site-and across-branch-heterogeneity (Blanquart and Lartillot 2008). Applying the CAT-BP model requires substantial time and computational overhead, and because it is implemented in a Bayesian framework, final interpretations of results are dependent on tree and parameter convergence (Williams et al. 2021). Similar caveats apply to NDCH and the site-heterogeneous CAT-GTR+Γmodel (Williams et al. 2021). Alternative ways to address branch-heterogeneity in amino acid composition have been through data curation by recoding amino acids into larger biochemical groups or removing very heterogeneous sites in a dataset (Embley et al. 2003; Susko and Roger 2007; Whelan et al. 2017; Martijn et al. 2018). Both approaches may improve model fit and parameter convergence for heterogeneous data (Feuda et al. 2017; Whelan et al. 2017). However, the efficacy of recoding approaches has been questioned (Li et al. 2021; Hernandez and Ryan 2021; Foster et al. 2022) and site-removal approaches may yield unexpected results (Francis and Canfield 2020).

GFmix is a novel site-and-branch-heterogeneous model implemented in a maximum-likelihood (ML) framework. GFmix was initially developed to resolve several conflicting relationships within the mitochondria-Alphaproteobacteria tree (Muñoz-Gómez et al. 2022). Mitochondria and multiple alphaproteobacterial phyla have undergone convergent genome reduction, which is correlated with increased genomic AT-content and elevated FYMINK-content in their proteomes (Ettema and Andersson 2009; Hershberg and Petrov 2010; Acosta et al. 2015). Other alphaproteobacterial phyla have comparatively GC-rich genomes and GARP-rich proteomes. This GARP/FYMINK heterogeneity was thought to drive conflicting placements of mitochondria with respect to Alphaproteobacteria, and the monophyly of FYMINK-rich alphaproteobacterial phyla (Martijn et al. 2018; Muñoz-Gómez et al. 2019; Fan et al. 2020). Application of the GFmix model to the mitochondria-Alphaproteobacteria tree favored placing mitochondria as sister to Alphaproteobacteria and multiple independent origins of FYMINK-rich proteomes within Alphaproteobacteria (Muñoz-Gómez et al. 2022). The model has subsequently been extended to other cases of compositional heterogeneity affecting the root of eukaryotes and relationships within Archaea (Baker et al. 2024; Baker et al. 2025; Williamson et al. 2025).

The GFmix model estimates a branch-specific parameter for each branch of a tree and uses this parameter to make branch-specific modifications to the mixture class frequencies of a previously-fitted profile mixture model (Muñoz-Gómez et al. 2022). The likelihood of the tree given this modified model is then re-estimated. Although GFmix improves tree inference in the presence of compositional heterogeneity, there are limitations to the original implementation. No re-optimization of model and branch length parameters occurs before likelihood re-estimation under GFmix. The branch-specific parameters used to modify the mixture model are estimated using the tree and the observed frequencies of amino acids for the taxa under study and impose parameter constraints (Muñoz-Gómez et al. 2022). The approach is thus not a full maximum likelihood implementation. Although this makes it computationally less intensive, which can be valuable for large data sets, it makes parameter estimation less accurate. Finally, in its original implementation, the model only accounts for GARP/FYMINK heterogeneity resulting from GC-content variation (Muñoz-Gómez et al. 2022).

Here, we present improved implementations of the GFmix model intended to loosen the constraints of the original implementation. All implementations now allow user-defined compositional heterogeneity to be modelled, and the most extensive implementations allow some or all model and tree parameters to be re-optimized in a maximum-likelihood framework. We assessed the performance of each implementation of the GFmix model using 16-taxa and downsampled 4-taxa datasets simulated under site-and-branch-heterogeneous GARP/FYMINK conditions. We find that the improved GFmix models more accurately estimate branch lengths and compositional shifts in 16-taxa simulations, and that this improved performance is also observed in more information-sparse 4-taxa simulations. Modeling GARP/FYMINK heterogeneity is appealing because of the connection that those groups of amino acids have to GC-content but, in some settings, there is reason to expect that the predominant heterogeneity will be for different groups (e.g. in Baker et al. (2024) the heterogeneity in halophilic Archaea is between DE/IK).Accordingly, we have developed several methods to identify the the most variable groups of amino acids. We find that the identification methods that use model-based optimization criteria more accurately identify the true groups of varying amino acid compositions in simulations, albeit with an associated increase in computational runtime.

Finally, we applied each GFmix implementation to a real dataset with known compositional heterogeneity arising from GC-content variation. We chose as our basis a 54-taxon dataset comprised of nuclear-encoded protein sequences from Rhodophyta and Viridiplantae and nucleomorph-encoded protein sequences from Cryptomonada (Novak et al. 2024). Nucleomorph genomes are highly reduced, and thus AT- and FYMINK-rich at the genomic and proteomic level, respectively (Moore and Archibald 2009). We downsampled the original dataset, retaining the *Cryptomonas paramecium* nucleomorph, and added nucleomorph-encoded protein sequences from the Rhizarian alga *Bigelowiella natans* (Gilson et al. 2006). We compared support for two hypotheses - one in which the *B. natans* and *Cryptomonas paramecium* nucleomorphs branch together at the base of Rhodophyta, and one in which the nucleomorphs branch separately within Rhodophyta and Viridiplantae based on the biological consensus of their respective origins (Ishida et al. 1999; Douglas et al. 2001). We find that the most extensive GFmix implementation consistently supports the latter hypothesis, with improved likelihoods under newer implementations of the model. Overall, our results indicate that GFmix improves likelihood and reduces compositional heterogeneity artifacts, and that newer implementations of the model show substantial improvements in performance over the initial implementation.

## Materials and Methods

### Implementations of GFmix

*The GFmix Model* The GFmix model for the evolution of an amino acid at a site along a tree builds on the usual base model where, conditional upon the ancestral amino acid, evolution along a branch occurs independently of neighbouring or ancestral branches and according to a continuous-time, time-reversible Markov chain. Time reversibility implies that the Markov chain rate matrix can be decomposed as *Q*_*ij*_ ∝*S*_*ij*_*π*_*j*_ for *i* ≠ *j*, where *π*_*j*_ is the stationary frequency of amino acid *j* and *S* is a symmetric exchangeability matrix with positive entries. Here *Q*_*ii*_ = −∑_*j*∣*j*≠ *i*_ *Q*_*ij*_ and the proportionality constant is determined by *−∑*_*i*_ *π*_*i*_*Q*_*ii*_ = 1 which leads to branch lengths being interpretable as expected numbers of substitutions. We use the fixed LG exchangeability matrix of Le and Gascuel (2008) in the GFmix model; no parameters in *S* require estimation. To emphasize dependence of *Q* on ***π*** we use the notation *Q*(***π***).

Let *z* denote the amino acids at all of the nodes for a site, including internal nodes, and let *p*(*e*) and *c*(*e*) denote the parent and child nodes for a branch *e*. Under the base model, the probability of *z* is a product over edges of matrix exponentials, multiplied by the frequency of the amino acid, *z*_*r*_, at the root,

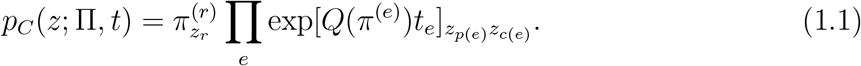

We have allowed that the frequencies Π = [*π*^(1)^ *… π*^(2*m−*2)^] differ over branches or at the root. In the base model these would be the same for each branch and any root location would give the same probability. Decomposing *z* as [*x, y*], where *y* are the amino acids at internal nodes, the probability of the observed data, *x*, is obtained by summing (1.1) over the unobserved internal node data,

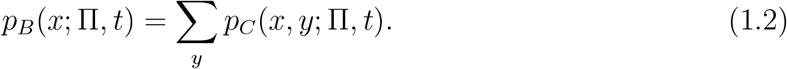

The pruning algorithm of Felsenstein (1973) is used to feasibly calculate (1.2).

Profile mixture models (Si Quang et al. 2008) and the discretized Gamma rate model of Yang (1994) allow variation of stationary frequencies and evolutionary rates over sites. They assume frequency classes and rates at sites are drawn from discrete distributions, independenty of each other and independently across sites. The frequency distribution usually has fixed frequency class profiles, *π*^(*c*)^, that are drawn with probability *w*_*c*_, the latter requiring estimation. We use the C-series frequencies (Si Quang et al. 2008) throughout our examples, but each GFmix implementation allows other profiles to be input. The rate distribution is the discretized Gamma distribution of Yang (1994) where rate *r*_*k*_(*α*) arises with probability 1*/K*; *α* is the shape parameter of the distribution and *K* (set to 4 in examples) the number of rates. Under these models, the conditional probability of the data at a site with rate class *k* and frequency class *c* is *p*_*M*_ (*x*| *c, k*; *π, t, α*) = *p*_*B*_(*x*; Π^(*c*)^, *r*_*k*_(*α*)*t*) where Π^(*c*)^ = [*π*^(*c*)^ *… π*^(*c*)^] and *p*_*B*_(*x*; Π^(*c*)^, *t*) is determined from (1.1)-(1.2). Since (*c, k*) are unobserved, the probability of the observed data *x* is calculated as

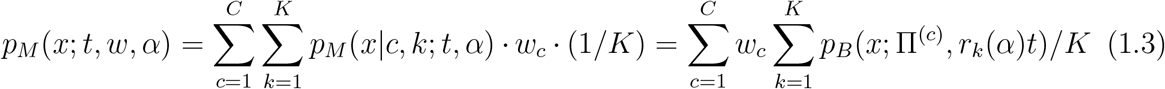

The model described has fixed stationary frequencies of amino acids over branches for any given profile class *c*. GFmix extends the model by allowing these frequencies to vary over the tree but in a specific manner

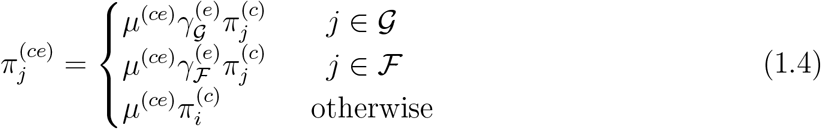

where 𝒢 and ℱ are groups of amino acids; 𝒢= *GARP* and ℱ= *FYMINK* in the original implementation. The additional branch-specific parameters 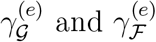 require estimation but the multiplicative constants 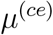 are implied by the constraint 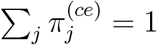. The additional parameters 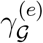 and 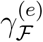 assumes that the biases towards amino acids in𝒢, *ℱ*or𝒪, biases that are determined by 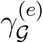 and 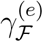, applies over all sites regardless of their rate and frequency profile. Let Π^(*c*)^ (*γ*) = [*π*^(*c*1)^*… π*^(*c*2*m−*2)^]; note that the root edge also has a *γ*^(*r*)^ parameter vector. The site likelihood is calculated similarly as in (1.3):

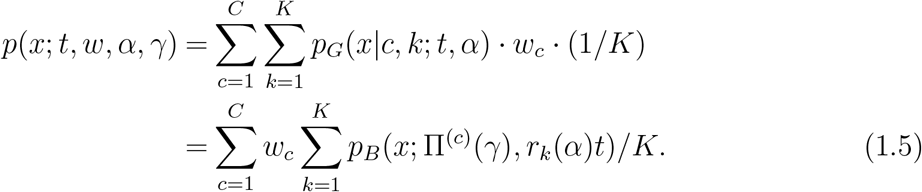

The GFmix methods described below differ largely in how they provide parameter estimates for the model. Given a set of parameters, the end result is the same, a log likelihood for the model described above. That log likelihood assumes evolution over sites is independent and thus is obtained as follows

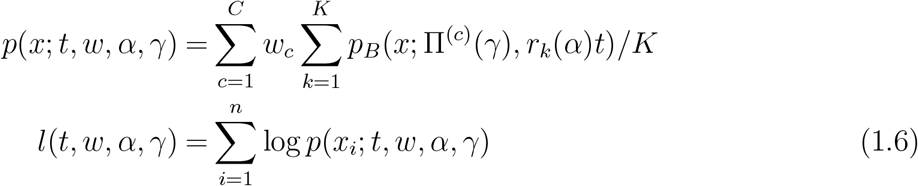

#### Maximum-likelihood implementations - GFP and GFF

The maximum likelihood implementations of GFmix are conceptually the most straightforward. They simply maximize the log likelihood (1.6) using the L-BFGS-B algorithm implementation of Zhu et al. (1997). GFF optimizes all parameters. Partial optimization over *γ* is done by GFP, holding all other parameters fixed. The other parameters are estimated by maximum likelihood using IQ-TREE under the corresponding profile mixture model (equivalently with 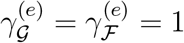).

#### Original GFmix implemention - OGF

The original GFmix implementation was first described in Muñoz-Gómez et al. (2022) for 𝒢*/*ℱ equal to GARP/FYMINK and modified in Baker et al. (2024) to accommodate arbitrary 𝒢 and ℱ. For a given branch *e*, assume that we know *b*_*e*_, the ratio of the total expected frequency ratio of ℱamino acids to the total frequency of 𝒢 amino acids; OGF estimates *b*_*e*_ from the vectors, over taxa, of observed frequencies of amino acids as described below. The 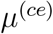 and 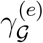 and 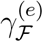 then satisfy the following equations

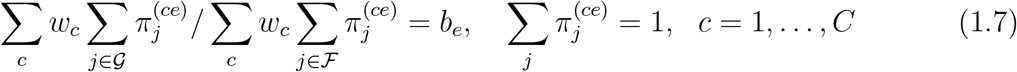

This gives *C* + 1 equations in *C* + 2 unknowns. The original GFmix implementation imposed the additional model condition

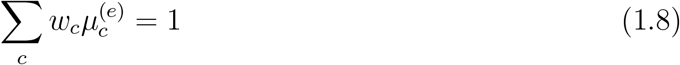

This constraint implies that the weighted average multiplier of the 𝒪 class is 1. That constraint is not essential and not present the maximum likelihood implementations GFP and GFF.

To estimate the *b*_*e*_, OGF aggregates all sequences that descend from branch *e* and then estimates *b*_*e*_ as the ratio of the total observed frequency of amino acids in 𝒢 to total observed frequency of amino acids in ℱ. It then solves the *C* +2 equations in *C* +2 unknowns given in (1.7)-(1.8) to obtain estimates of 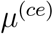 and 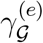 and 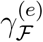.

#### Sum of squares difference between node and model frequencies - SSF

For a given frequency class *c* and rate class *k*, the conditional probability of amino acid *j* at the root is 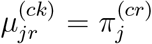. Given the conditional probabilities at the parental node *p*(*e*) of branch *e* probabilities at the child node c(*e*) can be calculated through

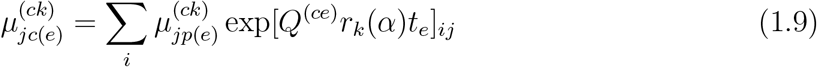

Starting with 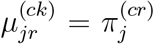, (1.9) can be used recursively, traversing edges in the tree from root to tip to calculate 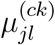 for all nodes *l*. The expected frequency of amino acid *j* is then obtained as 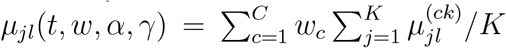, were we now explicitly indicate dependence on parameters.

We use the algorithm just described to get node frequencies in simulations where we know the parameter values. We also used it to obtain estimates of *γ* using least squares. Fixing 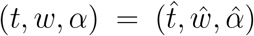 at maximum likelihood likelihood values obtained under the corresponding profile mixture model (equivalently with 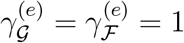), we minimize

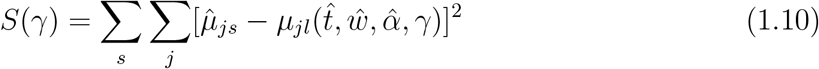

where sums are over terminal nodes *s* and amino acids *j*.

### Determining compositionally heterogeneous groups

The original GFmix model assumed fixed groups, 𝒢*/*ℱ = GARP/FYMINK. The rationale for this was both empirical, in that overall GARP and FYMINK frequencies often show a lot of variation over taxa, and because these two groups of amino acids are GC (GARP) and AT (FYMINK) rich (Muñoz-Gómez et al. 2022). The improved GFmix models now allow 𝒢*/*ℱ group determination to be based on the data at hand using methods that we now introduce.

#### Binomial test of two proportions

Compositional heterogeneity is first identified from data using a binomial test of two proportions (Baker et al. 2024; Williamson et al. 2025). This approach requires two apriori specified groups of taxa for which compositional heterogeneity between groups is suspected. For instance, Baker et al. (2024) separated taxa according to whether they exist in hypersaline environments or not. For a given dataset, taxa are divided into two groups. For each amino acid, the composition bias between the two taxon groups is computed as a Z-score in the following equation:

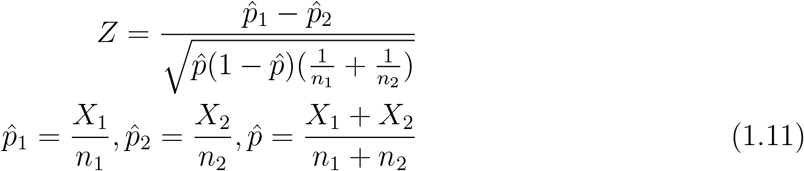

where *X*_1_, *X*_2_ are the total numbers of that amino acid and *n*_1_, *n*_2_ are the total number of all 20 amino acids across the two taxon groups respectively. This test assumes that the proportions of an amino acid across taxa is approximately normal, with the null hypothesis that *p*_1_ = *p*_2_. |*Z*| *>* 1.96 indicates rejection of the null hypothesis at significance level *p <* 0.05. Amino acids are divided into the classes 𝒢 (*Z >* 1.96), ℱ (*Z < −*1.96) or 𝒪 (|*Z*| ⩽ 1.96) on the basis of this *Z*-score.

#### General optimization routine

The main reason for the GFmix model adjustment (1.4) is to allow the frequencies of amino acids within groups to vary over taxa. So we seek groups where the average within-group 𝒢 (similarly ℱ) amino acid frequency shows evidence of substantial variation over edges. Since the simplifying assumption in (1.4) is that frequencies within a class are modified by the same scalar factor, we also seek choices of ℱ and ℱ classes so that frequencies of amino acids within the 𝒢 (or similarly the ℱ) class differ for an edge from the overall frequencies in a similar fashion. If, for instance, 𝒢*/ℱ* =GARP/FYMINK is a good choice, then when the frequency of *G* is elevated for a taxon relative to the overall frequencies, we also expect *A* to be elevated. In other words, the amino acids within a class should covary across edges.

Our approach to meeting the objectives of 𝒢*/ℱ* class selection outlined above is to choose the classes that minimizes a goodness of fit criterion. We describe three different possible criteria below. The results of the binomial approach give starting classes. Given a criterion, optimization of the class memberships follows a non-greedy approach, in which both the amino acids in the current classes and all permissible single-amino acid swaps between classes at a given point in the algorithm are assessed under some optimality criterion. Permissible swaps follow a bipartite relationship between classes: single-amino acid swaps are permitted between 𝒢 and 𝒪, ℱ and 𝒪, but not 𝒢 and ℱ. Once all swaps have been assessed, the best-performing swapped classes are compared to the current classes’ with respect to their optimality criteria. If the swapped classes improve upon the current classes, the swapped classes replace the current classes and the optimization routine repeats until no further improvement can be made. The resulting 𝒢,ℱ and 𝒪 are considered the final classes for this optimization routine.

#### Criterion 1: χ^2^ test of homogeneity

The *χ*^2^ test is a standard measure of compositional homogeneity in phylogenetics (Foster 2004). The test statistic is the standard *χ*^2^ test of homogeneity of multinomial frequencies over groups. In our application, each taxon plays the role of a group and the multinomial frequencies are the summed 𝒢, ℱ and 𝒪 for each taxon. In using the *χ*^2^ test as an optimization criterion, we attempt to identify 𝒢 and ℱ which maximizes the *χ*^2^ test statistic for a given alignment – in other words, the most compositionally-heterogeneous 𝒢 and ℱ. For a given 𝒢 and ℱ, the counts of all amino acids within each class are summed and the *χ*^2^ test is performed with the summed counts using the chisq.test() function in R (R Core Team 2025). Summation of the counts of 𝒢 and ℱ ensures that the *χ*^2^ test is always performed with two degrees of freedom, which means the test statistic is comparable for any size of 𝒢 or ℱ.

#### Criterion 2: Sum-of-squares difference using SSF implementation

SSF estimates branch-composition parameters that minimize the sum, over taxa and amino acids, of squared (SS) differences between the observed frequency of an observed amino acid for a taxa and the corresponding expected frequency under the model. It does this for a given choice of 𝒢 and ℱ but returns as a by-product, the optimal SS for that choice. That SS can also be used as a criterion. We attempt to identify 𝒢 and ℱ which minimize the sum-of-squares difference.

#### Criterion 3: Likelihood using OGF implementation

Finally, the original implementation of GFmix (OGF) can itself be used as optimization criteria. We attempt to identify 𝒢 and ℱ which maximizes the likelihood of a given alignment, tree and mixture model. For the original implementation of GFmix, 𝒢 and ℱ are used to estimate branch-specific composition parameters *b*_*e*_ as detailed above.

### Simulated data analysis

#### Generating 16-taxon datasets

Simulated site-and-branch-heterogeneous sequence data was generated using AliSim, a component of the IQ-TREE 3 software package (Ly-Trong et al. 2022; Wong et al. 2025). Data was generated using a balanced 16-taxon simulating tree, which was divided into four 4-taxon clades and rooted at the midpoint. Clades were labelled A-D with taxa in each clade labelled A1-A4 etc. Branch lengths are indicated in Figure 1. One hundred replicate sequence data sets were generated, each with a length of 10,000 sites with no gaps. Site-heterogeneity was simulated using an LG+C20+Γmodel with four discrete rate categories and *α* = 0.5 (Le and Gascuel 2008; Si Quang et al. 2008; Yang 1994). For branch-heterogeneity, all stem and tip branches in clades B and D were assigned a target *b*_e_ = 0.1, whereas all other branches were assigned a target *b*_e_ = 1 under the assumptions of the original GFmix model (Muñoz-Gómez et al. 2022). As simulating branch-heterogeneity is not a native feature of AliSim, a custom approach was used in which site-and-branch heterogeneous replicates were simulated on a class-by-class basis and these class sub-replicates were concatenated to form the final replicate for downstream analysis. Detailed explanation of this approach is provided in **Supplementary Information**.

**Fig. 1.**
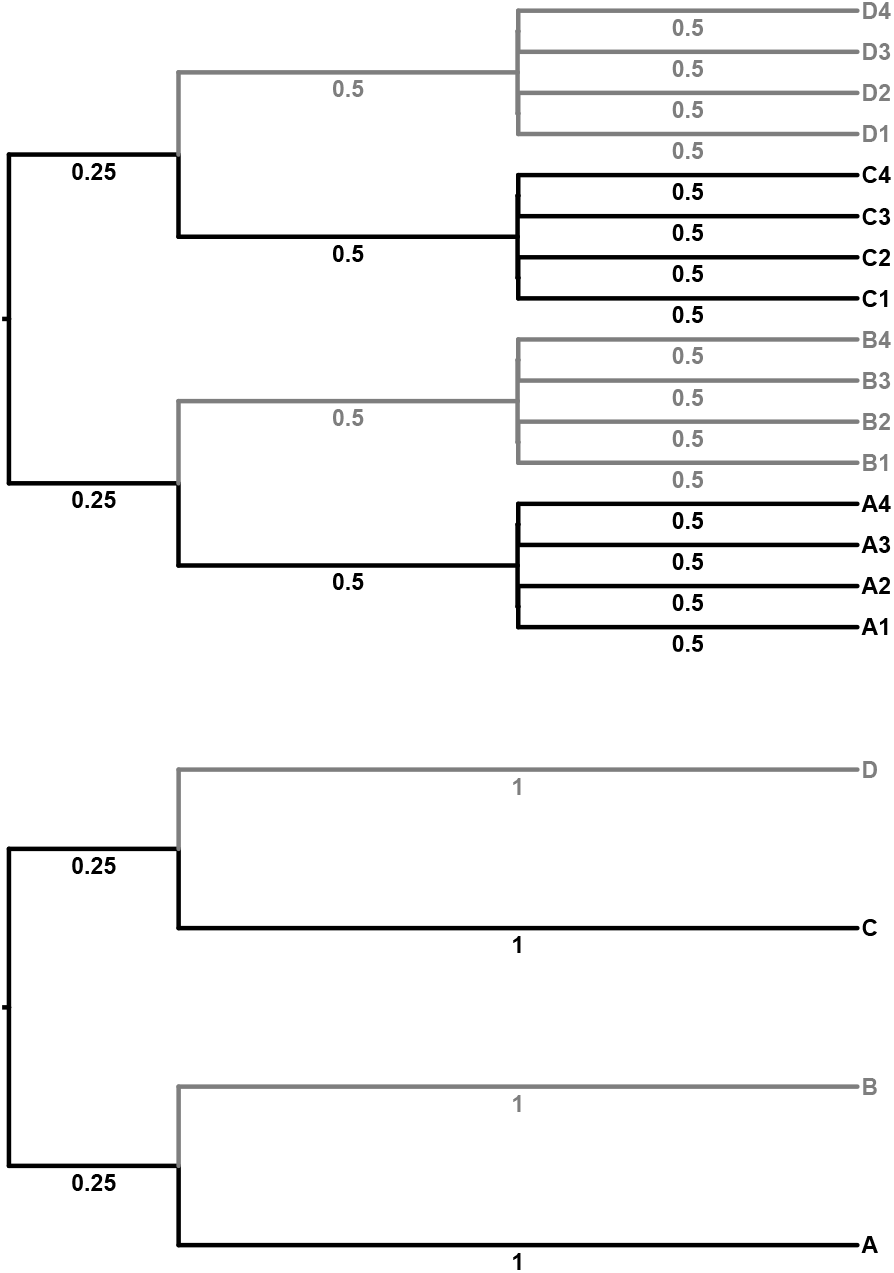
Tree topologies used for simulating and downsampled site-and-branch-heterogeneous sequence data. Branches and taxa coloured in grey have amino acid composition *π*(*GARP*) *< π*(*FY MINK*), branches and taxa coloured in black have amino acid composition *π*(*GARP*) = *π*(*FY MINK*). Simulating branch lengths indicated underneath branches. **Top:** 16-taxon tree. **Bottom:** 4-taxon tree.

#### Generating 4-taxon datasets

Phylogenetic models generally perform better when more taxa are available, particularly if these taxa can break long branches within a tree (Graybeal 1998). To assess model performance in more taxa-sparse cases, all 16-taxon replicates were downsampled to 4-taxa replicates by extracting one taxon from each clade A, B, C and D.

#### Site-heterogeneous model fitting

Site-heterogeneous model fitting was performed for all simulated data using IQ-TREE 3 (Wong et al. 2025), with a LG+C20+Γmodel with mixture weight optimization (Le and Gascuel 2008; Si Quang et al. 2008; Yang 1994) and using the simulating tree as a fixed tree.

#### Branch-heterogeneous model fitting

Branch-heterogeneous model fitting was performed for all simulated data using each of the four GFmix implementations outlined above (Muñoz-Gómez et al. 2022; Baker et al. 2024). For each implementation, the compositional heterogeneity to model was set to GARP/FYMINK.

### Assessment of GFmix model performance

#### Estimating branch lengths with and without maximum-likelihood optimization

We assessed the performance of GFF in re-estimating branch lengths by comparing the observed branch lengths estimated under LG+C20+Γand LG+C20+Γ+GFF against the expected simulating branch lengths. Branch lengths estimated under LG+C20+Γand LG+C20+Γ+GFF were extracted for each branch across all replicates. Root mean squared error (RMSE) for a given estimation method and model parameter is the square root of the average, over the 1000 simulated data sets, of the squared differences between the estimated parameter and the true one. For the 4-taxon data sets these were calculated separately for each external branch. The lengths of the two root branches were summed prior to calculation of a squared error so that the RMSEs were for branch lengths in the corresponding unrooted tree. For 16-taxon data sets, results were summarized for clades by taking averages of branch-specific RMSEs. For example, an RMSE reported for Clade A was calculated as the average of the RMSEs for the tip branches A1-A4 and the RMSE for the stem. The reason for this is that because of the symmetry of the tree and processes, the same behaviour is expected for the A1 and A4 branch, eliminating the need to consider them separately. Plots comparing the difference between simulating branch lengths and observed branch lengths in both 16-taxon and 4-taxon datasets were generated using ggplot2 (Wickham 2016).

#### Estimating node GARP/FYMINK frequency ratio

Under the GFmix model, expected amino acid frequencies vary continuously over the tree. We assessed the ability of methods to estimate the ratio of aggregate GARP/FYMINK frequencies by comparing those ratios calculated with estimated parameters against the true ratios calculated using simulated parameters.

Because of the symmetry of the processes and branch lengths for certain branches, long-run RMSEs of estimation are known to be the same, for instance, for a tip in the A clade and a tip in the C clade. Average RMSE were thus reported to simplify presentation. For the 16-taxa datasets, RMSEs were calculated as follows. For the four simulated clades, GARP/FYMINK ratio RMSEs were separately averaged over A and C tips (A+C tips), A and C stems (A+C stems), B and D tips (B+D tips) and B and D stems (B+D stems). For the two nodes closest to the root, the root mean squared error was calculated as the square root of the mean of the squared error of both nodes combined. RMSEs for the root node were taken from the square root of the mean of the squared error of every root node across all 100 replicates.

For the 4-taxa datasets, RMSEs were averaged over the A and C branches, and, separately, over the B and D branches. Similarly as for the 16-taxon data sets for the two internal nodes, the average RMSE - averaged over the two internal nodes - was calculated as well as the the RMSE for the root node. Plots comparing the difference between simulating GARP/FYMINK frequency ratios and observed GARP/FYMINK frequency ratios in both 16-taxon and 4-taxon datasets were generated using ggplot2 (Wickham 2016).

### Assessing methods of identifying compositional heterogeneity

We assessed the performance of our four methods for identifying compositional heterogeneity — the binomial test of two proportions and the three optimization criteria detailed above — across all 16-taxon and 4-taxon replicates. As the simulating heterogeneity for all replicates was GARP/FYMINK, our assessment was primarily based on how often a method correctly assigned 𝒢 = GARP and ℱ = FYMINK and, secondarily, how often a method misassigned 𝒢,𝒪 and ℱ. Each method of identification was performed across each of the 100 16-taxon and 4-taxon replicates, and the 𝒢,𝒪 and ℱ assignments of each method were tabulated. Unique patterns of 𝒢 /𝒪 /ℱ assignment across all methods were identified, and the number of times each method had made each assignment was tabulated. This 𝒢 / 𝒪 /ℱ assignment per-method information was visualized in an UpSet-like plot using ggplot2 (Lex et al. 2014; Wickham 2016). Additional assessment of each identification method is detailed in **Supplementary Information**.

### Applying GFmix to real data

#### Algal nuclear and nucleomorph dataset

Our real data analysis utilized a dataset designed to determine the position of Cryptomonad nucleomorphs relative to Rhodophyta (Novak et al. 2024), downsampled and modified to include Chlorarachniophyte nucleomorph sequences. Nucleomorphs are highly-reduced nuclear genomes that exist within plastids acquired through separate secondary endosymbosis events in Cryptomonads (Cryptista) and Chlorarachniophyceae (Rhizaria) (Moore and Archibald 2009). These nucleomorph genomes are AT-rich at the genomic level and FYMINK-rich at the proteomic level. Cryptomonad nucleomorphs are generally thought to be derived from a deep-branching Rhodophyte (Douglas et al. 2001) and Chlorarachniophycete nucleomorphs are thought to be derived from a Ulvophyte donor within Viridiplantae (Ishida et al. 1999). Novak et al. (2024) sampled 180 protein sequences across 54 taxa from Rhodophyta, Viridiplantae and four nucleomorph genomes from Cryptomonada. Novak et al. (2024) found strong support for the placement of Cryptomonad nucleomorphs as sister to extremophilic Cyanidiophytina at the base of Rhodophyta.

#### Adding B. natans nucleomorph data to the Novak et al. (2024) dataset

Novak et al. (2024) did not sample any Chlorarachniophycete nucleomorphs in their analysis. To create a data set where the long-branches and composition biases of the two kinds of nucleomorphs may lead to artefactual attraction, we added protein sequence data from the nucleomorph of the Chlorarachniophyceae alga *Bigelowiella natans* (Gilson et al. 2006) to the Novak et al. (2024) dataset as follows. Homologs from the *B. natans* nucleomorph genome were first identified using sequence similarity searches against *Cryptomonas paramecium* nucleomorph proteins in the dataset with BLASTp (e-value = 1*e*^*−*4^) (Altschul et al. 1990), and double-checked by sequence similarity searches against the non-redundant protein sequence database (Sayers et al. 2024). This retained 66 protein sequences from the Novak et al. (2024) dataset with homologous data from the *B. natans* nucleomorph. These sequences were re-aligned using MAFFT and all sequences were concatenated into a single 17,696-site dataset using R (Katoh and Standley 2013; R Core Team 2025).

#### Downsampling Novak et al. (2024) dataset and tree hypotheses

We downsampled the modified Novak et al. (2024) dataset based on phylogenetic diversity, attempting to include at least one member of each major clade from Rhodophyta and Viridiplantae sampled in the original dataset. We retained 16 taxa for our analysis - nine Rhodophyta, five Viridiplantae, and the nucleomorphs of *Cryptomonas paramecium* and *B. natans*.

We chose two tree hypothesis to test for our downsampled dataset following the phylogeny in Novak et al. (2024). The first tree (“NMs Apart”) places the two nucleomorphs as evolving independently: the *C. paramecium* nucleomorph as sister to *Cyanidioschyzon merolae* at the base of Rhodophyta, and the *B. natans* nucleomorph sister to the closest sampled relative of Ulvophyceae - *Volvox carteri* - within Viridiplantae. The second tree (“NMs Together”) places the two nucleomorphs as sister taxa adjacent to *C. merolae* at the base of Rhodophyta. We assume that “NMs Apart” is the correct tree based on biological consensus (Ishida et al. 1999; Douglas et al. 2001; Moore and Archibald 2009; Novak et al. 2024) whereas “NMs Together” is an artefact of long-branch attraction and shared compositional bias.

#### Site-heterogeneous model fitting

Site-heterogeneous model fitting was performed for each dataset using IQ-TREE 3 (Minh et al. 2020) with tree search constrained to the topologies detailed above. The LG+C20+Γmodel was fit for each tree using mixture weight optimization (i.e., the -mwopt flag). The estimated frequency class (+F) was not included in model fitting, due to its potential compromising effects on tree inference (Baños et al. 2023).

#### Branch-heterogeneous model fitting

Branch-heterogeneous model fitting was performed for each dataset using each of the four GFmix implementations outlined above. Trees were rooted between Viridiplantae and Rhodophyta. Initial model fitting was performed using the default 𝒢 / ℱ = GARP/FYMINK. Additional fitting was performed using 𝒢 / ℱ as determined by the bin optimization approach detailed above with the criterion function set to the OGF likelihood (Criterion 3).

## Results

### New implementations of GFmix

GFmix is a site-and-branch-heterogeneous model which modifies a site-heterogeneous profile mixture model to reflect branch-specific amino acid composition (Muñoz-Gómez et al. 2022). The general GFmix model assumes three sets of amino acids 𝒢, 𝒪 and ℱ.𝒢 and are ℱ amino acids which are assumed to be heterogeneous in composition across a tree, whereas 𝒪 is assumed to be homogeneous in composition. In this study, we assessed the performance of various implementations of the GFmix model:

- Original GFmix - OGF: the original implementation of GFmix first outlined in Muñoz- Gómez et al. (2022) and extended to model user-defined 𝒢 / ℱ compositional heterogeneity in Baker et al. (2024).
- SSFreq - SSF: estimating branch-composition parameters that minimize the sum-of-squares differences between observed and model amino acid frequencies over taxa.
- GFmix Partial - GFP: estimating separate branch-specific composition parameters for 𝒢 and ℱ amino acids, *γ*_*𝒢*_ and *γ*_*ℱ*_ respectively, using ML estimation
- GFmix Full - GFF: estimating (or re-estimating) all model, branch-specific composition and branch-length parameters using ML estimation

A brief comparison of each implementation is provided in **Table 1** and further information can be found in **Materials and Methods** and **Supplementary Information**.

**Table 1.** Summary of GFmix implementations assessed in this study, and abbreviations used throughout this study. 𝒢 and ℱ: classes of compositionally heterogeneous amino acids; *b*_*e*_: the estimated frequency ratio *b* of 𝒢 and ℱ amino acids at branch *e* of a tree; 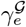 : the scaling constant for 𝒢 amino acids at branch *e* of a tree, 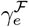 : the scaling constant for ℱ amino acids at branch *e* of a tree, *w*_*c*_: weights of a profile mixture model; *α*: shape parameter of the Γdistribution of site-rate categories.

| Method name | Abbreviation | Method description | Parameters estimated / optimized |
| --- | --- | --- | --- |
| Original GFmix | OGF | Estimate branch-specific composition parameters using non-linear equations | $b_e$ |
| SSFreq | SSF | Estimate branch-specific composition parameters that minimize sum-of-squares difference between observed and model frequencies | $b_e$ |
| GFmix Partial | GFP | Estimate separate branch-specific composition parameters for $\mathcal{G}$ and $\mathcal{F}$ amino acids | $\gamma_e^{\mathcal{G}}, \gamma_e^{\mathcal{F}}$ |
| GFmix Full | GFF | (Re-)estimate all available model and branch-length parameters | Branch lengths, $w_c, \alpha, \gamma_e^{\mathcal{G}}, \gamma_e^{\mathcal{F}}$ |

### Assessing GFmix performance

To assess the performance of each GFmix implementation, we simulated 100 16-taxon alignments under an LG+C20+Γsite-heterogenous model using AliSim (Ly-Trong et al. 2022) with a custom workflow for simulating GARP/FYMINK branch-heterogeneity (see **Materials and Methods** and **Supplementary Information**). We also downsampled these alignments to 4 taxa to assess model performance under more data-sparse conditions (Graybeal 1998). Information on the simulating and downsampled trees for all 16- and 4-taxon replicates is provided in **Figure 1**.

#### Estimation of branch lengths

Three of the four GFmix implementations compared in this study use branch lengths as estimated under the site-heterogeneous LG+C20+Γmodel by IQ-TREE, whereas the full GFmix model (GFF) is able to re-estimate these branch lengths through maximum-likelihood optimization. Thus, our comparison of branch length estimation for our simulated 16-taxon trees was between branch lengths estimated under an “uncorrected” maximum-likelihood model (hence referred to as UML) and branch lengths re-estimated under GFF. For 16-taxon datasets (**Figure 2**), UML consistently underestimates branch lengths for tip and stem branches in the high-FYMINK clades B and D, and for the two internal branches closest to the root. UML also overestimates branch lengths for stem branches in the equal-frequency clades A and C (**Figure 2**). GFF improves branch length estimation across all tip, stem and internal branches - with the tradeoff that there is a larger variance among branch length estimates for the internal branches (**Figure 2**). These trends can also be observed in the root mean squared error (RMSE) calculations for the 16-taxon trees, with the largest error for GFF observed at the internal branches (**Table 2**).

**Table 2.** Comparison of branch length estimation root mean squared error for 16-taxon trees simulated under LG+C20+Γwith GARP/FYMINK heterogeniety.

| Method | A/C | B/D | Internal |
| --- | --- | --- | --- |
| UML | 0.0352 | 0.0572 | 0.0347 |
| GFF | 0.0201 | 0.0195 | 0.101 |

**Fig. 2.**
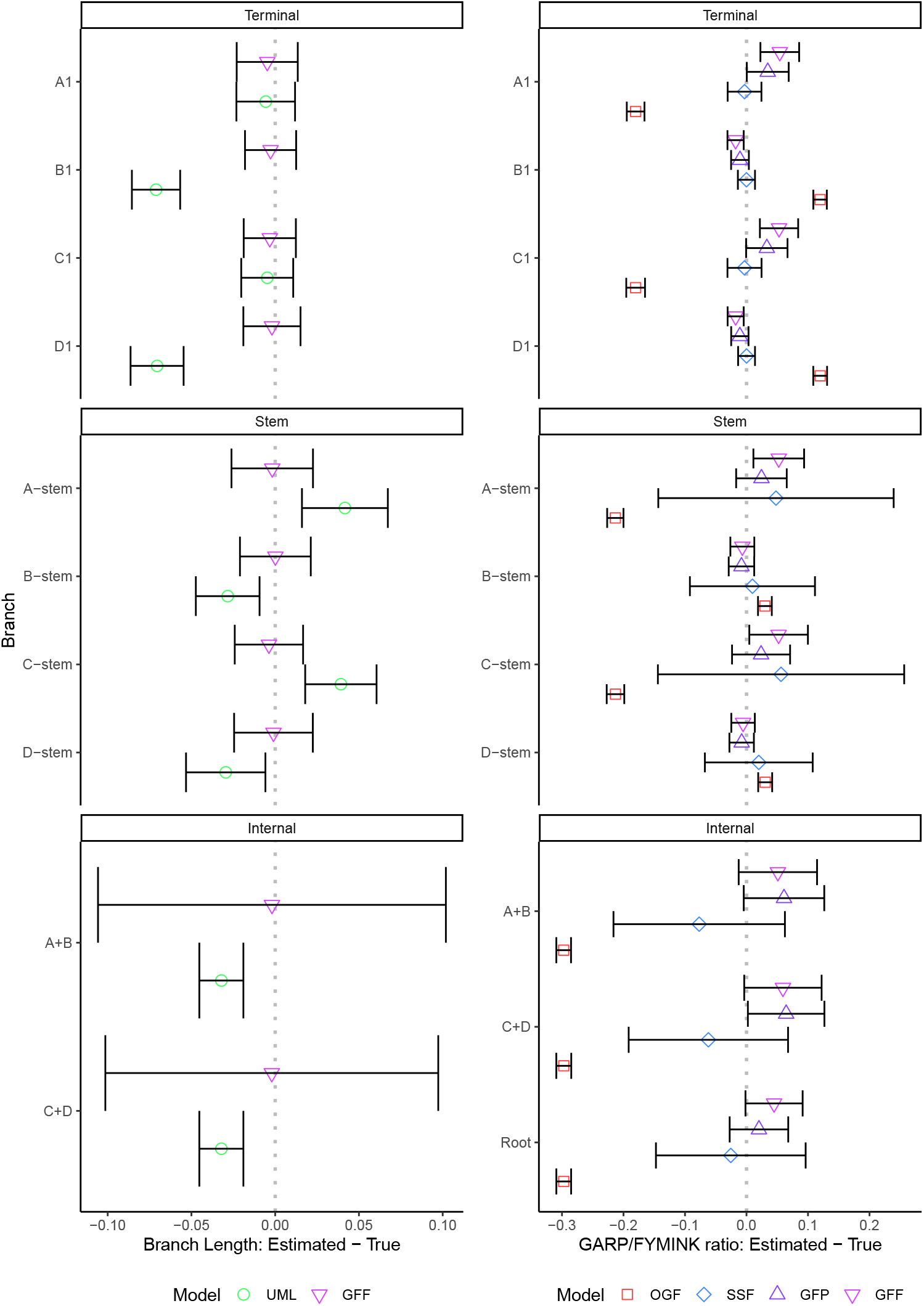
Performance of GFmix implementations across 100 16-taxon replicates simulated under LG+C20+Γ+*{*GARP/FYMINK*}*. Comparisons are between observed estimations and expected estimation based on simulating values. **Left:** Estimation of branch lengths. Comparison made between estimates under LG+C20+Γmodel by IQ-TREE and estimates under LG+C20+Γ+*{*GARP/FYMINK*}* model with maximum-likelihood optimization of all model and tree parameters by GFF. **Right:** Estimation of node GARP/FYMINK composition ratios. Comparison made between estimates by each GFmix implementation under LG+C20+Γ+*{*GARP/FYMINK*}* model. Dots: mean value, errorbars: *±*1*σ*, dotted line: observed and expected are identical.

For the downsampled 4-taxon datasets, UML overestimates the branch lengths for the equal-frequency taxa A and C (**Figure S1**). Estimation markedly improves for these taxa across replicates when branch lengths are re-optimized using GFF, alongside small improvements in estimation for the high-FYMINK taxa B and D (**Figure S1**). As with the 16-taxon datasets, there is a tradeoff in improving tip branch length estimation that results in larger variance in length estimates for internal branches (**Figure S1**). The larger degree of error observed in the 4-taxa RMSEs is likely a reflection of less information in the 4-taxa datasets relative to the 16-taxa datasets (**Table 3**).

**Table 3.**
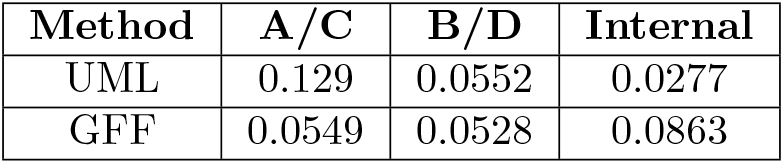
Comparison of branch length estimation root mean squared error for downsampled 4-taxon trees.

#### Estimation of branch GARP/FYMINK composition ratio

We assessed the performance of each of the four GFmix implementations at modelling compositional shifts across heterogeneous 16-taxon trees. This was done by comparing the GARP/FYMINK frequency ratio implied by the simulating LG+C20+Γ+GFmix model (where 𝒢 /ℱ = GARP/FYMINK) with those estimated under the alternative GFmix implementations. Performance varied for each implementation. OGF consistently underestimated GARP/FYMINK composition across the equal-frequency nodes, and overestimated GARP/FYMINK composition for all nodes in the high-FYMINK clades B and D (**Figure 2**). SSF displays improved performance for tip nodes but with substantially larger variance in estimates for stem, internal and root nodes (**Figure 2**). GFG and GFF estimate GARP/FYMINK compositions better and more consistently across all node types, with larger variance estimates for deeper nodes (**Figure 2**). These trends can also be observed in the RMSE calculations (**Table 4**). OGF performs the worst for all node types, SSF performs best in tip composition estimation, whereas GFG and GFF exhibit consistently lower errors across the tree (**Table 4**).

**Table 4.**
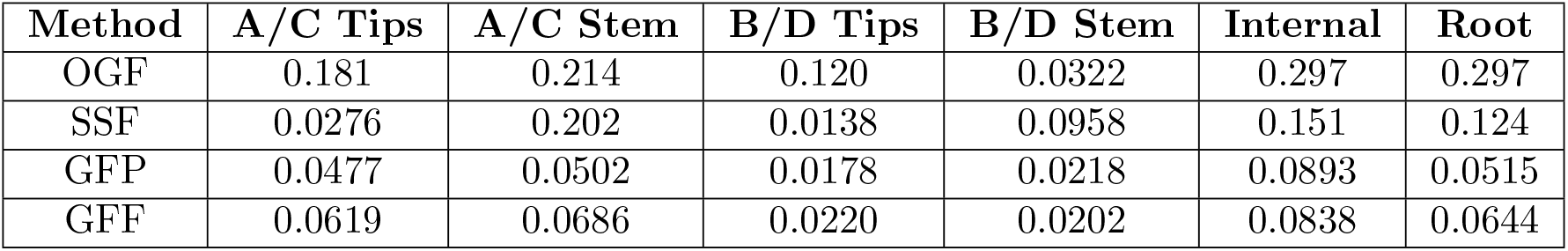
Comparison of GARP/FYMINK frequency ratio estimation root mean squared error for 16-taxon trees simulated under LG+C20+Γwith GARP/FYMINK heterogeniety.

Similar trends are observed in the downsampled 4-taxon datasets. OGF consistently gave biased branch GARP/FYMINK composition estimates across tip, internal and root nodes (**Figure S1**). SSF performs well at estimating tip compositions, which suggests better performance with less taxa, but fares poorer with internal and root composition estimation (**Figure S1**). GFG and GFF estimate GARP/FYMINK compositions better and more consistently across all node types, with GFF performing better for equal-frequency taxa A and C and both internal and root node compositions (**Figure S1**). These trends can also be observed in the RMSE calculations (**Table 5**). OGF performs worst for all node types, SSF performs best in tip composition estimation whereas GFP and GFF exhibit consistently lower errors across the tree (**Table 5**).

**Table 5.** Comparison of branch length estimation root mean squared error for downsampled 4-taxon trees.

| Method | A/C | B/D | Internal | Root |
| --- | --- | --- | --- | --- |
| OGF | 0.178 | 0.119 | 0.297 | 0.297 |
| SSF | 0.0270 | 0.0139 | 0.0959 | 0.110 |
| GFP | 0.0569 | 0.0216 | 0.140 | 0.0631 |
| GFF | 0.0624 | 0.0229 | 0.106 | 0.0737 |

### Identifying compositional heterogeneity

GFmix has been applied in cases of well-known amino acid compositional heterogeneity arising from GC-content variation (Muñoz-Gómez et al. 2022; Williamson et al. 2025; Baker et al. 2025) or adaptation to hypersaline environments (Baker et al. 2024). In the latter case, the amino acids most important to compositional heterogeneity were DE/IK rather than GARP/FYMNK. There may be other cases where underlying groups that are most important to compositional heterogeneity are either not known or not immediately apparent. In such cases, it would be desirable to identify the groups of amino acids directly from sequence data. We investigated how to identify these groups from data using our simulated datasets, which allows comparisons with truth, since GARP/FYMINK are known to be the heterogeneity-determining amino acids in the simulating model. We applied four methods of compositional heterogeneity identification - the binomial test of two proportions previously applied in conjuction with GFmix analysis (Baker et al. 2024; Williamson et al. 2025; Baker et al. 2025) and a new optimization-based procedure using three different statistical criteria. **Figure 3** outlines the workflow of each identification method, and further details are given in **Materials and Methods**. We tabulated the assignments of 𝒢, ℱ and 𝒪 for each method across our 16-taxon and 4-taxon datasets to assess how often each method correctly assigned 𝒢 /ℱ as GARP/FYMINK.

**Fig. 3.**
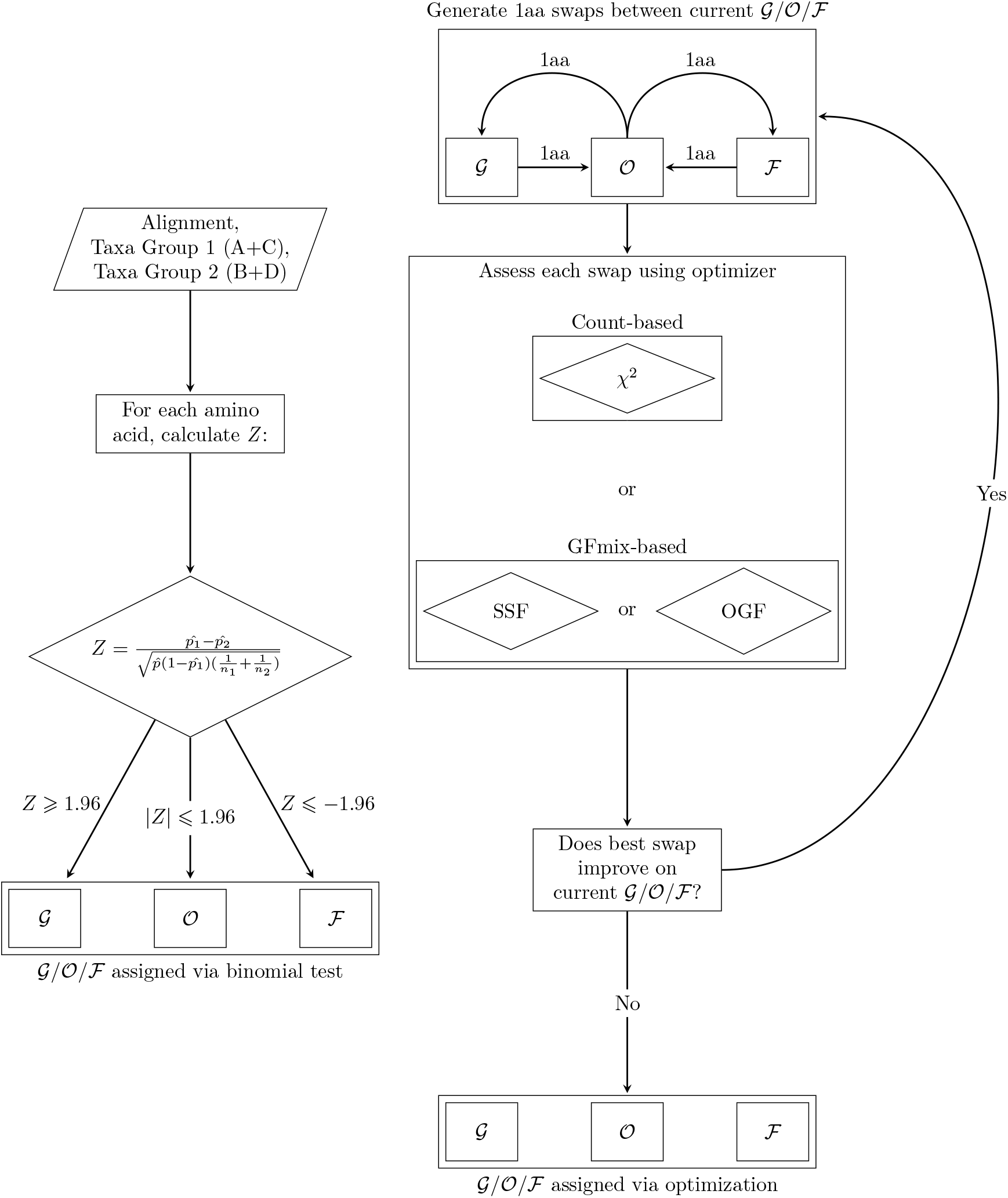
Workflow of 𝒢 /𝒪 /ℱ assignment from simulated sequence data. **Left:** Assignment using binomial test of two proportions. Given an input alignment, and two groups to which taxa are assigned, tabulate amino acid counts and calculate a *Z* -score for each amino acid across the two groups. Amino acids are then assigned into 𝒢, 𝒪 or ℱ on the basis of their *Z* -score. **Right:** Assignment using non-greedy optimization procedure. All possible 1-amino acid “swaps” are performed between 𝒢 / 𝒪 / ℱ, except any 𝒪 -to- ℱ swaps. Each new 𝒢 / 𝒪 / ℱ swap is assessed using one of three optimization criteria, and the best-performing swap is compared with the current 𝒢 /𝒪 /ℱassignment. If this swap improves the criterion, this swap is now the 𝒢 /𝒪 /ℱ assignment and the optimization procedure is repeated. If this swap does not improve the criterion, the optimization procedure is concluded and the current 𝒢 /𝒪 /ℱ assignment is considered optimized. Refer to **Methods and Materials** for further information.

#### Comparing method accuracy

The binomial test of two proportions is a simple onetime test for the equality of two proportions, in this case the proportion of each amino acid between two groups of taxa (Baker et al. 2024). For our 16- and 4-taxon datasets, this test was performed between the equal-frequency taxa (A* and C*) and the high-FYMINK taxa (B* and D*). Although the method does generally assign GARP to 𝒢 and FYMINK to ℱ it frequently assigns additional amino acids to the two classes which should be assigned 𝒪 to (**Figures 4 and S2**). Across almost all 16-taxon replicates, Leucine and Valine are always assigned to 𝒢 and Serine is always assigned to ℱ (**Figure 4**). For 29 of the 100 4-taxon replicates, the method assigns Tyrosine to 𝒪 instead of ℱ (**Figure S2**).

**Fig. 4.**
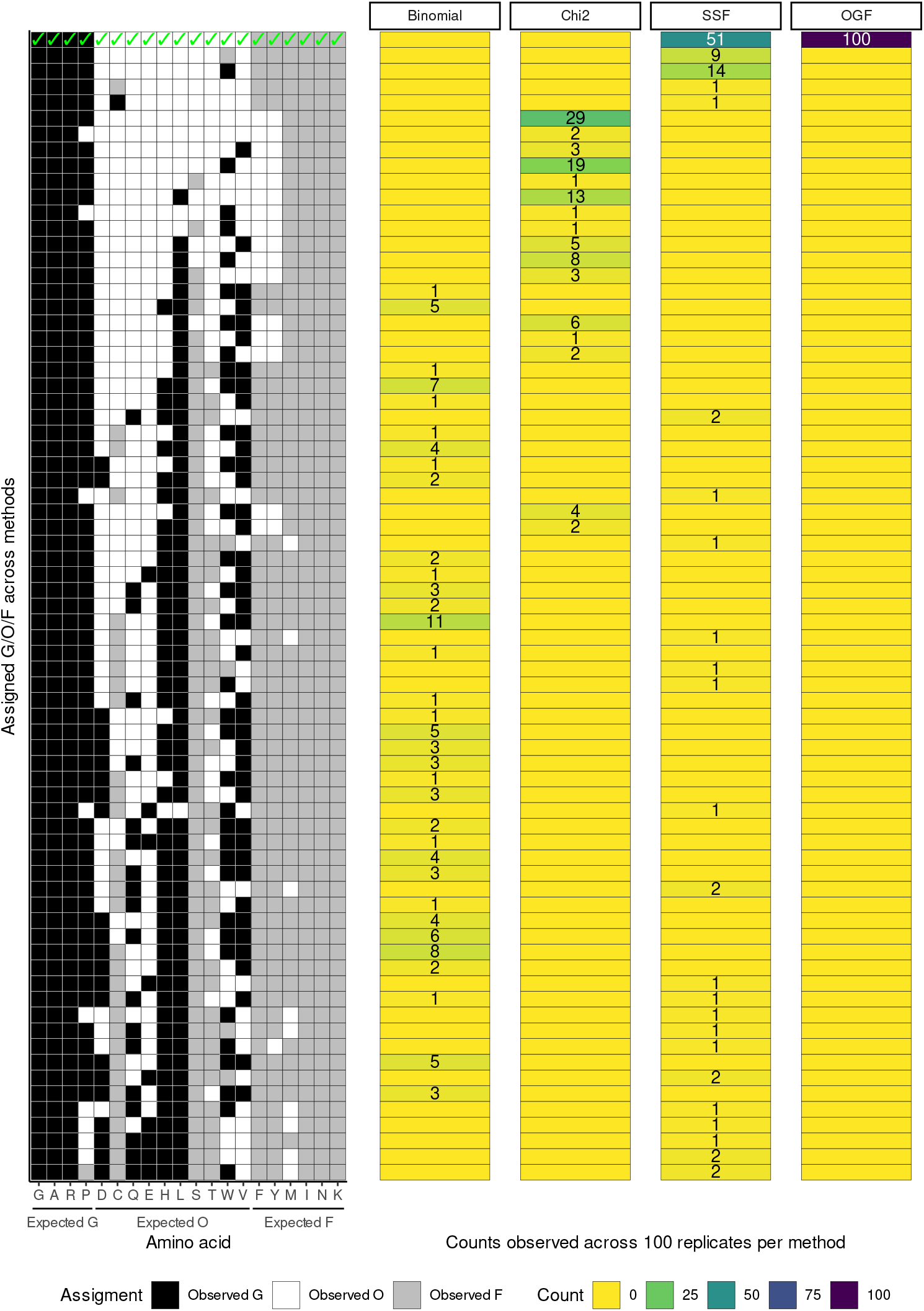
Comparison of 𝒢 /𝒪 /ℱassignment for four compositional heterogeneity identification methods across 100 16-taxon alignments simulated under a LG+C20+Γ+*{*GARP/FYMINK*}* model. **Left:** Assignments of 20 amino acids to𝒢, 𝒪andℱ, with tiles shaded by observed assignment and expected assignment of amino acids underneath the x-axis. Correct assignment - i.e. 𝒢= GARP, 𝒪= DCQEHLSTWV and ℱ= FYMINK - indicated by green ticks. Assignments ordered by from most to least correct. **Right:** Number of assignments by identification method across 100 16-taxon alignments. Blank cells indicate no assignment by that method.

The *χ*^2^ optimization procedure tries to select the 𝒢 and ℱ that minimize the *χ*^2^ statistic for an alignment, using the assignments of the binomial test as a starting point. This method noticeably improves upon the binomial test in terms of closeness to 𝒢 /ℱ = GARP/FYMINK, but still fails to correctly assign 𝒢, 𝒪 and ℱ for any 16-taxon or 4-taxon replicate (**Figures 4 and S2**). In the majority of both sets of replicates, Phenylalanine and Tyrosine are placed in 𝒪 instead of ℱ (**Figures 4 and S2**). In smaller numbers of 16-taxon replicates, Histidine and/or Tryptophan is assigned to 𝒢 (**Figure 4**). Leucine and Serine are assigned to 𝒢 and ℱrespectively in some of the 4-taxon replicates (**Figure S2**).

The final two optimization procedures use two of the GFmix implementations discussed in this study - SSF and OGF meaning that these optimization procedures incorporate model and tree information in their assessment. For SSF optimization, the procedures tries to select 𝒢 and ℱ which minimizes the sum-of-squares difference (SS) between observed and model amino acid frequencies for an alignment given a tree and model. For OGF optimization, the procedure tries to select 𝒢 and ℱ which maximizes the log-likelihood statistic estimated by the original GFmix model for an alignment, given a tree and underlying site-heterogeneous model (Muñoz-Gómez et al. 2022).

Both procedures show substantial improvement in 𝒢, 𝒪 and ℱ assignment over the other two methods (**Figures 4 and S2**). OGF optimization assigns 𝒢/ ℱ = GARP/FYMINK in all 100 16-taxon replicates (**Figure 4**). SSF optimization correctly assigns 𝒢/ ℱ = GARP/FYMINK in 51 of 100 replicates, and makes a single misassignment in another 25 16-taxon replicates (**Figure 4**). For the 4-taxon replicates, OGF makes the correct assignment in 81 out of 100 replicates and SSF makes the correct assignment in 45 out of 100 replicates (**Figure S2**).

In another 36 replicates, SSF misassigns one amino acid (**Figure S2**).

### Applying GFmix to real data

We applied each GFmix implementation to a modified version of a real dataset of protein sequences derived from algal nuclear and nucleomorph genomes (Novak et al. 2024). Nucleomorphs are vestigial nuclei found in membrane-delimited subcellular compartments that also contain plastids in Cryptomonada and Chlorarachniophyceae algae (Ishida et al. 1999; Douglas et al. 2001; Moore and Archibald 2009). These compartments are relics of “secondary symbionts”; that is eukaryotic algal symbionts that took up residence within a different eukaryotic host and became integrated organelles. Nucleomorphs have undergone extensive genome reduction, and are thus AT-rich at the genomic level and FYMINK-rich at the proteomic level (Moore and Archibald 2009). Novak et al. (2024) constructed a dataset of 180 protein sequences derived from 54 genomes from Rhodophyta, Viridiplantae and four nucleomorphs from Cryptomonada. Using this dataset, Novak et al. (2024) placed Cryptomonad nucleomorphs as sister to extremophilic Cyanidiophytina at the base of Rhodophyta.

We modified the Novak et al. (2024) dataset to include homologs from the genome of the *Bigelowiella natans* nucleomorph (Gilson et al. 2006), and downsampled the dataset to 66 genes and 17,696 sites across 16 representative taxa - 9 Rhodophyta, 5 Viridiplantae and the nucleomorph genomes of *Cryptomonas paramecium* and *B. natans* (see **Materials and Methods** for further details). We tested two hypotheses with this dataset - one in which both nucleomorphs branch sister to their closest sampled relative from the literature (Ishida et al. 1999; Douglas et al. 2001), and one in which the two nucleomorphs branch as sister taxa at the base of Rhodophyta as a result of long-branch attraction and shared amino acid composition bias. We designated these hypotheses as “NMs Apart” and “NMs Together”, respectively (**Figure 5**).

**Fig. 5.**
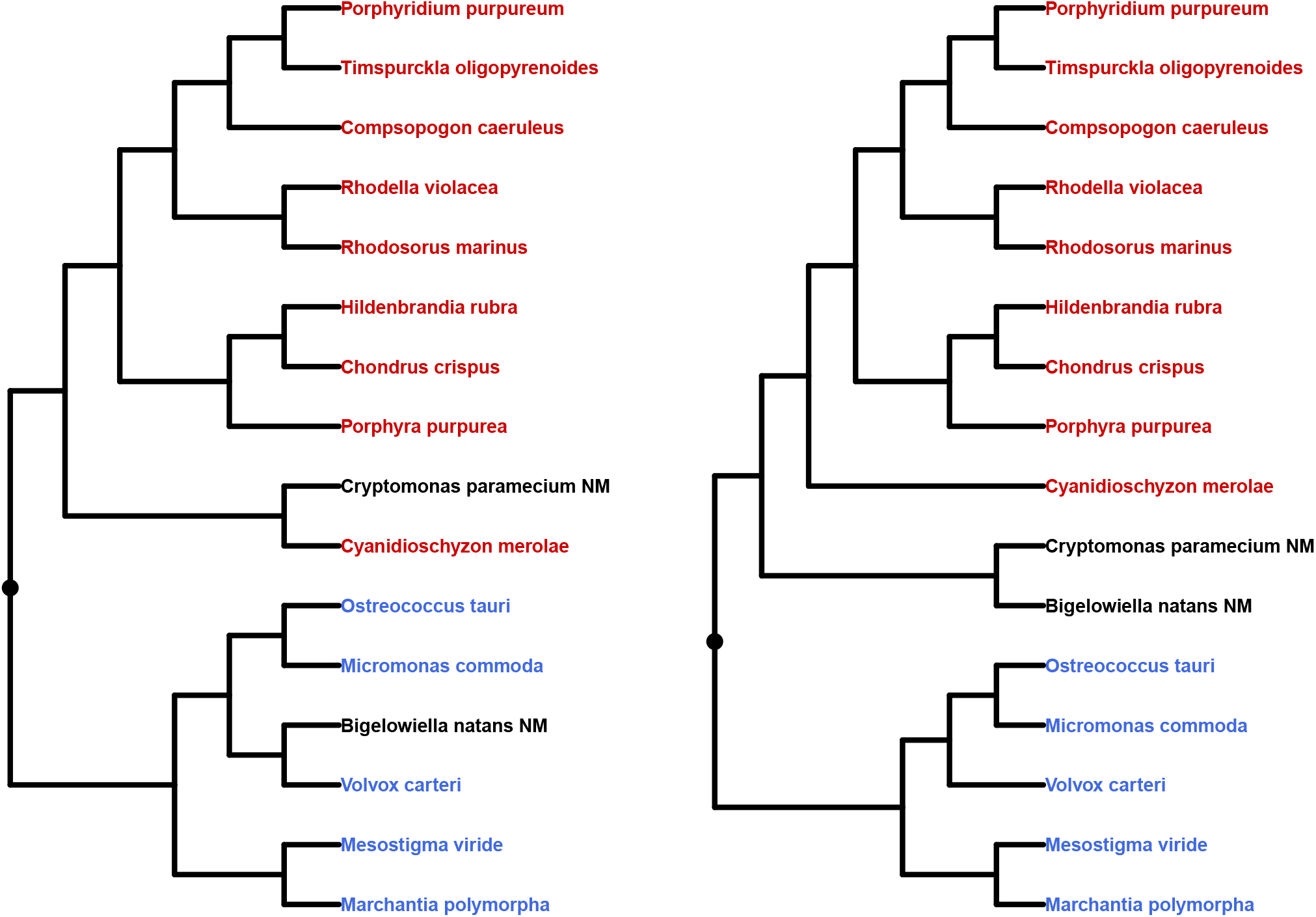
Tree topologies tested for modified Novak et al. (2024) dataset. The modified dataset contains 16 taxa - 9 Rhodophyta (red), 5 Viridiplantae (blue) and two nucleomorph genomes (black). **Left:** “NMs Apart” tree, in which the two nucleomorphs branch sister with their closest sampled relatives from the literature. **Right:** “NMs Together” tree, in which the two nucleomorphs branch as sister taxa at the base of Rhodophyta. “NMs Apart” is considered the “correct” tree on the basis of biological consensus (see references in text), whereas “NMs Together” is artefact of long-branch attraction due to the FYMINK-rich composition of both nucleomorph proteomes. Root position represented by a dot.

We first tested support for both trees under each GFmix implementation with the default 𝒢/𝒪 classes GARP/FYMINK (**Table 6**). We found consistent support for the “NMs Together” hypothesis under both the branch-homogeneous LG+C20+Γmodel and two of the four GFmix implementations tested. The correct “NMs Apart” hypothesis was favoured under the implementation of GFmix — LG+C20+Γ+GFF — that uses full maximum likelihood estimation and under LG+C20+Γ+SSF where 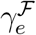 and 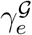 parameters are deter-mined by minimizing a sum of squares. Log-likelihoods improved by at least 1000 units for both hypotheses under each implementation of GFmix relative to LG+C20+Γ(**Table 6**). Calculating the Akaike and Bayesian information criteria for each model shows that LG+C20+Γ+GFF provides the best fit to data for each topology (**Tables S1 and S2**). Despite favouring the correct tree, LG+C20+Γ+SSF exhibits poorer fit to the data than the other GFmix models (**Tables 6**, **S1 and S2**).

**Table 6.** Phylogenetic analysis of two topologies for modified Novak et al. (2024) dataset. **“NMs Apart”**: nucleomorphs branch on either side of Rhodophyta/Viridiplantae split, **“NMs Together”**: nucleomorphs branch together as sister taxa at base of Rhodophyta. See **Fig. 5** for more details. LG+C20+Γfitting was performed using IQ-Tree 3 with mixture weight optimization. All other fittings performed using various implementations of GFmix with default 𝒢/ℱ (GARP/FYMINK) or 𝒢/ℱchosen by OGF optimization procedure.

| Model | $\mathcal{G}/\mathcal{F}$ | NMs Apart | NMs Together | $\Delta\ln(\mathbf{L})$ |
| --- | --- | --- | --- | --- |
| LG+C20+$\Gamma$ | None | -299706.2363 | -299631.303 | -74.9333 |
| LG+C20+$\Gamma$+OGF | GARP/FYMINK | -297920.1909 | -297846.1355 | -74.05535576 |
| LG+C20+$\Gamma$+SSF | GARP/FYMINK | -298319.0172 | -298552.7219 | 233.7046941 |
| LG+C20+$\Gamma$+GFP | GARP/FYMINK | -297173.5316 | -297081.9983 | -91.53331938 |
| LG+C20+$\Gamma$+GFF | GARP/FYMINK | -296945.0122 | -296990.5084 | 45.49624863 |
| LG+C20+$\Gamma$+OGF | GARPVMTHQ/FYCINK | -297276.6916 | -297197.6569 | -79.03475282 |
| LG+C20+$\Gamma$+SSF | GARPVMTHQ/FYCINK | -301322.876 | -300966.5262 | -356.3497707 |
| LG+C20+$\Gamma$+GFP | GARPVMTHQ/FYCINK | -296857.981 | -296783.9782 | -74.00278939 |
| LG+C20+$\Gamma$+GFF | GARPVMTHQ/FYCINK | -296587.1734 | -296624.529 | 37.35556237 |

We repeated this analysis with 𝒢/ℱ classes selected by the optimization procedure previously outlined. We selected OGF as the optimization criterion based on the results of our simulation analysis, and this selected 𝒢/ℱ= GARPVMTHQ/FYCINK. We observed a similar trend to the first analysis - the correct “NMs Apart” hypothesis is only favoured under LG+C20+G+GFF (**Table 6**). This suggests that optimizing the tree and model parame-ters - e.g. *α*, mixture weights and branch lengths - in the context of a branch-heterogeneous model is as important as accounting for branch heterogeneity itself. Log-likelihoods improved for both hypotheses with 𝒢/ℱ classes optimized over the default GARP/FYMINK assignments, with the exception of LG+C20+Γ+SSF which produces worse likelihoods than the base model. AIC and BIC also improved with 𝒢/ℱ classes optimized over the default GARP/FYMINK assignments, with the exception of LG+C20+Γ+SSF which produces worse fit than the base model (**Tables S1 and S2**). This poor performance is likely due to LG+C20+Γ+SSF assigning extreme *b*_*e*_ values to both tip branches and deep branches in both trees (see **Discussion** below for further discussion of this point).

## Discussion

### How well does GFmix perform on simulated data?

We simulated site-heterogeneous alignments with branch-heterogeneous GARP/FYMINK compositions, under the assumptions of the GFmix model, to assess the performance of each GFmix implementation discussed in this study. Given that we know the true simulating values, how well did each implementation perform at estimating branch lengths and node-specific GARP/FYMINK composition? What effect does downsampling from 16 to 4 taxa have on GFmix performance?

Only the most complex GFmix implementation assessed here — GFF — allows for re-estimation of branch lengths. Thus we compared the branch lengths estimated under the LG+C20+Γ+GFF model with those originally estimated by IQ-TREE under the branch-homogeneous LG+C20+Γmodel (or UML for “uncorrected ML” in our notation). Notably, the UML model had large biases in branch length estimates across most branches in heterogeneous trees. This includes underestimating the lengths of branches undergoing a shift from equal-frequency to high-FYMINK composition, and overestimating the lengths of adjacent branches which are more compositionally-homogeneous. This is an error which LG+C20+Γ+GFF largely corrects, although this results in larger estimation variance at branches close to the root.

Assessment of node-specific GARP/FYMINK composition estimation was performed by comparing the expected GARP/FYMINK frequency ratio at each node under the simulating model with those estimated at each node by each GFmix implementation. OGF shows a substantial bias in estimation. SSF shows little bias in estimation but often has the largest standard deviation. Because it reduces the data to tip frequencies prior to estimation, some increase in variance was to be expected relative to ML methods. GFP and GFF perform well across all node types, with larger estimation variance at branches close to the root. These two methods are comparable in their biases and variance of frequency estimation. They do, however, tend to have biases that are larger than SSF. The reasons for this is not clear.

The performance trends we observe for 16-taxon datasets largely hold for 4-taxon datasets. The noteworthy exception is the larger variance in composition estimation at internal nodes for 4-taxon datasets relative to the 16-taxa datasets. This can be explained by the relative lack of information available for parameter estimation for models in 4-taxon datasets relative to 16-taxon datasets.

Overall, LG+C20+Γ+GFF outperforms LG+C20+Γin estimating branch lengths for heterogeneous trees and the two ML-based GFmix implementations (GFP and GFF) give better root mean squared errors of node-specific composition estimation than the non-ML GFmix implementations (OGF and SSF). However, because of their computational costs, these approaches, particularly GFF, may not always be feasible for taxon-rich or site-rich datasets. The “simpler” OGF or SSF implementations may have utility in these cases.

#### How well can we identify compositional heterogeneity?

Although originally implemented to model GARP/FYMINK heterogeneity, the GFmix implementations assessed in this study are capable of modeling user-defined 𝒢/ℱ heterogeneity. For some datasets, the optimal heterogeneity to model may be known *a priori* -e.g. GARP/FYMINK as a result of GC-content variation - but a method of identifying the groups of amino acids most important to heterogeneity is desirable. We devised four methods for identifying compositional heterogeneity directly from sequence data and assessed their performance in identifying 𝒢/ℱ = GARP/FYMINK from simulated site-and-branch-heterogeneous data.

The least computationally-intensive methods — the one-time binomial test of two proportions and 𝒢/ℱclass optimization using the *χ*^2^ test statistic — are also the least accurate in identifying the true heterogeneity from simulated data. The more computationally-intensive, GFmix-based methods —𝒢/ℱclass optimization using either the sum-of-squares difference between observed and model amino acid frequencies estimated by SSF, or calculating the likelihood of the tree under OGF with𝒢/ℱ— can use additional information from the model or tree for 𝒢/ℱ assignment. Perhaps unsuprisingly, these methods perform far better at identifying true heterogeneity from simulated data. Comparing OGF and SSF, OGF had a far larger frequency of correct 𝒢/ℱ identification. It is worth noting, however, that for SSF the most frequent mistake was not correctly assigning a single amino acid, either C or W, both of which are relatively rare in terms of overall frequencies.

The nature of common mistakes in 𝒢/ℱ estimation led us to recognize some unexpected behaviour of the GFmix model. GFmix models sequence evolution along a branch according to a Markov process with stationary amino acid frequencies of mixture classes determined by branch-specific 𝒢, ℱ and rate multipliers associated with these classes. Although the model along a branch is stationary, the marginal frequencies of amino acids at the ancestral node for the branch will not be the same as the stationary frequencies for the branch except in special cases such as when the 𝒢 and ℱ rate multipliers are constant throughout the tree. Consequently, the marginal frequencies change continuously along a branch from what they were at the ancestral node and tend toward the stationary frequencies for the branch as it gets longer. However, since branch lengths are finite, frequencies at the end node of the branch can differ substantially from the stationary frequencies that would be approximated with very large branch lengths. **Figure 6** illustrates how amino acid frequencies averaged over mixture classes behave under the default LG+C20+Γ+GFmix model, as the GARP/FYMINK frequency ratio shifts from those expected under the standard LG+C20+Γmodel to 0.1 times that ratio, as a function of branch length. As expected, the frequencies of GARP decrease as the frequencies of FYMINK increase, with all frequencies stabilizing as branch length increases.

**Fig. 6.**
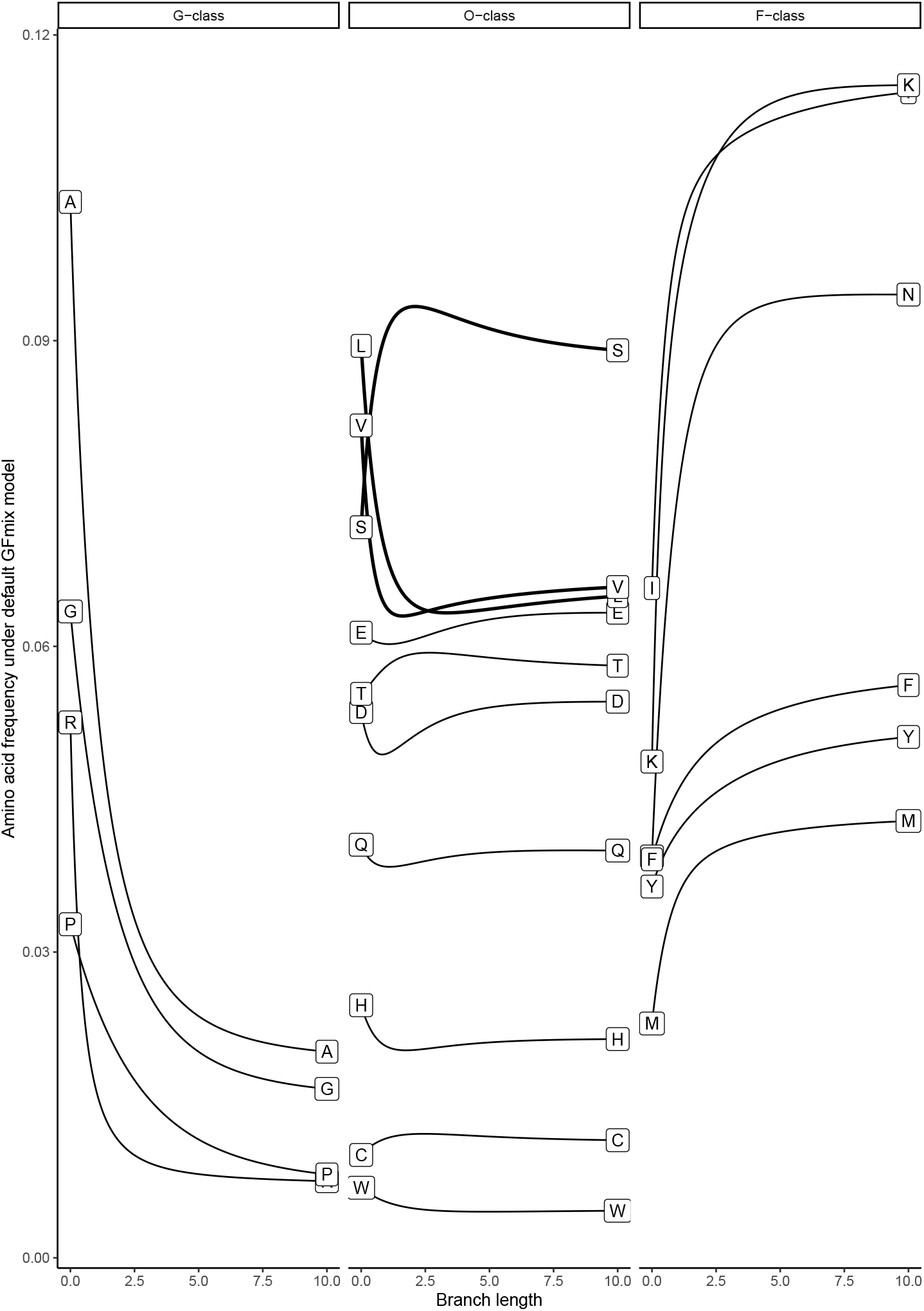
Average amino acid frequencies as a function of branch length under LG+C20+Γ+GARP/FYMINK. As the GARP/FYMINK frequency ratio decreases from those of the LG+C20+Γmodel to 0.1 times that ratio – meaning *π*(*GARP*) *< π*(*FY MINK*) – the frequencies of GARP decrease and the frequencies of FYMINK increase. This effect stabilizes as branch lengths increase. At shorter branch lengths (*<* 2.5), amino acids in the homogeneous 𝒪class also increase or decrease in frequency before returning to equilibrium at longer branch lengths. The exceptions (in **bold**) are Leucine, Serine and Valine. The behaviour of amino acids under the GFmix model at shorter branch lengths may explain misassignments of𝒢, 𝒪and ℱfor some identification methods.

The behaviour of 𝒪 amino acids in **Figure 6** is more surprising. Many 𝒪 amino acids also increase/decrease in frequency at shorter branch lengths, before returning to their original frequency at branch lengths above 2.5. Leucine, Serine and Valine do not follow this behaviour at longer branch lengths: L and V remain at decreased frequencies and S remains at an increased frequency (**Figure 6**). This is mixture-model-dependent behaviour — C20 includes classes in which, across classes, the frequency of Leucine and Valine are positively correlated with the the frequency of some FYMINK amino acids (particularly Phenylalanine and Isoleucine). Similarly, Serine usage is postively correlated with the use of GARP amino acids (particularly Alanine). Thus, when the frequencies of FYMINK amino acids in these classes are shifted this causes an inverse shift of L and V to compensate. Similarly, decreasing GARP causes an increase in S. This behaviour likely explains the consistent misassignment of L, S and V under more simplistic class membership optimization criteria like the binomial approach that chooses 𝒢/ℱ based on differences in observed amino acids across groups without adjusting for branch-lengths. The behaviour of the other 𝒪 amino acids, which need not be close to limiting (large branch length) values, particularly at branch lengths similar to those used to simulate heterogeneous data in this study, is also a likely cause of misassignment. A possible explanation is that as GARP amino acids shift towards increasing usage of FYMINK, these transitions occur via intermediates in the 𝒪 class, leading to temporal increases (and compensatory increases) in some frequencies.

In terms of required runtime, the tree/model-dependent methods of identification are slower than the *χ*^2^ method as is to be expected. For 4-taxon datasets SSF is faster than OGF. This is due to the nature of the sum-of-squares calculation, which is faster for smaller datasets than the standard likelihood calculation. SSF, however, was slower for 16-taxon data sets. Likelihood calculation is slower than sum-of-squares calculation but, for each bin choice, OGF calculates a single likelihood using *b*_*e*_ parameters determined by a hierarchical clustering procedure, SSF chooses *b*_*e*_ parameters to minimize a sum of squares which can require many sum-of-squares and derivative calculations. It is possible that many parameter combinations give similar sum-of-squares, thus leading to flatter surfaces and longer optimization times.

Overall, optimization using likelihood *via* OGF is the best performing method of identifying amino acids belonging to 𝒢 and ℱ classes from sequence data and has acceptable runtime and scaling. The other methods assessed here may have some utility for less heterogeneous datasets or smaller datasets, but should be used with some caveats regarding their performance and/or compared with *a priori* knowledge of compositional heterogeneity for a given dataset if available.

#### How well does GFmix perform on real data?

GFmix is intended for use in phylogenetic analysis where compositional heterogeneity is suspected. Artefacts arising from compositional heterogeneity have been suspected in a number of deep-time phylogenetic analyses (Williams et al. 2021). However, artefacts can occur in any dataset with an appreciably diverse sampling of taxa (Muñoz-Gómez et al. 2019) or particularly divergent protein sequences (Foster and Hickey 1999). We applied each GFmix implementation to a modified version of a dataset comprised of protein sequences derived from algal nuclear and nucleomorph genomes, adding data from the *B. natans* nucleomorph genome and downsampling to 16 representative taxa including the nucleomorphs of *C. paramecium* and *B. natans* (Gilson et al. 2006; Novak et al. 2024). The Novak et al. (2024) dataset was selected as a basis for our real data analysis as nucleomorph genomes are highly-reduced and thus FYMINK-rich at the proteomic level (Moore and Archibald 2009), whereas most of the nuclear genomes retained in the downsampled dataset were comparatively homogeneous with respect to GARP and FYMINK. We tested two hypotheses with this dataset - one in which the two sampled nucelomorphs in this dataset branch with their closest sampled relatives according to the literature (Ishida et al. 1999; Douglas et al. 2001), and one in which the two sampled nucleomorphs branch as a sister taxa at the base of Rhodophyta due to long-branch attraction.

We found that only full ML estimation with GFmix (LG+C20+Γ+GFF) consistently favoured the independent placement of the two nucleomorphs within Rhodophyta and Viridiplantae respectively. This result is consistent with the accepted biological consensus for the origin of nucleomorphs in Cryptomonada and Chlorarachniophyceae (Moore and Archibald 2009). The log-likelihoods evaluated for each tree and the fit to data according to AIC and BIC improve under each GFmix implementation relative to the base LG+C20+Γmixture model, with one exception, and substantially so for LG+C20+Γ+GFP and LG+C20+Γ+GFF. This suggests that directly estimating branch-specific amino acid composition parameters improves log-likelihood evaluation over the indirect estimation as used in the original implementation of GFmix tested here as LG+C20+Γ+OGF (Muñoz-Gómez et al. 2022; Baker et al. 2024). The further improvement of LG+C20+Γ+GFF over LG+C20+Γ+GFP in favouring the “correct” tree is evidence of the utility of re-estimating additional parameters such as branch lengths and mixture weights in the context of a branch-heterogeneous evolutionary process. Finally, we see improved likelihoods and fit to data when using 𝒢 and ℱ classes determined by our optimization procedure over the default GARP/FYMINK assignments. This further supports the validity of inferring compositional heterogeneity from sequence data as we have attempted with our optimization procedure.

The exception to this trend is the unexpected behaviour of the LG+C20+Γ+SSF model. When 𝒢/ℱ = GARP/FYMINK, the model favours the correct tree but when 𝒢/ℱ = GARPVMTHQ/FYCINK the model favours the incorrect tree with a greater likelihood difference than the base model. Furthermore, when = GARPVMTHQ/FYCINK the model also produces worse likelihoods than the base model for both trees. The fit of the model to data is the worst of each GFmix model when 𝒢/ℱ= GARP/FYMINK, and worse than the base model when 𝒢/ℱ= GARPVMTHQ/FYCINK. Examining the *b*_*e*_ parameters estimated by the model (**Supplementary Information, Supplementary Material**), it appears that this may be a result of the model estimating extremely large *b*_*e*_ parameters at the tips and extremely small *b*_*e*_ parameters along internal branches in both trees. The SSF implementation of the GFmix model attempts to minimize the differences between the amino acid frequencies observed across all terminal nodes with the stationary frequencies implied by the base model. As such, it is sensitive to branch lengths estimated under the base model. As many of the internal branches of the tree have small estimated branch lengths, extreme *b*_*e*_ values are sometimes assigned by SSF in order to change frequencies over edges to best fit taxon frequencies. It is possible that penalizing large or small *b*_*e*_ values could lead to better estimation. At present, we would caution that although use of the SSF will give frequencies that better match observed frequencies, it can also lead to poor tree likelihoods.

#### Other considerations and future directions

There are limitations to the GFmix model which apply to all implementations tested here. For example, our software implementation of GFmix does not perform tree searching. A set of candidate trees for likelihood evaluations must be determined by the user using IQ-TREE or other means. Additionally, GFmix assumes rate heterogeneity categories are derived from the Gamma (Γ) distribution (Yang 1994) and thus is not currently compatible with the FreeRate model of rate heterogeneity (Soubrier et al. 2012). Finally, it cannot yet be used in combination with protein evolution models which accommodate other complex features, such as the GHOST model of heterotachy (Crotty et al. 2019) or MAST for non-treelike evolution (Wong et al. 2024).

GFmix was originally implemented in Muñoz-Gómez et al. (2022) to model amino acid composition biases arising from genome-wide GC-content variation amongst species which drives changes in protein sequence composition such as GARP vs. FYMINK alterations. However, the model can also be applied to cases of selection biases in preferred amino acids as shown in Baker et al. (2024). GFmix assumes that whatever amino acid composition bias exists for a given taxa (or group), it applies to the entire protein sequence(s) under consideration. For certain trees of interest, compositional heterogeneity may involve several different sets of amino acids in different parts of the tree due to lineage-specific adaptations. This can be observed across Archaea, where different amino acid groups are associated with lineages that have undergone genome reduction (resulting in GARP/FYMINK changes), adaptation to hypersaline environments (leading to DE/IK changes) or adaptation to extremely hot environments (the proteomic fraction of IVYWREL or ILVWYGERKP amino acids) (Baker et al. 2024; Baker et al. 2025; Eme et al. 2023; Zeldovich et al. 2007). For such trees of interest, identifying the optimal composition bias to model using the methods outlined here may be difficult. Further curation of data, such as subsampling trees or partitioning based on shared amino acid composition or other information may be advisable. Another alternative, as used in Williamson et al. (2025), is to partition sites on the basis of compositional similarity and to apply the GFmix model separately to each partition. Future elaborations of the GFmix approach could investigate the use of different 𝒢 and ℱ amino acid groups for different parts of a tree.

## Conclusions

Accurate inference of deep phylogenies requires models that can accommodate heterogeneous amino acid composition at both the site and branch levels. In this study, we presented new implementations of the site-and-branch-heterogeneous GFmix model. The simpler implementations estimate branch-specific model parameters independently and then use the GFmix model for likelihood inference with these parameters. More complex implementations of the model can estimate some or all of the model and tree parameters in a maximum-likelihood framework. We assessed the performance of these implementations when applied to data simulated under GARP/FYMINK compositional heterogeneity and to real data subject to similar composition biases.

Applied to simulated data, we showed that our newer implementations of GFmix perform substantially better at estimating branch-specific amino acid composition changes, and the most extensive implementation is capable of correcting branch length estimation errors made by branch-homogeneous profile mixture models. We also demonstrate that GFmix can be used to identify the underlying amino acid groups most responsible for compositional heterogeneity from sequence data.

Applied to an algal nuclear and nucleomorph dataset derived from Novak et al. (2024), GFmix improves likelihoods and fit to data over branch-homogeneous models and does so substantially for the most extensive implementations. We find that the most extensive implementation supports the independent origins of the nucleomorphs of *Bigelowiella natans* and *Cryptomonas paramecium*, in Viridiplantae and Rhodophyta respecitvely, over the artefactual long-branch attraction tree placing the two nucleomorphs as sister taxa. This demonstrates the efficacy of the GFmix model in handling known compositional heterogeneity artefacts within real phylogenetic data.

This study demonstrates the utility of modelling both site- and branch-level heterogeneity in phylogenetics. The GFmix model should be considered for use by researchers in cases where compositional heterogeneity may adversely affect phylogenetic tree inference.

## Acknowledgements

This work and C.G.P.M. were supported by the Moore-Simons Project Call on the Origin of the Eukaryotic Cell, Simons Foundation Grant 735923LPI (https://doi.org/10.46714/735923LPI) and by NSERC Discovery Grants awarded to A.J.R. and E.S.. The authors thank Trong Hahn Ly and Minh Bui (Australian National University) for their guidance on simulating site-and-branch-heterogeneous data with AliSim, and Ryo Harada (Dalhousie University) for identifying and adding *Bigelowiella natans* nucleomorph protein sequence data to the algal dataset analyzed in this study. C.G.P.M. thanks Hector Baños (Dalhousie University and California State University, San Bernardino), Kelsey Williamson (Dalhousie University) and Brittany Baker (Université Paris-Saclay and Institut Pasteur) for advice and feedback in other projects which informed this work.

## Author contributions

A.J.R., E.S. and C.G.P.M. designed and led the study. E.S. designed and implemented all models and parameter estimation methods. C.G.P.M. implemented the analysis code and performed all simulation, downsampling, phylogenetics and data analysis. C.G.P.M. drafted the initial manuscript and all authors contributed to editing the paper.

## Supplementary Material

Additional information for generating simulated site-and-branch-heterogeneous sequence data and additional assessments of compositional heterogeneity identification methods is provided in **Supplementary Information**.

Tree parameter estimations for real dataset analysis is provided in **Supplementary Material**.

Simulated and real datasets, and the code used to perform the analysis and generate figures provided in this study is provided in **Supplementary Data** available on Dryad: Link TBD.

