## Supplementary Information for "Modeling site-and-branch-heterogeneity with GFmix"

### Simulating site-and-branch heterogeneous data

Simulated data was generated using a balanced 16-taxa tree containing four 4-taxon clades. Each clade was assigned the label A, B, C and D, and each taxon was assigned the label A1, A2, etc. All tip branches and all clade stem branches were assigned a branch length of 0.5. Internal branches within each clade (e.g. the branch joining A1 and A2) were assigned arbitrarily small values (0.001) as all GFmix implementations assume a fully-bifuracting tree. Both internal branches on each side of the midpoint root were assigned a branch length of 0.25. All branches in the clades B and D were assigned as having a "high FYMINK" amino acid composition, with a target GARP/FYMINK composition ratio of 0.1 under the assumptions of the original GFmix implementation (Muñoz-Gómez et al. 2022). All other branches were assigned an "equal frequency" amino acid composition with a target GARP/FYMINK composition ratio of 1.

LG was chosen as the default substitution matrix for all simulations (Le and Gascuel 2008). The C20 profile mixture model was chosen to generate site-heterogeneous substitution patterns (Si Quang et al. 2008). Rate heterogeneity was simulated using 4 rate categories with values taken from the Gamma ($\Gamma$) distribution with the $\alpha$ shape parameter set to 0.5 (Yang 1994). Rate categories were generated using a method derived from the discrete.gamma() function in the R package Phangorn (Schliep 2010). The target alignment length was 10,000 sites with no gaps.

At the time of writing, simulating sequence data undergoing both site- and branch-specific processes with AliSim requires custom user-defined input files (Ly-Trong et al. 2022). We simulated site- and branch-heterogeneous GARP/FYMINK composition for a full sequence alignment using a double-partitioned approach where sites are simulated on a profile-by-profile basis, and for a given profile sites are simulated on a rate category-by-category basis. This was to ensure that sites simulated by AliSim are consistently assigned to the same rate- and site-profile across changing GARP/FYMINK compositions in a tree, as should be the case when sites are assigned to a mixture profile in real phylogenetic analysis.

For the simulating C20 profile mixture model, the profile weights of the model as defined by IQ-TREE 3 (Wong et al. 2025) were resampled using a random multinomial distribution. For each profile, the resampled weight for that profile determined how many sites of the full alignment were simulated under that profile - e.g. if a profile had a weight of 0.1 that profile was simulated for 1,000 sites. For each profile, the weights of each Gamma rate category were also resampled from a random multinomial distribution from the default weights (0.25 for each category). The resampled weight of a rate category determined how many sites in a profile were simulated under that category's rate - using the previous example if the weights of each category were = (0.4, 0.3, 0.2, 0.1) then the corresponding rates for each category were simulated across 400, 300, 200 and 100 sites respectively.

For each profile in the simulating mixture model, a Nexus-formatted model file and Newick-formatted tree file were generated as input for AliSim. The Nexus file defined the frequencies of that profile when the GARP/FYMINK ratio = 1 and the Gamma rate-category partitioning for that profile. The Newick file defined the tree topology to simulate under, and the frequencies of that profile when GARP/FYMINK = 0.1 (all B and D branches) and GARP/FYMINK = 1 (all other branches). AliSim generated each simulated profile alignment in FASTA format. All profile alignments are then parsed and concatenated into a full alignment using R (R Core Team 2025).

### Results for 4-taxon simulated datasets

#### *GFmix performance*

The following **Figure S1** displays the performance of each GFmix implementation across 100 downsampled 4-taxon datasets. Refer to the main text for more information.

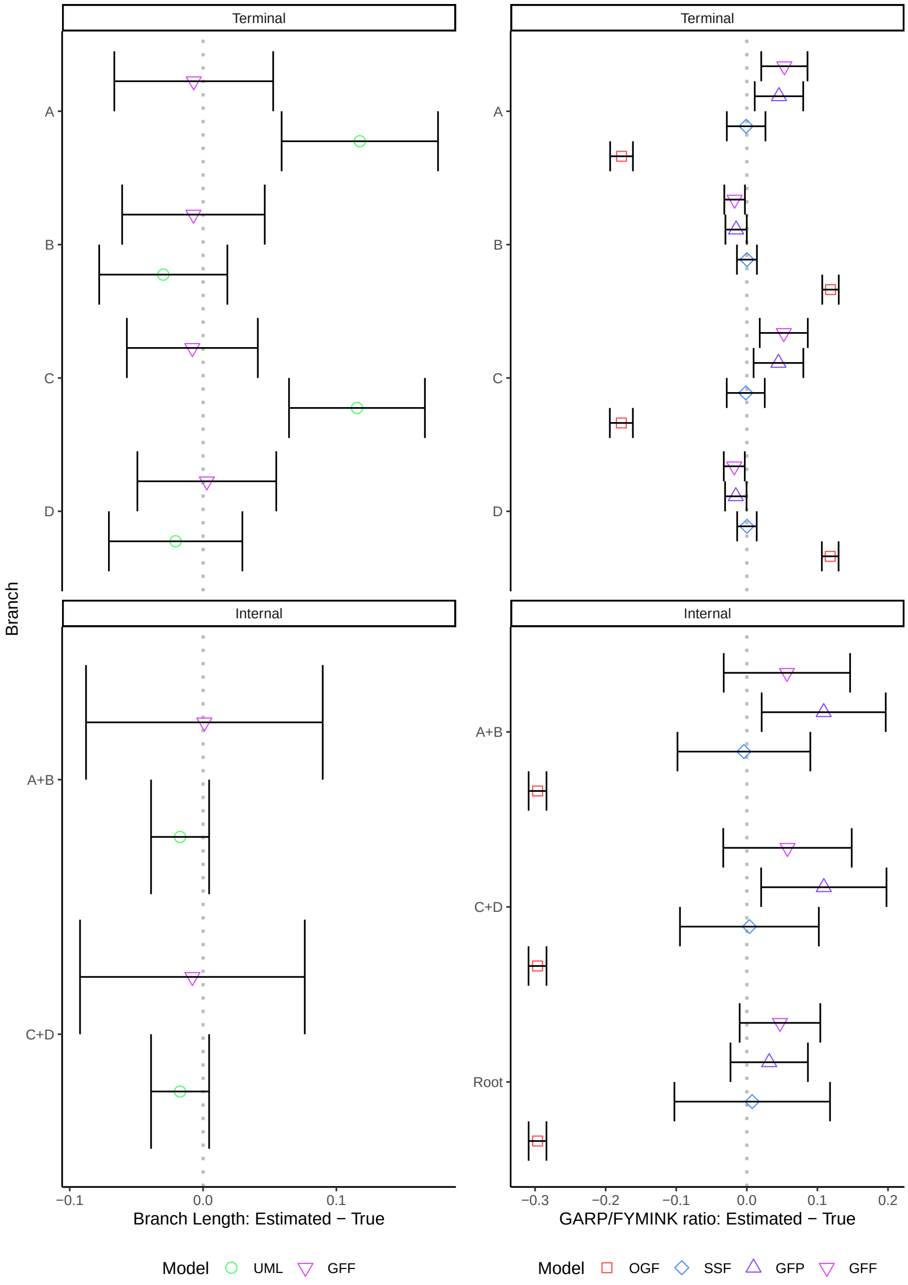


Fig. S1. Performance of GFmix implementations across 100 4-taxon replicates simulated under LG+C20+Γ+{GARP/FYMINK}. Comparisons are between observed estimations and expected estimation based on simulating values. **Left:** Estimation of branch lengths. Comparison made between estimates under LG+C20+Γ model by IQ-TREE and estimates by GFF under LG+C20+Γ+{GARP/FYMINK} model with maximum-likelihood optimization of all model and tree parameters. **Right:** Estimation of node GARP/FYMINK composition ratios. Comparison made between estimates by each GFmix implementation under LG+C20+Γ+{GARP/FYMINK} model. Dots: mean value, errorbars: ±1$\sigma$, dotted line: observed and expected are identical.

#### *Identifying compositional heterogeneity*

The following **Figure S2** displays the performance of each compositional heterogeneity identification method across 100 downsampled 4-taxon datasets. Refer to the main text for more information.

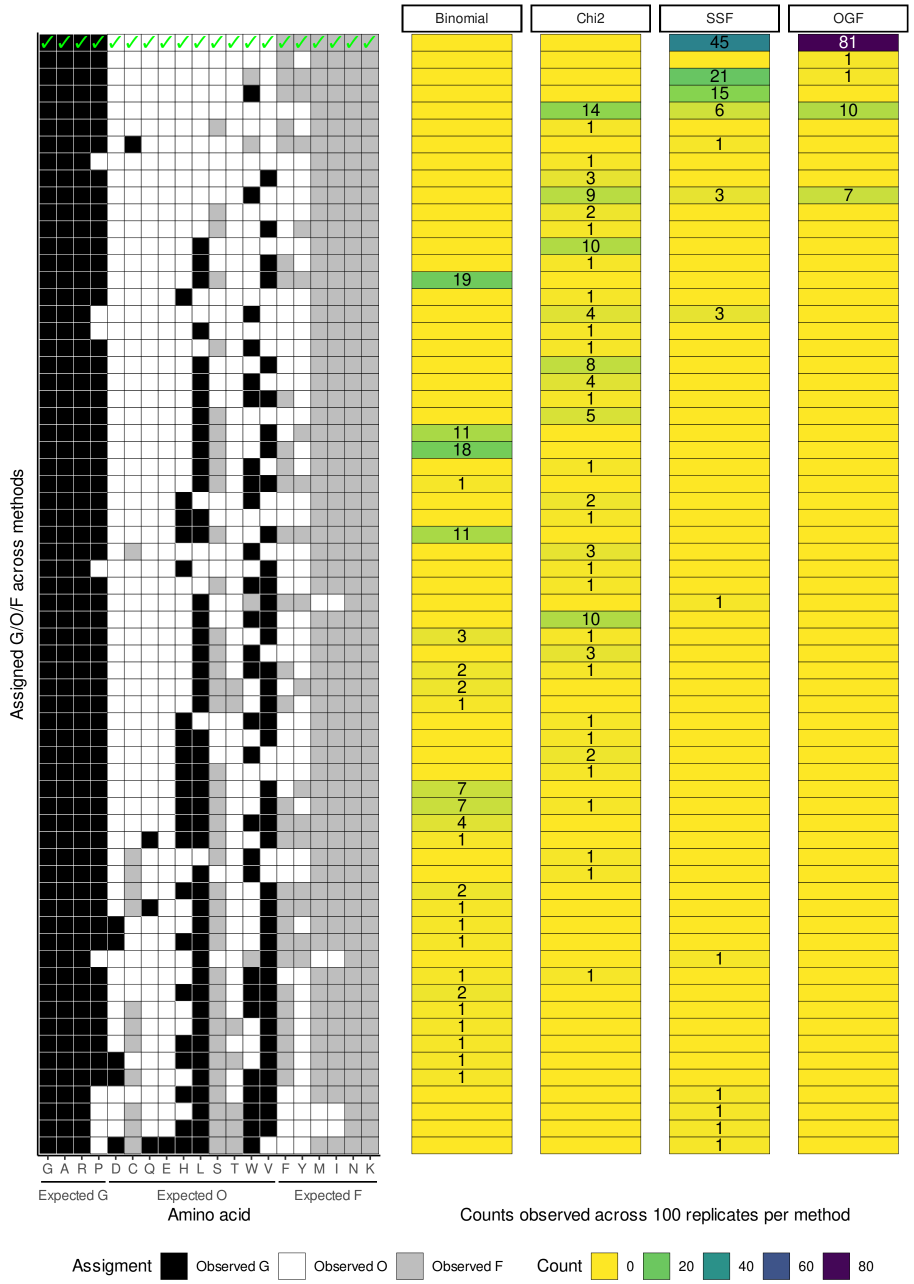


Fig. S2. Comparison of $\mathcal{G}/\mathcal{O}/\mathcal{F}$ assignment for four compositional heterogeneity identification methods across 100 downsampled 4-taxon alignments simulated under a LG+C20+Γ+{GARP/FYMINK} model. **Left:** Assignments of 20 amino acids to $\mathcal{G}$, $\mathcal{O}$ and $\mathcal{F}$, with tiles shaded by observed assignment and expected assignment of amino acids underneath the x-axis. Correct assignment - i.e. $\mathcal{G}$ = GARP, $\mathcal{O}$ = DCQEHLSTWV and $\mathcal{F}$ = FYMINK - indicated by green ticks. Assignments ordered by from most to least correct. **Right:** Number of assignments by identification method across 100 16-taxon alignments. Blank cells indicate no assignment by that method.

### Assessing optimization routine runtimes and iterations

An additional consideration for each identification method was the run time (in seconds) and numbers of optimization routine iterations of each approach. As the binomial test of two proportions is a one-time calculation which is trivial to perform, our comparisons are between the three optimization criteria tested. All optimization analyses for simulated data were performed on Intel Xeon Silver 4510 CPUs using 10 cores for simultaneous $\mathcal{G}/\mathcal{O}/\mathcal{F}$ swap assessment *via* R's parallel library (R Core Team 2025). Across all 16-taxon and 4-taxon replicates, we tabulated the number of iterations for each entire optimization procedure, the run time for each optimization procedure, the run time of the final iteration of the optimization procedure and the run times of each $\mathcal{G}/\mathcal{O}/\mathcal{F}$ swap assessment during the final iteration. Information for each optimization-dependent method, such as the number of iterations a method took to reach final $\mathcal{G}/\mathcal{O}/\mathcal{F}$ assignments and timings of various depths of the optimization procedure was also recorded. This information was visualized using ggplot2 (Wickham 2016) (**Figures S3 and S4**).

In terms of iterations required to make a final $\mathcal{G}/\mathcal{O}/\mathcal{F}$ assignment, all three optimization criteria required on average ∼6-8 iterations for 16-taxon replicates and ∼5-6 iterations for 4-taxon replicates (**Figures S3 and S4**). In terms of runtime, the $\chi^2$ optimization procedure is far faster than the other two optimization procedures due to its relative lack of overhead (e.g. implemented solely within R, lack of additional information requirements such as model or tree) (**Figures S3 and S4**). For the 16-taxon replicates, optimization by likelihood is faster on average than optimization by SSF across each depth of the optimization procedures (**Figure S3**). For the 4-taxon replicates, optimization by SSF is faster on average than optimization by OGF across each depth of the optimization procedures (**Figure S4**).

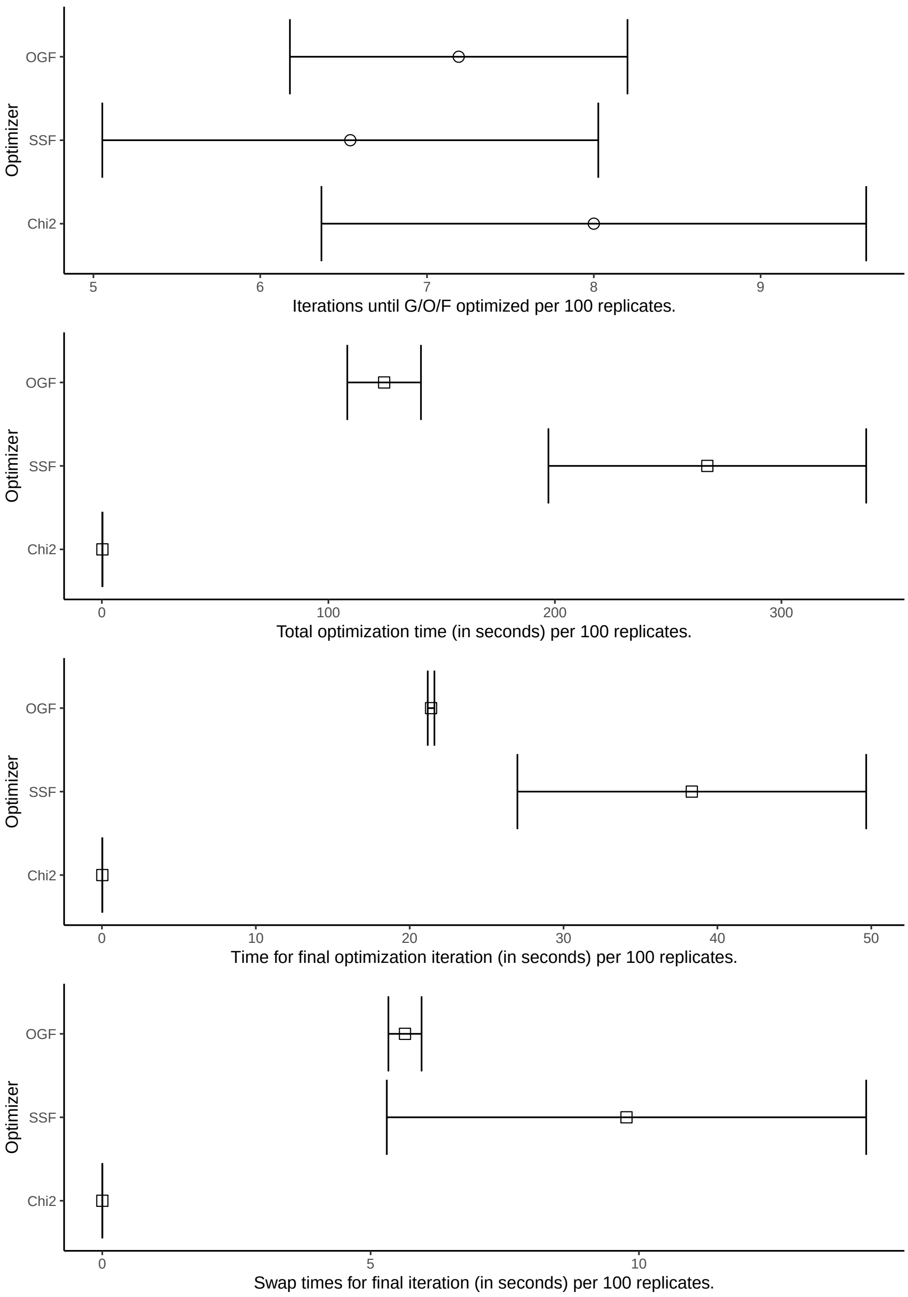


Fig. S3. Comparison of optimization compositional heterogeneity identification methods across 100 16-taxon alignments simulated under a LG+C20+$\Gamma$+{GARP/FYMINK} model. **Top:** Iterations required for method to reach "optimum" $\mathcal{G}/\mathcal{O}/\mathcal{F}$ assignment across 100 replicates. Circle: mean number of iterations, errorbars: $\pm 1\sigma$. **Underneath:** runtime for each method across 100 replicates for full optimization procedure (**Middle top**), the final iteration of the optimization procedure (**Middle bottom**) and assessment of all 1-amino acid swaps during the final iteration of the optimization procedure (**Bottom**). Square: mean runtime, errorbars: $\pm 1\sigma$.

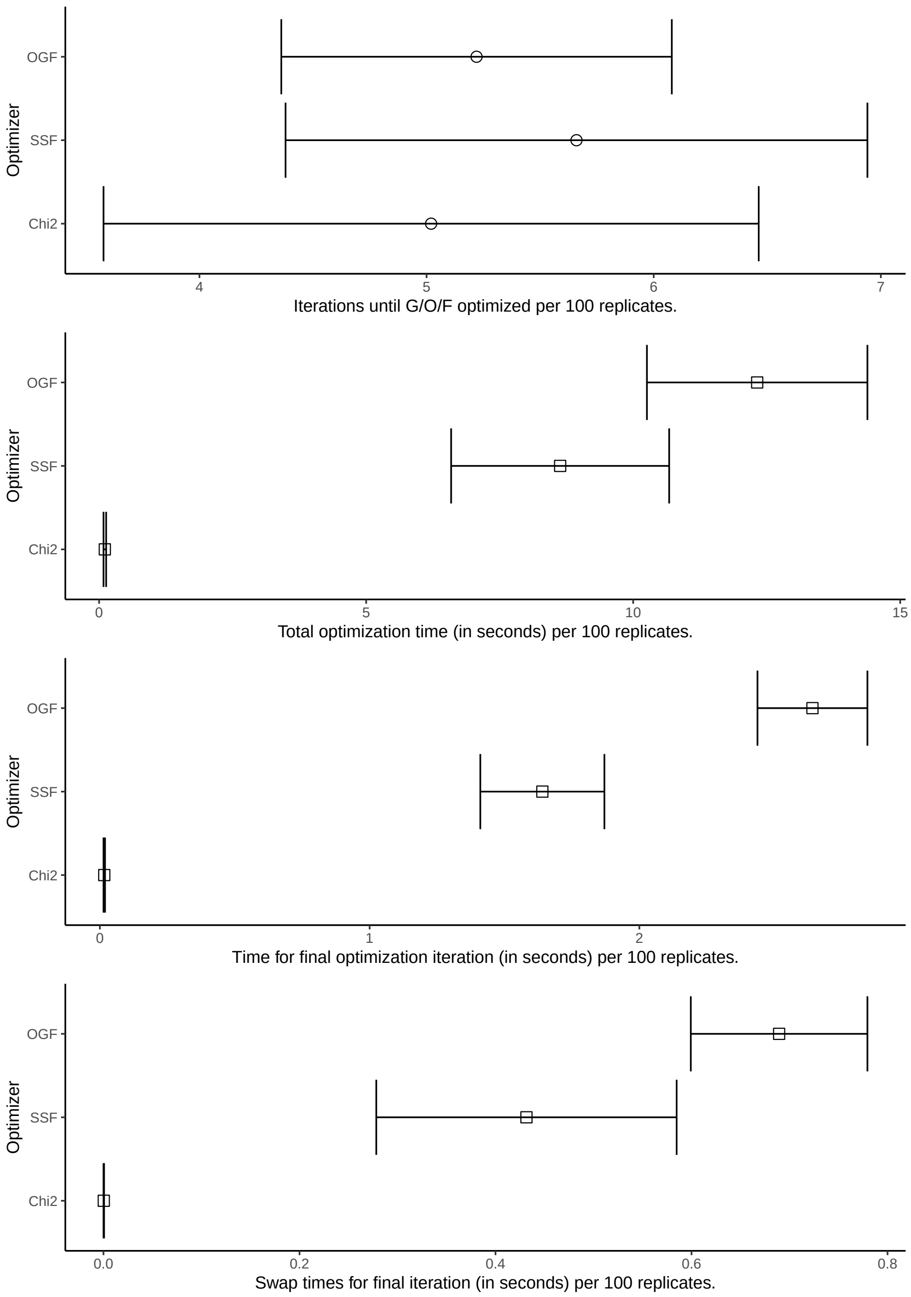


Fig. S4. Comparison of optimization compositional heterogeneity identification methods across 100 4-taxa alignments simulated under a LG+C20+Γ+{GARP/FYMINK} model. **Top:** Iterations required for method to reach "optimum" $\mathcal{G}/\mathcal{O}/\mathcal{F}$ assignment across 100 replicates. Circle: mean number of iterations, errorbars: $\pm 1\sigma$. **Underneath:** runtime for each method across 100 replicates for full optimization procedure (**Middle top**), the final iteration of the optimization procedure (**Middle bottom**) and assessment of all 1-amino acid swaps during the final iteration of the optimization procedure (**Bottom**). Square: mean runtime, errorbars: $\pm 1\sigma$.

### Phylogenetic analysis of modified Novak et al. (2024) dataset

The Akaike information criterion and Bayesian information criterion for each model assessed in **Table 6** in the main text was calculated. Each criterion was calculated using 49 parameters for LG+C20+Γ (29 branches, 19 free weights and $\alpha$), 80 parameters for LG+C20+Γ+OGF and 81 parameters for LG+C20+Γ+SSF (adding an additional parameter for the root branch and 31 branch-compostion parameters) and 112 parameters for LG+C20+Γ+GFP and LG+C20+Γ+GFF (adding an additional parameter for the root branch and 62 branch-composition parameters). Results are presented separately for "NMs Apart" (**Table S1**) and "NMs Together" (**Table S2**).

| **Model** | $\mathcal{G}/\mathcal{F}$ | **Parameters** | **AIC** | **BIC** | **ΔAIC** | **ΔBIC** |
|---|---|---|---|---|---|---|
| LG+C20+Γ | None | 49 | 599510.5 | 599891.7 | 6112.1258 | 5621.9169 |
| LG+C20+Γ+OGF | GARP/FYMINK | 81 | 596002.4 | 596632.7 | 2604.0350 | 2362.8211 |
| LG+C20+Γ+SSF | GARP/FYMINK | 81 | 596800.0 | 597430.3 | 3401.6876 | 3160.4737 |
| LG+C20+Γ+GFP | GARP/FYMINK | 112 | 594571.1 | 595442.5 | 1172.7164 | 1172.7164 |
| LG+C20+Γ+GFF | GARP/FYMINK | 112 | 594114.0 | 594985.5 | 715.6776 | 715.6776 |
| LG+C20+Γ+OGF | GARPVMTHQ/FYCINK | 81 | 594715.4 | 595345.7 | 1317.0364 | 1075.8225 |
| LG+C20+Γ+SSF | GARPVMTHQ/FYCINK | 81 | 602807.8 | 603438.0 | 9409.4052 | 9168.1913 |
| LG+C20+Γ+GFP | GARPVMTHQ/FYCINK | 112 | 593940.0 | 594811.4 | 541.6152 | 541.6152 |
| LG+C20+Γ+GFF | GARPVMTHQ/FYCINK | 112 | 593398.3 | 594269.8 | 0.0000 | 0.0000 |

Table S1. Akaike information criterion, Bayesian information criterion, and ΔAIC / ΔBIC for each model fitted to the "NMs Apart" tree for the modified Novak et al. (2024) dataset. ΔAIC: AIC - minimum(AIC), ΔBIC: BIC - minimum(BIC). See main text for likelihoods and more information.

| **Model** | $\mathcal{G}/\mathcal{F}$ | **Parameters** | **AIC** | **BIC** | **ΔAIC** | **ΔBIC** |
|---|---|---|---|---|---|---|
| LG+C20+Γ | None | 49 | 599360.6 | 599741.9 | 5887.5480 | 5397.3391 |
| LG+C20+Γ+OGF | GARP/FYMINK | 81 | 595854.3 | 596484.5 | 2381.2130 | 2139.9991 |
| LG+C20+Γ+SSF | GARP/FYMINK | 81 | 597267.4 | 597897.7 | 3794.3858 | 3553.1719 |
| LG+C20+Γ+GFP | GARP/FYMINK | 112 | 594388.0 | 595259.5 | 914.9386 | 914.9386 |
| LG+C20+Γ+GFF | GARP/FYMINK | 112 | 594205.0 | 595076.5 | 731.9588 | 731.9588 |
| LG+C20+Γ+OGF | GARPVMTHQ/FYCINK | 81 | 594557.3 | 595187.6 | 1084.2558 | 843.0419 |
| LG+C20+Γ+SSF | GARPVMTHQ/FYCINK | 81 | 602095.1 | 602725.3 | 8621.9944 | 8380.7805 |
| LG+C20+Γ+GFP | GARPVMTHQ/FYCINK | 112 | 593792.0 | 594663.4 | 318.8984 | 318.8984 |
| LG+C20+Γ+GFF | GARPVMTHQ/FYCINK | 112 | 593473.1 | 594344.5 | 0.0000 | 0.0000 |

Table S2. Akaike information criterion, Bayesian information criterion and ΔAIC / ΔBIC for each model fitted to the "NMs Together" tree for the modified Novak et al. (2024) dataset. ΔAIC: AIC - minimum(AIC), ΔBIC: BIC - minimum(BIC). See main text for likelihoods and more information.

For both topologies LG+C20+Γ+GFF and LG+C20+Γ+GFP with optimized $\mathcal{G}/\mathcal{F}$ provide the best fit to data under both criteria, followed by LG+C20+Γ+GFF and LG+C20+Γ+GFP with the default $\mathcal{G}/\mathcal{F}$. LG+C20+Γ+SSF has the worst performance out of each GFmix implementation for both topologies, particularly with optimized $\mathcal{G}/\mathcal{F}$.

The section following the bibliography contains all phylogenetic trees evaluated by each GFmix implementation tested in this study. Branches are coloured by branch-specifc composition parameters estimated by the GFmix implementation (either $b_e$ or $\gamma_e^{\mathcal{G}}$ / $\gamma_e^{\mathcal{F}}$). **Figures S5-S12** represent evaluated trees where $\mathcal{G}/\mathcal{F}$ = GARP/FYMINK and **Figures**

**S13-S20** represent evaluated trees where $\mathcal{G}/\mathcal{F}$ = GARPVMTHQ/FYCINK. All tree figures were generated and annotated using ggtree (Xu et al. 2022). Refer to the main text for the log-likelihoods of each tree. Further information is also provided in **Supplementary Material**.

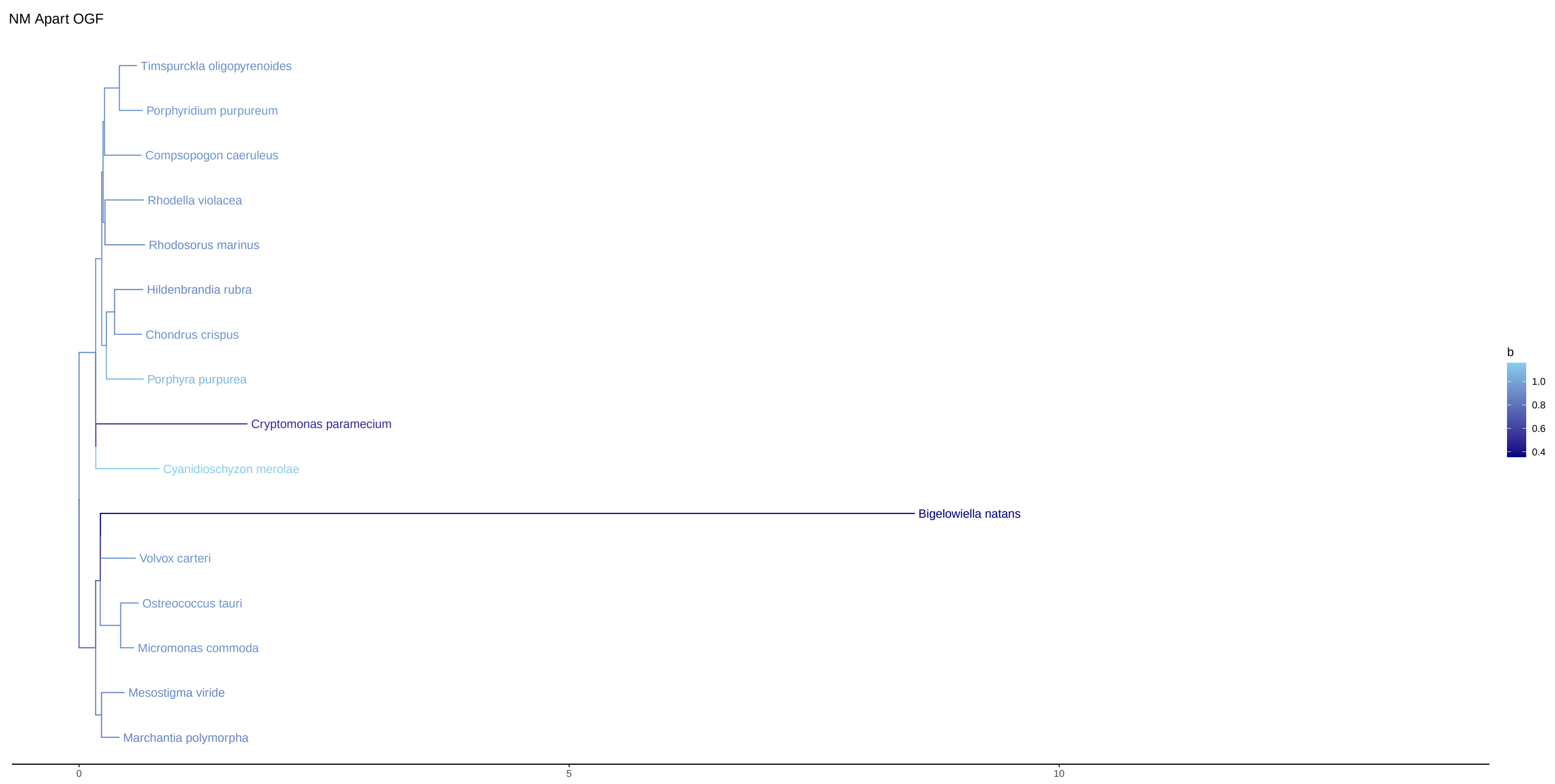


Fig. S5. NM Apart LG+C20+G+OGF, $\mathcal{G}/\mathcal{F}$ = GARP/FYMINK

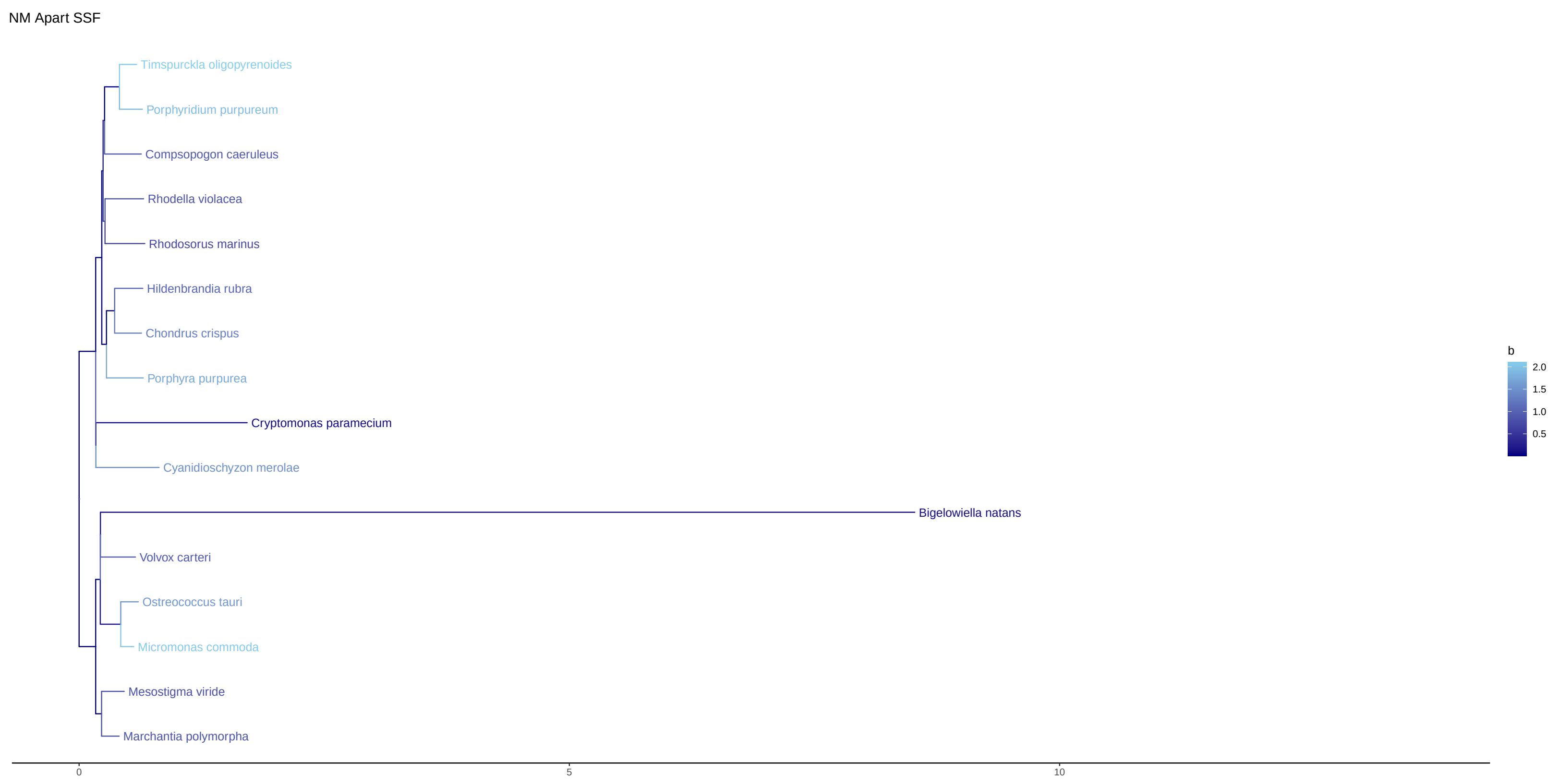


Fig. S6. NM Apart LG+C20+G+SSF, $\mathcal{G}/\mathcal{F}$ = GARP/FYMINK

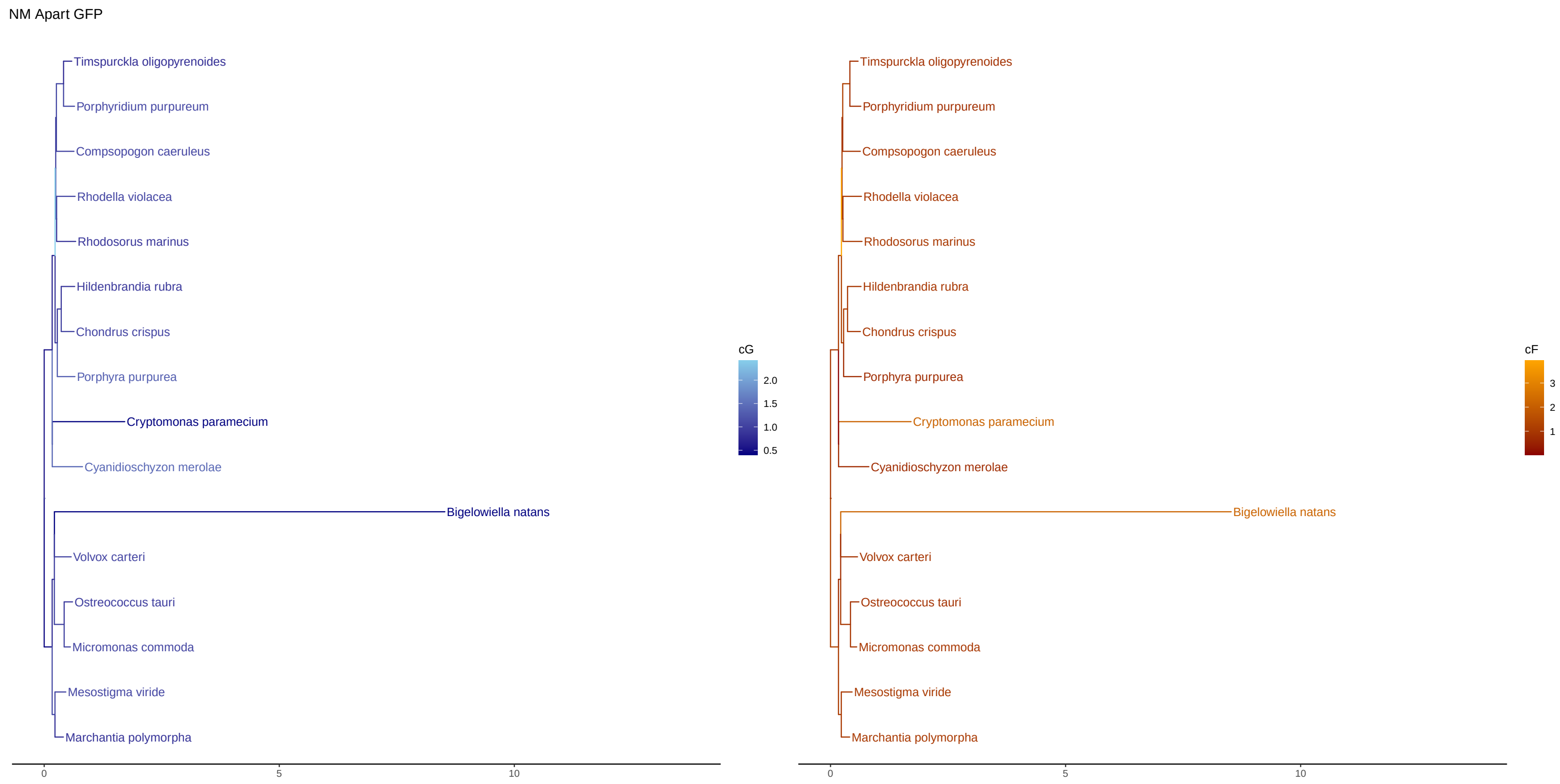


Fig. S7. NM Apart LG+C20+G+GFP $\mathcal{G}/\mathcal{F}$ = GARP/FYMINK

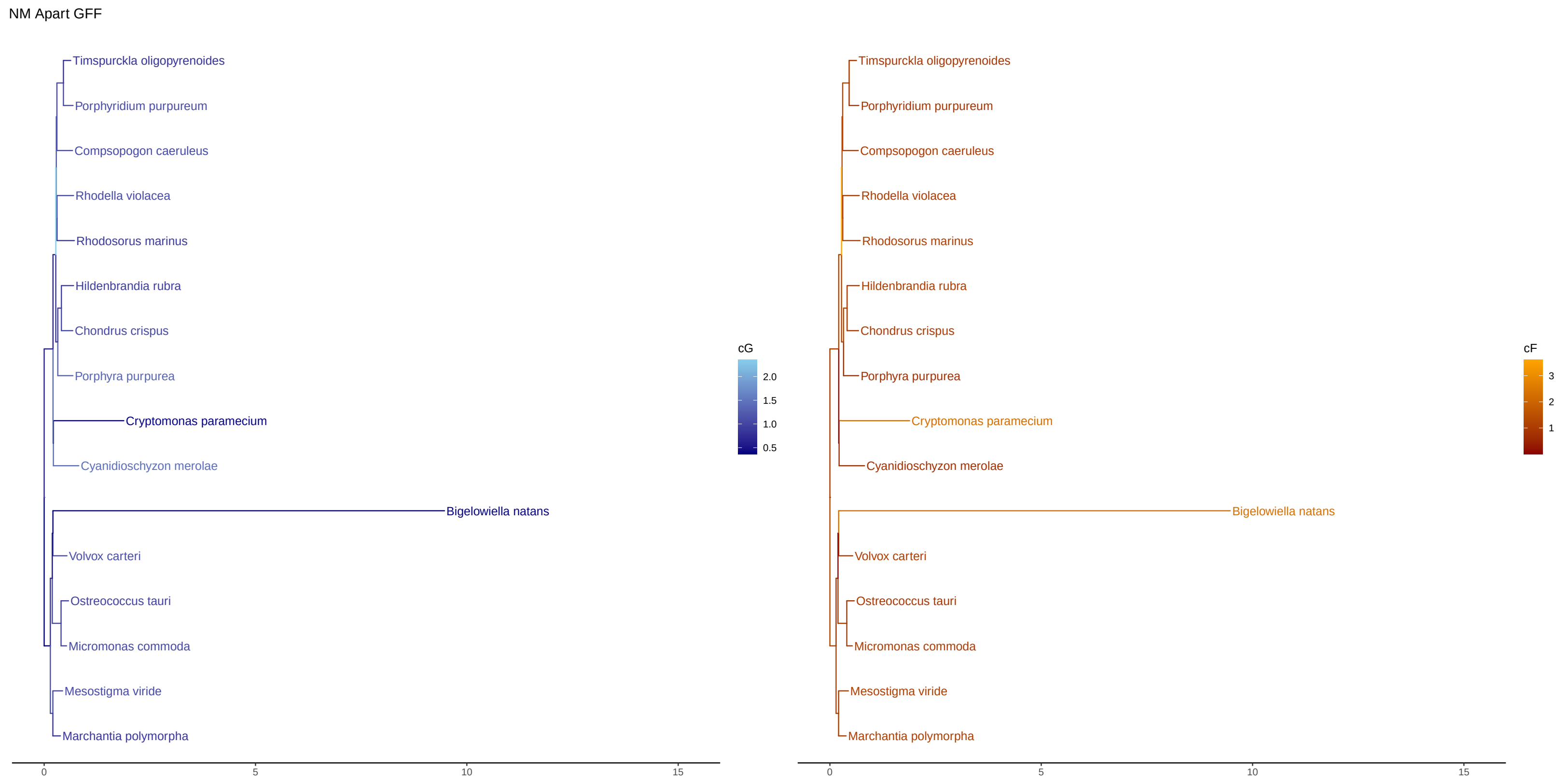


Fig. S8. NM Apart LG+C20+G+GFF, $\mathcal{G}/\mathcal{F}$ = GARP/FYMINK

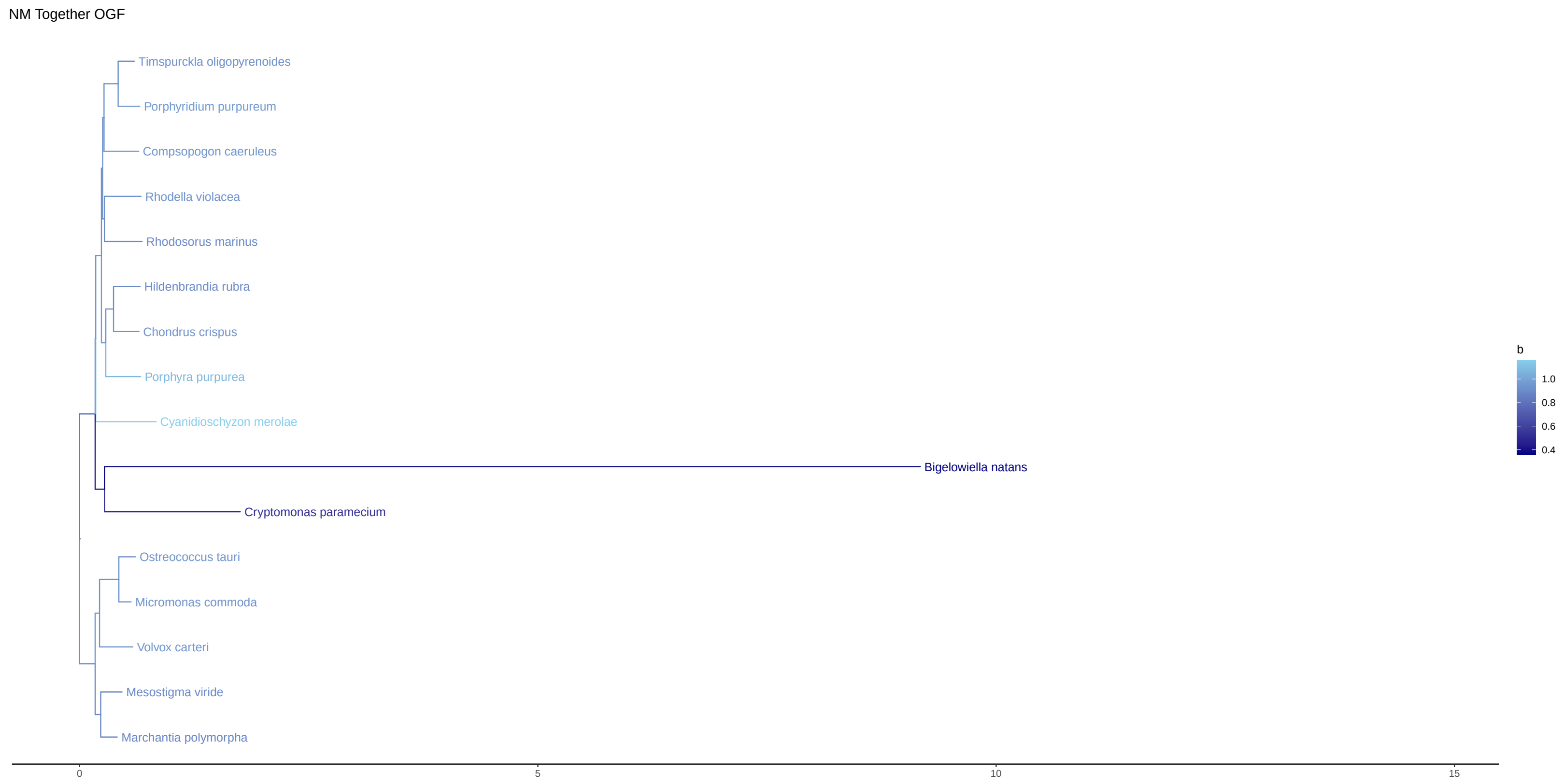


Fig. S9. NM Together LG+C20+G+OGF, $\mathcal{G}/\mathcal{F}$ = GARP/FYMINK

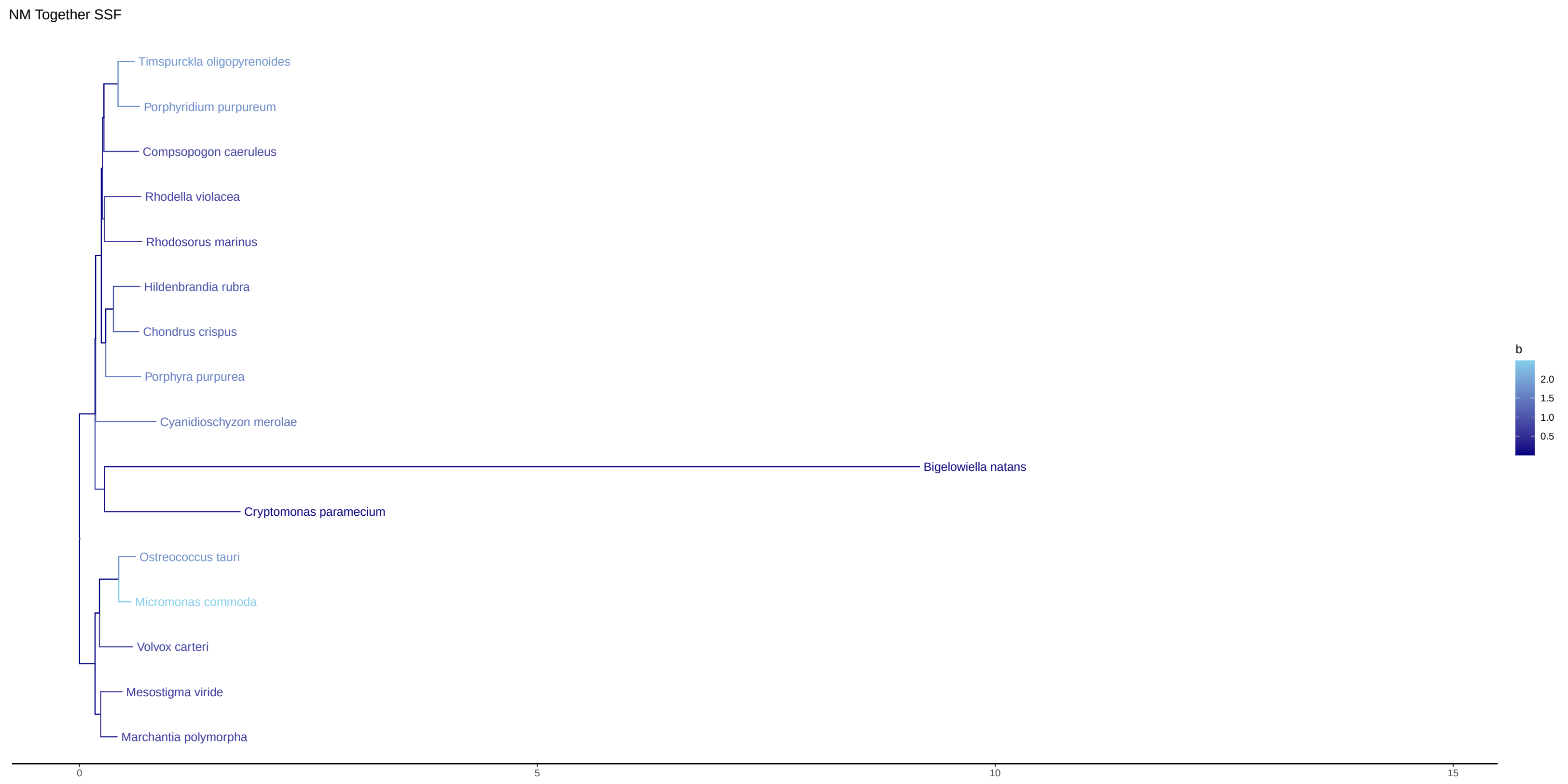


Fig. S10. NM Together LG+C20+G+SSF, $\mathcal{G}/\mathcal{F}$ = GARP/FYMINK

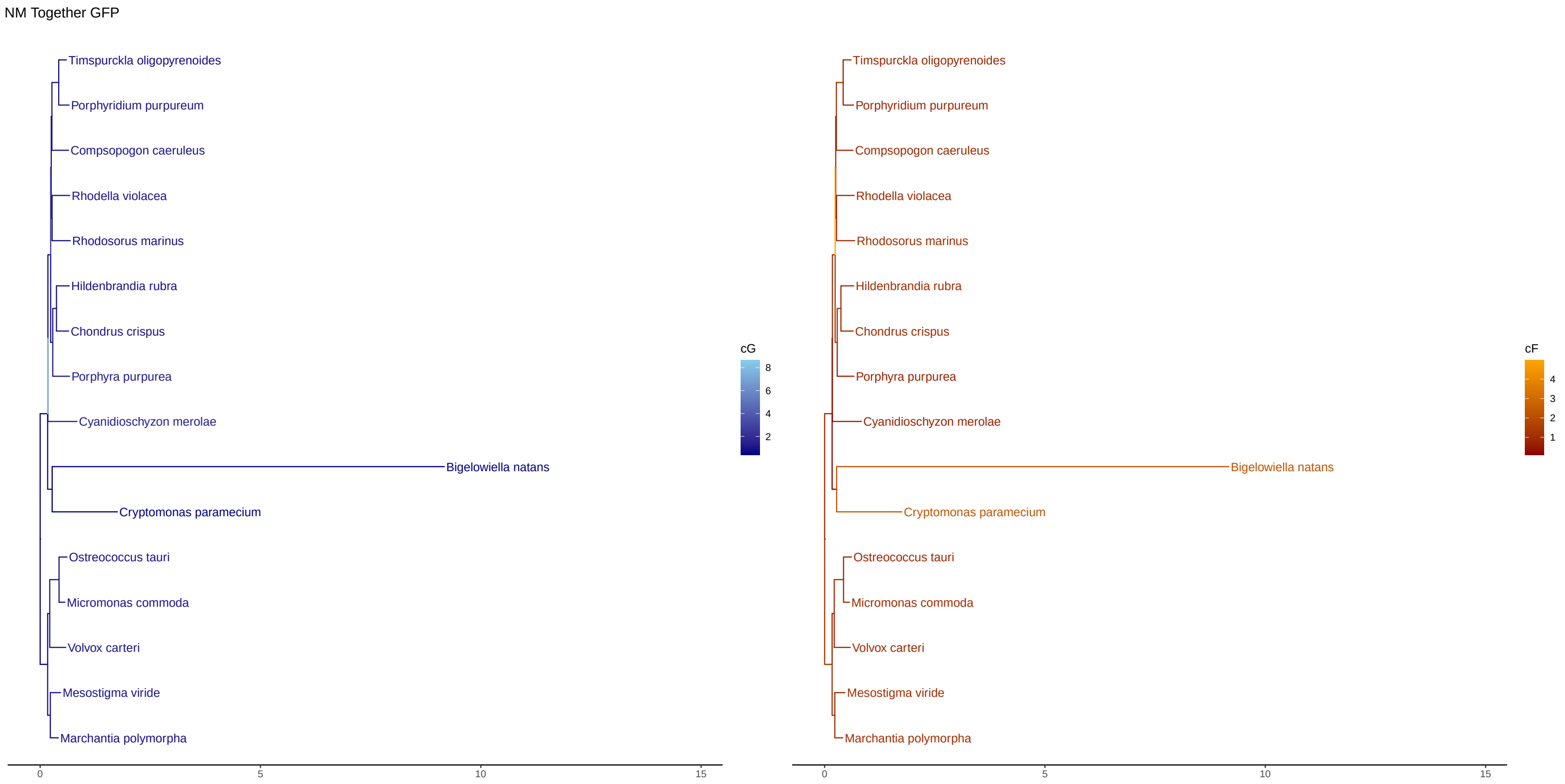


Fig. S11. NM Together LG+C20+G+GFP $\mathcal{G}/\mathcal{F}$ = GARP/FYMINK

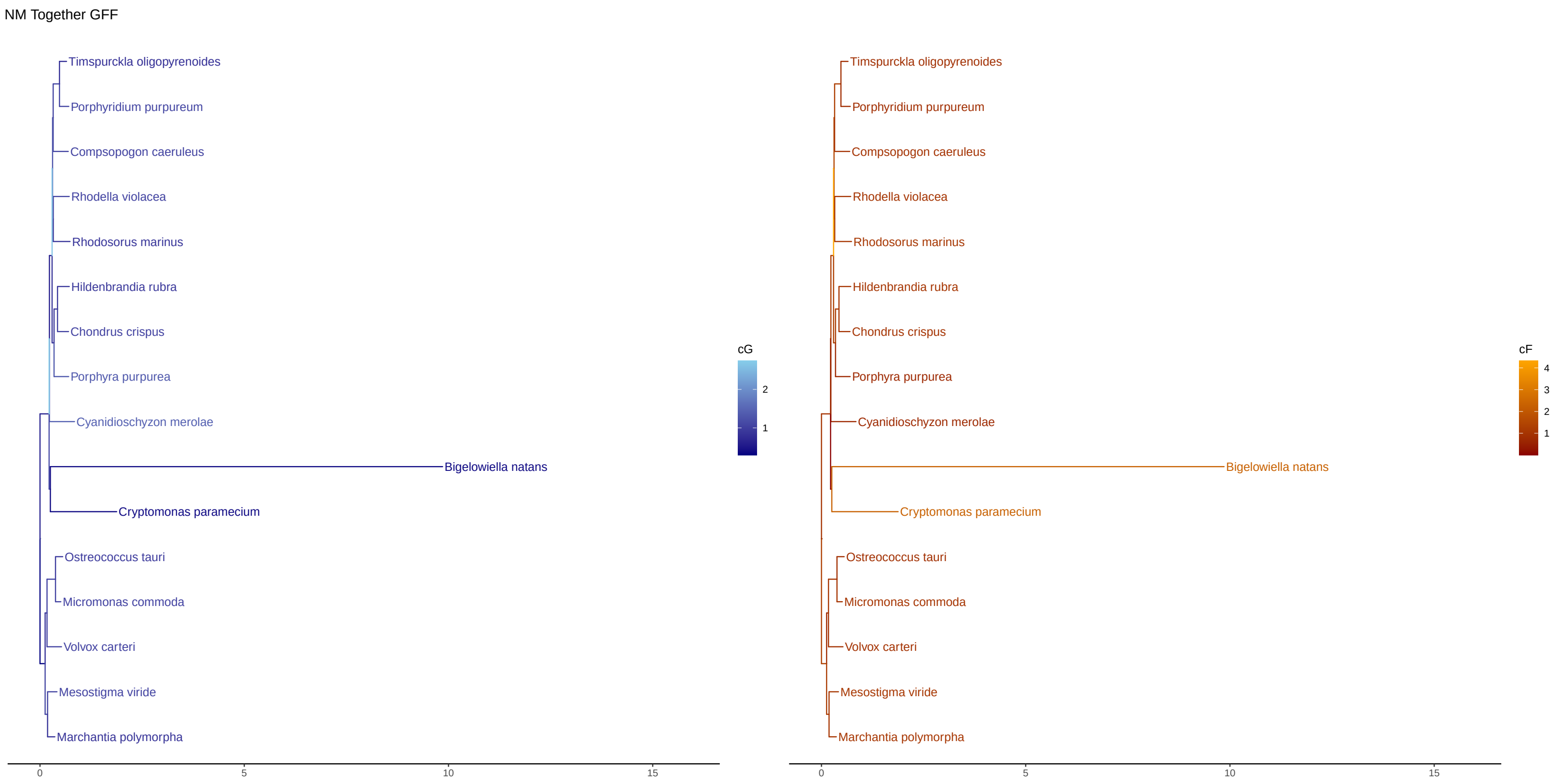


Fig. S12. NM Together LG+C20+G+GFF, $\mathcal{G}/\mathcal{F}$ = GARP/FYMINK

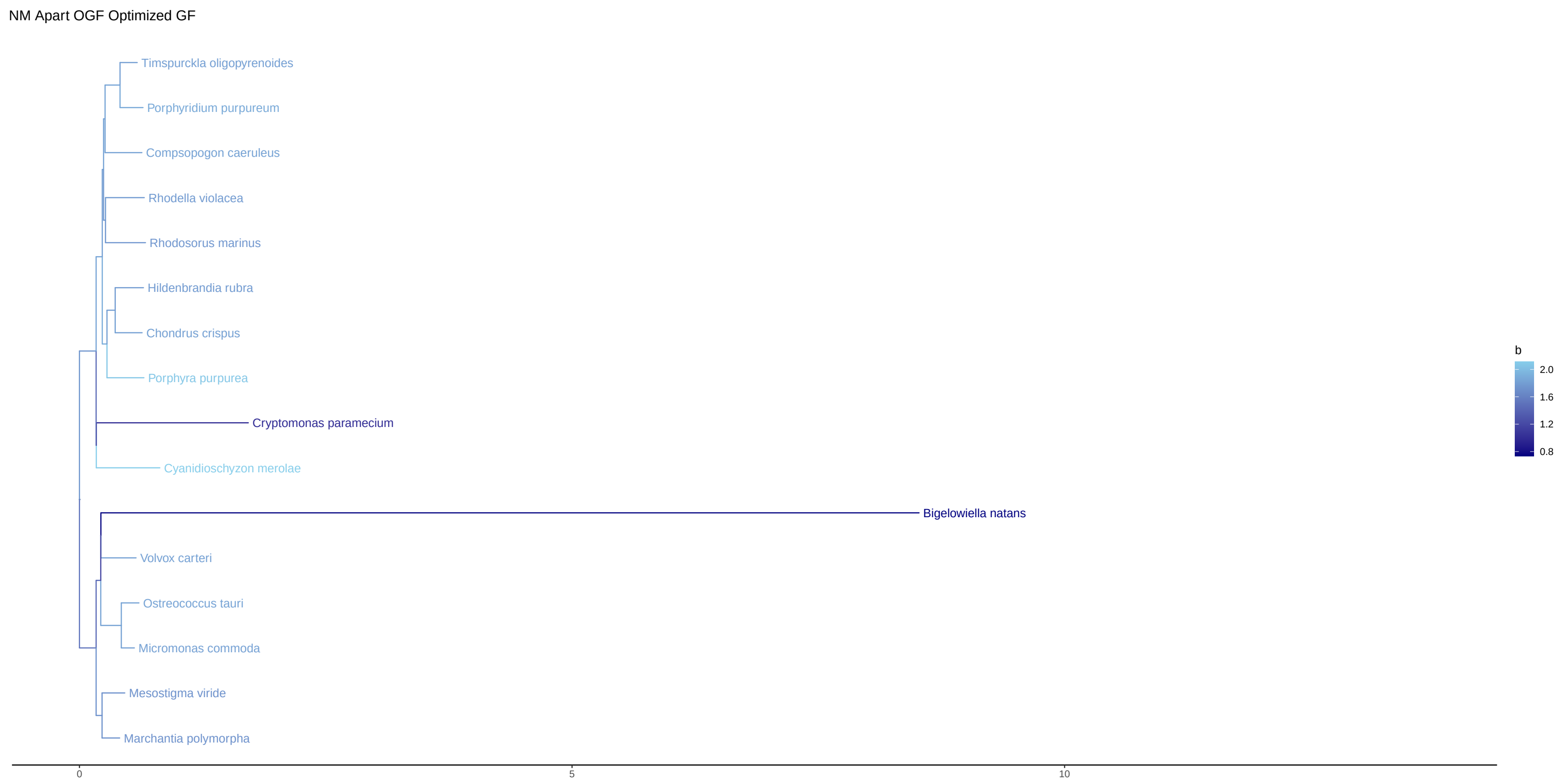


Fig. S13. NM Apart LG+C20+G+OGF, $\mathcal{G}/\mathcal{F}$ = GARPVMTHQ/FYCINK

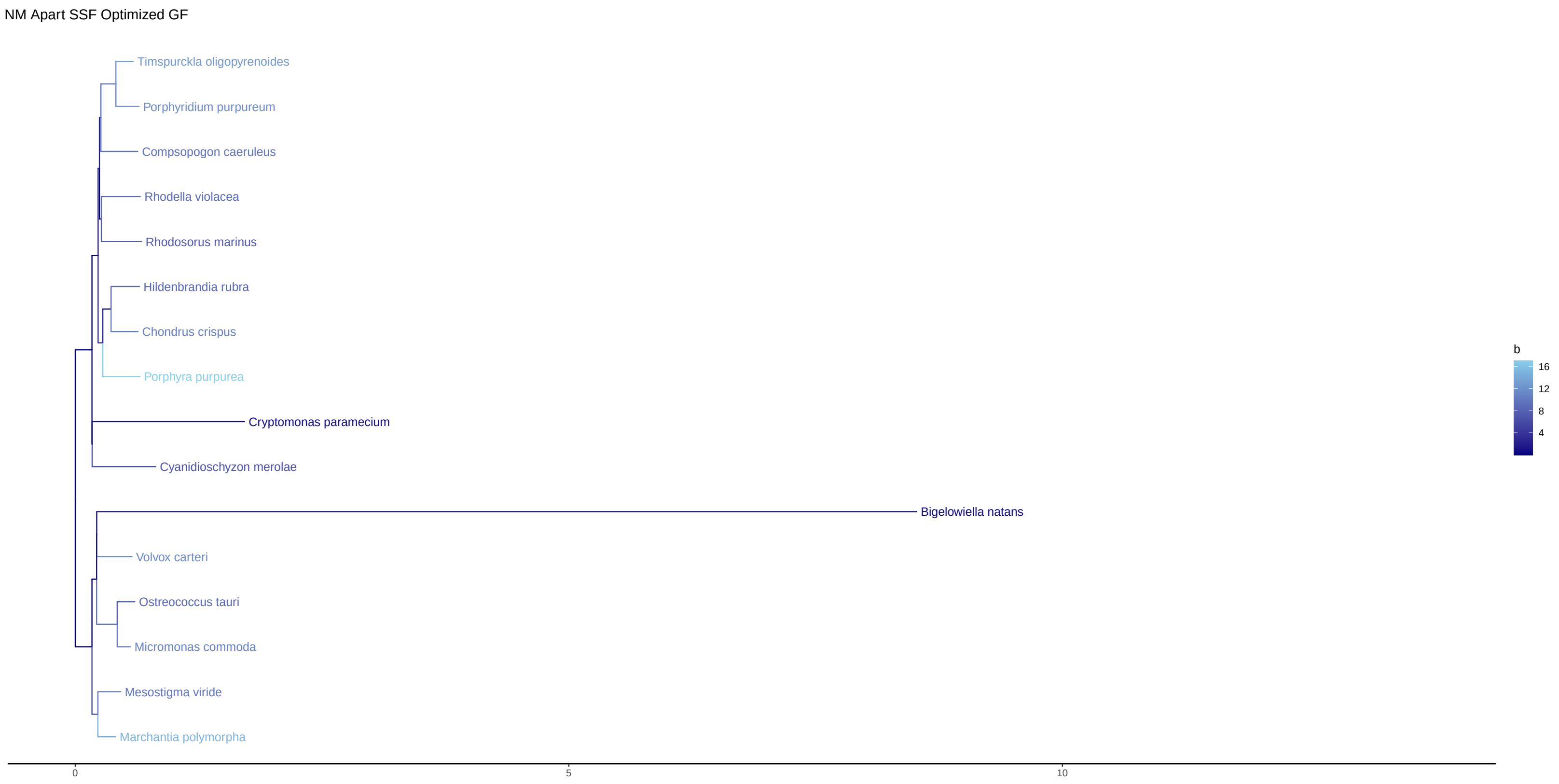


Fig. S14. NM Apart LG+C20+G+SSF, $\mathcal{G}/\mathcal{F}$ = GARPVMTHQ/FYCINK

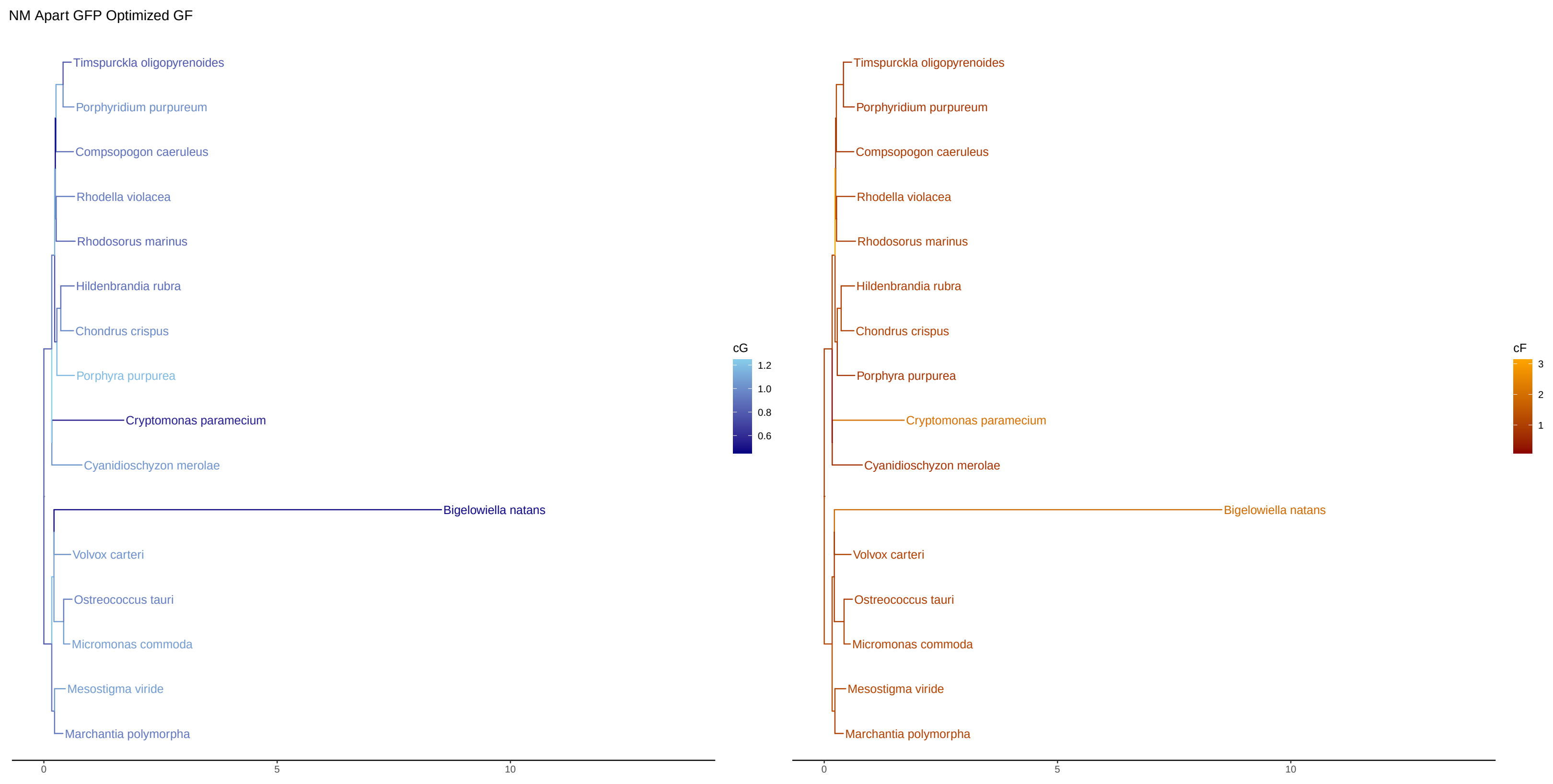


Fig. S15. NM Apart LG+C20+G+GFP $\mathcal{G}/\mathcal{F}$ = GARPVMTHQ/FYCINK

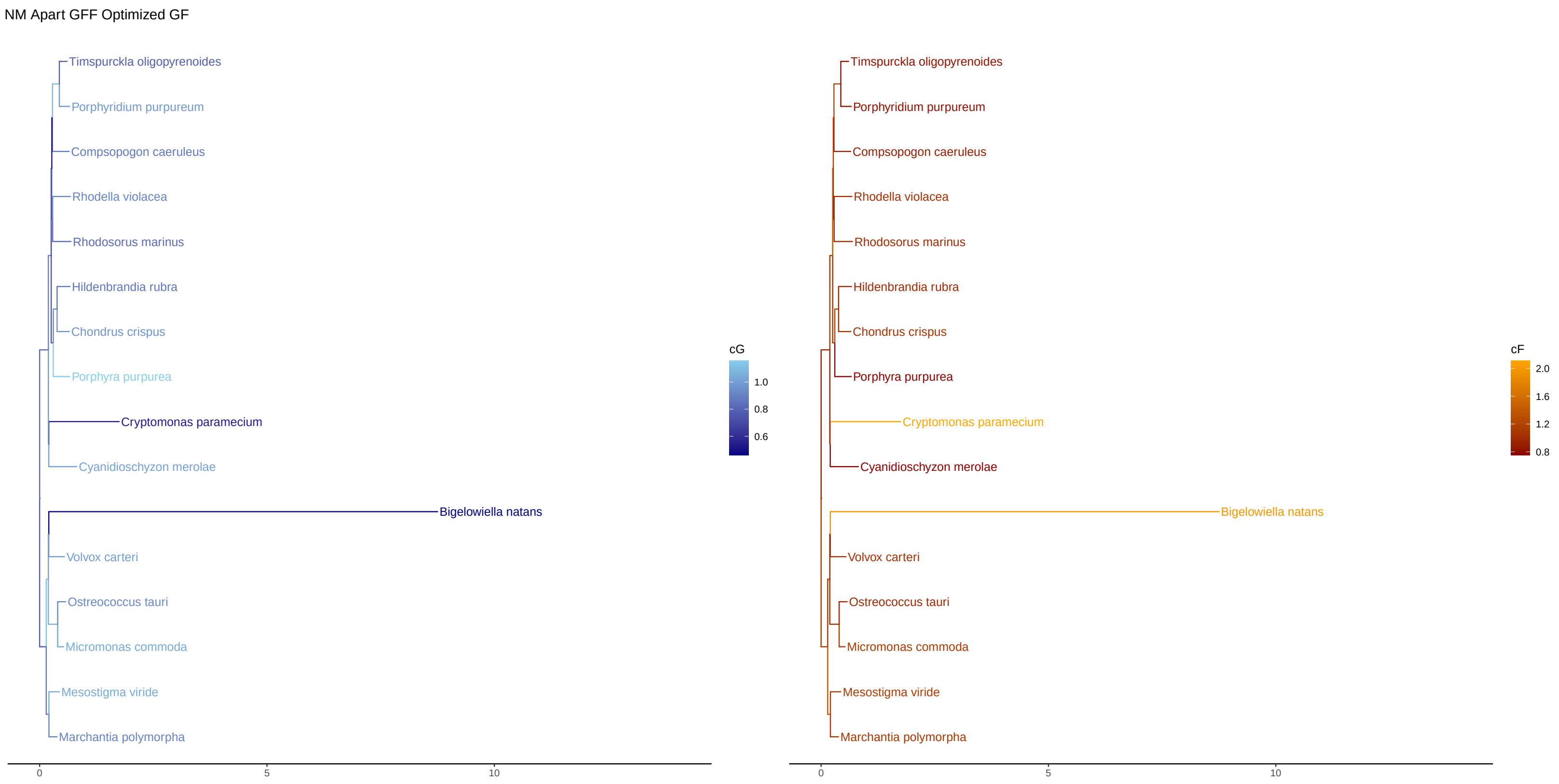


Fig. S16. NM Apart LG+C20+G+GFF, $\mathcal{G}/\mathcal{F}$ = GARPVMTHQ/FYCINK

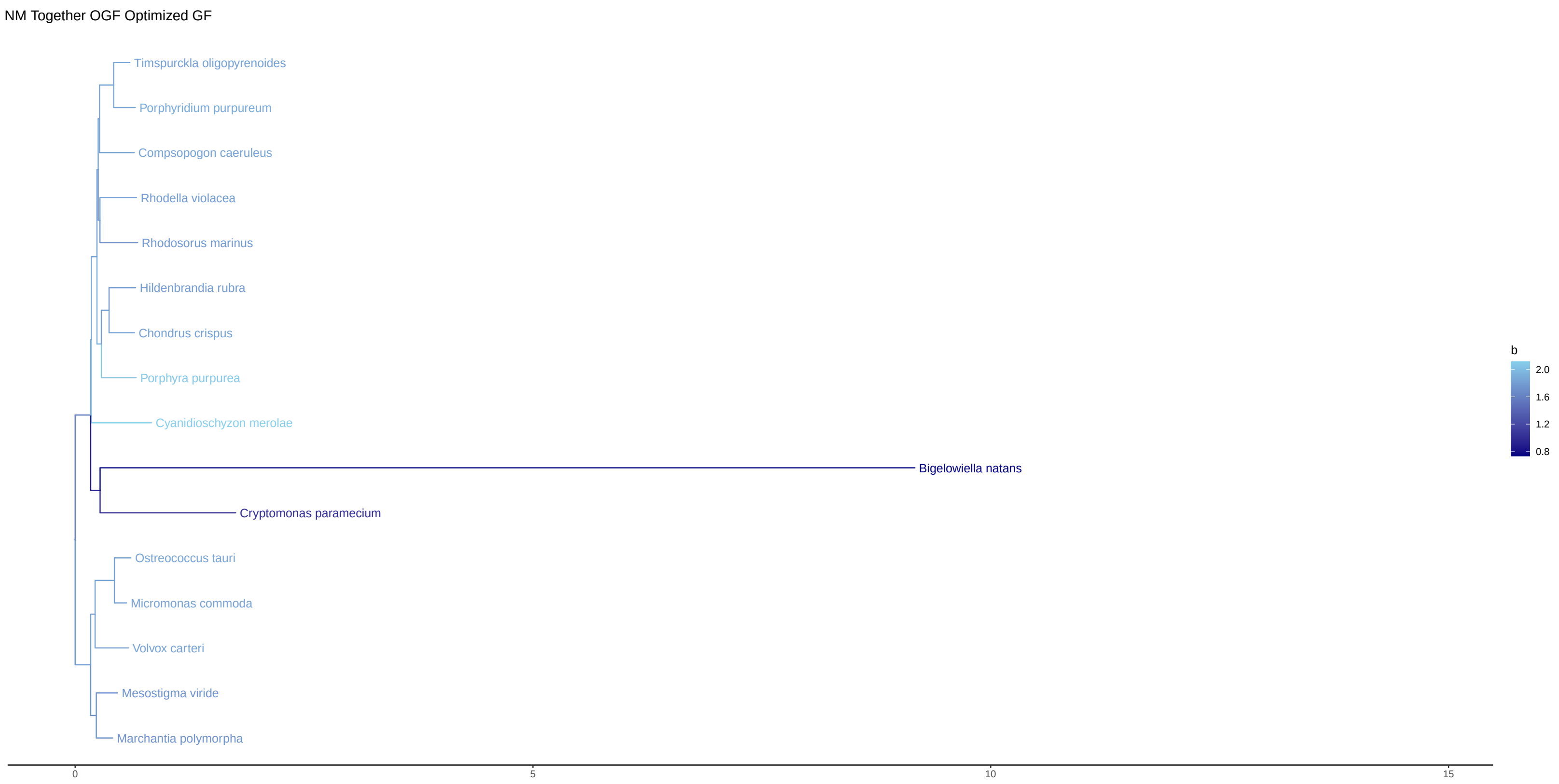


Fig. S17. NM Together LG+C20+G+OGF, $\mathcal{G}/\mathcal{F}$ = GARPVMTHQ/FYCINK

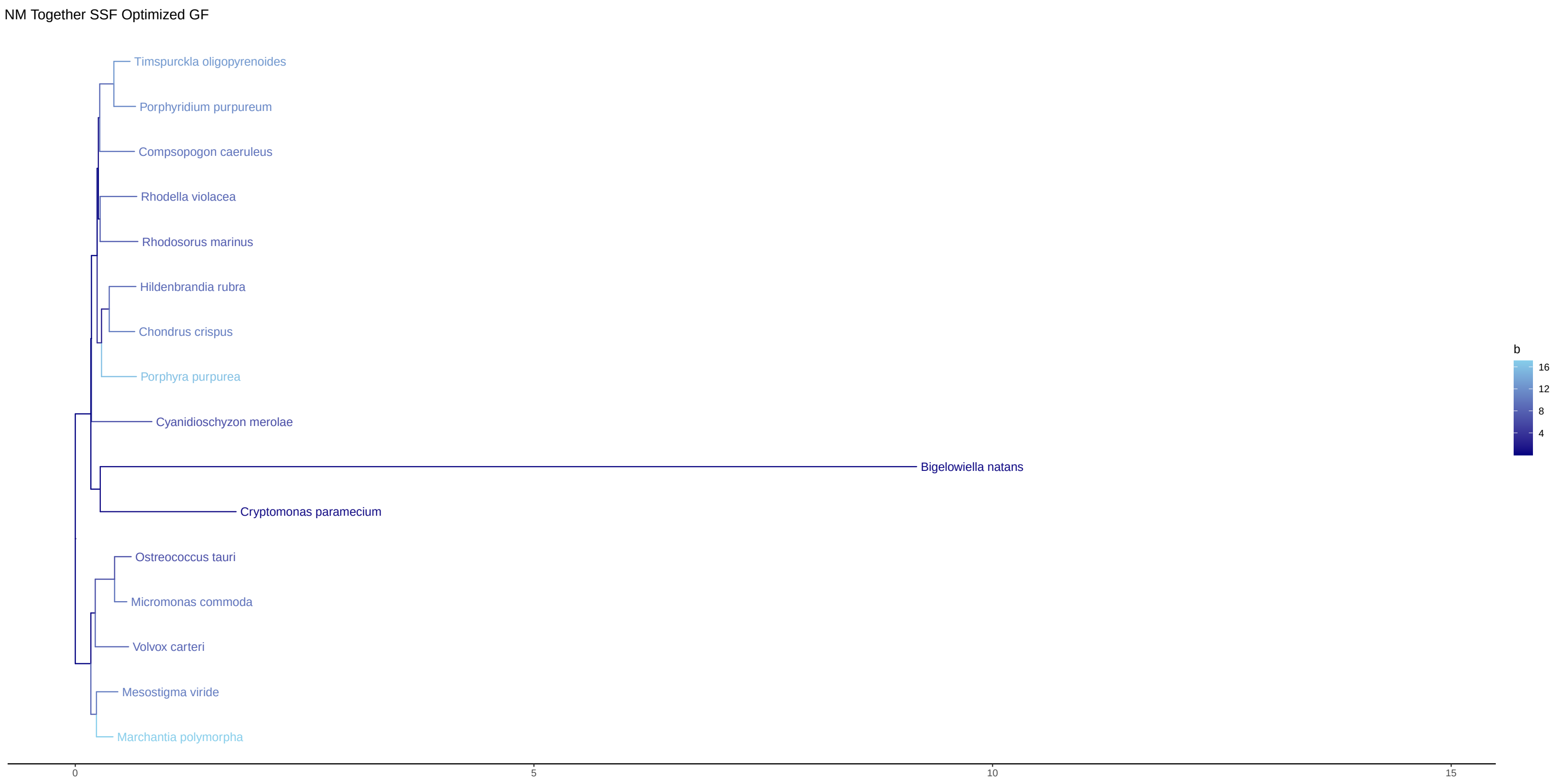


Fig. S18. NM Together LG+C20+G+SSF, $\mathcal{G}/\mathcal{F}$ = GARPVMTHQ/FYCINK

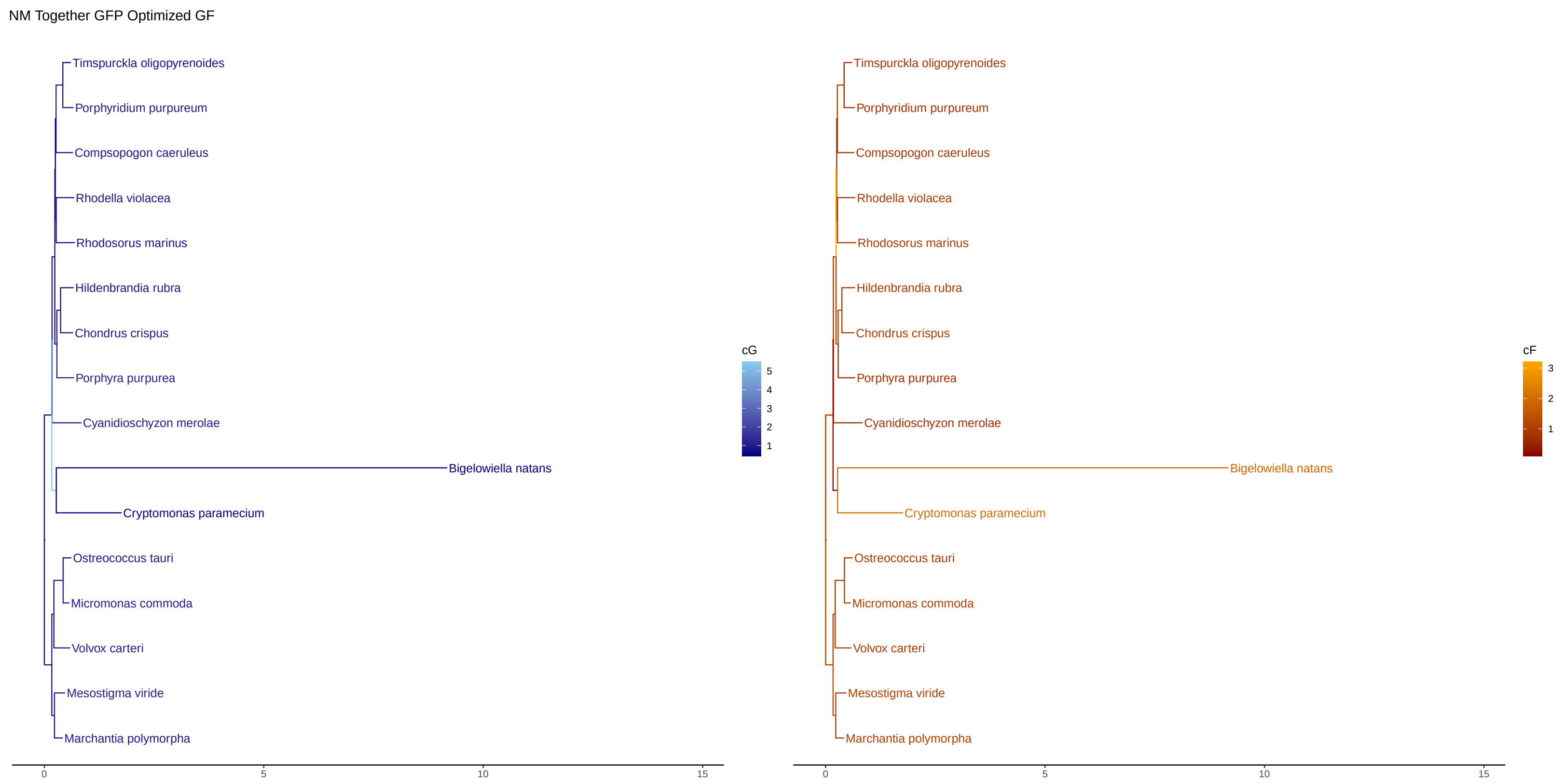


Fig. S19. NM Together LG+C20+G+GFP $\mathcal{G}/\mathcal{F}$ = GARPVMTHQ/FYCINK

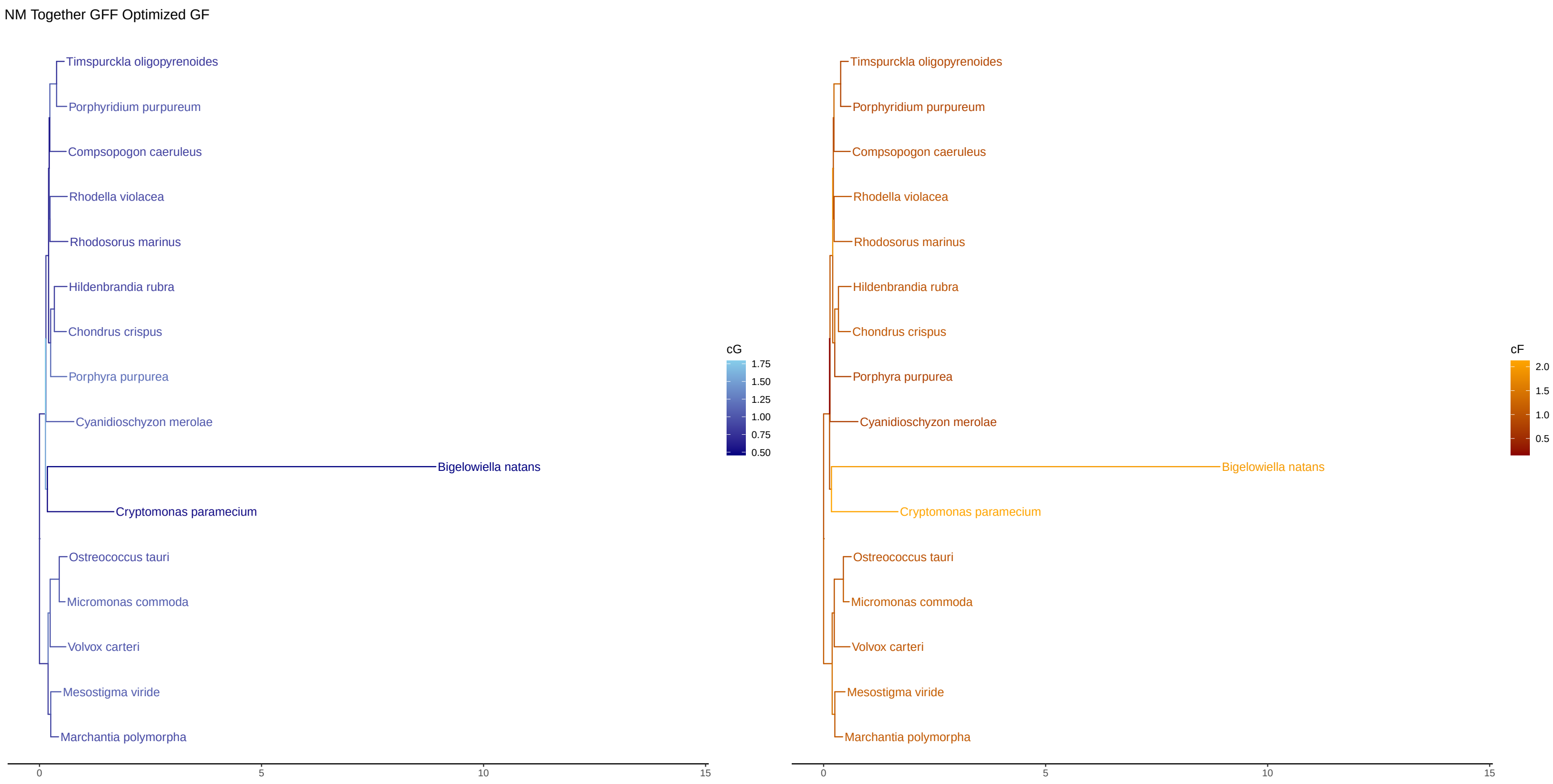


Fig. S20. NM Together LG+C20+G+GFF, $\mathcal{G}/\mathcal{F}$ = GARPVMTHQ/FYCINK
