## Supplementary material for "Modeling site-and-branch-heterogeneity with GFmix": aa_empirical.h

```
/*----------------------------------------------------------------------------*/
double pamprobmat[400] =
  {
      -0.01522976,-0.00746819,-0.13934468, 0.11755315,-0.00212101,
        0.01558456,-0.07408235,-0.00322387, 0.01375826, 0.00448826,
        0.00154174, 0.02013313,-0.00159183,-0.00069275,-0.00399898,
        0.08414055,-0.01188178,-0.00029870, 0.00220371, 0.00042546,
      -0.07765582,-0.00712634,-0.03683209,-0.08065755,-0.00462872,
       -0.03791039, 0.10642147,-0.00912185, 0.01436308,-0.00133243,
        0.00166346, 0.00624657,-0.00003363,-0.00128729,-0.00690319,
        0.17442028,-0.05373336,-0.00078751,-0.00038151, 0.01382718,
      -0.08810973,-0.04081786,-0.04066004,-0.04736004,-0.03275406,
       -0.03761164,-0.05047487,-0.09086213,-0.03269598,-0.03558015,
       -0.08407966,-0.07970977,-0.01504743,-0.04011920,-0.05182232,
       -0.07026991,-0.05846931,-0.01016998,-0.03047472,-0.06280511,
       0.02513756,-0.00578333, 0.09865453, 0.01322314,-0.00310665,
        0.05880899,-0.09252443,-0.02986539,-0.03127460, 0.01007539,
       -0.00360119,-0.01995024, 0.00094940,-0.00145868,-0.01388816,
        0.11358341,-0.12127513,-0.00054696,-0.00055627, 0.00417284,
       0.16517316,-0.00254742,-0.03318745,-0.01984173, 0.00031890,
       -0.02817810, 0.02661678,-0.01761215, 0.01665112, 0.10513343,
       -0.00545026, 0.01827470,-0.00207616,-0.00763758,-0.01322808,
       -0.02202576,-0.07434204, 0.00020593, 0.00119979,-0.10827873,
       0.16088826, 0.00056313,-0.02579303,-0.00319655, 0.00037228,
       -0.03193150, 0.01655305,-0.03028640, 0.01367746,-0.11248153,
        0.00778371, 0.02675579, 0.00243718, 0.00895470,-0.01729803,
       -0.02686964,-0.08262584, 0.00011794,-0.00225134, 0.09415650,
      -0.01739295, 0.00572017,-0.00712592,-0.01100922,-0.00870113,
       -0.00663461,-0.01153857,-0.02248432,-0.00382264,-0.00358612,
       -0.00139345,-0.00971460,-0.00133312, 0.01927783,-0.01053838,
       -0.00911362,-0.01010908, 0.09417598, 0.01763850,-0.00955454,
       0.01728888, 0.01344211, 0.01200836, 0.01857259,-0.17088517,
        0.01457592, 0.01997839, 0.02844884, 0.00839403, 0.00196862,
        0.01391984, 0.03270465, 0.00347173,-0.01940984, 0.01233979,
        0.00542887, 0.01008836, 0.00126491,-0.02863042, 0.00449764,
      -0.02881366,-0.02184155,-0.01566086,-0.02593764,-0.04050907,
       -0.01539603,-0.02576729,-0.05089606,-0.00597430, 0.02181643,
        0.09835597,-0.04040940, 0.00873512, 0.12139434,-0.02427882,
       -0.02945238,-0.01566867,-0.01606503, 0.09475319, 0.02238670,
       0.04080274,-0.02869626,-0.05191093,-0.08435843, 0.00021141,
        0.13043842, 0.00871530, 0.00496058,-0.02797641,-0.00636933,
        0.02243277, 0.03640362,-0.05735517, 0.00196918,-0.02218934,
       -0.00608972, 0.02872922, 0.00047619, 0.00151285, 0.00883489,
      -0.02623824, 0.00331152, 0.03640692, 0.04260231,-0.00038223,
       -0.07480340,-0.01022492,-0.00426473, 0.01448116, 0.01456847,
        0.05786680, 0.03368691,-0.10126924,-0.00147454, 0.01275395,
        0.00017574,-0.01585206,-0.00015767,-0.00231848, 0.02310137,
      -0.00846258,-0.01508106,-0.01967505,-0.02772004, 0.01248253,
       -0.01331243,-0.02569382,-0.04461524,-0.02207075, 0.04663443,
        0.19347923,-0.02745691, 0.02288515,-0.04883849,-0.01084597,
       -0.01947187,-0.00081675, 0.00516540,-0.07815919, 0.08035585,
      -0.06553111, 0.09756831, 0.00524326,-0.00885098, 0.00756653,
        0.02783099,-0.00427042,-0.16680359, 0.03951331,-0.00490540,
        0.01719610, 0.15018204, 0.00882722,-0.00423197,-0.01919217,
       -0.02963619,-0.01831342,-0.00524338, 0.00011379,-0.02566864,
      -0.07494341,-0.11348850, 0.00241343,-0.00803016, 0.00492438,
        0.00711909,-0.00829147, 0.05793337, 0.02734209, 0.02059759,
       -0.02770280, 0.14128338, 0.01532479, 0.00364307, 0.05968116,
       -0.06497960,-0.08113941, 0.00319445,-0.00104222, 0.03553497,
       0.05948223,-0.08959930, 0.03269977,-0.03272374,-0.00365667,
       -0.03423294,-0.06418925,-0.05902138, 0.05746317,-0.02580596,
        0.01259572, 0.05848832, 0.00672666, 0.00233355,-0.05145149,
        0.07348503, 0.11427955, 0.00142592,-0.01030651,-0.04862799,
      -0.01606880, 0.05200845,-0.01212967,-0.06824429,-0.00234304,
        0.01094203,-0.07375538, 0.08808629, 0.12394822, 0.02231351,
       -0.03608265,-0.06978045,-0.00618360, 0.00274747,-0.01921876,
       -0.01541969,-0.02223856,-0.00107603,-0.01251777, 0.05412534,
       0.01688843, 0.05784728,-0.02256966,-0.07072251,-0.00422551,
       -0.06261233,-0.08502830, 0.08925346,-0.08529597, 0.01519343,
       -0.05008258, 0.10931873, 0.00521033, 0.02593305,-0.00717855,
        0.02291527, 0.02527388,-0.00266188,-0.00871160, 0.02708135,
      -0.04233344, 0.00076379, 0.01571257, 0.04003092, 0.00901468,
        0.00670577, 0.03459487, 0.12420216,-0.00067366,-0.01515094,
        0.05306642, 0.04338407, 0.00511287, 0.01036639,-0.17867462,
       -0.02289440,-0.03213205, 0.00017924,-0.01187362,-0.03933874,
       0.01284817,-0.01685622, 0.00724363, 0.01687952,-0.00882070,
       -0.00555957, 0.01676246,-0.05560456,-0.00966893, 0.06197684,
       -0.09058758, 0.00880607, 0.00108629,-0.08308956,-0.08056832,
       -0.00413297, 0.02973107, 0.00092948, 0.07010111, 0.13007418,
       0.00700223,-0.01347574, 0.00691332, 0.03122905, 0.00310308,
        0.00946862, 0.03455040,-0.06712536,-0.00304506, 0.04267941,
       -0.10422292,-0.01127831,-0.00549798, 0.11680505,-0.03352701,
       -0.00084536, 0.01631369, 0.00095063,-0.09570217, 0.06480321
    }; /* transpose of pamprobmat of proml */
/*----------------------------------------------------------------------------*/
double pameigmat[20] = 
    {
-2.2091252,-1.9297602,0.000476,-1.7477817,-1.6575549,
-1.5504543,-0.2112213,-0.2685727,-0.2976402,-1.3440755,
-1.2926992,-0.4293227,-0.5356688,-1.1064786,-1.0480731,
-0.8760449,-0.7142318,-0.7381851,-0.7806557,-0.8127024
    }; /* pameigmat of proml * 100 */
/*----------------------------------------------------------------------------*/
double jtteigmat[20] =
  {
    0,-0.7031123,-0.6484345,-0.6086499,-0.5514432,
    -0.772664,-0.8643413,-1.062076,-0.9965552,-1.167181,
    -1.222242,-0.4589201,-1.310371,-1.404804,-0.3170582,
    -0.347935,-1.531168,-1.602119,-1.799145,-1.891189
  };/* new jtt matrix decomposition - Elisabeth  Tillier */
/*----------------------------------------------------------------------------*/
double jttprobmat[400] =
{
 0.076999996, 0.051000003, 0.043000004, 0.051999998, 0.019999996, 0.041,
  0.061999994, 0.073999997, 0.022999999, 0.052000004, 0.090999997, 0.058999988,
  0.024000007, 0.04, 0.050999992, 0.069, 0.059000006, 0.014000008, 0.032000004,
  0.066000005,
 0.015604455, -0.068062363, 0.020106264, 0.070723273, 0.011702977, 0.009674053,
  0.074000798, -0.169750458, 0.005560808, -0.008208636, -0.012305869,
 -0.063730179, -0.005674643, -0.02116828, 0.104586169, 0.016480839, 0.016765139,
  0.005936994, 0.006046367, -0.0082877,
 -0.049778281, -0.007118197, 0.003801272, 0.070749616, 0.047506147,
   0.006447017, 0.090522425, -0.053620432, -0.008508175, 0.037170603,
   0.051805545, 0.015413608, 0.019939916, -0.008431976, -0.143511376,
  -0.052486072, -0.032116542, -0.000860626, -0.02535993, 0.03843545,
 -0.028906423, 0.092952047, -0.009615343, -0.067870117, 0.031970392,
   0.048338335, -0.054396304, -0.135916654, 0.017780083, 0.000129242,
   0.031267424, 0.116333586, 0.007499746, -0.032153596, 0.033517051,
  -0.013719269, -0.00347293, -0.003291821, -0.02158326, -0.008862168,
 0.037181176, -0.023106564, -0.004482225, -0.029899635, 0.118139633,
 -0.032298569, -0.04683198, 0.05566988, -0.012622847, 0.002023096,
 -0.043921088, -0.04792557, -0.003452711, -0.037744513, 0.020822974,
  0.036580187, 0.02331425, -0.004807711, -0.017504496, 0.01086673,
 0.044754061, -0.002503471, 0.019452517, -0.015611487, -0.02152807,
 -0.013131425, -0.03465365, -0.047928912, 0.020608851, 0.067843095,
 -0.122130014, 0.002521499, 0.013021646, -0.082891087, -0.061590119,
  0.016270856, 0.051468938, 0.002079063, 0.081019713, 0.082927944,
 0.058917882, 0.007320741, 0.025278141, 0.000357541, -0.002831285,
 -0.032453034, -0.010177288, -0.069447924, -0.034467324, 0.011422358,
 -0.128478324, 0.04309667, -0.015319944, 0.113302422, -0.035052393,
  0.046885372, 0.06185183, 0.00175743, -0.06224497, 0.020282093,
 -0.014562092, 0.022522921, -0.007094389, 0.03480089, -0.000326144,
 -0.124039037, 0.020577906, -0.005056454, -0.081841576, -0.004381786,
  0.030826152, 0.091261631, 0.008878828, -0.02829487, 0.042718836,
 -0.011180886, -0.012719227, -0.000753926, 0.048062375, -0.009399129,
 0.033789571, -0.013512235, 0.088010984, 0.017580292, -0.006608005,
 -0.037836971, -0.061344686, -0.034268357, 0.018190209, -0.068484614,
  0.120024744, -0.00319321, -0.001349477, -0.03000546, -0.073063759,
  0.081912399, 0.0635245, 0.000197, -0.002481798, -0.09108114,
 -0.113947615, 0.019230545, 0.088819683, 0.064832765, 0.001801467,
 -0.063829682, -0.072001633, 0.018429333, 0.057465965, 0.043901014,
 -0.048050874, -0.001705918, 0.022637173, 0.017404665, 0.043877902,
 -0.017089594, -0.058489485, 0.000127498, -0.029357194, 0.025943972,
 0.01512923, 0.023603725, 0.006681954, 0.012360216, -0.000181447,
 -0.023011838, -0.008960024, -0.008533239, 0.012569835, 0.03216118,
  0.061986403, -0.001919083, -0.1400832, -0.010669741, -0.003919454,
 -0.003707024, -0.026806029, -0.000611603, -0.001402648, 0.065312824,
 -0.036405351, 0.020816769, 0.011408213, 0.019787053, 0.038897829,
   0.017641789, 0.020858533, -0.006067252, 0.028617353, -0.064259496,
  -0.081676567, 0.024421823, -0.028751676, 0.07095096, -0.024199434,
  -0.007513119, -0.028108766, -0.01198095, 0.111761119, -0.076198809,
 0.060831772, 0.144097327, -0.069151377, 0.023754576, -0.003322955,
 -0.071618574, 0.03353154, -0.02795295, 0.039519769, -0.023453968,
 -0.000630308, -0.098024591, 0.017672997, 0.003813378, -0.009266499,
 -0.011192111, 0.016013873, -0.002072968, -0.010022044, -0.012526904,
 -0.050776604, 0.092833081, 0.044069596, 0.050523021, -0.002628417,
   0.076542572, -0.06388631, -0.00854892, -0.084725311, 0.017401063,
  -0.006262541, -0.094457679, -0.002818678, -0.0044122, -0.002883973,
   0.028729685, -0.004961596, -0.001498627, 0.017994575, -0.000232779,
 -0.01894566, -0.007760205, -0.015160993, -0.027254587, 0.009800903,
  -0.013443561, -0.032896517, -0.022734138, -0.001983861, 0.00256111,
   0.024823166, -0.021256768, 0.001980052, 0.028136263, -0.012364384,
  -0.013782446, -0.013061091, 0.111173981, 0.021702122, 0.00046654,
 -0.009444193, -0.042106824, -0.02535015, -0.055125574, 0.006369612,
  -0.02945416, -0.069922064, -0.067221068, -0.003004999, 0.053624311,
   0.128862984, -0.057245803, 0.025550508, 0.087741073, -0.001119043,
  -0.012036202, -0.000913488, -0.034864475, 0.050124813, 0.055534723,
 0.145782464, -0.024348311, -0.031216873, 0.106174443, 0.00202862,
  0.02653866, -0.113657267, -0.00755018, 0.000307232, -0.051241158,
  0.001310685, 0.035275877, 0.013308898, 0.002957626, -0.002925034,
 -0.065362319, -0.071844582, 0.000475894, -0.000112419, 0.034097762,
 0.079840455, 0.018769331, 0.078685899, -0.084329807, -0.00277264,
 -0.010099754, 0.059700608, -0.019209715, -0.010442992, -0.042100476,
 -0.006020556, -0.023061786, 0.017246106, -0.001572858, -0.006703785,
  0.056301316, -0.156787357, -0.000303638, 0.001498195, 0.051363455,
 0.049628261, 0.016475144, 0.094141653, -0.04444633, 0.005206131,
 -0.001827555, 0.02195624, 0.013066683, -0.010415582, -0.022338403,
  0.007837197, -0.023397671, -0.002507095, 0.005177694, 0.017109561,
 -0.202340113, 0.069681441, 0.000120736, 0.002201146, 0.004670849,
 0.089153689, 0.000233354, 0.010826822, -0.004273519, 0.001440618,
  0.000436077, 0.001182351, -0.002255508, -0.000700465, 0.150589876,
 -0.003911914, -0.00050154, -0.004564983, 0.00012701, -0.001486973,
 -0.018902754, -0.054748555, 0.000217377, -0.000319302, -0.162541651
};
/*------------------------------------------------------------------------*/
double wagprobmat[400] =
  {
    8.6627904331395514e-02, 4.3972002198600091e-02, 3.9089401954470158e-02,
    5.7045102852255107e-02, 1.9307800965390060e-02, 3.6728101836405132e-02,
    5.8058902902945193e-02, 8.3251804162590315e-02, 2.4431301221565036e-02,
    4.8466002423300139e-02, 8.6209004310450116e-02, 6.2028603101430192e-02,
    1.9502700975135050e-02, 3.8431901921595082e-02, 4.5763102288155187e-02,
    6.9517903475895371e-02, 6.1012703050635379e-02, 1.4385900719295025e-02,
    3.5274201763710086e-02, 7.0895603544780264e-02, 3.5561607230076675e-02,
    2.1452034798925650e-02, 2.5289700297807173e-02, 5.4263632130575258e-02,
    -8.2227713139344395e-03, 1.8809713405152310e-02, 4.5250485177498877e-02,
    1.1396137747298690e-01, 2.2695735884200692e-03,-5.3709625003230864e-02,
    -1.1682537170907210e-01, 3.7849568737497064e-02,-1.8090959226090730e-02,
    -8.2134946483552643e-02, 2.8286855860628530e-02, 3.2195382055601636e-02,
    1.5315516810597547e-02,-3.6410812788007695e-02,-5.7594741008656421e-02,
    -5.7516220033223035e-02,-1.3908508233028910e-02, 1.0741058910567900e-02,
    7.7142089656707467e-03, 1.5297298394271509e-02,-7.2692369772449856e-03,
    1.8756949033626091e-03, 6.6642638253604945e-03, 5.6835688582020170e-02,
    1.1043132925221123e-02,-5.6826449999744365e-02,-8.7968406260232349e-02,
    4.6971770804279701e-03,-1.7222872652551572e-02, 2.0989711261972341e-02,
    -7.8464103074134157e-03, 2.9840656912852133e-03,-1.6256261561745401e-02,
    9.4377548761120852e-02, 4.6171262350368288e-02,-7.2092965659688091e-02,
    -9.7473641707852988e-03, 9.5017223630150319e-03, 8.0373879255763742e-03,
    1.9992975630317825e-02,-1.3370527648434624e-01, 1.2188355967379839e-02,
    1.9340811379916827e-02,-2.1308575843419986e-02, 1.0519629982552198e-02,
    6.4157143850267911e-03, 2.2175620508616836e-02, 1.8379806007210828e-02,
    3.6995050827064472e-03, 2.0647226873777309e-02, 1.2527476168275313e-02,
    -6.4803924513660328e-03, 1.2234185725789516e-04,-8.5073058616038921e-03,
    1.7744672681986148e-02,-1.5443320020940997e-03, 1.7001972980605972e-02,
    -6.4899067797287296e-03,-1.4677169244502764e-02,-2.3288867015549917e-02,
    -2.2344319072353971e-02,-8.8507859286603794e-03,-1.4074219549986493e-02,
    4.1660841243721145e-02,-3.0194652331642903e-02, 4.9405058377634432e-02,
    6.7156610040389128e-02,-1.0997957541837343e-02, 1.4151027692304656e-02,
    -8.4658234578690303e-02,-1.3575378347862733e-03,-4.8548192818665065e-03,
    9.5855773184325577e-03, 5.8067161411780553e-02,-1.0143174601847883e-01,
    6.6191966113215886e-02, 5.5940563437940623e-04, 3.8468104953025675e-02,
    1.1444860698120021e-02, 2.6194302421481647e-02, 1.2403239604190725e-02,
    2.7633006762398871e-02, 3.6460369427774607e-02,-2.2632388532992354e-01,
    1.3241927942500996e-02,-1.2651903029776661e-02,-2.6892304398426634e-02,
    5.0471948264076055e-02,-2.8051221154347500e-03,-3.9648723892390714e-02,
    7.6543445589337844e-02, 9.7970457414629560e-03, 1.9668413939263687e-02,
    2.0842116738760217e-02,-2.2660972140595224e-02,-1.2745276810225298e-02,
    2.1380209161766427e-02,-3.4252993456739905e-02,-2.4153790058581974e-02,
    -5.6900611243356061e-02,-4.3030665245530836e-03,-2.5505194038801030e-02,
    -4.8250760732301883e-02, 4.5499525791566957e-02,-1.4454806196411856e-02,
    -9.5258032419141773e-03,-8.2878799321754893e-03,-5.0304500044025466e-02,
    -4.5270227357394853e-03, 1.9332947733615973e-02, 1.8449411930725712e-01,
    1.0377118224484075e-02,-1.3477904016439130e-03,-1.5276819290023075e-03,
    1.0873853651013947e-02,-8.6158733344581327e-03, 3.1100196651221757e-02,
    -1.2976272983272708e-01, 2.2634933891269058e-02, 1.2962746125401634e-01,
    -5.2009218084257528e-04,-2.3617222520153473e-02, 6.2868517095620302e-02,
    -3.7127268321369790e-02,-2.3181990407100242e-02, 1.5819209122190911e-02,
    -3.3730072385244267e-02,-8.9541716031825591e-02,-5.5384754870589194e-03,
    4.9888520706533887e-03, 4.1515481016138101e-04, 1.4760563566797643e-02,
    1.6493326334473216e-02, 6.3476703985155987e-03, 4.8024236271855328e-03,
    3.3161258344216696e-02,-9.3397798665151521e-02, 2.3762091403094370e-03,
    -1.3365608713028249e-03, 7.6254108489326455e-02, 1.4229002852796149e-02,
    1.8923156288257432e-02, 5.0117035684255032e-02, 3.3388769595753244e-02,
    1.3562926640855594e-03,-3.7089728993583766e-02, 1.5135867581356804e-01,
    1.3749492828968005e-03, 9.9685669536694350e-03, 2.6115791194251045e-02,
    3.7042081516679919e-02,-6.5569109873164430e-02,-9.9371842878282005e-02,
    2.0237874125610749e-04,-4.3787598558267790e-02,-8.2154378377351428e-02,
    5.0858849149674236e-02, 2.0749290641652009e-02,-9.4142728743790947e-03,
    -7.2050029432379923e-03,-2.9056780975804837e-03,-5.8328604459562637e-03,
    1.0895984289926943e-02,-2.1484675813403159e-02,-8.4324050465665421e-02,
    -2.9082525315236033e-02,-2.5629155791891959e-02, 3.5899110364035841e-02,
    -4.1581098434880521e-03, 1.1756461558453200e-01,-1.6372687851347458e-02,
    3.6899540291678225e-02, 4.3577632946987375e-02, 1.0052496452281861e-03,
    -7.9765256989734579e-02,-3.1275996481794736e-02, 6.4648213300009832e-02,
    -4.5286836663019657e-02, 2.2896515019620048e-02,-3.0613993299013528e-02,
    -4.7249975002003576e-03,-6.7850482701700609e-03,-4.0142303469059344e-02,
    -2.5295996131222860e-02, 3.3258862145181561e-02,-7.3493254404135344e-02,
    1.2744615710956228e-01,-3.5954598776571818e-02, 1.2075402272480619e-02,
    -2.3149650954854401e-02,-2.8060336091316512e-02, 8.6902702943959576e-02,
    8.9133169653849195e-02, 8.9689278536865199e-04,-6.9083221755479015e-03,
    -1.1684257749492037e-01,-1.3188712339839277e-02,-2.5201546418154058e-02,
    1.5471166235240700e-02,-1.4048232951367973e-02, 4.6376166126662970e-04,
    1.2962754944902356e-02,-1.9256689075660898e-02, 4.2002592451201972e-03,
    1.0955954320107231e-01, 1.9302206061051940e-02,-6.1048245337646030e-02,
    -3.0302612215077924e-02,-8.2483661742359469e-03, 7.1711114402665671e-02,
    1.0333159680581243e-03,-4.1002195974806765e-03,-9.4410693690943936e-04,
    4.0336913416456170e-03,-9.4394860332760094e-02, 3.1995778318108868e-02,
    -1.4297590994888315e-01, 6.0680069216446461e-02, 6.9733590321782835e-02,
    7.9385622621616220e-02, 7.4498967202616556e-04,-7.6929394578439836e-02,
    -9.6124188201433924e-02, 5.0463102117680016e-03,-8.8133248206431775e-03,
    3.2895233231778044e-02,-3.0372506779427549e-03,-1.7858974511085993e-02,
    -2.0502868715590932e-03, 2.2060400797727824e-03, 1.2470540858441161e-02,
    3.8547511534476366e-02, 6.3566948157559172e-02,-1.0329335178926749e-03,
    -6.7824023202805693e-03,-9.6721904575058585e-03,-1.5461363123684130e-01,
    -9.7966066871409394e-02, 1.8215739459980639e-02,-5.7082699302948479e-02,
    4.4578879211827991e-03, 5.1207951215528992e-02, 3.0743349448865920e-02,
    2.5680743430060361e-02,-1.3277648416397577e-02, 2.4994769795659139e-02,
    -2.3452872369277612e-02, 9.6343766531534869e-02, 2.3243876300254276e-02,
    2.7382354224967173e-04, 1.0476648413065818e-02,-1.0957598150792867e-02,
    8.8334756509341320e-02, 1.6910539222818770e-03, 4.0262744885943974e-03,
    -2.2340124630932757e-02, 2.9323802928908304e-02,-5.6629668320205995e-02,
    5.6923395200187223e-02,-1.1678029942639301e-02,-5.3902282183251491e-04,
    -2.5949292388769449e-02,-4.3300904000318892e-02,-6.8531829578852470e-03,
    -2.5826092550178853e-03,-1.9840557360383025e-03, 5.9597976961404744e-02,
    1.1363016187151703e-01,-9.4159710366617327e-02,-2.1204494101847555e-03,
    -5.9149756224307335e-04, 1.4424533762850252e-02,-5.9906144009341321e-02,
    1.0052536287630033e-03,-4.6915230824837368e-04, 3.1858594725711925e-02,
    7.1271610647088077e-02,-4.0083782421349475e-02, 4.9505006163231856e-02,
    1.6180042487677501e-02,-3.6424677513288337e-04,-3.9192606765110376e-02,
    -6.5903383940019974e-02,-1.1654251492515226e-02, 1.5754244584066880e-03,
    -1.3775083805893790e-03,-4.9027613056858627e-02, 8.7935375742901292e-02,
    9.0038005689940109e-02, 1.5668629704927934e-03, 1.5071587512597738e-03,
    2.0990694072840705e-03,-9.4421508671830648e-02, 8.0999281239943499e-04,
    -2.9920209955590594e-03,-1.7471626631715847e-02, 3.1827000793682704e-02,
    -8.9801507968671531e-03,-8.1412546908303754e-02, 4.0176295130136160e-02,
    1.7340891665920245e-03,-1.0103712010989639e-01, 1.8677421679886945e-02,
    1.0982061460000879e-02, 3.0943297608324503e-02,-3.1274338581654446e-02,
    1.4546895520865988e-02, 8.3820295163042610e-02, 9.1514099727366789e-04,
    7.5019946230749275e-04, 3.9872439811473809e-03,-1.1566371882457203e-01,
    9.0469726577489334e-02, 5.8630468948720231e-04, 8.6721774251191685e-04,
    8.0846852485450312e-03, 1.3252155931930222e-02, 2.0087005643411766e-03,
    -3.0844331179362222e-02,-2.7207484416003614e-02, 7.8063211256813099e-04,
    -5.8892343648989209e-02, 7.2506273771614727e-02,-3.5140587683799166e-03,
    1.2342251847686935e-02, 1.4106845444653088e-01,-1.1455607779569450e-02,
    1.3280996346457177e-02,-1.1501732586922284e-02,-2.0526810655552255e-03,
    -2.4905301264579274e-03, 6.5487659964863984e-02,-3.3456428537590670e-02,
    1.2797423882205969e-04,-1.8469921022642227e-03,-1.3759290901372054e-01,
    7.5959464423422482e-02,-7.8079495263044972e-03,-1.5795332034633989e-02,
    6.6873552357592098e-02, 2.0832985715545219e-03, 8.3761342494846791e-02,
    -1.1563215140324860e-01, 2.7686944696295687e-03,-9.0610814116160506e-03,
    7.9088716942001827e-02,-7.7083591132996098e-03, 2.9720288262336573e-03,
    -1.5633366477761715e-02, 9.7597088374988625e-04, 1.3625200711270035e-03,
    -1.0130997497900149e-01, 3.9905002719784068e-02, 7.2947724062451724e-04,
    3.0357030316817512e-03,-8.6567557086382335e-02, 5.4297039373508757e-02,
    2.1867453707076000e-02, 1.2564670875940023e-01,-7.1811633558895860e-02,
    1.8167330492839856e-03,-2.7826226325727101e-02, 6.6492461594168092e-02,
    5.6451618942812058e-04,-9.2930305715977039e-03, 1.3365448768529766e-02,
    2.4829592264043522e-03,-4.7050626252299381e-02,-2.0413104847365296e-04,
    2.1653157344373665e-03, 5.5416494397296989e-03,-1.4203499202174025e-01,
    3.2785279562103635e-02, 5.2415732948228751e-04,-4.0891908103757503e-04,
    -2.8920163873781084e-02
  };
/*----------------------------------------------------------------------------*/
double wageigmat[20] =
  {
    0.0,-3.5122707074250598e-01,-4.5515856739462934e-01,
    -4.8893257594445549e-01,-5.5037844703306915e-01,-5.5499787467373596e-01,
    -6.2967289889813272e-01,-7.9074063942384476e-01,-8.5502559203244555e-01,
    -9.3428366563267362e-01,-1.0050214079557256e+00,-1.0712190618081447e+00,
    -1.2188397247689513e+00,-1.2762113612487385e+00,-1.3890446164301007e+00,
    -1.4070264931532848e+00,-1.5483111871701543e+00,-1.6655486317312065e+00,
    -1.7075342548435470e+00,-1.7791695407992054e+00
  };
/*----------------------------------------------------------------------------*/

double mtREV24probmat[400] =
  {
    7.2000000000000106e-02, 1.9000000000000055e-02, 3.8999999999999965e-02,
    1.9000000000000052e-02, 5.9999999999999932e-03, 2.5000000000000005e-02,
    2.4000000000000014e-02, 5.5999999999999980e-02, 2.8000000000000004e-02,
    8.7999999999999884e-02, 1.6899999999999996e-01, 2.3000000000000000e-02,
    5.3999999999999944e-02, 6.1000000000000006e-02, 5.3999999999999965e-02,
    7.1999999999999925e-02, 8.6000000000000076e-02, 2.9000000000000008e-02,
    3.3000000000000002e-02, 4.3000000000000010e-02, 1.3942001744884941e-02,
    1.7481249715609417e-03, 7.2098267845172922e-03, 4.0055323235337129e-03,
    6.6957331542989848e-04, 5.6165699885365209e-03, 7.3764646088821974e-03,
    1.4741162797926062e-02, 5.4438517761326750e-03, 1.4728635694750780e-02,
    2.1184759765052821e-02, 3.9988470264509272e-03, 7.9223463271922232e-03,
    9.4675726067677511e-03, 1.1973580294682632e-02, 1.0954529915146383e-02,
    1.4613059560863745e-02,-1.6765340899576234e-01, 4.5570674834516248e-03,
    7.4999020099992689e-03,-1.7008160256183784e-02, 1.3169101574322731e-01,
    2.3918502205267883e-03, 4.8146728619157348e-03, 3.7079431166765108e-04,
    1.7025537256968083e-02, 1.4177856654347076e-02,-8.4415873772621314e-04,
    1.2885369346771012e-02,-2.9626896928023543e-02,-5.5187110546076557e-02,
    9.7631034102268859e-03,-1.6808738207762866e-02,-1.9077642848672336e-02,
    -4.2709200403695472e-03,-1.2145410691462724e-02,-2.0509119935462321e-02,
    -6.8854541823322403e-04,-2.0674863427758024e-03,-1.4886009852902164e-02,
    -5.5087216993520722e-03, 9.5078228021539465e-03,-1.5999136307979834e-03,
    -9.4971684157689057e-03, 1.0060701165645351e-03, 4.5327612602892231e-03,
    -1.3873027077998460e-02,-2.2153992091216332e-01, 7.5948692295378373e-03,
    3.5415710400438824e-02, 7.8417538041002613e-02, 6.0613184721215838e-04,
    2.0687251140545902e-02, 3.5346711238379298e-02, 1.6360804460343031e-02,
    -2.7833337409606364e-03, 1.7272364656047253e-02,-4.6369055719090316e-03,
    1.5636303114467818e-02, 1.7054652741968001e-02, 2.1321984749694576e-02,
    2.7513605788343960e-02,-2.6385479666787059e-02,-5.1872425915470408e-02,
    9.2757207130785544e-04,-2.8498775070132234e-02,-1.2101936870418238e-01,
    4.4346493116370400e-02,-1.9621870912195003e-02, 3.6534564673249011e-02,
    7.2697883853844736e-02,-2.5087872319727464e-02, 1.9813099505476339e-02,
    2.2877523824216123e-02,-4.2062383154004073e-03, 3.0203529210638767e-03,
    1.7109263481935550e-02,-3.8689488111286448e-03,-4.2266092170117047e-03,
    1.8625244946533049e-02, 8.0295085911015508e-03,-7.7652703001077020e-03,
    1.2908802291884083e-02,-3.2254439516033475e-03, 5.1636076420900792e-04,
    1.8406862126309893e-02,-3.0967062226226003e-02,-7.7751353319695680e-03,
    9.6382460916077384e-03,-3.8369235165013602e-02,-8.7859473708315400e-02,
    9.1729837057034792e-03,-2.0310651942508497e-02,-5.6990742028812734e-02,
    2.0537260105731484e-01, 2.1040143481269598e-02, 3.5154329962107629e-03,
    2.2169292229626096e-03,-1.7802963484865878e-02,-1.9751892189150723e-02,
    -4.2352085504018908e-02,-1.3831405098563140e-02, 3.0784177175352114e-02,
    1.0363158716562954e-02, 6.9111014195040125e-03, 3.1983357784537668e-02,
    -4.9791242820397770e-02, 1.1460162816995668e-02, 8.1452029005183849e-02,
    -5.5344377122843152e-02,-7.1286350215609959e-02, 1.4651214381280650e-02,
    -2.6922558358635457e-02, 7.1073171178518399e-02,-2.3867301165933139e-02,
    -9.9885267116555793e-03,-3.3589613884042011e-02,-5.0515675541924689e-04,
    1.0292480608708640e-01,-3.4124560927903308e-02,-4.3058586023707560e-02,
    9.0621842363354771e-04,-5.3857856866972954e-02,-9.0862868379940939e-02,
    -1.5458541767351432e-03, 3.0992307207084349e-03, 6.4623925473236671e-02,
    2.4305785287191808e-02,-5.5704599277222391e-03,-1.0764393638331027e-02,
    2.9214249503415501e-02,-1.5620414729831871e-02,-5.0382005100120678e-03,
    8.5459916005763428e-02, 5.7244011536436198e-02,-3.6356956001437547e-02,
    -3.3243427600992484e-02, 9.2261841663216144e-04, 3.9654670712339114e-02,
    -9.5116082236727621e-03,-3.5607518230276948e-02, 1.2653699435606456e-02,
    -1.0273526643183511e-02, 7.2619293146026789e-02,-4.7511211249969725e-03,
    -7.4929766636335118e-02, 2.8442952487810307e-03, 6.7643140941294260e-03,
    -3.3296478375647545e-02,-1.2457130295849526e-02, 2.4223195942225113e-02,
    -6.1677621895427845e-02,-6.6009948577442452e-03, 9.1868572637285081e-02,
    6.0561241106405560e-02,-2.1628970506541147e-02,-2.8897196285090968e-02,
    -4.3914194458858941e-04, 3.0258819509548073e-02,-1.1233964324325392e-02,
    3.4626972906581573e-02, 1.1244088398481194e-03,-3.0889794439813148e-03,
    -8.4811822919813908e-03, 7.2320795746895136e-02,-2.9105052707592012e-02,
    1.2952503184698753e-02,-9.7471108874533346e-03,-4.9761270105372294e-03,
    -1.0207539277238262e-02,-4.9923364468588922e-02,-2.1425462100839177e-02,
    -7.6389181920920183e-03,-2.2107826969034853e-02,-5.5175592443221086e-03,
    2.7402432269926273e-02, 1.6479428281597576e-02,-5.4329898568029401e-04,
    1.2183933474128661e-02,-4.3280531243350406e-03, 1.0602749098786254e-01,
    4.0128397827780696e-03, 9.2848833258419115e-03,-2.4613176198757528e-02,
    -2.4037334282228290e-02,-5.9876740977274398e-02, 2.1551781525034834e-02,
    -1.5609621864944849e-02, 5.3774376706539610e-02, 2.1863060121364930e-03,
    -8.7191642562251190e-02,-5.1747895291181040e-02,-3.7239969486428533e-03,
    -8.2231025099897048e-02,-1.2088423805397772e-02, 5.0568832799796209e-02,
    4.3439598244909415e-02, 6.5245453947046039e-04, 5.9441556281580786e-02,
    1.0179736824624509e-02, 3.4610302584949001e-02, 3.1756027236521361e-03,
    3.4581996335759611e-02,-1.6586402168475813e-02,-5.1091630535929624e-03,
    -7.7016155520247986e-02,-3.6310068850195796e-04,-1.0146050073386733e-02,
    -4.9657715596630334e-02,-3.4480618074316231e-02,-7.0512634585505343e-02,
    1.0444781268045485e-01,-1.2761881055841825e-02, 4.7839354834689915e-02,
    -2.0159701467678949e-03, 3.8132066591278033e-02, 1.6018278133794386e-02,
    -6.8216185785896976e-04, 2.1880650414200151e-02,-2.1354211477652020e-02,
    -1.3013249542137970e-01, 8.0018571797155555e-05, 3.5772514434765450e-03,
    -6.2756603772158354e-04, 6.0294547532934832e-03,-6.2266735954623863e-02,
    3.5498766526103143e-03, 1.6905481648381820e-02, 4.9273577159003906e-02,
    5.7824919862743331e-02, 1.2708953922286503e-01, 3.7543913257366612e-02,
    2.4243645463124013e-02,-1.0648893312728998e-01, 2.1371258890977293e-02,
    -6.4477842532674370e-02,-3.6047303344615998e-02,-1.0940878466621628e-03,
    2.1855103247642713e-02, 3.1790924091685858e-02,-2.2343787182364434e-03,
    -1.2636605967299571e-03, 2.8808968596202134e-02,-1.3523997513060375e-02,
    1.4315577673815762e-03,-4.4157092566691002e-02, 5.9440163004523851e-03,
    -1.6562715031051087e-03, 9.5776455651077397e-02,-2.7828922856678019e-02,
    8.6145214353677176e-03,-5.1623358621970799e-03,-6.6346672579203122e-03,
    8.6575397291795683e-02,-3.1393957301753222e-03, 2.0358860403482150e-02,
    -2.2456609104123857e-03, 3.4263259980758223e-04,-1.1603755948364916e-01,
    -2.3968467046711416e-02, 7.1066901202629124e-02,-1.4271703625776857e-03,
    -9.9874046051312587e-02, 2.2764176855852036e-02, 2.3701497172434155e-03,
    -4.4598512665113077e-03,-4.0016464019557163e-03, 2.9684599054359633e-03,
    3.3307425403905112e-02, 8.1052148456567497e-02,-1.5990964460323667e-01,
    3.3941971870815116e-02,-9.1713307324679597e-03, 3.9706406317883892e-02,
    1.3214302707161263e-02,-8.9310890002969046e-02,-1.4207647108859564e-02,
    1.8513418110384045e-03,-8.4501824968164357e-03, 8.8569124778174815e-02,
    -1.0384303053719209e-01, 1.4945299803994084e-03, 9.7503378661016590e-02,
    -1.9189279950865397e-02,-8.8977049096963074e-04, 4.0169407532188745e-03,
    2.4213652886624099e-03, 2.3717360736080816e-03,-2.2079482368213013e-02,
    9.4232077079897641e-02,-1.8703447035227672e-01,-2.5524526429220588e-02,
    -6.7580013165596982e-03, 3.2074780712461753e-02,-1.0396598316003496e-03,
    2.7482863507617403e-02, 1.8416877078642240e-02, 1.0343936515667725e-03,
    -7.0757720582876752e-03, 9.2385050548094358e-02,-1.1453232083409967e-01,
    1.8977185196520364e-04,-1.0471639046860522e-01, 1.2198886143628882e-02,
    -3.0428410454664246e-03,-1.6100636443911209e-03,-2.2204838200661783e-03,
    -1.4572816466059981e-03, 1.1414846414323448e-02, 4.9141967181568873e-03,
    -5.4543157928328222e-02, 8.9976551629593875e-03, 9.8445554726762832e-03,
    -9.3495446201879583e-04,-1.0708150645770305e-02, 1.7284917890894769e-01,
    8.2938928021369790e-02,-8.9304452131118471e-04, 4.8551542726285086e-04,
    -9.1748451046270180e-03, 3.0336879081727046e-02,-3.3752782498249785e-04,
    -4.4059345017784237e-03, 1.0240344160820778e-04,-1.0598757545741607e-05,
    1.9664660976771850e-03,-9.0624253038513612e-04,-4.6743856534761461e-03,
    -6.3660743309883449e-05,-9.7255099816571811e-03, 8.8990614582936431e-02,
    3.9394654110374815e-03,-9.6045316425721408e-02,-4.8558234849316005e-03,
    3.3181794752759576e-04, 1.1918289922406770e-01,-1.9617451101509251e-01,
    -5.3603847646895020e-04, 6.0184630165408990e-04, 7.2283157307114085e-02,
    3.0915198312213075e-03, 8.9828113085556355e-06, 2.2151533200917461e-04,
    -9.0083112991316849e-05,-4.6989452619435236e-04, 4.5042205022524087e-04,
    1.3410987231912623e-04,-2.5965099740437480e-05,-2.7861981259965290e-04,
    1.8804011837507170e-01, 2.1584987967896159e-02, 6.8567480468941584e-04,
    -1.4508856573559828e-01,-2.3913452365155012e-03,-1.0532936244651600e-03,
    -1.0665377888514476e-02, 3.5527843983730517e-02, 3.0612878384504076e-04,
    1.6864723527011745e-04,-9.0156806010967053e-02,-2.0190518755886382e-02,
    -5.8106231531653369e-05,-9.4389850367775747e-04, 3.3245766282020236e-05,
    -2.4816427980418719e-04,-1.2394776729695335e-03,-4.1111871090684719e-05,
    1.6012017637935276e-03,-4.9472824145910672e-06,-1.2418938572281396e-01,
    8.9794919118293196e-02,-2.2835875595985760e-04,-1.3169253693631700e-01,
    9.8376612991560812e-05,-2.4745373526791421e-03,-4.4029462774714243e-02,
    1.5300915574941021e-01,-3.4711885975447832e-06, 2.8490565994530799e-04,
    8.0522172657739963e-02
  };
/*----------------------------------------------------------------------------*/
double mtREV24eigmat[20] =
  {
    0.0,-1.3445101019750322e-01,-1.9640582075338989e-01,
    -2.1859267849476735e-01,-2.6734649499091523e-01,-4.1814449164662365e-01,
    -4.5354323573511868e-01,-5.4376979448246754e-01,-5.8943563448558001e-01,
    -6.5770168183810263e-01,-7.4644979931723210e-01,-7.6174493673799415e-01,
    -8.6283905007117379e-01,-9.6276523514824786e-01,-1.1479972243712393e+00,
    -1.2134870690943689e+00,-1.5089420026765514e+00,-1.9135822015049038e+00,
    -2.2822225988851370e+00,-2.2983211626674187e+00
  };
/*------------------------------------------------------------------------*/
double lgprobmat[400] =
  {
    7.9065960467029794e-02,5.5940972029520968e-02,4.1976979011515751e-02,
    5.3051973474019826e-02,1.2936993531504846e-02,4.0766979616515264e-02,
    7.1585964207026773e-02,5.7336971331521512e-02,2.2354988822508373e-02,
    6.2156968921523233e-02,9.9080950459537159e-02,6.4599967700024241e-02,
    2.2950988524508609e-02,4.2301978849015841e-02,4.4039977980016479e-02,
    6.1196969401522971e-02,5.3286973356519933e-02,1.2065993967004524e-02,
    3.4154982922512789e-02,6.9146965426525986e-02,-1.9743224774156552e-02,
    -4.1235252562365945e-02,-3.3496747119987749e-02,-6.2892059201797426e-02,
    3.8731320965339415e-03,-2.7543007703670509e-02,-7.6258019828254725e-02,
    -5.7368794671777856e-02,-5.3800277513103220e-03,7.7601572397712454e-02,
    1.3690930635304319e-01,-5.3546553429465876e-02,2.3430119605645262e-02,
    7.1550613600477653e-02,-3.5625636540491348e-02,-3.0046711214500164e-02,
    -9.6217117037292934e-03,2.5479000507884191e-02,4.6700480485725768e-02,
    6.7213521454485309e-02,2.6504276628120039e-02,-2.6439951483312348e-02,
    -1.3807247342127406e-02,-2.3644047102187635e-02,2.7688546587510956e-03,
    -1.1850508234544262e-02,-2.6805923516874122e-02,-5.7137652919102399e-03,
    -1.7488171499646061e-02,6.0468097251606054e-02,7.3761580986359362e-02,
    -2.4286210017683753e-02,1.4059178616348485e-02,-3.7383710777615799e-02,
    6.6153484963771117e-02,6.3610653317193490e-03,1.5194039068913407e-02,
    -7.6897008794620955e-02,-6.3794765097258746e-02,6.2840731652192283e-02,
    -1.3924798929662594e-03,2.2277504319773021e-02,1.4238449496380388e-02,
    2.4866757319979695e-02,1.4091222028743120e-03,1.2297207373985385e-02,
    3.1946366192707007e-02,2.1224409913014407e-02,1.8874098618876986e-03,
    2.4685238469787551e-02,3.1655374885580713e-02,2.7244998847681882e-02,
    7.3898289069546487e-03,-9.1227581256849918e-03,-1.8701939727078809e-01,
    -1.1350151232212111e-03,6.1712183960446771e-03,-3.5252796557877643e-02,
    -1.5938741115001074e-02,2.2567301898887795e-02,-5.0607384730302074e-02,
    6.6965132373474498e-02,2.6087712901631602e-03,1.8629569513318806e-02,
    -8.6467387457103695e-03,3.1864734721956993e-02,5.1773311550857119e-02,
    -1.9539682047243975e-01,1.4470832593984143e-02,-5.0651408259902604e-04,
    1.2214216746626502e-02,7.1233729160128198e-02,2.3282320953969681e-03,
    9.3707091100948929e-03,1.7616958153928983e-02,-2.5257337704169266e-02,
    -6.9404237778183091e-03,-1.8680139258307346e-02,1.5152533678480387e-02,
    -8.1933722170642164e-03,1.0334055890435737e-02,2.2464574514470176e-02,
    -8.0459621179691920e-03,-5.4906210364740480e-03,-1.5195271027692535e-03,
    6.0401363738881129e-03,1.0262335444285732e-02,-3.6797272898241438e-02,
    -2.1606665751450472e-02,4.1983267089029039e-02,3.9786510441593696e-02,
    2.4552071257281857e-02,9.3721430999609903e-03,-8.6022049297201780e-02,
    -1.2845401733821734e-02,-2.7137515001761787e-04,1.3089444518316281e-02,
    6.1026909215920598e-02,-1.1136039522223506e-01,4.5047822464998159e-02,
    -3.3423158971180372e-03,-1.2785866274405969e-01,1.8814544266344779e-02,
    1.2634991683034563e-01,-3.0646529292579629e-03,-1.0037760431385144e-02,
    1.1662705399002882e-01,-4.4878518627956339e-02,-9.3799660148480395e-03,
    1.1086416066472222e-02,1.2690678240714624e-02,-1.0069230855233753e-01,
    5.1515903801084399e-04,4.0220030434445784e-03,-2.2734652667434686e-04,
    -1.1086000553335532e-03,-2.2568947509838616e-03,5.9107974603170603e-03,
    -3.7824173539668629e-03,1.0612874948243303e-02,-1.0238493146402072e-01,
    2.8115232265673012e-02,-2.0870317589153468e-02,2.6666213963477230e-02,
    -5.7331530940844276e-02,1.0147077538497260e-02,4.8995599993771753e-02,
    8.3833793682422608e-02,-7.4704177977297989e-03,1.2430959839507953e-02,
    1.0218461053252755e-01,2.7576212175343714e-02,1.1061865604850112e-02,
    1.9629115665778189e-02,2.6026497645435048e-02,-8.6016566366654737e-02,
    -8.2767596546007599e-02,-7.4139290453363713e-04,-1.6466140848131122e-02,
    -2.2618284450208494e-02,-3.6940702059647779e-02,2.0774590010512517e-02,
    -4.1569695644095718e-02,6.7906281059486647e-03,9.2312795900265560e-02,
    1.8860436804987015e-03,5.6923069941605446e-02,1.9667608528084694e-02,
    -2.6440872200438673e-02,-2.1978854615431587e-02,4.0324261052825050e-02,
    1.4315546065513354e-02,1.7145647588164498e-03,2.8615424058016564e-02,
    9.8976902750234335e-03,-4.7309364083796344e-02,-6.3492890028298762e-02,
    -1.4420890489906465e-03,-2.3050102757108143e-02,-3.0997651939302689e-02,
    8.5369641897319881e-02,-1.0344850877214385e-02,-9.5442921130530350e-02,
    -7.1498315367308288e-02,-2.2946474391185009e-02,1.6036749911130511e-02,
    1.2486956963527344e-01,1.2039796572661696e-05,-6.1832529704366285e-02,
    1.0406947136678604e-02,-8.4923011498886272e-02,2.3111294346449776e-02,
    -9.9662410326114212e-03,4.0286542468940222e-02,-7.3858329110240949e-03,
    2.6677637543966030e-04,1.4792590332698518e-02,-1.1141455332939535e-03,
    1.1264402303267225e-02,3.9037768242649193e-02,3.5505049368858906e-02,
    -5.4089551651873003e-04,2.3596323778714755e-02,1.1726185115758698e-02,
    -1.4402422444254385e-02,-1.5846217302207748e-02,-3.4642719342232522e-02,
    -2.2457888742201840e-02,-4.4200450497823375e-02,-7.9318292084089401e-02,
    8.3267718521137693e-02,1.3417205420190703e-02,1.0840331839991488e-02,
    1.0481026619859968e-01,-7.5522760910446789e-03,6.1417288671761265e-02,
    4.6172185246599280e-02,-1.0030731537077158e-03,-7.0710502975695019e-02,
    -1.0007781601183727e-01,-3.5748509014860146e-02,5.2177993011422154e-02,
    2.3797848130597526e-02,9.4856941342795945e-02,4.2286805213227475e-04,
    -7.8287286438624315e-02,-6.9602061460999246e-02,2.0045666959955261e-03,
    -6.3071429238677895e-02,5.8827130821609917e-02,-1.1084291723041498e-01,
    3.9561145230447514e-02,-1.8662182283830858e-02,4.1780174528076343e-02,
    6.9759696579535955e-03,-1.7738120201316432e-02,-5.8365632853970041e-03,
    -2.2335858108895961e-04,1.2707663187208208e-03,7.8337023945458384e-02,
    -3.0711017761146795e-02,-1.9429206132990762e-02,2.2414649038326611e-02,
    -3.1213731027018361e-02,2.0606443143171864e-03,4.5827524358180341e-02,
    6.8222725128459262e-03,6.1228926693107808e-03,6.7356430841227138e-02,
    3.7842501224517759e-02,-1.1089073986242590e-01,-1.9667151431996866e-02,
    -1.6171456077779634e-02,1.0376346661705753e-01,2.1599415941928558e-03,
    -2.1528200579094138e-02,-1.5063884083750017e-02,8.8589796344873449e-04,
    -8.6432367146452430e-02,5.5851532969229851e-02,-1.3677226585164581e-01,
    -1.1144594129379914e-01,6.5155493669261197e-02,-6.0088216909408784e-02,
    9.1911453315601330e-03,2.4914452898557250e-02,2.1245542441495426e-02,
    1.6942520552400368e-02,-3.5712914056423808e-02,1.7713176328241223e-02,
    -1.1351072659671366e-02,1.1023895218597228e-01,4.7601461180599037e-03,
    -7.8362517597880925e-03,5.9363371721151406e-03,6.3960746227408700e-03,
    6.4337547373787066e-02,7.7442644277571945e-04,1.4240787762115573e-02,
    1.3600596316548602e-03,-1.4146261223625758e-01,9.1692834867631787e-02,
    -3.3531434155057202e-02,-9.1479143326067842e-03,2.8325446037017568e-03,
    -4.6308370862400738e-04,4.3817371297379372e-02,1.8305875415739581e-02,
    -5.1992706297516941e-03,7.9450609237370351e-03,-8.6516101851676873e-03,
    -1.2091584404144733e-01,3.7172011335702103e-03,9.6549460085257116e-04,
    3.1548698650327874e-03,2.9202708524359657e-02,1.2435392860075287e-01,
    -7.3194151462194483e-04,5.9201628153075683e-04,-6.4761953107536716e-03,
    -1.4269537701996638e-02,-3.6571763866311738e-02,-7.8500394831457432e-02,
    1.6887632598201072e-02,5.5836695991927897e-04,-1.2771096947577978e-01,
    3.4340211812727152e-02,6.9688153774306767e-03,6.6247660295992689e-02,
    -9.0346538653875742e-03,1.6511999983580872e-02,9.1014928368926884e-02,
    -2.9473276027534279e-03,3.7120755679848150e-03,-4.5229744093102601e-05,
    -1.7938289918888208e-03,5.9458903280459982e-02,4.1260996116465072e-04,
    -1.5128554823731913e-02,-1.0110943302987636e-02,-1.4217855024277115e-02,
    -4.3670322437586555e-02,-1.0914866236520412e-01,9.3678867148642486e-02,
    1.6753244231121033e-03,1.0333043840654101e-01,-1.0746356877695834e-01,
    8.2144233576585679e-03,6.8113848783505954e-04,-5.7172767054854750e-03,
    -1.8927526539122906e-02,3.9595559258491404e-02,1.6282700474742548e-02,
    1.4899911760650364e-03,-1.3358370494686654e-03,-1.3397352453260873e-02,
    4.9717298163168493e-02,-1.0132583671049620e-05,1.2570882572579261e-03,
    -2.0342952184793598e-03,-1.6049285841559395e-02,-5.3721484625182392e-03,
    -1.5159297375746849e-02,5.1331298212743490e-03,-4.2741567653879482e-04,
    2.0878714422447590e-02,-9.5155170929635147e-03,-1.9919796689139361e-04,
    -1.5876116660969337e-03,5.7549171176067794e-03,9.5182544456575743e-02,
    6.7229728814970739e-03,-1.4018788545377800e-01,-1.7119561571186167e-03,
    -2.0338802637462688e-03,3.8777807341765495e-02,3.7578434067920793e-04,
    2.7171172890092757e-04,1.7436376182556825e-03,1.7402976227955107e-02,
    7.3397548761988438e-02,6.9139580094934429e-03,4.1174696402484320e-02,
    -1.6428089882195474e-03,2.8513430186209126e-03,3.3821358870368681e-03,
    -2.3283455591082843e-03,5.2104268600209813e-03,-3.4245424589693209e-03,
    -4.7678515571643786e-02,2.6516320385331280e-02,-1.0426461692773748e-02,
    -2.0972481140015885e-02,1.5590147644764284e-03,3.4032600256674247e-03,
    -1.9567230365291013e-01,1.0188436403222668e-01,7.1814592684183863e-05,
    7.0002914938278047e-04,1.5080547174227162e-02,4.0686919810066551e-02,
    8.1037430420548707e-04,2.2356264663481945e-03,2.7283254694700232e-04,
    2.7621851367426583e-03,1.6560039350386069e-03,-2.8746136938476299e-04,
    -9.8183613038329686e-04,-5.5518112011235153e-05,1.7717917967046645e-01,
    -2.8111564830023320e-02,-1.1797655776843486e-03,-1.3220218846432447e-02,
    1.5517366789327764e-04,6.1981750500204164e-04,-2.8514397554629870e-02,
    1.9414603269827264e-02,2.4009317809974297e-04,1.9216177241647748e-04,
    -1.7387420884250429e-01
  };
/*------------------------------------------------------------------------*/
double lgeigmat[20] =
  {
    0.0,-2.6718046750031804e-01,-4.0553164685962512e-01,
    -4.3276501116235572e-01,-4.8767792703894219e-01,-5.4617220395411259e-01,
    -6.0825341318450166e-01,-7.5391442704973044e-01,-9.1682365683555578e-01,
    -9.8565522505531034e-01,-1.0048426435720317e+00,-1.0633771740982703e+00,
    -1.1683943738967237e+00,-1.2517432888728204e+00,-1.3370743802202905e+00,
    -1.4335179353250611e+00,-1.4587299196517403e+00,-1.6534242622834008e+00,
    -1.7991131391748842e+00,-2.1161585752491985e+00
  };
/*------------------------------------------------------------------------*/
double lgpaml[] =
  {
    0.425093,
0.276818,0.751878,
0.395144,0.123954,5.076149,
2.489084,0.534551,0.528768,0.062556,
0.969894,2.807908,1.695752,0.523386,0.084808,
1.038545,0.363970,0.541712,5.243870,0.003499,4.128591,
2.066040,0.390192,1.437645,0.844926,0.569265,0.267959,0.348847,
0.358858,2.426601,4.509238,0.927114,0.640543,4.813505,0.423881,0.311484,
0.149830,0.126991,0.191503,0.010690,0.320627,0.072854,0.044265,0.008705,0.108882,
0.395337,0.301848,0.068427,0.015076,0.594007,0.582457,0.069673,0.044261,0.366317,4.145067,
0.536518,6.326067,2.145078,0.282959,0.013266,3.234294,1.807177,0.296636,0.697264,0.159069,0.137500,
1.124035,0.484133,0.371004,0.025548,0.893680,1.672569,0.173735,0.139538,0.442472,4.273607,6.312358,0.656604,
0.253701,0.052722,0.089525,0.017416,1.105251,0.035855,0.018811,0.089586,0.682139,1.112727,2.592692,0.023918,1.798853,
1.177651,0.332533,0.161787,0.394456,0.075382,0.624294,0.419409,0.196961,0.508851,0.078281,0.249060,0.390322,0.099849,0.094464,
4.727182,0.858151,4.008358,1.240275,2.784478,1.223828,0.611973,1.739990,0.990012,0.064105,0.182287,0.748683,0.346960,0.361819,1.338132,
2.139501,0.578987,2.000679,0.425860,1.143480,1.080136,0.604545,0.129836,0.584262,1.033739,0.302936,1.136863,2.020366,0.165001,0.571468,6.472279,
0.180717,0.593607,0.045376,0.029890,0.670128,0.236199,0.077852,0.268491,0.597054,0.111660,0.619632,0.049906,0.696175,2.457121,0.095131,0.248862,0.140825,
0.218959,0.314440,0.612025,0.135107,1.165532,0.257336,0.120037,0.054679,5.306834,0.232523,0.299648,0.131932,0.481306,7.803902,0.089613,0.400547,0.245841,3.151815,
2.547870,0.170887,0.083688,0.037967,1.959291,0.210332,0.245034,0.076701,0.119013,10.649107,1.702745,0.185202,1.898718,0.654683,0.296501,0.098369,2.188158,0.189510,0.249313};
/*------------------------------------------------------------------------*/
double jttpaml[]=
  {
    58,                                                                        
 54,  45,                                                                    
 81,  16, 528,                                                                
 56, 113,  34,  10,                                                            
 57, 310,  86,  49,   9,                                                        
105,  29,  58, 767,   5, 323,                                                    
179, 137,  81, 130,  59,  26, 119,                                                
 27, 328, 391, 112,  69, 597,  26,  23,                                            
 36,  22,  47,  11,  17,   9,  12,   6,  16,                                        
 30,  38,  12,   7,  23,  72,   9,   6,  56, 229,                                    
 35, 646, 263,  26,   7, 292, 181,  27,  45,  21,  14,                                
 54,  44,  30,  15,  31,  43,  18,  14,  33, 479, 388,  65,                            
 15,   5,  10,   4,  78,   4,   5,   5,  40,  89, 248,   4,  43,                        
194,  74,  15,  15,  14, 164,  18,  24, 115,  10, 102,  21,  16,  17,                    
378, 101, 503,  59, 223,  53,  30, 201,  73,  40,  59,  47,  29,  92, 285,                
475,  64, 232,  38,  42,  51,  32,  33,  46, 245,  25, 103, 226,  12, 118, 477,            
  9, 126,   8,   4, 115,  18,  10,  55,   8,   9,  52,  10,  24,  53,   6,  35,  12,        
 11,  20,  70,  46, 209,  24,   7,   8, 573,  32,  24,   8,  18, 536,  10,  63,  21,  71,    
    298,  17,  16,  31,  62,  20,  45,  47,  11, 961, 180,  14, 323,  62,  23,  38, 112,  25,  16};
/*------------------------------------------------------------------------*/
double wagpaml[]=
  {
0.551571,
0.509848,0.635346,
0.738998,0.147304,5.429420,
1.027040,0.528191,0.265256,0.0302949,
0.908598,3.035500,1.543640,0.616783,0.0988179,
1.582850,0.439157,0.947198,6.174160,0.021352,5.469470,
1.416720,0.584665,1.125560,0.865584,0.306674,0.330052,0.567717,
0.316954,2.137150,3.956290,0.930676,0.248972,4.294110,0.570025,0.249410,
0.193335,0.186979,0.554236,0.039437,0.170135,0.113917,0.127395,0.0304501,0.138190,
0.397915,0.497671,0.131528,0.0848047,0.384287,0.869489,0.154263,0.0613037,0.499462,3.170970,
0.906265,5.351420,3.012010,0.479855,0.0740339,3.894900,2.584430,0.373558,0.890432,0.323832,0.257555,
0.893496,0.683162,0.198221,0.103754,0.390482,1.545260,0.315124,0.174100,0.404141,4.257460,4.854020,0.934276,
0.210494,0.102711,0.0961621,0.0467304,0.398020,0.0999208,0.0811339,0.049931,0.679371,1.059470,2.115170,0.088836,1.190630,
1.438550,0.679489,0.195081,0.423984,0.109404,0.933372,0.682355,0.243570,0.696198,0.0999288,0.415844,0.556896,0.171329,0.161444,
3.370790,1.224190,3.974230,1.071760,1.407660,1.028870,0.704939,1.341820,0.740169,0.319440,0.344739,0.967130,0.493905,0.545931,1.613280,
2.121110,0.554413,2.030060,0.374866,0.512984,0.857928,0.822765,0.225833,0.473307,1.458160,0.326622,1.386980,1.516120,0.171903,0.795384,4.378020,
0.113133,1.163920,0.0719167,0.129767,0.717070,0.215737,0.156557,0.336983,0.262569,0.212483,0.665309,0.137505,0.515706,1.529640,0.139405,0.523742,0.110864,
0.240735,0.381533,1.086000,0.325711,0.543833,0.227710,0.196303,0.103604,3.873440,0.420170,0.398618,0.133264,0.428437,6.454280,0.216046,0.786993,0.291148,2.485390,
2.006010,0.251849,0.196246,0.152335,1.002140,0.301281,0.588731,0.187247,0.118358,7.821300,1.800340,0.305434,2.058450,0.649892,0.314887,0.232739,1.388230,0.365369,0.314730};
/*------------------------------------------------------------------------*/
```
