## Supplementary material for "Modeling site-and-branch-heterogeneity with GFmix": alpha_est_mix_rt.c

```
#include "alpha_est_mix_rtf.h"
int main(int argc, char **argv)
{
  FILE *freqfile,*seqfile,*utreecfile,*wtfile,*treefile,*sitellfile;
  char (*names)[11],value[1000],opt;
  int no_opt,*seq,ntaxa,nsite,*locs,*count,npatt,nclass=0,i,nchar=20,fso;
  int nrate=4,smodel=9,optim=1,garp=0,wtopt=1,pr=0;
  double *utreec,S[400],alpha=1.0,lnl,*fr,*wtc,*lp,*liks;
  /* command line options */
  freqfile=NULL; seqfile=NULL; utreecfile=NULL; wtfile=NULL; treefile=NULL;
  sitellfile=NULL;
  argc--;
  while((opt=get_param(&argc,argv,value)) != 'z'){
    no_opt=1;
    if(opt=='f'){
      freqfile=fopene(value,"r","freqfile not available"); no_opt=0;
    }
    if(opt=='i'){
      seqfile=fopene(value,"r","seqfile not available"); no_opt=0;
    }
    if(opt=='u'){
      utreecfile=fopene(value,"r","utreecfile not available"); no_opt=0;
    }
    if(opt=='c'){ nclass=atoi(value); no_opt=0;}
    if(opt=='s'){ smodel=atoi(value); no_opt=0; }
    if(opt=='n'){ optim=0; no_opt=0; }
    if(opt=='w'){
      wtfile=fopene(value,"r","wtfile not available"); no_opt=0;
    }
    if(opt=='t'){
      treefile=fopene(value,"r","treefile not available"); no_opt=0;
    }
    if(opt=='a'){ alpha=atof(value); no_opt=0;}
    if(opt=='p'){ pr=1; no_opt=0; }
    if(opt=='l'){
      sitellfile=fopene(value,"wb","site log likelihood file not available");
      no_opt=0;
    }
    if(no_opt) {printf("Option %c not available\n",opt); exit(1);} 
  }
  if(freqfile==NULL){printf("freqfile needed: -f freqfile\n"); exit(1);}
  if(seqfile==NULL){printf("seqfile needed: -i seqfile\n"); exit(1);}
  if(utreecfile==NULL && treefile==NULL){
    printf("utreecfile needed: -u utreecfile\n"); exit(1);
  }
  if(utreecfile!=NULL && treefile!=NULL){
    printf("treefile or utreecfile should be specified, not both\n"); exit(1);
  }
  if(nclass==0){printf("number of classes >0 needed: -c nclass\n"); exit(1);}
  if(smodel!=5 && smodel!=9 && smodel !=6){
    printf("smodel=5 (JTT), 6 (WAG) or 9(LG)\n"); exit(1);}
  /* sequence data */
  seq=read_seq(seqfile,nchar,0,&ntaxa,&nsite,&names);
  /* for(i=0; i<nsite; i++) count[i] = 1.0; */
  locs=(int *) malloc((size_t) nsite*sizeof(int));
  count=(int *) malloc((size_t) nsite*sizeof(int));
  npatt=nsite;
  npatt = patt_freq_locs(ntaxa,nsite,seq,count,locs);
  /* frequencies & weights */
  fr=(double *) malloc((size_t) nchar*(2*ntaxa-1)*nclass*sizeof(double));
  wtc=(double *) malloc((size_t) nclass*sizeof(double));
  for(i=0; i<nchar*(2*ntaxa-1)*nclass; i++){
    fso=fscanf(freqfile,"%lf",&fr[i]);
    if(fso<=0){
      printf("freqfile does not have enough entries nclass=%i\n",nclass);
      exit(1);
    }}
  if(wtfile==NULL){
    for(i=0; i<nclass; i++) wtc[i] = 1.0/nclass;
  }
  else{
    wtopt=0;
    for(i=0; i<nclass; i++){
      fso=fscanf(wtfile,"%lf",&wtc[i]);
      if(fso<=0){ printf("Need %i entries in weightfile\n",nclass); exit(1);}
    }
    for(lnl=0.0, i=0; i<nclass; i++) lnl += wtc[i]; /* rnded wts eg. iqtree */
    for(i=0; i<nclass; i++) wtc[i] /= lnl;
  }
  /* tree */
  utreec=(double *) malloc((size_t) (ntaxa-1)*4*sizeof(double));
  if(utreecfile!=NULL){
    for(i=0; i<(ntaxa-1)*4; i++){
      fso=fscanf(utreecfile,"%lf",&utreec[i]);
      if(fso<=0){
	printf("utreecfile does not have enough entries ntaxa=%i\n",ntaxa);
	exit(1);
      }}
  }
  if(treefile!=NULL) read_treecsnl(treefile,ntaxa,names,utreec,1);
  /* echangeability matrix */
  if(smodel==9) LG_exchangeability(S);
  if(smodel==5) JTT_exchangeability(S);
  if(smodel==6) WAG_exchangeability(S);
  
  lp=(double *) malloc((size_t) (2*ntaxa-2+nclass+1)*sizeof(double));
  liks=(double *) malloc((size_t) nsite*sizeof(double));
  if(!optim){
    lnl=fmix_garp_lnld(nchar,ntaxa,npatt,seq,count,utreec,nclass,fr,wtc,S,
  		       nrate,alpha,lp,0,liks);
  }
  /* else{ */
  /*   lnl=fmix_garp_lnld_check(nchar,ntaxa,npatt,seq,count,utreec,nclass,fr,wtc, */
  /* 			     S,nrate,alpha,lp,liks); */
  /* } */
  printf("%.16e %.16e\n",lnl,alpha);
  for(i=0; i<nclass; i++) printf("%.16e ",wtc[i]); printf("\n");
  if(sitellfile!=NULL)
    for(i=0; i<nsite; i++) fwrite(&liks[locs[i]],sizeof(double),1,sitellfile);
  pr_utreec(stdout,ntaxa,utreec);
}
```
