## Supplementary material for "Modeling site-and-branch-heterogeneity with GFmix": alpha_est_mix_rtf.c

```
#include "alpha_est_mix_rtf.h"
#include "aa_empirical.h"
void Mmake_rt_mix(int nchar, int ntaxa, double *utreec,
		  int nclass, double *fr, double *S,
		  int nrate, double *rated, 
		  double *M, double *Mp, int deriv){
  int ic,idx,idf,nb,r,j,k,ir,nc2,l;
  double W[400],Q[400],Lam[20];
  
  nb=2*ntaxa-2;
  nc2=nchar*nchar;
  for(ic=0; ic<nclass; ic++){
    for(j=0; j<ntaxa-1; j++)
      for(k=0; k<2; k++){
	r=(int) utreec[k+j*4];
	idf=r*nchar+ic*nchar*(nb+1);
	GetWLam(nchar,S,&fr[idf],Q,W,Lam);
	idx=ic*nchar*nchar*nb*nrate;
	for(ir = 0; ir < nrate; ir++){
	  idx=r*nc2+ir*nc2*nb+ic*nc2*nb*nrate;
	  if(deriv>0){
	    Mijdp(nchar,rated[ir]*utreec[k+2+j*4],&fr[idf],W,Lam,&M[idx],&Mp[idx]);
	    for(l=0; l<nc2; l++) Mp[l+idx] *= rated[ir];
	  }
	  else
	    Mij_nod(nchar,rated[ir]*utreec[k+2+j*4],&fr[idf],W,Lam,&M[idx]);
	}
      }}
}
double fmix_garp_lnld(int nchar, int ntaxa, int npatt, int *seq, int *count,
		      double *utreec,
		      int nclass, double *fr, double *wtc,
		      double *S, int nrate, double alpha, double *lp,
		      int deriv, double *liks){
  int nb,ic,idx,i,*s,j,nsite,np,nc2,lderiv,safe=1,nd;
  double *wt,*rated,*M,*Mp,lik,*likc,*llikc,likcm,*lps,*llps,lnl,lpc,likm;

  wt=(double *) malloc((size_t) nrate*sizeof(double));
  if(deriv>=2){
    rated=(double *) malloc((size_t) nrate*2*sizeof(double));
    /* DiscreteGammad(nrate,alpha,wt,rated); /\* rates and derivs *\/ */
    DiscreteGammada(nrate,alpha,wt,rated,1.0e-6); /* rates and derivs */
  }
  else{
    rated=(double *) malloc((size_t) nrate*sizeof(double));
    DiscreteGamma(wt,rated,alpha,alpha,nrate,0);
    /* DiscreteGammad(nrate,alpha,wt,rated); /\* rates and derivs *\/ */
  }
  
  nb=2*ntaxa-2;
  nc2=nchar*nchar;
  if(deriv==0) lderiv=0; 
  if(deriv==1){lderiv=1; np=nb;}
  if(deriv==2){lderiv=2; np=nb+1;}
  if(deriv==3){lderiv=2; np=nb+1+nclass;}

  nsite=0;
  for(i=0; i<npatt; i++) nsite += count[i];
  
  if(deriv>0){
    lps=(double *) malloc((size_t) nclass*(nb+1)*sizeof(double));
    llps=(double *) malloc((size_t) nclass*(nb+1)*sizeof(double));
    for(j=0; j<np; j++) lp[j]=0.0;
    Mp=(double *) malloc((size_t) nclass*nc2*nb*nrate*sizeof(double));
  }

  M=(double *) malloc((size_t) nclass*nc2*nb*nrate*sizeof(double));
  Mmake_rt_mix(nchar,ntaxa,utreec,nclass,fr,S,nrate,rated,M,Mp,deriv);

  s=(int *) malloc((size_t) ntaxa*sizeof(int));
  likc=(double *) malloc((size_t) nclass*sizeof(double));
  llikc=(double *) malloc((size_t) nclass*sizeof(double));
  lnl=0.0;
    
  for(i=0; i<npatt; i++){

    /* likelikelihoods and derivatives (log for safe version) for each class */
    for(j = 0; j < ntaxa; j++) s[j] = seq[i+j*npatt];
    for(ic=0; ic<nclass; ic++){
      idx=ic*nc2*nb*nrate;
      j=nb*nchar+ic*nchar*(nb+1);
      /* if(!safe){ */
      /* 	likc[ic]=lik_deriv_site_rt(nchar,ntaxa,s,&fr[j],utreec, */
      /* 				   nrate,rated,wt,&M[idx],&Mp[idx], */
      /* 				   &lps[ic*(nb+1)],lderiv,1); */
      /* }else{ */
      likc[ic]=lik_deriv_site_rt(nchar,ntaxa,s,&fr[j],utreec,
				 nrate,rated,wt,&M[idx],&Mp[idx],&llikc[ic],
				 &lps[ic*(nb+1)],&llps[ic*(nb+1)],deriv,1);
      /* } */
      /* printf("%.5e %.5e\n",lps[ic*(nb+1)]*exp(likc[ic]),exp(likc[ic])); */
    }

    /* update likelihood and derivative calculations */
    if(safe){
      likm=llikc[0];
      for(ic=1; ic<nclass; ic++) if(llikc[ic]>likm) likm=llikc[ic];
      for(lik=0.0, ic=0; ic<nclass; ic++){
	lik += wtc[ic]*likc[ic]*exp(llikc[ic]-likm);
      }
      /* printf("%i %.5e %.5e\n",i,lik,likm); */
      liks[i] = likm+log(lik);
      lnl += count[i]*liks[i];
      if(deriv>0){
	nd=(deriv==1)?nb:(nb+1);
	for(j=0; j<nd; j++){
	  for(lpc=0.0, ic=0; ic<nclass; ic++)
	    if(lps[j+ic*(nb+1)] != 0.0)
	      lpc += wtc[ic]*lps[j+ic*(nb+1)]*exp(llps[j+ic*(nb+1)]-liks[i]);
	  lp[j] += count[i]*lpc;
	}}
      if(deriv==3) /* weight contribution */
	for(ic=0; ic<nclass; ic++)
	  lp[nb+1+ic] += count[i]*likc[ic]*exp(llikc[ic]-liks[i]); 
    }
    else{
      for(lik=0.0, ic=0; ic<nclass; ic++) lik += wtc[ic]*likc[ic];
      liks[i] = log(lik);
      lnl += count[i]*liks[i];
      if(deriv>0){
	nd=(deriv==1)?nb:(nb+1);
	for(j=0; j<nd; j++){
	  for(lpc=0.0, ic=0; ic<nclass; ic++) lpc += wtc[ic]*lps[j+ic*(nb+1)];
	  lp[j] += count[i]*lpc/lik;
	}}
      if(deriv==3) /* weight contribution */
	for(ic=0; ic<nclass; ic++) lp[nb+1+ic] += count[i]*likc[ic]/lik; 
    }
    
  }
  /* for(i=0; i<nb; i++) printf("%.16e\n",lp[i]); printf("\n"); */

  /* Lagrange: lnl - n(sum_j wtc[j] - 1) is quantity optimized */
  lnl += nsite;
  for(ic=0; ic<nclass; ic++){
    lnl -= nsite*wtc[ic];
    if(deriv==3) lp[nb+1+ic] -= nsite;
  }

  free(wt); free(rated); free(M); free(s); free(likc); free(llikc);
  if(deriv>0){
    free(Mp); free(lps); free(llps);
  }
  return(lnl);
}
void fmix_garp_param2x(int ntaxa, int nclass, double *utreec, double *wtc,
		       double alpha, double *x){
  int nb,np,j;

  nb=2*ntaxa-2;
  np=nb+nclass+1;
  for(j=0; j<ntaxa-1; j++){ /* rooted tree */
    x[(int) utreec[j*4]]=utreec[2+j*4];
    x[(int) utreec[1+j*4]]=utreec[3+j*4];
  }
  x[nb]=alpha;
  for(j=0; j<nclass; j++) x[nb+1+j]=wtc[j];
}
void fmix_garp_x2param(int ntaxa, int nclass, double *x, double *utreec,
		       double *wtc, double *alpha){
  int nb,np,j;

  nb=2*ntaxa-2;
  np=nb+nclass+1;
  for(j=0; j<ntaxa-1; j++){ /* rooted tree */
    utreec[2+j*4]=x[(int) utreec[j*4]];
    utreec[3+j*4]=x[(int) utreec[1+j*4]];
  }
  *alpha=x[nb];
  for(j=0; j<nclass; j++) wtc[j]=x[nb+1+j];
}
FILE *fopene(char *filename, char *rw, char *err){
  FILE *fp;
  if((fp=fopen(filename,rw))==NULL){
    printf("%s\n",err);
    exit(1);
  }
  return(fp);
}
char get_param(int *k, char **arg, char *value){ 
  char opt='z';

  *value='\0';
  if(*k==0) return('z');
  if(arg[(*k)][0]!='-'){ /* not an option, rather a value */
    sprintf(value,"%s",arg[(*k)]);
    *k = (*k)-1;
    if(arg[(*k)][0]=='-') opt=arg[(*k)][1];
  }
  else{ /* is an option */
    opt=arg[(*k)][1];
    if(arg[(*k)][1]!='\0' && arg[(*k)][2]!='\0') sprintf(value,"%s",&arg[(*k)][2]);
  }
  *k = (*k)-1;
  return(opt);
}
char ctl_paraml(FILE *fp, char *opt, char *value, int *is_option){
  /* return last char on line */
  char c;
  int i;

  *is_option=0;
  while(((c = getc(fp)) == ' ' || c == '\t' || c=='\r') && c != EOF && c != '\n');
  if(c == EOF || c == '\n') return(c);
  if(c == '*'){ /* comment */
    while((c = getc(fp)) != '\n' && c != EOF) ;
    return(c);
  } 

  /* option name */
  *is_option=1;
  for(i=0; c !=' ' && c != '\t' && c != '\r'; i++){
    opt[i]=c;
    c=getc(fp);
  }
  opt[i] = '\0';
  /* skip '=' */
  while((c=getc(fp))==' ' || c=='\t' || c == '=') ;
  for(i=0; c != ' ' && c != '\t' && c != '\r' && c != '\n' && c != EOF; i++){
    value[i]=c;
    c=getc(fp);
  }
  value[i]= '\0';
  while(c != '\n' && c != EOF) c=getc(fp);
  return(c);
}
int pmodel2smodel(int model, char *aaRmat, int nchar){
  if (nchar==4){
    if(model==0) return(0); /* JC */
    if(model==2) return(1); /* F81 */
    if(model==3) return(2); /* F84 */
    if(model==4) return(8); /* HKY */
    if(model==7) return(3); /* GTR */
  }
  if (nchar != 4){
    if(model==0) return(0); /* JC */
    if(model==1) return(1); /* F81 */
    if(model==8) return(3); /* GTR */
  }
  if (*aaRmat!='\0'){
    if(strcmp(aaRmat,"dayhoff.dat")==0) return(4); /* PAM */
    if(strcmp(aaRmat,"jones.dat")==0) return(5); /* JTT */
    if(strcmp(aaRmat,"wag.dat")==0) return(6); /* WAG */
    if(strcmp(aaRmat,"mtREV24.dat")==0) return(7); /* mtREV24 */
    if(strcmp(aaRmat,"lg.dat")==0) return(9); /* LG */
  }
  printf("No substitution model available\n");
  exit(1);
}
int std_paramf(FILE *ctlfile, FILE **treefile, FILE **utreecfile, FILE **seqfile,
	   FILE **Qfile, FILE **ratefile, char *aaRmat, 
	   int *model, int *nchar, int *nsite, int *ntaxa, int *nrate, 
	   int *ntrees, double *alpha, double *kappa){
  char c,opt[100],value[100];
  int is_option,done=0;
  *treefile=NULL;
  *utreecfile=NULL;
  *seqfile=NULL;
  *Qfile=NULL;
  *ratefile=NULL;
  *aaRmat='\0';
  *model=0;
  *nchar=4;
  *nsite=0;
  *ntaxa=0;
  *ntaxa=0;
  *nrate=4;
  *ntrees=1;
  *alpha=-1;
  *kappa=-1; 
  while(!done){
    c = ctl_paraml(ctlfile,opt,value,&is_option);
    if(is_option){
      if(strcmp(opt,"treefile")==0)
	*treefile=fopene(value,"r","treefile not available");
      if(strcmp(opt,"utreecfile")==0)
	*treefile=fopene(value,"r","utreecfile not available");
      if(strcmp(opt,"seqfile")==0) 
	*seqfile=fopene(value,"r","seqfile not available");
      if(strcmp(opt,"Qfile")==0) 
	*Qfile=fopene(value,"r","Qfile not available");
      if(strcmp(opt,"ratefile")==0)
	*ratefile=fopene(value,"r","ratefile not available");
      if(strcmp(opt,"aaRatefile")==0) sprintf(aaRmat,"%s",value);
      if(strcmp(opt,"model")==0) *model=atoi(value);
      if(strcmp(opt,"nchar")==0) *nchar=atoi(value);
      if(strcmp(opt,"nsite")==0) *nsite=atoi(value);
      if(strcmp(opt,"ntaxa")==0) *ntaxa=atoi(value);
      if(strcmp(opt,"nrate")==0) *nrate=atoi(value);
      if(strcmp(opt,"ncatG")==0) *nrate=atoi(value);
      if(strcmp(opt,"ntrees")==0) *ntrees=atoi(value);
      if(strcmp(opt,"alpha")==0) *alpha=atof(value);
      if(strcmp(opt,"kappa")==0) *kappa=atof(value);
      if(strcmp(opt,"ttratio")==0) *kappa=-atof(value);
      
    }
    if(c==EOF) done=1;
  }
  return(0);
}
void rmgap(int numsp, int *nsite, char *seqc){
  int i,j,k,sp,newnsite;

  newnsite = *nsite;
  for(i = *nsite-1; i >= 0; i--)
    for(j = 0; j < numsp; j++)
      if (seqc[i + j*(*nsite)] == '-'){
	for(k = i; k < newnsite-1; k++)
	  for(sp = 0; sp < numsp; sp++)
	    seqc[k + sp*(*nsite)] = seqc[k+1 + sp*(*nsite)];
	newnsite--;
	break;
      }

  for(j = 1; j < numsp; j++)
    for(i = 0; i < newnsite; i++)
      seqc[i + j*newnsite] = seqc[i + j*(*nsite)];

  *nsite = newnsite;
}
void letter(int numsp, int nsite, char *seq)
{
  int i, j;
  for(i = 1; i < numsp; i++)
    for(j = 0; j < nsite; j++)
      if (seq[j + i*nsite] == '.') seq[j + i*nsite]=seq[j];
}
int l2i(char c){
  if (c == 'a' || c == 'A') return(0);
  if (c == 'c' || c == 'C') return(1);
  if (c == 'g' || c == 'G') return(2);
  if (c == 't' || c == 'T') return(3);
  if (c == '-') return(5);
  return(6);
}
char i2l(int i){
  switch(i){
  case 0:
    return('A');
  case 1:
    return('C');
  case 2:
    return('G');
  case 3:
    return('T');
  }
  return('E');
}
char i2lp(int a) {
  switch (a) {
  case alanine:        return('A'); /* 0 */
  case arginine:       return('R'); /* 1 */
  case asparagine:     return('N'); /* 2 */
  case aspartic:       return('D'); /* 3 */
  case cysteine:       return('C'); /* 4 */
  case glutamine:      return('Q'); /* 5 */
  case glutamic:       return('E'); /* 6 */
  case glycine:        return('G'); /* 7 */
  case histidine:      return('H'); /* 8 */
  case isoleucine:     return('I'); /* 9 */
  case leucine:        return('L'); /* 10 */
  case lysine:         return('K'); /* 11 */
  case methionine:     return('M'); /* 12 */
  case phenylalanine:  return('F'); /* 13 */
  case proline:        return('P'); /* 14 */
  case serine:         return('S'); /* 15 */
  case threonine:      return('T'); /* 16 */
  case tryptophan:     return('W'); /* 17 */
  case tyrosine:       return('Y'); /* 18 */
  case valine:         return('V'); /* 19 */
  case 21:             return('-');
  case 22:             return('X');
  case 23:             return('?');
  }

  return('?');
}
int l2ip(char c) {
  switch (c) {
  case 'A': return (alanine);
  case 'R': return (arginine);
  case 'N': return (asparagine);
  case 'D': return (aspartic);
  case 'C': return (cysteine);
  case 'Q': return (glutamine);
  case 'E': return (glutamic);
  case 'G': return (glycine);
  case 'H': return (histidine);
  case 'I': return (isoleucine);
  case 'L': return (leucine);
  case 'K': return (lysine);
  case 'M': return (methionine);
  case 'F': return (phenylalanine);
  case 'P': return (proline);
  case 'S': return (serine);
  case 'T': return (threonine);
  case 'W': return (tryptophan);
  case 'Y': return (tyrosine);
  case 'V': return (valine);
  case '-': return (21);
  case 'X': return (22);
  case '?': return (23);
  }

  return (100);
}
int rseqf(FILE *fp, char *seq){
  int rsite = 0;
  char c;
  
  while((c = getc(fp)) != '\n' &&  c != EOF){
    if (c != ' ' && c != '\r' && c!= '\t'){
      *seq = c;
      seq++;
      rsite++;
    }
  }

  return(rsite);
}
char ignore_whitespace_seqfile(FILE *infile){
  char c;
  while((c = getc(infile)) == ' ' || c == '\n' || c == '\r' || c == '\t') ;
  if(c == EOF){
    printf("seqfile: EOF before entire sequence read");
    exit(1);
  }
  return(c);
}
int rinterleave_block(FILE *infile, int numsp, int nsite, char name[][11], 
		       char *seq, int *initb){
  char c;
  int i,j,lsite,lsiteo;
  
  /* printf("%i\n",numsp); */
  for(i = 0; i < numsp; i++){
    c = ignore_whitespace_seqfile(infile);
    if((*initb) == 1){
      name[i][0] = c;
      for(j = 1; j < 10; j++)name[i][j] = getc(infile);
      name[i][10] = '\0';
      c = ignore_whitespace_seqfile(infile);      
      /* printf("%s\n",name[i]); */
    }
    seq[i*nsite] = c;
    lsite = rseqf(infile,&seq[1+i*nsite])+1;
    /* printf("%i\n",lsite); */
    if(i==0) lsiteo=lsite;

    if(i > 0 && lsite != lsiteo){
      printf("seqfile: number of character states should be same for all taxa in a block\n");
      exit(1);
    }
  }
  *initb = 0;
  
  return(lsite);
}
void rinterleavef_nolim(FILE *infile, const int *numsp, const int *nsite, 
			char name[][11], char *seq)
{
  int lsite = 0, csite = 0, initb = 1;

  while(csite < *nsite){
    lsite = rinterleave_block(infile,*numsp,*nsite,name,&seq[csite],&initb);
    csite += lsite;
  }
}
void rinterleave(int *numsp, int *nsite, char name[][11], char *seq)
{
  int fso;
  fso=scanf("%i %i", numsp, nsite);
  if(fso<=0){printf("Error reading sequence file\n"); exit(1);}
  rinterleavef_nolim(stdin,numsp,nsite,name,seq);
}
void rinterleavef(FILE *infile, int *numsp, int *nsite, char name[][11], 
		  char *seq)
{
  int fso;
  fso=fscanf(infile, "%i %i", numsp, nsite); 
  if(fso<=0){printf("Error reading sequence file\n"); exit(1);}
  rinterleavef_nolim(infile,numsp,nsite,name,seq);
}
int char_freq(int nchar, int numsp, int nsite, int *seq, double *freq)
{
  int i,j,nc=0;

  for(j = 0; j < nchar; j++){
    freq[j] = 0.0;
  }
  for(i = 0; i < nsite*numsp; i++){
    if(seq[i] < nchar){
      for(j = 0; j < nchar; j++){
	if (seq[i] == j) freq[j]++;
      }
      nc++;
    }
  }

  for(j = 0; j < nchar; j++){
    freq[j] /= nc;
  }
  return(nc);
}
int char_freq_count(int nchar, int numsp, int endsite, int *seq, int *count, 
		     double *freq)
{
  int i,j,nc=0,k;

  for(j = 0; j < nchar; j++){
    freq[j] = 0.0;
  }

  for(i = 0; i < endsite; i++){
    for(k = 0; k < numsp; k++)
      if(seq[i+k*endsite] < nchar)
	for(j = 0; j < nchar; j++)
	  if (seq[i+k*endsite] == j){
	    freq[j] += count[i];
	    nc += count[i];
	  }
  }

  for(j = 0; j < nchar; j++) freq[j] /= nc;
  return(nc);
}
int patt_freq(int numsp, int nsite, int *seq, int *count)
{
  int *s;
  int endsite,i,j,k,match;

  s = (int *) malloc((size_t) nsite*numsp*sizeof(int));
  endsite = 0;
  for(i = 0; i < nsite; i++){
    /* check for a match */
    match = 0;
    for(k = 0; k < endsite; k++){
      match = 1;
      for(j = 0; j < numsp; j++){
	if (seq[i+j*nsite] != s[j+k*numsp]){
	  match = 0;
	  break;
	}
      }
      if (match){
	count[k] += 1;
	break;
      }
    }
    
    /* if not a match, new element */
    if (!match){
      for(j = 0; j < numsp; j++) s[j+endsite*numsp] = seq[i+j*nsite];
      count[endsite] = 1;
      endsite++;
    }
  }
  

  for(j = 0; j < numsp; j++)
    for(i = 0; i < endsite; i++)
      seq[i+j*endsite] = s[j+i*numsp];

  free(s);
  return(endsite);
}
int patt_freq_locs(int numsp, int nsite, int *seq, int *count, int *locs)
{
  /* locs[i] gives the location of ith site in compressed seq matrix */
  int *s;
  int endsite,i,j,k,match;

  s = (int *) malloc((size_t) nsite*numsp*sizeof(int));
  for(i = 0; i < nsite; i++) count[i]=0;
  endsite = 0;
  for(i = 0; i < nsite; i++){
    /* check for a match */
    match = 0;
    for(k = 0; k < endsite; k++){
      match = 1;
      for(j = 0; j < numsp; j++){
	if (seq[i+j*nsite] != s[j+k*numsp]){
	  match = 0;
	  break;
	}
      }
      if (match){
	count[k] += 1;
	locs[i]=k;
	break;
      }
    }
    
    /* if not a match, new element */
    if (!match){
      for(j = 0; j < numsp; j++) s[j+endsite*numsp] = seq[i+j*nsite];
      count[endsite] = 1;
      locs[i]=endsite;
      endsite++;
    }
  }
  

  for(j = 0; j < numsp; j++)
    for(i = 0; i < endsite; i++)
      seq[i+j*endsite] = s[j+i*numsp];

  free(s);
  return(endsite);
}
void pr_utreec(FILE *fp, int ntaxa, double *utreec){
  int i;
  for(i = 0; i < ntaxa-1; i++) 
    fprintf(fp, "%i %i %.10f %.10f\n", (int) utreec[i*4], (int) utreec[1+i*4],
	   utreec[2+i*4], utreec[3+i*4]);
}
char *read_seqc(FILE *fp, int *ntaxa, int *nsite, char (**name)[11]){
  char *seqc;
  int fso;

  fso=fscanf(fp, "%i %i",ntaxa,nsite);
  if(fso<=0){printf("Error reading sequence file"); exit(1);}
  *name=(char (*)[11]) malloc((size_t) (*ntaxa)*sizeof(**name));
  seqc = (char *) malloc((size_t) (*ntaxa)*(*nsite)*sizeof(char));
  rinterleavef_nolim(fp,ntaxa,nsite,*name,seqc);
  return(seqc);
}
int *read_seq(FILE *fp, int nchar, int rm_gap, int *ntaxa, int *nsite, 
	      char (**name)[11]){
  int i,*seq;
  char *seqc;
  
  seqc = read_seqc(fp,ntaxa,nsite,name);
  letter(*ntaxa,*nsite,seqc);
  if (rm_gap) rmgap(*ntaxa,nsite,seqc);
  seq = (int *) malloc((size_t) (*ntaxa)*(*nsite)*sizeof(int));
  if(nchar==20)
    for(i = 0; i < (*nsite)*(*ntaxa); i++) seq[i] = l2ip(seqc[i]);
  else
    for(i = 0; i < (*nsite)*(*ntaxa); i++) seq[i] = l2i(seqc[i]);
  free(seqc);
  return(seq);
}
int DiscreteGamma (double freqK[], double rK[], double alpha, double beta, int K, int dgammamean)
{
/* discretization of gamma distribution with equal proportions in each 
   category.
*/
   int i;
   double t, factor=alpha/beta*K, lnga1;

   if (dgammamean==0) {
      lnga1=LnGamma(alpha+1);
      for (i=0; i<K-1; i++) /* cutting points, Eq. 9 */
         freqK[i]=InverseCDFGamma((i+1.0)/K, alpha, beta);
      for (i=0; i<K-1; i++) /* Eq. 10 */
         freqK[i]=IncompleteGamma(freqK[i]*beta, alpha+1, lnga1);

      for(i=0; i<K; i++) rK[i]=freqK[i];

      rK[0] = freqK[0]*factor;
      rK[K-1] = (1-freqK[K-2])*factor;
      for (i=1; i<K-1; i++)  rK[i] = (freqK[i]-freqK[i-1])*factor;
   }
   else {
      for(i=0; i<K; i++) rK[i]=InverseCDFGamma((i*2.+1)/(2.*K), alpha, beta);
      for(i=0,t=0; i<K; i++) t+=rK[i];
      for(i=0; i<K; i++) rK[i]*=factor/t;
   }
   for (i=0; i<K; i++) freqK[i]=1.0/K;

   return (0);
}
int DiscreteGammad(int K, double alpha, double freqK[], double rK[])
{
  int i,j;
  double Ic[6],da,q,te;

   for(i=0; i<K-1; i++){
     q=InverseCDFGamma((i+1.0)/K,alpha,alpha);
     digami(alpha*q,alpha,Ic);
     da = (-q*Ic[0]-Ic[2])/(alpha*Ic[0]);
     te = (da*alpha+q);
     
     digami(alpha*q,alpha+1.0,Ic);
     rK[i]=Ic[5];
     rK[i+K]=te*Ic[0]+Ic[2];
   }
   rK[K-1]=(1-rK[K-2])*K;
   rK[K-1+K]=-rK[K-2+K]*K;
   for(i=K-2; i>=1; i--)
     for(j=0; j<2; j++) 
       rK[i+j*K]=(rK[i+j*K]-rK[i-1+j*K])*K;
   rK[0] *= K; rK[K] *= K;

   for (i=0; i<K; i++) freqK[i]=1.0/K;

   return (0);
}
int DiscreteGammada(int K, double alpha, double freqK[], double rK[], double h){
  int i;
  DiscreteGamma(freqK,rK,alpha,alpha,K,0);
  DiscreteGamma(freqK,&rK[K],alpha+h,alpha+h,K,0);
  for (i=0; i<=K-1; i++)
    rK[i+K]=(rK[i+K]-rK[i])/h;
  return(0);
}
void error2 (char * message)
{ printf("\nError: %s.\n", message); exit(-1); }
long factorial (int n)
{
   long f=1, i;
   if (n>10) error2("n>10 in factorial");
   for (i=2; i<=(long)n; i++) f *= i;
   return (f);
}
double LnGamma (double x)
{
/* returns ln(gamma(x)) for x>0, accurate to 10 decimal places.
   Stirling's formula is used for the central polynomial part of the procedure.

   Pike MC & Hill ID (1966) Algorithm 291: Logarithm of the gamma function.
   Communications of the Association for Computing Machinery, 9:684
*/
   double f=0, fneg=0, z, lng;
   int nx=(int)x;

   if((double)nx==x && nx>1 && nx<12)
      lng=log((double)factorial(nx-1));
   else {
      if(x<=0) {
         error2("LnGamma not implemented for x<0");
         if((int)x-x==0) { puts("lnGamma undefined"); return(-1); }
         for (fneg=1; x<0; x++) fneg/=x;
         if(fneg<0) error2("strange!! check lngamma");
         fneg=log(fneg);
      }
      if (x<7) {
         f=1;  z=x-1;
         while (++z<7)  f*=z;
         x=z;   f=-log(f);
      }
      z = 1/(x*x);
      lng = fneg+ f + (x-0.5)*log(x) - x + .918938533204673 
             + (((-.000595238095238*z+.000793650793651)*z-.002777777777778)*z
                  +.083333333333333)/x;
   }
   return  lng;
}
double InverseCDFNormal (double prob)
{
/* returns z so that Prob{x<z}=prob where x ~ N(0,1) and (1e-12)<prob<1-(1e-12)
   returns (-9999) if in error
   Odeh RE & Evans JO (1974) The percentage points of the normal distribution.
   Applied Statistics 22: 96-97 (AS70)

   Newer methods:
     Wichura MJ (1988) Algorithm AS 241: the percentage points of the
       normal distribution.  37: 477-484.
     Beasley JD & Springer SG  (1977).  Algorithm AS 111: the percentage 
       points of the normal distribution.  26: 118-121.
*/
   double a0=-.322232431088, a1=-1, a2=-.342242088547, a3=-.0204231210245;
   double a4=-.453642210148e-4, b0=.0993484626060, b1=.588581570495;
   double b2=.531103462366, b3=.103537752850, b4=.0038560700634;
   double y, z=0, p=prob, p1;

   p1 = (p<0.5 ? p : 1-p);
   if (p1<1e-20) z=999;
   else {
      y = sqrt (log(1/(p1*p1)));   
      z = y + ((((y*a4+a3)*y+a2)*y+a1)*y+a0) / ((((y*b4+b3)*y+b2)*y+b1)*y+b0);
   }
   return (p<0.5 ? -z : z);
}
double InverseCDFChi2 (double prob, double v)
{
/* returns z so that Prob{x<z}=prob where x is Chi2 distributed with df=v
   returns -1 if in error.   0.000002<prob<0.999998
   RATNEST FORTRAN by
       Best DJ & Roberts DE (1975) The percentage points of the 
       Chi2 distribution.  Applied Statistics 24: 385-388.  (AS91)
   Converted into C by Ziheng Yang, Oct. 1993.
*/
   double e=.5e-6, aa=.6931471805, p=prob, g, small=1e-6;
   double xx, c, ch, a=0,q=0,p1=0,p2=0,t=0,x=0,b=0,s1,s2,s3,s4,s5,s6;

   if (p<small)   return(0);
   if (p>1-small) return(9999);
   if (v<=0)      return (-1);

   g = LnGamma (v/2);
   xx=v/2;   c=xx-1;
   if (v >= -1.24*log(p)) goto l1;

   ch=pow((p*xx*exp(g+xx*aa)), 1/xx);
   if (ch-e<0) return (ch);
   goto l4;
l1:
   if (v>.32) goto l3;
   ch=0.4;   a=log(1-p);
l2:
   q=ch;  p1=1+ch*(4.67+ch);  p2=ch*(6.73+ch*(6.66+ch));
   t=-0.5+(4.67+2*ch)/p1 - (6.73+ch*(13.32+3*ch))/p2;
   ch-=(1-exp(a+g+.5*ch+c*aa)*p2/p1)/t;
   if (fabs(q/ch-1)-.01 <= 0) goto l4;
   else                       goto l2;
  
l3: 
   x = InverseCDFNormal(p);
   p1 = 0.222222/v;
   ch = v*pow((x*sqrt(p1)+1-p1), 3.0);
   if (ch>2.2*v+6)
      ch = -2*(log(1-p)-c*log(.5*ch)+g);
l4:
   q=ch;   p1=.5*ch;
   if ((t=IncompleteGamma (p1, xx, g))<0)
      error2 ("\nIncompleteGamma");
   p2=p-t;
   t=p2*exp(xx*aa+g+p1-c*log(ch));   
   b=t/ch;  a=0.5*t-b*c;

   s1=(210+a*(140+a*(105+a*(84+a*(70+60*a))))) / 420;
   s2=(420+a*(735+a*(966+a*(1141+1278*a))))/2520;
   s3=(210+a*(462+a*(707+932*a)))/2520;
   s4=(252+a*(672+1182*a)+c*(294+a*(889+1740*a)))/5040;
   s5=(84+264*a+c*(175+606*a))/2520;
   s6=(120+c*(346+127*c))/5040;
   ch+=t*(1+0.5*t*s1-b*c*(s1-b*(s2-b*(s3-b*(s4-b*(s5-b*s6))))));
   if (fabs(q/ch-1) > e) goto l4;

   return (ch);
}
double IncompleteGamma (double x, double alpha, double ln_gamma_alpha)
{
/* returns the incomplete gamma ratio I(x,alpha) where x is the upper 
           limit of the integration and alpha is the shape parameter.
   returns (-1) if in error
   ln_gamma_alpha = ln(Gamma(alpha)), is almost redundant.
   (1) series expansion,     if (alpha>x || x<=1)
   (2) continued fraction,   otherwise
   RATNEST FORTRAN by
   Bhattacharjee GP (1970) The incomplete gamma integral.  Applied Statistics,
   19: 285-287 (AS32)
*/
   int i;
   double p=alpha, g=ln_gamma_alpha;
   double accurate=1e-10, overflow=1e60;
   double factor, gin=0, rn=0, a=0,b=0,an=0,dif=0, term=0, pn[6];

   if (x==0) return (0);
   if (x<0 || p<=0) return (-1);

   factor=exp(p*log(x)-x-g);   
   if (x>1 && x>=p) goto l30;
   /* (1) series expansion */
   gin=1;  term=1;  rn=p;
 l20:
   rn++;
   term *= x/rn;   gin += term;
   if (term > accurate) goto l20;
   gin *= factor/p;
   goto l50;
 l30:
   /* (2) continued fraction */
   a = 1-p;   b = a+x+1;  term = 0;
   pn[0] = 1;  pn[1] = x;  pn[2] = x+1;  pn[3] = x*b;
   gin = pn[2]/pn[3];
 l32:
   a++;  
   b += 2;
   term++;
   an = a*term;
   for (i=0; i<2; i++) 
      pn[i+4] = b*pn[i+2] - an*pn[i];
   if (pn[5] == 0) goto l35;
   rn = pn[4]/pn[5];
   dif = fabs(gin-rn);
   if (dif > accurate) goto l34;
   if (dif <= accurate*rn) goto l42;
 l34:
   gin = rn;
 l35:
   for (i=0; i<4; i++) pn[i] = pn[i+2];
   if (fabs(pn[4]) < overflow) goto l32;
   for (i=0; i<4; i++) pn[i] /= overflow;
   goto l32;
 l42:
   gin = 1-factor*gin;

 l50:
   return (gin);
}
double trigamma(double x){
  double a=1.0e-4,b2=0.1666666667,b4=-0.03333333333,b6=0.02380952381;
  double b8=-0.03333333333,b=5.0,z,trigam,y;
  
  if(x<=a) return(1.0/(x*x));
  z=x; trigam=0.0;
  while(z<b){
    trigam += 1.0/(z*z);
    z += 1.0;
  }
  y = 1.0/(z*z);
  trigam += 0.5*y + (1.0+y*(b2+y*(b4+y*(b6+y*b8))))/z;
  return(trigam);
}
double digamma(double x){
  double s=1.0e-5,c=8.5,s3=8.333333333e-02,s4=8.333333333e-03;
  double s5=3.968253968e-03,d1=-0.5772156649,y,digama=0,r;
  
  y=x;
  if(x <= s) return(d1-1.0/y);
  while(y<c){
    digama -= 1.0/y;
    y += 1.0;
  }
  r=1.0/y;
  digama += log(y)-r*0.5;
  r=r*r;
  digama -= r*(s3-r*(s4-r*s5));
  return(digama);
}
int digami(double x, double p, double *d){
  int i,i2;
  double pn[6],dp[6],dpp[6],e=1.0e-6,oflo=1.0e+30,tmax=1000.0,vsmall=1.0e-30;
  double gplog,gp1log,psip,psip1,psidp,psidp1,pm1,xlog,f,dfp,dfpp,tmaxp,c,s,cp;
  double cpp,dsp,dspp,a,cpc,b,term,s0,an;

  gplog=LnGamma(p);
  gp1log=log(p)+gplog;
  psip=digamma(p);
  psip1=1/p + psip;
  psidp=trigamma(p);
  psidp1=psidp-1/(p*p);

  pm1=p-1.0;
  xlog=log(x);
  d[0]=exp(-gplog + pm1*xlog - x);
  d[1]=d[0]*(pm1/x-1.0);
  d[4]=d[0]*(xlog-psip);

  if(x<=1.0 || x<p){
    f=exp(p*xlog-gp1log-x);
    dfp=f*(xlog-psip1);
    dfpp=dfp*dfp/f-f*psidp1;

    tmaxp=tmax+p;
    c=1.0; s=1.0; cp=0.0; cpp=0.0; dsp=0.0; dspp=0.0;
    a=p;
    do{
      a += 1.0;
      cpc=cp/c;
      cp=cpc-1.0/a;
      cpp=cpp/c-cpc*cpc+1.0/(a*a);
      c=c*x/a;
      cp *= c;
      cpp=cpp*c+cp*cp/c;
      s += c;
      dsp += cp;
      dspp += cpp;
    }while(a<=tmaxp && c>e*s);
    if(a>tmaxp){
      return(1);
    }
    else{
      d[5]=s*f;
      d[2]=s*dfp+f*dsp;
      d[3]=s*dfpp+2.0*dfp*dsp+f*dspp;
      return(0);
    }
  }
  else{
    f=exp(p*xlog-gplog-x);
    dfp=f*(xlog-psip);
    dfpp=dfp*dfp/f-f*psidp;
    
    a=pm1;
    b=x+1.0-a;
    term=0.0;
    pn[0]=1.0;
    pn[1]=x;
    pn[2]=x+1.0;
    pn[3]=x*b;
    s0=pn[2]/pn[3];
    for(i=0; i<4; i++){
      dp[i]=0.0;
      dpp[i]=0.0;
    }
    dp[3]=-x;
    
    do{
      a -= 1.0;
      b += 2.0;
      term += 1.0;
      an=a*term;
      pn[4]=b*pn[2]+an*pn[0];
      pn[5]=b*pn[3]+an*pn[1];
      dp[4]=b*dp[2]-pn[2]+an*dp[0]+pn[0]*term;
      dp[5]=b*dp[3]-pn[3]+an*dp[1]+pn[1]*term;
      dpp[4]=b*dpp[2]+an*dpp[0]+2.0*(term*dp[0]-dp[2]);
      dpp[5]=b*dpp[3]+an*dpp[1]+2.0*(term*dp[1]-dp[3]);
      /* printf("%f %f\n",pn[4],pn[5]); */

      if(fabs(pn[5])>=vsmall){
	s=pn[4]/pn[5];
	c=fabs(s-s0);
	/* printf("%.16f %.16f %.4e %.4e %.4e\n",s0,s,c,p,e); */
	/* printf("%.16f %.e\n",c*p,e); */
	if(c*p<=e && c <= e*s) break;
      }
      s0=s;

      for(i=0; i<4; i++){
	i2=i+2;
	dp[i]=dp[i2];
	dpp[i]=dpp[i2];
	pn[i]=pn[i2];
      }
      if(term>tmax) return(1);
      if(fabs(pn[4]) >= oflo){
	for(i=0; i<4; i++){
	  dp[i] /= oflo;
	  dpp[i] /= oflo;
	  pn[i] /= oflo;
	}}
    }while(term<=tmax);

    d[5]=1.0-f*s;
    dsp=(dp[4]-s*dp[5])/pn[5];
    dspp=(dpp[4]-s*dpp[5]-2.0*dsp*dp[5])/pn[5];
    d[2]=-f*dsp-s*dfp;
    d[3]=-f*dspp-2.0*dsp*dfp-s*dfpp;
    return(0);
  }
}
void makeprotfreqs(double *fr, double *W, double *Lam) 
{
  long i, mineig,nchar=20;
  mineig = 0;
  for (i = 0; i < nchar; i++)
    if (fabs(Lam[i]) < fabs(Lam[mineig]))
      mineig = i;
  for(i = 0; i < nchar; i++) fr[i] =  fabs(W[i+mineig*nchar]);
}
void Mijp(double t, double *fr, double *W, double *Lam, double *f){
  /* transition probs
   * adapted from proml make_pmatrix 
   */
  static double elambdat[NCHAR];
  double q, p0;
  int i, j, k;

  if (t == 0.0){
    for(i = 0; i < NCHAR; i++)
      for(j = 0; j < NCHAR; j++) f[i + j*NCHAR] = (i == j)?1.0:0.0;
    return;
  }

  for(k = 0; k < NCHAR; k++) elambdat[k] = exp(t * Lam[k]);
  for(i = 0; i < NCHAR; i++)
    for(j = 0; j < NCHAR; j++){      
      p0 = 0.0;
      for(k = 0; k < NCHAR; k++){
	q = W[i + k*NCHAR] * W[j+ k*NCHAR];
	p0 += (q * elambdat[k]);
      }
      f[i + j*NCHAR] = p0/fr[i];
    }

}
void Mijpd(double t, double *fr, double *W, double *Lam, double *M, double *Mp, 
	   double *Mpp){
  /* transition probs
   * adapted from proml make_pmatrix 
   * alpha not used
   */
  static double elambdat[NCHAR];
  double q, p0, p0p, p0pp;
  int i, j, k;

/*   if (t == 0.0){ */
/*     for(i = 0; i < NCHAR; i++) */
/*       for(j = 0; j < NCHAR; j++) M[i + j*NCHAR] = (i == j)?1.0:0.0; */
/*     return; */
/*   } */

  for(k = 0; k < NCHAR; k++) elambdat[k] = exp(t * Lam[k]);
  for(i = 0; i < NCHAR; i++)
    for(j = 0; j < NCHAR; j++){      
      p0 = 0.0; p0p = 0.0; p0pp = 0.0;
      for(k = 0; k < NCHAR; k++){
	q = W[i + k*NCHAR] * W[j+ k*NCHAR];
	p0 += (q * elambdat[k]);
	p0p += (q * Lam[k] * elambdat[k]);
	p0pp += (q * Lam[k] * Lam[k] * elambdat[k]);
      }
      M[i + j*NCHAR] = p0/fr[i];
      Mp[i + j*NCHAR] = p0p/fr[i];
      Mpp[i + j*NCHAR] = p0pp/fr[i];
    }
}
void WLamfr2exchange(int nchar, double *W, double *Lam, double *fr, double *R){
  int i,j,k;
  for(i = 0; i < nchar; i++)
    for(j = (i+1); j < nchar; j++){
      R[i+j*nchar] = 0.0;
      for(k = 0; k < nchar; k++) 
	R[i+j*nchar] += W[i+k*nchar]*W[j+k*nchar]*Lam[k];
    }
  
  for(i = 0; i < nchar; i++) 
    for(j = (i+1); j < nchar; j++) R[i+j*nchar] /= (fr[i]*fr[j]);
  for(i = 0; i < nchar; i++) 
    for(j = 0; j < i; j++) R[i+j*nchar] = R[j+i*nchar];
  for(i = 0; i < nchar; i++)R[i+i*nchar] = 0.0;
}
void chQ_freq(int nbin, double *W, double *Lam, double *fr, double *fro)
{
  /* binning/udistfbf.c 
   *     Q = Pi^(-1) W Lam W' 
   * decomposition used to obtain P(t) as
   *     P(t) = Pi^(-1) W exp(Lam t) W'
   * where W is determined from eigen-decomp
   *     Pi^(1/2) Q Pi^(-1/2) = U' Lam U,   W = (Pi)^(1/2) U'
   * To adjust for frequencies Pi*:
   * obtain Q_ij* = Q_ij pi_j* / pi_j
   *        Q_ii* = - sum_{j neq j} Q_ij*
   *        M_ij* =  sqrt(pi_i*) Q_ij* / sqrt(pi_j*)
   *        U and Lam from M = U' Lam U (eigen-decomp)
   *        W = (Pi*)^(1/2) U' */
  int i,j,l;
  double R[400], Offdiag[400], Rt, mu;

/*   for(i = 0; i < nbin; i++) */
/*     for(j = 0; j < nbin; j++){ */
/*       Rt = 0.0; */
/*       for(l = 0; l < nbin; l++) */
/* 	Rt += W[i+l*nbin]*W[j+l*nbin]*Lam[l]; */
/*       Rt /= fro[i]; */
/*       if (i != j) */
/* 	printf("%f\n", Rt); */
/*     } */
/*   exit(0); */

  for(i = 0; i < nbin; i++){ /* R_ij = Q_ij / (pi_i pi_j) */
    for(j = (i+1); j < nbin; j++){
      Rt = 0.0;
      for(l = 0; l < nbin; l++){
	Rt += W[i+l*nbin]*W[j+l*nbin]*Lam[l];
      }
      Rt /= (fro[i]*fro[j]);
      R[i+j*nbin] = Rt; R[j+i*nbin] = Rt;
    }
  }

  for(i = 0; i < nbin; i++){ /* R_ii = Q_ii* */
    Rt = 0.0;
    for(j = 0; j < nbin; j++){
      if (j != i)
	Rt -= R[i+j*nbin]*fr[j];
    }
    R[i+i*nbin] = Rt/fr[i]; /* will cancel with fr[i] below */
  }

/*   for(i = 0; i < nbin; i++) */
/*     for(j = 0; j < nbin; j++) printf("%f\n",R[i+j*nbin]*fr[j]); exit(0); */

  for(i = 0; i < nbin; i++){ /* R_ij = entry of M_ij* above */
    for(j = i; j < nbin; j++){
      R[i+j*nbin] *= sqrt(fr[i]*fr[j]);
      if (j != i) R[j+i*nbin] = R[i+j*nbin];
    }
  }

  eigenRealSym(R, nbin, Lam, Offdiag); /* eigen-decomp of M* */
  for(i = 0; i < nbin; i++){ /* W = (Pi*)^(1/2) U' */
    for(j = 0; j < nbin; j++){
      W[i+j*nbin] = R[j+i*nbin] * sqrt(fr[i]);
    }
  }

  /* rescale eigenvalues so that -sum_i pi_i Q_ii = 1 */
  mu = 0.0;
  for(i = 0; i < nbin; i++)
    for(l = 0; l < nbin; l++) mu -= Lam[l]*W[i+l*nbin]*W[i+l*nbin];
  for(i = 0; i < nbin; i++) Lam[i] /= mu;
}
double int_elamrt(double t, double lam, double alpha, int deriv, double *de){
  double te,dt;
  te = -lam*t/alpha;
  dt= exp(-alpha*log(1+te));
  if(deriv==0) return(dt);
  if(deriv==1 || deriv==2){
    de[0]=lam*exp(-(alpha+1)*log(1+te));
    if(deriv==2) de[1]=(alpha+1)*lam*lam*exp(-(alpha+2)*log(1+te))/alpha;
  }
  if(deriv== 3 || deriv==4){
    de[0] = dt*(te/(1+te)-log(1+te));
    if(deriv==4) 
      de[1]=de[0]*de[0]/dt + 
	dt*(te*te/(alpha*(1+te)*(1+te))-te/(alpha*(1+te)));
  }
  return(dt);
}
double pijJCg(int x, int y, double t, int nchar){
  double m;
  m = nchar/(nchar - 1.0);
  if (x == y) 
    return((1.0/nchar) + exp(-t*m)/m);
  else
    return((1.0/nchar) - (1.0/nchar)*exp(-t*m));
}
double pijJCg_gen(int x, int y, double t, int nchar, double *fr, double *W, 
		  double *Lam){
  return(pijJCg(x,y,t,nchar));
}
double ppijJCg(int x, int y, double t, int nchar){ 
  double m;
  m = nchar/(nchar - 1.0);
  if (x == y) 
    return(-exp(-t*m));
  else
    return(exp(-t*m)/(nchar - 1.0));
}
double ppijJCg_gen(int x, int y, double t, int nchar, double *fr,
		   double *W, double *Lam){ 
  return(ppijJCg(x,y,t,nchar));
}
double pp2ijJCg(int x, int y, double t, int nchar){
  double m;
  m = nchar/(nchar - 1.0);
  if (x == y) 
    return(m*exp(-t*m));
  else
    return(-m*exp(-t*m)/(nchar - 1.0));
}
void pJCm(double t, int nchar, double *fr, int deriv, 
	    double *M, double *Mp, double *Mpp){
  int i,j;
  double m,emt;
  m = nchar/(nchar - 1.0);
  emt=exp(-t*m);
  for(i = 0; i < nchar; i++)
    for(j = 0; j < nchar; j++){
      if(i==j){
	M[i+j*nchar]=1.0/nchar+(1.0-1.0/nchar)*emt;
	if(deriv==1 || deriv==2) Mp[i+j*nchar]=-m*(1.0-1.0/nchar)*emt;
	if(deriv==2) Mpp[i+j*nchar]=m*m*(1.0-1.0/nchar)*emt;
      }
      if(i!=j){
	M[i+j*nchar]=1.0/nchar*(1.0-exp(-m*t));
	if(deriv==1 || deriv==2) Mp[i+j*nchar]=m*1.0/nchar*emt;
	if(deriv==2) Mpp[i+j*nchar]=-m*m*1.0/nchar*emt;
      }
    }
}
void pJCm_gen(double t, int nchar, double *fr, double *W, double *Lam, 
		int deriv, double *M, double *Mp, double *Mpp){
  pJCm(t,nchar,fr,deriv,M,Mp,Mpp);
}
void pJCma(double t, int nchar, double *fr, int deriv, double alpha,
	   double *M, double *Mp, double *Mpp){
  int i,j;
  double mu,emut,demu[2];
  mu = nchar/(nchar - 1.0);
  emut=int_elamrt(t,-1.0*mu,alpha,deriv,demu);

  for(i = 0; i < nchar; i++)
    for(j = 0; j < nchar; j++){
      if(i==j){
	M[i+j*nchar]=1.0/nchar+(1.0-1.0/nchar)*emut;
	if(deriv > 0) 
	  Mp[i+j*nchar]=(1.0-1.0/nchar)*demu[0];
	if(deriv==2 || deriv==4) Mpp[i+j*nchar]=(1.0-1.0/nchar)*demu[1];
      }
      if(i!=j){
	M[i+j*nchar]=1.0/nchar-emut/nchar;
	if(deriv > 0) 
	  Mp[i+j*nchar]=-demu[0]/nchar;
	if(deriv==2 || deriv==4) Mpp[i+j*nchar]=-demu[1]/nchar;
      }
    }
}
void pJCma_gen(double t, int nchar, double *fr, double *W, double *Lam, 
	       int deriv, double alpha, double *M, double *Mp, double *Mpp){
  pJCma(t,nchar,fr,deriv,alpha,M,Mp,Mpp);
}
double muF81f(int nchar, double *fr){
  int l;
  double mu = 0.0;
  for(l = 0; l < nchar; l++) mu += fr[l]*(1.0-fr[l]);
  return(1.0/mu);
}
double pijF81(int i, int j, double t, int nchar, double mu, double *fr){
  if (i == j) return(fr[j]+(1.0-fr[j])*exp(-mu*t));
  if (i != j) return(fr[j]*(1.0-exp(-mu*t)));
  return(fr[j]*(1.0-exp(-mu*t)));
}
double pijF81_gen(int i, int j, double t, int nchar, double *fr, double *mu,
		  double *Lam){
  return(pijF81(i,j,t,nchar,*mu,fr));
}
double ppijF81(int i, int j, double t, int nchar, double mu, double *fr){
  if (i == j) return(-mu*(1.0-fr[j])*exp(-mu*t));
  if (i != j) return(mu*exp(-mu*t)*fr[j]);
  return(mu*exp(-mu*t)*fr[j]);
}
double ppijF81_gen(int i, int j, double t, int nchar, double *fr, double *mu,
		   double *Lam){
  return(ppijF81(i,j,t,nchar,*mu,fr));
}
void pF81m(double t, int nchar, double *fr, double mu, int deriv, 
	   double *M, double *Mp, double *Mpp){
  int i,j;
  double emut;
  emut=exp(-mu*t);
  for(i = 0; i < nchar; i++)
    for(j = 0; j < nchar; j++){
      if(i==j){
	M[i+j*nchar]=fr[j]+(1.0-fr[j])*emut;
	if(deriv==1 || deriv==2) Mp[i+j*nchar]=-mu*(1.0-fr[j])*emut;
	if(deriv==2) Mpp[i+j*nchar]=mu*mu*(1.0-fr[j])*emut;
      }
      if(i!=j){
	M[i+j*nchar]=fr[j]*(1.0-exp(-mu*t));
	if(deriv==1 || deriv==2) Mp[i+j*nchar]=mu*fr[j]*emut;
	if(deriv==2) Mpp[i+j*nchar]=-mu*mu*fr[j]*emut;
      }
    }
}
void pF81m_gen(double t, int nchar, double *fr, double *mu, double *Lam, 
		 int deriv, double *M, double *Mp, double *Mpp){
  pF81m(t,nchar,fr,*mu,deriv,M,Mp,Mpp);
}
void pF81ma(double t, int nchar, double *fr, double mu, int deriv, double alpha,
	   double *M, double *Mp, double *Mpp){
  int i,j;
  double emut,demu[2];
  emut=int_elamrt(t,-1.0*mu,alpha,deriv,demu);
  for(i = 0; i < nchar; i++)
    for(j = 0; j < nchar; j++){
      if(i==j){
	M[i+j*nchar]=fr[j]+(1.0-fr[j])*emut;
	if(deriv > 0) Mp[i+j*nchar]=(1-fr[j])*demu[0];
	if(deriv==2 || deriv==4) Mpp[i+j*nchar]=(1-fr[j])*demu[1];
      }
      if(i!=j){
	M[i+j*nchar]=fr[j]-fr[j]*emut;
	if(deriv > 0) Mp[i+j*nchar]=-fr[j]*demu[0];
	if(deriv==2 || deriv==4) Mpp[i+j*nchar]=-fr[j]*demu[1];
      }
    }
}
void pF81ma_gen(double t, int nchar, double *fr, double *mu, double *Lam, 
		int deriv, double alpha, double *M, double *Mp, double *Mpp){
  pF81ma(t,nchar,fr,*mu,deriv,alpha,M,Mp,Mpp);
}
int is_transitionf(int i, int j){
  return((i==0 && j==2) || (i==2 && j==0) || (i==1 && j==3) || (i==3 && j==1));
}
double muF84f(double K, double *fr){
  double mu = 0.0;
  int l;
/*   printf("%f\n",K); */
  for(l = 0; l < 4; l++) mu += fr[l]*(1.0-fr[l]);
  mu += 2*K*( fr[0]*fr[2]/(fr[0]+fr[2]) + fr[1]*fr[3]/(fr[1]+fr[3]) );
  return(1.0/mu);
}
double pijF84(int i, int j, double t, double mu, double K, double *fr){
  double Pj;
  int is_transition;
/*   printf("1.0/(K+1):%f\n",1.0/(K+1)); */
/*   printf("(K+1)*mu*t:%f\n",(K+1)*mu*t); */
/*   printf("mu*t:%f\n",mu*t); */
  if (j == 0 || j == 2) 
    Pj=fr[0]+fr[2];
  else
    Pj=fr[1]+fr[3];

  if(i == j)
    return(fr[j]+fr[j]*(1/Pj - 1)*exp(-mu*t)+(1-fr[j]/Pj)*exp(-mu*(K+1)*t));

  if (i != j){
    is_transition = is_transitionf(i,j);
    if (is_transition)
      return(fr[j]+fr[j]*(1/Pj - 1)*exp(-mu*t)-(fr[j]/Pj)*exp(-mu*(K+1)*t));
    else
      return(fr[j]*(1.0-exp(-mu*t)));
  }
  return(-1.0);
}
double pijF84_gen(int i, int j, double t, int nchar, double *fr, double *mu,
		  double *K){
  return(pijF84(i,j,t,*mu,*K,fr));
}
double ppijF84(int i, int j, double t, double mu, double K, double *fr){
  double Pj,f;
  int is_transition;
  if (j == 0 || j == 2) 
    Pj=fr[0]+fr[2];
  else
    Pj=fr[1]+fr[3];
  
  if(i == j)
    f=-mu*fr[j]*(1/Pj - 1)*exp(-mu*t) + 
      -1.0*mu*(K+1)*(1-fr[j]/Pj)*exp(-mu*(K+1)*t);
  
  if (i != j){
    is_transition = is_transitionf(i,j);
    if (is_transition)
      f = -mu*fr[j]*(1/Pj - 1)*exp(-mu*t)+
	mu*(K+1)*(fr[j]/Pj)*exp(-mu*(K+1)*t);
    else
      f = mu*fr[j]*exp(-mu*t);
  }
  return(f);
}
double ppijF84_gen(int i, int j, double t, int nchar, double *fr, double *mu,
		  double *K){
  return(ppijF84(i,j,t,*mu,*K,fr));
}
double p2ijF84(int i, int j, double t, double mu, double K, double *fr){
  double Pj,f;
  int is_transition;
  if (j == 0 || j == 2) 
    Pj=fr[0]+fr[2];
  else
    Pj=fr[1]+fr[3];
  
  if(i == j)
    f=mu*mu*fr[j]*(1/Pj - 1)*exp(-mu*t) + 
      mu*(K+1)*mu*(K+1)*(1-fr[j]/Pj)*exp(-mu*(K+1)*t);
  
  if (i != j){
    is_transition = is_transitionf(i,j);
    if (is_transition)
      f = mu*mu*fr[j]*(1/Pj - 1)*exp(-mu*t) -
	mu*(K+1)*mu*(K+1)*(fr[j]/Pj)*exp(-mu*(K+1)*t);
    else
      f = -mu*mu*fr[j]*exp(-mu*t);
  }
  return(f);
}
double p2ijF84_gen(int i, int j, double t, int nchar, double *fr, double *mu,
		  double *K){
  return(p2ijF84(i,j,t,*mu,*K,fr));
}
void pF84m(double t, double mu, double K, double *fr, int deriv, 
	   double *M, double *Mp, double *Mpp){
  int i,j;
  double Pr,Py,Pj,muK,emuKt,emut;
  Pr=fr[0]+fr[2];
  Py=fr[1]+fr[3];
  muK=mu*(K+1); emuKt=exp(-muK*t); emut=exp(-mu*t);

  for(j = 0; j < 4; j++){
    if (j == 0 || j == 2) Pj=Pr;
    if (j == 1 || j == 3) Pj=Py;
    for(i = 0; i < 4; i++){
      if(i==j){
	M[i+j*4]=fr[j]+fr[j]*(1/Pj-1)*emut+(1-fr[j]/Pj)*emuKt;
	if(deriv==1 || deriv==2)
	  Mp[i+j*4]=-mu*fr[j]*(1/Pj-1)*emut-muK*(1-fr[j]/Pj)*emuKt;
	if(deriv==2)
	  Mpp[i+j*4]=mu*mu*fr[j]*(1/Pj-1)*emut + muK*muK*(1-fr[j]/Pj)*emuKt;
      }
      if(i!=j){
	if(is_transitionf(i,j)){
	  M[i+j*4]=fr[j]+fr[j]*(1/Pj-1)*emut-(fr[j]/Pj)*emuKt;
	  if(deriv==1 || deriv==2)
	    Mp[i+j*4]=-mu*fr[j]*(1/Pj-1)*emut+muK*(fr[j]/Pj)*emuKt;
	  if(deriv==2)
	    Mpp[i+j*4]=mu*mu*fr[j]*(1/Pj-1)*emut-muK*muK*(fr[j]/Pj)*emuKt;
	}
	else{
	  M[i+j*4]=fr[j]*(1.0-emut);
	  if(deriv==1 || deriv==2) Mp[i+j*4] = mu*fr[j]*emut;
	  if(deriv==2) Mpp[i+j*4]=-mu*mu*fr[j]*emut;
	}
      }
    }
  }
}
void pF84m_gen(double t, int nchar, double *fr, double *mu, double *K, 
	       int deriv, double *M, double *Mp, double *Mpp){
  pF84m(t,*mu,*K,fr,deriv,M,Mp,Mpp);
}
void pF84ma(double t, double mu, double K, double *fr, int deriv, double alpha,
	    double *M, double *Mp, double *Mpp){
  int i,j;
  double Pr,Py,Pj,emuKt,emut,demuK[2],demu[2];
  Pr=fr[0]+fr[2];
  Py=fr[1]+fr[3];
  emut=int_elamrt(t,-1.0*mu,alpha,deriv,demu);
  emuKt=int_elamrt(t,-1.0*mu*(K+1),alpha,deriv,demuK);
/*   if (deriv==4) printf("%f %f\n",emut,emuKt); */

  for(j = 0; j < 4; j++){
    if (j == 0 || j == 2) Pj=Pr;
    if (j == 1 || j == 3) Pj=Py;
    for(i = 0; i < 4; i++){
      if(i==j){
	M[i+j*4]=fr[j]+fr[j]*(1/Pj-1)*emut+(1-fr[j]/Pj)*emuKt;
	if(deriv > 0)
	  Mp[i+j*4]=fr[j]*(1/Pj-1)*demu[0]+(1-fr[j]/Pj)*demuK[0];
	if(deriv==2 || deriv==4)
	  Mpp[i+j*4]=fr[j]*(1/Pj-1)*demu[1]+(1-fr[j]/Pj)*demuK[1];
      }
      if(i!=j){
	if(is_transitionf(i,j)){
	  M[i+j*4]=fr[j]+fr[j]*(1/Pj-1)*emut-(fr[j]/Pj)*emuKt;
	  if(deriv > 0)
	    Mp[i+j*4]=fr[j]*(1/Pj-1)*demu[0]-(fr[j]/Pj)*demuK[0];
	  if(deriv==2 || deriv==4)
	    Mpp[i+j*4]=fr[j]*(1/Pj-1)*demu[1]-(fr[j]/Pj)*demuK[1];
	}
	else{
	  M[i+j*4]=fr[j]-fr[j]*emut;
	  if(deriv > 0) Mp[i+j*4] = -fr[j]*demu[0];
	  if(deriv==2 || deriv==4) Mpp[i+j*4]=-fr[j]*demu[1];
	}
      }
    }
  }
}
void pF84ma_gen(double t, int nchar, double *fr, double *mu, double *K, 
		int deriv, double alpha, double *M, double *Mp, double *Mpp){
  pF84ma(t,*mu,*K,fr,deriv,alpha,M,Mp,Mpp);
}
double tt2K(double R, double *fr){
  double pY,pR,K;
  pY=fr[1]+fr[3]; pR=fr[0]+fr[2];
  K=(R*pR*pY-fr[0]*fr[2]-fr[1]*fr[3])/(fr[0]*fr[2]/pR+fr[1]*fr[3]/pY);
  return(K);
}
double K2tt(double K, double *fr){
  double pY,pR,R;
  pY=fr[1]+fr[3]; pR=fr[0]+fr[2];
  R=K*(fr[0]*fr[2]/pR+fr[1]*fr[3]/pY)/(pR*pY) + 
    (fr[0]*fr[2]+fr[1]*fr[3])/(pR*pY);
  return(R);
}
double muHKYf(double kappa, double *fr){
  double mu;
  mu=2.0*(kappa*(fr[0]*fr[2]+fr[1]*fr[3])+(fr[0]+fr[2])*(fr[1]+fr[3]));
  return(1.0/mu);
}
double pijHKY(int i, int j, double t, double mu, double kappa, double *fr){
  int is_transition;
  double A,Pi_j;
  is_transition=is_transitionf(i,j);
  if (i !=j && !is_transition) 
    return(fr[j]*(1.0-exp(-mu*t)));

  if (j == 0 || j == 2) Pi_j=fr[0]+fr[2];
  if (j == 1 || j == 3) Pi_j=fr[1]+fr[3];
  A = 1+Pi_j*(kappa-1);
  if (i == j){
    return(fr[j]+fr[j]*(1/Pi_j - 1)*exp(-mu*t) +(1-fr[j]/Pi_j)*exp(-mu*t*A));
  }
  /* must be transition */
  return(fr[j]+fr[j]*(1/Pi_j - 1)*exp(-mu*t) -fr[j]*exp(-mu*t*A)/Pi_j);
}
double pijHKY_gen(int i, int j, double t, int nchar, double *fr, double *mu,
		  double *kappa){
  return(pijHKY(i,j,t,*mu,*kappa,fr));
}
double ppijHKY(int i, int j, double t, double mu, double kappa, double *fr){
  int is_transition;
  double A,Pi_j;
  is_transition=is_transitionf(i,j);
  if (i !=j && !is_transition) 
    return(mu*fr[j]*exp(-mu*t));

  if (j == 0 || j == 2) Pi_j=fr[0]+fr[2];
  if (j == 1 || j == 3) Pi_j=fr[1]+fr[3];
  A = 1+Pi_j*(kappa-1);
  if (i == j){
    return(-mu*fr[j]*(1/Pi_j-1)*exp(-mu*t)-mu*A*(1-fr[j]/Pi_j)*exp(-mu*t*A));
  }
  /* must be transition */
  return(-mu*fr[j]*(1/Pi_j - 1)*exp(-mu*t)+mu*A*fr[j]*exp(-mu*t*A)/Pi_j);
}
double ppijHKY_gen(int i, int j, double t, int nchar, double *fr, double *mu,
		   double *kappa){
  return(ppijHKY(i,j,t,*mu,*kappa,fr));
}
double p2ijHKY(int i, int j, double t, double mu, double kappa, double *fr){
  double A,Pi_j;
  if (i !=j && !is_transitionf(i,j)) 
    return(-mu*mu*fr[j]*exp(-mu*t));
  if (j == 0 || j == 2) Pi_j=fr[0]+fr[2];
  if (j == 1 || j == 3) Pi_j=fr[1]+fr[3];
  A = 1+Pi_j*(kappa-1);
  if (i == j){
    return(mu*mu*fr[j]*(1/Pi_j-1)*exp(-mu*t)-mu*A*mu*A*(1-fr[j]/Pi_j)*exp(-mu*t*A));
  }
  /* must be transition */
  return(mu*mu*fr[j]*(1/Pi_j - 1)*exp(-mu*t)-mu*A*mu*A*fr[j]*exp(-mu*t*A)/Pi_j);
}
double p2ijHKY_gen(int i, int j, double t, int nchar, double *fr, double *mu,
		   double *kappa){
  return(p2ijHKY(i,j,t,*mu,*kappa,fr));
}
void pHKYm(double t, int nchar, double *fr, double mu, double kappa,
	   int deriv, double *M, double *Mp, double *Mpp){
  int i,j;
  double Pr,Py,A,emuAt,emut,muA,Pj;

  Pr=fr[0]+fr[2];
  Py=fr[1]+fr[3];
  emut=exp(-mu*t);

  for(j = 0; j < 4; j++){

    if (j == 0 || j == 2) Pj=Pr;
    if (j == 1 || j == 3) Pj=Py;
    A = 1+Pj*(kappa-1); muA=mu*A;
    emuAt=exp(-muA*t);

    for(i = 0; i < 4; i++){
      if(i==j){
	M[i+j*4] = fr[j]+fr[j]*(1/Pj-1)*emut+(1-fr[j]/Pj)*emuAt;
	if(deriv==1 || deriv==2)
	  Mp[i+j*4] = -mu*fr[j]*(1/Pj-1)*emut-muA*(1-fr[j]/Pj)*emuAt;
	if(deriv==2)
	  Mpp[i+j*4] = mu*mu*fr[j]*(1/Pj-1)*emut+muA*muA*(1-fr[j]/Pj)*emuAt;
      }
      if(i!=j){
	if(is_transitionf(i,j)){
	  M[i+j*4] = fr[j]+fr[j]*(1/Pj-1)*emut-fr[j]*emuAt/Pj;
	  if(deriv==1 || deriv==2)
	    Mp[i+j*4] = -mu*fr[j]*(1/Pj-1)*emut+muA*fr[j]*emuAt/Pj;
	  if(deriv==2)
	    Mpp[i+j*4] = mu*mu*fr[j]*(1/Pj-1)*emut-muA*muA*fr[j]*emuAt/Pj;
	}
	else{
	  M[i+j*4] = fr[j]-fr[j]*emut;
	  if(deriv==1 || deriv==2) Mp[i+j*4] = mu*fr[j]*emut;
	  if(deriv==2) Mpp[i+j*4] = -mu*mu*fr[j]*emut;
	}
      }
    }}
}
void pHKYm_gen(double t, int nchar, double *fr, double *mu, double *kappa, 
	       int deriv, double *M, double *Mp, double *Mpp){
  pHKYm(t,nchar,fr,*mu,*kappa,deriv,M,Mp,Mpp);
}
void pHKYma(double t, int nchar, double *fr, double mu, double kappa,
	    int deriv, double alpha, double *M, double *Mp, double *Mpp){
  int i,j;
  double Pr,Py,A,emuAt,demuA[2],emut,demu[2],muA,Pj;

  Pr=fr[0]+fr[2];
  Py=fr[1]+fr[3];
  emut=int_elamrt(t,-1.0*mu,alpha,deriv,demu);

  for(j = 0; j < 4; j++){

    if (j == 0 || j == 2) Pj=Pr;
    if (j == 1 || j == 3) Pj=Py;
    A = 1+Pj*(kappa-1); muA=mu*A;
    emuAt=int_elamrt(t,-1.0*muA,alpha,deriv,demuA);

    for(i = 0; i < 4; i++){
      if(i==j){
	M[i+j*4] = fr[j]+fr[j]*(1/Pj-1)*emut+(1-fr[j]/Pj)*emuAt;
	if(deriv > 0)
	  Mp[i+j*4] = fr[j]*(1/Pj-1)*demu[0]+(1-fr[j]/Pj)*demuA[0];
	if(deriv==2 || deriv==4)
	  Mpp[i+j*4] = fr[j]*(1/Pj-1)*demu[1]+(1-fr[j]/Pj)*demuA[1];
      }
      if(i!=j){
	if(is_transitionf(i,j)){
	  M[i+j*4] = fr[j]+fr[j]*(1/Pj-1)*emut-fr[j]*emuAt/Pj;
	  if(deriv > 0)
	    Mp[i+j*4] = fr[j]*(1/Pj-1)*demu[0]-fr[j]*demuA[0]/Pj;
	  if(deriv==2 || deriv==4)
	    Mpp[i+j*4] = fr[j]*(1/Pj-1)*demu[1]-fr[j]*demuA[1]/Pj;
	}
	else{
	  M[i+j*4] = fr[j]-fr[j]*emut;
	  if(deriv > 0) Mp[i+j*4] = -fr[j]*demu[0];
	  if(deriv==2 || deriv==4) Mpp[i+j*4] = -fr[j]*demu[1];
	}
      }
    }}
}
void pHKYma_gen(double t, int nchar, double *fr, double *mu, double *kappa, 
	       int deriv, double alpha, double *M, double *Mp, double *Mpp){
  pHKYma(t,nchar,fr,*mu,*kappa,deriv,alpha,M,Mp,Mpp);
}
double tt2kappa(double R, double *fr){
  double pR,pY;
  pR=fr[0]+fr[2];
  pY=fr[1]+fr[3];
  return(R*pR*pY/(fr[0]*fr[2]+fr[1]*fr[3]));
}
double kappa2tt(double kappa,  double *fr){
  double pR,pY;
  pR=fr[0]+fr[2];
  pY=fr[1]+fr[3];
  return(kappa*(fr[0]*fr[2]+fr[1]*fr[3])/(pR*pY));
}
void rmnegpv(int m, double *pv, int incp){
  int i,mi;
  double sum=0.0;
  mi=m*incp;
  for(i=0; i<mi; i+=incp)
    if(pv[i] < 0.0)
      pv[i] = 0.0;
    else
      sum += pv[i];
  for(i=0; i<mi; i+=incp) pv[i] /= sum;
}
void rmnegtrans(int m, double *P){
  int i,j;

  for(i=0; i<m; i++)
    for(j=0; j<m; j++)
      if(P[i+j*m] < 0.0){
	rmnegpv(m,&P[i],m);
	break;
      }
  return;
}
void Q2frWLam(int nbin, double *Q, double *fr, double *W, double *Lam){
  int i,j;
  double R[400], Offdiag[400], mu;

  /* pi_j = [1 + sum_i neq j Q_ji/Q_ij]**(-1) */
  for(j = 0; j < nbin; j++){
    fr[j] = 1.0;
    for(i = 0; i < nbin; i++)
      if (j != i) fr[j] += Q[j+i*nbin]/Q[i+j*nbin];
    fr[j] = 1.0 / fr[j];
  }
  for(i = 0; i < nbin; i++)
    for(j = 0; j < nbin; j++) /* R =  Pi^(1/2) Q Pi^(-1/2) */
      R[i+j*nbin] = sqrt(fr[i])*Q[i+j*nbin]/sqrt(fr[j]);

  /* eigen-decomp of Pi^(1/2) Q Pi^(-1/2) */
  eigenRealSym(R, nbin, Lam, Offdiag); 

  for(i = 0; i < nbin; i++){ /* W = (Pi*)^(1/2) U' */
    for(j = 0; j < nbin; j++)
      W[i+j*nbin] = R[j+i*nbin] * sqrt(fr[i]);
  }

  /* rescale eigenvalues so that -sum_i pi_i Q_ii = 1 */
  mu = 0.0;
  for(i = 0; i < nbin; i++)
    for(j = 0; j < nbin; j++) mu -= Lam[j]*W[i+j*nbin]*W[i+j*nbin];
  for(i = 0; i < nbin; i++) Lam[i] /= mu;
}
void GetWLam(int nchar,double *S, double *fr, double *Q, 
	     double *W, double *Lam){
  int i,j;					
  double mu;
  
  for(i=0; i<nchar; i++)
    for(j=0; j<nchar; j++) Q[i+j*nchar]=S[i+j*nchar]*fr[j];
  for(i=0; i<nchar; i++) 
    for(j=0; j<nchar; j++) if(j!=i) Q[i+i*nchar] -= Q[i+j*nchar];
  mu=0.0;
  for(i=0; i<nchar; i++) mu -= fr[i]*Q[i+i*nchar];
  for(i=0; i<400; i++) Q[i] /= mu;
  
  Q2frWLam(nchar,Q,fr,W,Lam);
  return;
}
double pijGTR(int i, int j, double t, int nchar, double *fr, double *W, 
	      double *Lam){
  int k;
  double f;
  if (t == 0.0){
    if (i == j) return(1.0);
    if (i != j) return(0.0);
  }
  
  f=0.0;
  /* printf("%i\n", nchar); */
  for(k = 0; k < nchar; k++){
    /* printf("%f %f\n", W[i + k*nchar], W[j+ k*nchar]); */
    /* printf("%f\n", Lam[k]); */
    f += W[i + k*nchar] * W[j+ k*nchar] * exp(t*Lam[k]);
  }
  return(f/fr[i]);
}
double ppijGTR(int i, int j, double t, int nchar, double *fr, double *W, 
	       double *Lam){
   int k;
   double f;
   f=0.0;
   for(k = 0; k < nchar; k++) 
     f += Lam[k]*W[i + k*nchar] * W[j+ k*nchar] * exp(t*Lam[k]);
   return(f/fr[i]);
}
double p2ijGTR(int i, int j, double t, int nchar, double *fr, double *W, 
	       double *Lam){
   int k;
   double f;
   f=0.0;
   for(k = 0; k < nchar; k++) 
     f += Lam[k]*Lam[k]*W[i + k*nchar] * W[j+ k*nchar] * exp(t*Lam[k]);
   return(f/fr[i]);
}
void pGTRm(double t, int nchar, double *fr, double *W, double *Lam, 
	   int deriv, double *M, double *Mp, double *Mpp){
  int i,j,k;
  double w;
  for(i = 0; i < nchar*nchar; i++) M[i]=0.0;
  if(deriv==1 || deriv==2) for(i = 0; i < nchar*nchar; i++) Mp[i]=0.0;
  if(deriv==2) for(i = 0; i < nchar*nchar; i++) Mpp[i]=0.0;
  
  for(i = 0; i < nchar; i++)
    for(j = 0; j < nchar; j++){
      for(k = 0; k < nchar; k++){
	w=W[i + k*nchar] * W[j+ k*nchar] * exp(Lam[k]*t);
	M[i+j*nchar] += w;
	if(deriv==1 || deriv==2) Mp[i+j*nchar] += Lam[k]*w;
	if(deriv==2) Mpp[i+j*nchar] += Lam[k]*Lam[k]*w;
      }
      M[i+j*nchar] /= fr[i];
      if(deriv==1 || deriv==2) Mp[i+j*nchar] /= fr[i];
      if(deriv==2) Mpp[i+j*nchar] /= fr[i];
    }
  rmnegtrans(nchar,M);
}
void pGTRma(double t, int nchar, double *fr, double *W, double *Lam, 
	    int deriv, double alpha, double *M, double *Mp, double *Mpp){
  int i,j,k;
  double dt,de[2];
  for(i = 0; i < nchar*nchar; i++) M[i]=0.0;
  if(deriv > 0) for(i = 0; i < nchar*nchar; i++) Mp[i]=0.0;
  if(deriv==2 || deriv==4) for(i = 0; i < nchar*nchar; i++) Mpp[i]=0.0;
  
  for(i = 0; i < nchar; i++)
    for(j = 0; j < nchar; j++){
      for(k = 0; k < nchar; k++){
	dt=int_elamrt(t,Lam[k],alpha,deriv,de);
	M[i+j*nchar] += W[i + k*nchar] * W[j+ k*nchar] * dt;
	if(deriv > 0){
	  Mp[i+j*nchar] += W[i + k*nchar] * W[j+ k*nchar] * de[0];
	}
	if(deriv==2 || deriv==4)
	  Mpp[i+j*nchar] += W[i + k*nchar] * W[j+ k*nchar] * de[1];
      }
      M[i+j*nchar] /= fr[i];
      if(deriv > 0) Mp[i+j*nchar] /= fr[i];
      if(deriv==2 || deriv==4) Mpp[i+j*nchar] /= fr[i];
    }
  rmnegtrans(nchar,M);
}
void LG_exchangeability(double *S){
  int i,j,idx=0;
  for(i=0; i<20; i++) S[i+i*20]=0.0;
  for(i=0; i<20; i++)
    for(j=0; j<i; j++){
      S[i+j*20] = lgpaml[idx++];
      S[j+i*20]=S[i+j*20];
    }
}
void JTT_exchangeability(double *S){
  int i,j,idx=0;
  for(i=0; i<20; i++) S[i+i*20]=0.0;
  for(i=0; i<20; i++)
    for(j=0; j<i; j++){
      S[i+j*20] = jttpaml[idx++];
      S[j+i*20]=S[i+j*20];
    }
}
void WAG_exchangeability(double *S){
  int i,j,idx=0;
  for(i=0; i<20; i++) S[i+i*20]=0.0;
  for(i=0; i<20; i++)
    for(j=0; j<i; j++){
      S[i+j*20] = wagpaml[idx++];
      S[j+i*20]=S[i+j*20];
    }
}
int subst_model_setp(int smodel,
		     double (*(*pij))(int i, int j, double t, int nchar,
				     double *fr, double *W, double *Lam),
		     double (*(*ppij))(int i, int j, double t, int nchar,
				       double *fr, double *W, double *Lam)){
  /* IN:
   * smodel - 0:JC, 1:F81, 2:F84, 3:GTR, 4:PAM, 5:JTT, 6:WAG, 7:mtREV24, 8:HKY,
   *          9:LG
   *
   * OUT:
   * pij,ppij - transition function and first derivative
   */
  switch(smodel){
  case 0:
    *pij=&pijJCg_gen; *ppij=&ppijJCg_gen;
    break;
  case 1:
    *pij=&pijF81_gen; *ppij=&ppijF81_gen;
    break;
  case 2:
    *pij=&pijF84_gen; *ppij=&ppijF84_gen;
    break;
  case 3: case 4: case 5: case 6: case 7: case 9:
    *pij=&pijGTR; *ppij=&ppijGTR;
    break;
  case 8:
    *pij=&pijHKY_gen; *ppij=&ppijHKY_gen;
    break;
  }
  return(1);
}
int subst_model_setpm(int smodel,
		      void (*(*pm))(double t, int nchar, double *fr, 
				    double *W, double *Lam, int deriv, 
				    double *M, double *Mp, 
				    double *Mpp)){
  /* IN:
   * smodel - 0:JC, 1:F81, 2:F84, 3:GTR, 4:PAM, 5:JTT, 6:WAG, 7:mtREV24, 8:HKY,
   *          9:LG
   *
   * OUT:
   * pm - transition matrix function 
   */
  switch(smodel){
  case 0:
    *pm=&pJCm_gen; 
    break;
  case 1:
    *pm=&pF81m_gen; 
    break;
  case 2:
    *pm=&pF84m_gen; ;
    break;
  case 3: case 4: case 5: case 6: case 7: case 9:
    *pm=&pGTRm; 
    break;
  case 8:
    *pm=&pHKYm_gen;
    break;
  }
  return(0);
}
int subst_model_setpma(int smodel,
		       void (*(*pm))(double t, int nchar, double *fr, 
				     double *W, double *Lam, int deriv, 
				     double alpha,
				     double *M, double *Mp, double *Mpp)){
  /* IN:
   * smodel - 0:JC, 1:F81, 2:F84, 3:GTR, 4:PAM, 5:JTT, 6:WAG, 7:mtREV24, 8:HKY,
   *          9:LG
   *
   * OUT:
   * pm - transition matrix function 
   */
  switch(smodel){
  case 0:
    *pm=&pJCma_gen; 
    break;
  case 1:
    *pm=&pF81ma_gen; 
    break;
  case 2:
    *pm=&pF84ma_gen; ;
    break;
  case 3: case 4: case 5: case 6: case 7: case 9:
    *pm=&pGTRma; 
    break;
  case 8:
    *pm=&pHKYma_gen;
    break;
  }
  return(0);
}
int subst_model_setparam(int nchar, int smodel, double *fr, double *Qk,
			 double **W, double **Lam){
  /* IN:
   * nchar - number of character states
   * smodel - 0:JC, 1:F81, 2:F84, 3:GTR, 4:PAM, 5:JTT, 6:WAG, 7:mtREV24, 8:HKY,
   *          9:LG
   * Qk - rate matrix for GTR, ttratio, K or Kappa for F84, HKY, < 0 => ttratio; 
   * not used otherwise. 
   *
   * IN/OUT:
   * fr - frequencies; reset for JC, determined from Q for GTR
   *      if *fr < 0 for aa models, model-based frequencies used
   * OUT:
   * W,Lam - contain parameters for substitution model
   *
   * RETURN:
   * 2,1 or 0 according to whether both W and Lam were allocated, just W or
   * neither
   */
  int allocWLam=0,aamodel=0,i;
  double *fro,*Wo,*Lamo;
  if (smodel==4 || smodel==5 || smodel==6 || smodel==7 || smodel==9) aamodel=1;

  if (smodel==0){ /* JC */
    for(i = 0; i < nchar; i++) fr[i] = 1.0/nchar;
  }
  if (smodel==1){ /* F81 */
    *W=(double *) malloc((size_t) sizeof(double));
    **W=muF81f(nchar,fr);
/*     for(i = 0; i < nchar; i++) printf("%f\n",fr[i]); printf("\n"); */
/*     printf("%f\n",**W); */
    allocWLam=1;
  }
  if (smodel==2){ /* F84 */
    *W=(double *) malloc((size_t) sizeof(double));
    *Lam=(double *) malloc((size_t) sizeof(double));
/*     printf("%.16e\n",*Qk); */
/*     for(i = 0; i < nchar; i++) printf("%f\n",fr[i]); printf("\n"); */
    **Lam=*Qk;
    if(*Qk < 0) 
      **Lam=tt2K(-(*Qk),fr);
    **W= muF84f(**Lam,fr);
/*     printf("%f %f\n",**W,**Lam); */
    allocWLam=2;
  }
  if (smodel==3){ /* GTR */
    *W = (double *) malloc((size_t) nchar*nchar*sizeof(double));
    *Lam = (double *) malloc((size_t) nchar*sizeof(double));
    Q2frWLam(nchar,Qk,fr,*W,*Lam);
    allocWLam=2;
  }
  if (smodel==4){ /* PAM */
    Wo = pamprobmat; Lamo = pameigmat;
  }
  if (smodel==5){ /* JTT */
    Wo = jttprobmat; Lamo = jtteigmat;
  }
  if (smodel==6){ /* WAG */
    Wo = wagprobmat; Lamo = wageigmat;
  }
  if (smodel==7){ /* mtREV24 */
    Wo = mtREV24probmat; Lamo = mtREV24eigmat;
  }
  if (smodel==8){ /* HKY */
    *W=(double *) malloc((size_t) sizeof(double));
    *Lam=(double *) malloc((size_t) sizeof(double));
    **Lam=*Qk;
    if(*Qk < 0) 
      **Lam=tt2kappa(-(*Qk),fr);
    **W = muHKYf(**Lam,fr);
    allocWLam=2;
  }
  if(smodel==9){ /* LG */
    Wo = lgprobmat; Lamo = lgeigmat;
  }
  if (aamodel){  /* amino acid models */
    if (*fr > 0){ /* different frequencies than for model */
      *W = (double *) malloc((size_t) nchar*nchar*sizeof(double));
      *Lam = (double *) malloc((size_t) nchar*sizeof(double));
      fro = (double *) malloc((size_t) nchar*sizeof(double));
      makeprotfreqs(fro,Wo,Lamo);
      memcpy(*W,Wo,(size_t) nchar*nchar*sizeof(double));
      memcpy(*Lam,Lamo,(size_t) nchar*sizeof(double));
      chQ_freq(nchar,*W,*Lam,fr,fro);
      allocWLam=2;
    }
    else{
      *W=Wo; *Lam=Lamo;
      makeprotfreqs(fr,*W,*Lam);
    }
  }
  return(allocWLam);
}
int subst_model_set(int nchar, int smodel, double *fr, double *Qk, 
		    double (*(*pij))(int i, int j, double t, int nchar, 
				     double *fr, double *W, double *Lam),
		    double (*(*ppij))(int i, int j, double t, int nchar, 
				      double *fr, double *W, double *Lam),
		    double **W, double **Lam)
{
  /* IN:
   * nchar - number of character states
   * smodel - 0:JC, 1:F81, 2:F84, 3:GTR, 4:PAM, 5:JTT, 6:WAG, 7:mtREV24, 8:HKY,
   *          9:LG
   * Qk - rate matrix for GTR, K or Kappa for F84, HKY; not used otherwise
   *
   * IN/OUT:
   * fr - frequencies; reset for JC
   *      if *fr < 0 for aa models, model-based frequencies used
   * OUT:
   * pij,ppij - transition function and first derivative
   * W,Lam - contain parameters for substitution model
   *
   * RETURN:
   * 2,1 or 0 according to whether both W and Lam were allocated, just W or 
   * neither
   */
  subst_model_setp(smodel,pij,ppij);
  return(subst_model_setparam(nchar,smodel,fr,Qk,W,Lam));
}
int subst_model_setm(int nchar, int smodel, double *fr, double *Qk, 
		     void (*(*pm))(double t, int nchar, double *fr, 
				   double *W, double *Lam, int deriv, 
				   double *M, double *Mp, 
				   double *Mpp),
		     double **W, double **Lam)
{
  /* IN:
   * nchar - number of character states
   * smodel - 0:JC, 1:F81, 2:F84, 3:GTR, 4:PAM, 5:JTT, 6:WAG, 7:mtREV24, 8:HKY,
   *          9:LG
   * Qk - rate matrix for GTR, K or Kappa for F84, HKY, < 0 => ttratio; 
   * not used otherwise. 
   *
   * IN/OUT:
   * fr - frequencies; reset for JC, determined from Q for GTR
   *      if *fr < 0 for aa models, model-based frequencies used
   * OUT:
   * pm - transition matrix function 
   * W,Lam - contain parameters for substitution model
   *
   * RETURN:
   * 2,1 or 0 according to whether both W and Lam were allocated, just W or 
   * neither
   */
  subst_model_setpm(smodel,pm);
  return(subst_model_setparam(nchar,smodel,fr,Qk,W,Lam));
}
void trans_gen(int nchar, int ntaxa, double *utreec, double *fr, double *W, 
	       double *Lam, 		      
	       double pij(int i, int j, double t, int nchar, 
			  double *fr, double *W, double *Lam),
	       double ppij(int i, int j, double t, int nchar, 
			     double *fr, double *W, double *Lam),
	       double *M, double *Mp)
{
  /* IN:
   * utreec: the tree, nutreec format, 0 last edge
   * OUT:
   * M, Mp - transition matrices and first derivatives for each of the edges
   */
  int nchar2,join,kup,k,r,x,y;
  double tr;

  nchar2 = nchar*nchar;
  for(join = 0; join < ntaxa-1; join++){
    kup = (join < ntaxa-2)?2:1;
    for(k = 0; k < kup; k++){
      r = (int) utreec[k+join*4];
      tr = utreec[k+2+join*4];
      for(x = 0; x < nchar; x++)
	for(y = 0; y < nchar; y++){
	  M[x+y*nchar+r*nchar2]=pij(x,y,tr,nchar,fr,W,Lam);
	  Mp[x+y*nchar+r*nchar2]=ppij(x,y,tr,nchar,fr,W,Lam);
	}
      /* printf("r: %i, tr: %f\n", r, tr); */
      /* wmat(stdout,nchar,nchar,&M[r*nchar2]); */
/*       wmat(stdout,nchar,nchar,&M[r*nchar2]); */
      /* wmat(stdout,nchar,nchar,&Mp[r*nchar2]); */
    }
  }
}
void trans_gen_rate(int nchar, int ntaxa, double *utreec, 
		    int nrate, double *rate, double *wt,
		    double *fr, double *W, double *Lam, 		      
		    double pij(int i, int j, double t, int nchar, 
			       double *fr, double *W, double *Lam),
		    double ppij(int i, int j, double t, int nchar, 
				double *fr, double *W, double *Lam),
		    double *M, double *Mp)
{
  /* IN:
   * utreec: the tree, nutreec format, 0 last edge
   * OUT:
   * M, Mp - transition matrices and first derivatives for each edges x rate
   *         nchar x nchar x nbranch x nrate 
   *         derivatives are not multiplied by rate
   */
  int nchar2,nb,j,l,i;
  double *utreecn;
  
  nchar2 = nchar*nchar;
  nb = 2*ntaxa-3;
  utreecn = (double *) malloc((size_t) (ntaxa-1)*4*sizeof(double));
  memcpy(utreecn,utreec,(ntaxa-1)*4*sizeof(double));
  for(j = 0; j < nrate; j++){
    for(i = 0; i < ntaxa-1; i++)
      for(l = 0; l < 2; l++) utreecn[l+2+i*4] = rate[j]*utreec[l+2+i*4];
    trans_gen(nchar,ntaxa,utreecn,fr,W,Lam,pij,ppij,&M[j*nchar2*nb],
	      &Mp[j*nchar2*nb]);
  }
  free(utreecn);
  return;
}
void trans_gen_rates(int nchar, int ntaxa, double *utreec, 
		    int nrate, double *rate, double *wt,
		    double *fr, double *W, double *Lam, 		      
		    double pij(int i, int j, double t, int nchar, 
			       double *fr, double *W, double *Lam),
		    double ppij(int i, int j, double t, int nchar, 
				double *fr, double *W, double *Lam),
		     double *M, double *Mp){
  int i,j,nel;
  trans_gen_rate(nchar,ntaxa,utreec,nrate,rate,wt,fr,W,Lam,pij,ppij,M,Mp);
  nel=(2*ntaxa-3)*nchar*nchar;
  for(i=0; i<nrate; i++) 
    for(j=0; j<nel; j++) Mp[j+i*nel] *= rate[i];
  return;
}
void transm_rate(int smodel, int nchar, int ntaxa, double *utreec, 
		 int nrate, double *rate, double alpha,
		 double *fr, double *W, double *Lam, int deriv, 
		 double *M, double *Mp, double *Mpp){
  int nchar2,nb,j,join,kup,r,i,k;
  double tr,one,*Mpt,*Mppt;
  void (*pm)(double t, int nchar, double *fr, double *W, double *Lam, 
	     int deriv, double *M, double *Mp, double *Mpp);
  void (*pma)(double t, int nchar, double *fr, double *W, double *Lam, 
	      int deriv, double alpha, double *M, double *Mp, double *Mpp);
  if (alpha < 0)
    subst_model_setpm(smodel,&pm);
  if (alpha > 0) 
    subst_model_setpma(smodel,&pma);
  nchar2 = nchar*nchar;
  nb = 2*ntaxa-3;
  for(j = 0; j < nrate; j++){
    for(join = 0; join < ntaxa-1; join++){
      kup = (join < ntaxa-2)?2:1;
      for(k = 0; k < kup; k++){
	r = (int) utreec[k+join*4];
	tr = utreec[k+2+join*4]*rate[j];
	Mpt=&one; Mppt=&one;
	if(deriv > 0) Mpt=&Mp[r*nchar2+j*nchar2*nb];
	if(deriv==2||deriv==4) Mppt=&Mpp[r*nchar2+j*nchar2*nb];
	if(alpha < 0)
	  (*pm)(tr,nchar,fr,W,Lam,deriv,&M[r*nchar2+j*nchar2*nb],Mpt,Mppt);
	if(alpha > 0)
	  (*pma)(tr,nchar,fr,W,Lam,deriv,alpha,&M[r*nchar2+j*nchar2*nb],
		 Mpt,Mppt);	
	if(deriv==1 || deriv==2)
	  for(i = 0; i < nchar2; i++) 
	    Mp[i+r*nchar2+j*nchar2*nb] *= rate[j];
	if(deriv==2)
	  for(i = 0; i < nchar2; i++) 
	    Mpp[i+r*nchar2+j*nchar2*nb] *= rate[j]*rate[j];
      }}}
}
int eigenRealSym(double A[], int n, double Root[], double Offdiag[])
{
/* This finds the eigen solution of a real symmetrical matrix A[n*n].  In return, 
   A has the right vectors and Root has the eigenvalues. work[n] is the working space.
   The matrix is first reduced to a tridiagonal matrix using HouseholderRealSym(), 
   and then using the QL algorithm with implicit shifts.  

   Adapted from routine tqli in Numerical Recipes in C, with reference to LAPACK
   Ziheng Yang, 23 May 2001
*/
   int status=0;
   HouseholderRealSym(A, n, Root, Offdiag);
   status=EigenTridagQLImplicit(Root, Offdiag, n, A);
   EigenSort(Root, A, n);

   return(status);
}
void EigenSort(double d[], double U[], int n)
{
/* this sorts the eigen values d[] and rearrange the (right) eigen vectors U[]
*/
   int k,j,i;
   double p;

   for (i=0;i<n-1;i++) {
      p=d[k=i];
      for (j=i+1;j<n;j++)
         if (d[j] >= p) p=d[k=j];
      if (k != i) {
         d[k]=d[i];
         d[i]=p;
         for (j=0;j<n;j++) {
            p=U[j*n+i];
            U[j*n+i]=U[j*n+k];
            U[j*n+k]=p;
         }
      }
   }
}
void HouseholderRealSym(double a[], int n, double d[], double e[])
{
/* This uses HouseholderRealSym transformation to reduce a real symmetrical matrix 
   a[n*n] into a tridiagonal matrix represented by d and e.
   d[] is the diagonal (eigends), and e[] the off-diagonal.
*/
   int m,k,j,i;
   double scale,hh,h,g,f;

   for (i=n-1;i>=1;i--) {
      m=i-1;
      h=scale=0;
      if (m > 0) {
         for (k=0;k<=m;k++)
            scale += fabs(a[i*n+k]);
         if (scale == 0)
            e[i]=a[i*n+m];
         else {
            for (k=0;k<=m;k++) {
               a[i*n+k] /= scale;
               h += a[i*n+k]*a[i*n+k];
            }
            f=a[i*n+m];
            g=(f >= 0 ? -sqrt(h) : sqrt(h));
            e[i]=scale*g;
            h -= f*g;
            a[i*n+m]=f-g;
            f=0;
            for (j=0;j<=m;j++) {
               a[j*n+i]=a[i*n+j]/h;
               g=0;
               for (k=0;k<=j;k++)
                  g += a[j*n+k]*a[i*n+k];
               for (k=j+1;k<=m;k++)
                  g += a[k*n+j]*a[i*n+k];
               e[j]=g/h;
               f += e[j]*a[i*n+j];
            }
            hh=f/(h*2);
            for (j=0;j<=m;j++) {
               f=a[i*n+j];
               e[j]=g=e[j]-hh*f;
               for (k=0;k<=j;k++)
                  a[j*n+k] -= (f*e[k]+g*a[i*n+k]);
            }
         }
      } 
      else
         e[i]=a[i*n+m];
      d[i]=h;
   }
   d[0]=e[0]=0;

   /* Get eigenvectors */
   for (i=0;i<n;i++) {
      m=i-1;
      if (d[i]) {
         for (j=0;j<=m;j++) {
            g=0;
            for (k=0;k<=m;k++)
               g += a[i*n+k]*a[k*n+j];
            for (k=0;k<=m;k++)
               a[k*n+j] -= g*a[k*n+i];
         }
      }
      d[i]=a[i*n+i];
      a[i*n+i]=1;
      for (j=0;j<=m;j++) a[j*n+i]=a[i*n+j]=0;
   }
}
int EigenTridagQLImplicit(double d[], double e[], int n, double z[])
{
/* This finds the eigen solution of a tridiagonal matrix represented by d and e.  
   d[] is the diagonal (eigenvalues), e[] is the off-diagonal
   z[n*n]: as input should have the identity matrix to get the eigen solution of the 
   tridiagonal matrix, or the output from HouseholderRealSym() to get the 
   eigen solution to the original real symmetric matrix.
   z[n*n]: has the orthogonal matrix as output

   Adapted from routine tqli in Numerical Recipes in C, with reference to
   LAPACK fortran code.
   Ziheng Yang, May 2001
*/
   int m,j,iter,niter=30, status=0, i,k;
   double s,r,p,g,f,dd,c,b, aa,bb;

   for (i=1;i<n;i++) e[i-1]=e[i];  e[n-1]=0;
   for (j=0;j<n;j++) {
      iter=0;
      do {
         for (m=j;m<n-1;m++) {
            dd=fabs(d[m])+fabs(d[m+1]);
            if (fabs(e[m])+dd == dd) break;  /* ??? */
         }
         if (m != j) {
            if (iter++ == niter) {
               status=-1;
               break;
            }
            g=(d[j+1]-d[j])/(2*e[j]);

            /* r=pythag(g,1); */

            if((aa=fabs(g))>1)  r=aa*sqrt(1+1/(g*g));
            else                r=sqrt(1+g*g);

            g=d[m]-d[j]+e[j]/(g+SIGN(r,g));
            s=c=1;
            p=0;
            for (i=m-1;i>=j;i--) {
               f=s*e[i];
               b=c*e[i];

               /*  r=pythag(f,g);  */
               aa=fabs(f); bb=fabs(g);
               if(aa>bb)       { bb/=aa;  r=aa*sqrt(1+bb*bb); }
               else if(bb==0)             r=0;
               else            { aa/=bb;  r=bb*sqrt(1+aa*aa); }

               e[i+1]=r;
               if (r == 0) {
                  d[i+1] -= p;
                  e[m]=0;
                  break;
               }
               s=f/r;
               c=g/r;
               g=d[i+1]-p;
               r=(d[i]-g)*s+2*c*b;
               d[i+1]=g+(p=s*r);
               g=c*r-b;
               for (k=0;k<n;k++) {
                  f=z[k*n+i+1];
                  z[k*n+i+1]=s*z[k*n+i]+c*f;
                  z[k*n+i]=c*z[k*n+i]-s*f;
               }
            }
            if (r == 0 && i >= j) continue;
            d[j]-=p; e[j]=g; e[m]=0;
         }
      } while (m != j);
   }
   return(status);
}
double under_adj(int n, double *f){
  int i;
  double fmax,SMALL=1.0e-150;
  if(fabs(f[0])>SMALL) return(0.0); /* no adjustment needed */
  fmax=f[0];
  for(i=1; i<n; i++){
    if(fabs(f[i])>SMALL) return(0.0);
    if(fabs(f[i])>fmax) fmax=f[i];
  }
  if(fmax<=0){return(0);}
  for(i=0; i<n; i++) f[i] /= fmax;
  return(log(fmax));
}
void bprob_edge(int nchar, int ntaxa, int j, int r, double *M, double *fb,
		  double lfb, double *b, double *lb){
  int k,i;
  for(k=0; k<nchar; k++){
    b[k+r*nchar]=0;
    for(i=0; i<nchar; i++)
      b[k+r*nchar] += b[i+j*nchar]*M[i+k*nchar]*fb[k];
  }
  lb[r]=under_adj(nchar,&b[r*nchar])+lfb+lb[j];
  return;
}
void backward_probs(int nchar, int ntaxa, double *utreec, double *fr,
		    double *M, double *fb, double *lfb, double *f, double *lf,
		    double *b, double *lb){
  int i,j,r,rp,l,nc2;
  nc2=nchar*nchar;

  r=(int) utreec[(ntaxa-2)*4]; l=(int) utreec[1+(ntaxa-2)*4];
  for(i=0; i<nchar; i++) b[i+r*nchar]=fr[i]*fb[i+l*nchar];
  lb[r]=under_adj(nchar,&b[r*nchar])+lfb[l];
  for(i=0; i<nchar; i++) b[i+l*nchar]=fr[i]*fb[i+r*nchar];
  lb[l]=under_adj(nchar,&b[l*nchar])+lfb[r];
  
  for(j=ntaxa-3; j>=0; j--){
    r=(int) utreec[j*4]; l=(int) utreec[1+j*4];
    bprob_edge(nchar,ntaxa,j+ntaxa,r,&M[(j+ntaxa)*nc2],&fb[l*nchar],lfb[l],
	       b,lb);
    bprob_edge(nchar,ntaxa,j+ntaxa,l,&M[(j+ntaxa)*nc2],&fb[r*nchar],lfb[r],
	       b,lb);
  }
}
double fprob_edge(int nchar, int ntaxa, int r, int *s, double *M, double *f,
		  double lf, double *L){
  int i,k;

  if(r<ntaxa && s[r]>=nchar){
    for(i=0; i<nchar; i++) L[i]=1;
    return(0.0);
  }
  if(r<ntaxa && s[r]<nchar){
    for(i=0; i<nchar; i++) L[i]=M[i+s[r]*nchar];
    return(under_adj(nchar,L));
  }
  if(r>=ntaxa){
    for(i=0; i<nchar; i++)
    for(L[i]=0.0, k=0; k<nchar; k++) L[i] += M[i+k*nchar]*f[k];
    return(under_adj(nchar,L)+lf);
  }
}
void forward_probs(int nchar, int ntaxa, int *s, double *utreec, double *M,
		   double *f, double *lf, double *fb, double *lfb){
  int i,j,r,l,nc2;
  nc2=nchar*nchar;
  for(j=0; j<ntaxa-1; j++){
    r=(int) utreec[j*4]; l=(int) utreec[1+j*4];
    lfb[r]=fprob_edge(nchar,ntaxa,r,s,&M[r*nc2],&f[(r-ntaxa)*nchar],lf[r-ntaxa],
		      &fb[r*nchar]);
    lfb[l]=fprob_edge(nchar,ntaxa,l,s,&M[l*nc2],&f[(l-ntaxa)*nchar],lf[l-ntaxa],
		      &fb[l*nchar]);
    for(i=0; i<nchar; i++) f[i+j*nchar]=fb[i+r*nchar]*fb[i+l*nchar];
    lf[j]=under_adj(nchar,&f[j*nchar])+lfb[r]+lfb[l];
  }
}
double Mfn_tedge_site(int nchar, int s, double *b, double lb, double *Mf,
		      double *lMf){
  int i;
  double Mfs=0.0;
  if(s<nchar){
    for(i=0; i<nchar; i++) Mfs += b[i]*Mf[i+s*nchar];
    *lMf=lb;
  }
  else{
    *lMf=0.0;
  }
  return(Mfs);
}
double Mfn_iedge_site(int nchar, double *b, double lb,
		     double *f, double lf, double *Mf, double *lMf){
  int i,k;
  double Mfs=0.0;
  for(i=0; i<nchar; i++)
    for(k=0; k<nchar; k++) Mfs += b[i]*f[k]*Mf[i+k*nchar];
  *lMf=lb+lf;
  return(Mfs);
}
void lik_deriv_calc(int nchar, int ntaxa, int *s, double *fr, double *utreec,
		    double *M, double *Mp, 
		    double *f, double *lf, double *fb, double *lfb,
		    double *lp, double *llp){
  int i,k,nc2,r;
  double *b,*lb;
  nc2=nchar*nchar;
  b=(double *) malloc((size_t) (2*ntaxa-2)*nchar*sizeof(double));
  lb=(double *) malloc((size_t) (2*ntaxa-2)*sizeof(double));
  backward_probs(nchar,ntaxa,utreec,fr,M,fb,lfb,f,lf,b,lb);
  for(r=0; r<ntaxa; r++)
    lp[r]=Mfn_tedge_site(nchar,s[r],&b[r*nchar],lb[r],&Mp[r*nc2],&llp[r]);
  for(r=ntaxa; r<2*ntaxa-2; r++)
    lp[r]=Mfn_iedge_site(nchar,&b[r*nchar],lb[r],&f[(r-ntaxa)*nchar],
			 lf[r-ntaxa],&Mp[r*nc2],&llp[r]);
  free(b); free(lb);
  return;
}
double lik_deriv_site_1rate(int nchar, int ntaxa, int *s, double *fr,
			    double *utreec, double *M, double *Mp,
			    double *lp, double *llp, double *llik, int deriv){
  int i;
  double lik;
  double *f,*lf,*fb,*lfb;
  
  f = (double *) malloc((size_t) (ntaxa-1)*nchar*sizeof(double));
  lf = (double *) malloc((size_t) (ntaxa-1)*sizeof(double));
  fb = (double *) malloc((size_t) (2*ntaxa-2)*nchar*sizeof(double));
  lfb = (double *) malloc((size_t) (2*ntaxa-2)*sizeof(double));
  
  forward_probs(nchar,ntaxa,s,utreec,M,f,lf,fb,lfb);
  for(lik=0.0, i = 0; i < nchar; i++) lik += fr[i]*f[i+(ntaxa-2)*nchar];
  *llik = lf[ntaxa-2];
  /* printf("%.16e\n",log(lik)); */

  if(deriv>=1)
    lik_deriv_calc(nchar,ntaxa,s,fr,utreec,M,Mp,f,lf,fb,lfb,lp,llp);
  free(f); free(lf); free(fb); free(lfb);
  return(lik);
}
double lik_deriv_site_rt(int nchar, int ntaxa, int *s, double *fr,
			 double *utreec, int nrate, double *rate, double *wt, 
			 double *M, double *Mp, double *lliks,
			 double *lp, double *llp, int deriv, int rt){
  int r,i,k,ir,nchar2,nb,np,idx;
  double liks,*lik,*llik,*lpt,*llpt,te;

  if(((int) utreec[1+(ntaxa-2)*4])!=2*ntaxa-3){ /* testing */
    printf("utreec[1,p]=2p-3 needed for rooted routine\n");exit(1);
  }
  lik=(double *) malloc((size_t) nrate*sizeof(double));
  llik=(double *) malloc((size_t) nrate*sizeof(double));
  nchar2 = nchar*nchar;
  nb = 2*ntaxa-2; 
  if(deriv>0){
    np=(deriv>=2)?(nb+1):nb;
    lpt = (double *) malloc((size_t) np*nrate*sizeof(double));
    llpt = (double *) malloc((size_t) np*nrate*sizeof(double));
  }
  for(ir = 0; ir < nrate; ir++){
    idx=ir*nchar2*nb;
    lik[ir]=lik_deriv_site_1rate(nchar,ntaxa,s,fr,utreec,&M[idx],&Mp[idx],
				 &lpt[ir*np],&llpt[ir*np],&llik[ir],deriv);
    /* printf("%.5e %.5e\n",log(lik[ir]),llik[ir]); */
  }
  /* exit(0); */
  
  (*lliks)=llik[0];
  for(ir = 1; ir < nrate; ir++) if(llik[ir]>(*lliks)) (*lliks)=llik[ir];
  for(liks=0, ir=0; ir<nrate; ir++)
    liks += wt[ir]*lik[ir]*exp(llik[ir]-(*lliks));
  /* if(isnan(liks)){ */
  /*   for(ir=0; ir<nrate; ir++) printf("%.5e ",llik[ir]); printf("\n");} */
  
  if (deriv>0)
    for(k = 0; k < nb; k++){
      llp[k]=llpt[0];
      for(ir=1; ir<nrate; ir++) if(llpt[k+ir*np]>llp[k]) llp[k]=llpt[k+ir*np];
      lp[k]=0.0;
      for(ir=0; ir<nrate; ir++)
	lp[k] += wt[ir]*lpt[k+ir*np]*exp(llpt[k+ir*np]-llp[k]);
    }
  if(deriv>=2){
    llp[nb]=llp[0];
    for(k=1; k<nb; k++) if(llp[k]>llp[nb]) llp[nb]=llp[k];
    lp[nb]=0;
    for(ir=0; ir<nrate; ir++){ 
      for(te=0.0, k=0; k<ntaxa-1; k++)
	for(i=0; i<2; i++){
	  r=utreec[i+k*4]; 
	  te += utreec[2+i+k*4]*lpt[r+ir*np]*exp(llpt[r+ir*np]-llp[nb]);
	}
      lp[nb] += wt[ir]*rate[ir+nrate]*te/rate[ir];
    }
  }
  /* exit(1); */
  if (deriv>0){ free(lpt); free(llpt); }
  free(lik); free(llik);
  return(liks);
}
void Mijdp(int nchar, double t, double *fr, double *W, double *Lam,
		double *f,double *fd){
  static double elambdat[61];
  double q, p0, p1;
  int i, j, k;

  for(k = 0; k < nchar; k++) elambdat[k] = exp(t * Lam[k]);
  for(i = 0; i < nchar; i++)
    for(j = 0; j <= i; j++){
      p0 = 0.0; p1 = 0.0;
      for(k = 0; k < nchar; k++){
	q = W[i + k*nchar] * W[j+ k*nchar] * elambdat[k];
	p0 += q;
	p1 +=  (q * Lam[k]);
      }
      f[i + j*nchar] = p0;
      fd[i + j*nchar] = p1;
    }
  
  for(i=0; i<nchar; i++)
    for(j=0; j<i; j++){
      f[j+i*nchar] = f[i+j*nchar]/fr[j];
      fd[j+i*nchar] = fd[i+j*nchar]/fr[j];
    }
  
  for(i=0; i<nchar; i++)
    for(j=0; j<=i; j++){
      f[i+j*nchar] /= fr[i];
      fd[i+j*nchar] /= fr[i];
    }
}
void Mij_nod(int nchar, double t, double *fr, double *W, double *Lam,
	     double *f){
  double elambdat[61];
  double q, p0;
  int i, j, k;

  for(k = 0; k < nchar; k++) elambdat[k] = exp(t * Lam[k]);
  for(i = 0; i < nchar; i++)
    for(j = 0; j <= i; j++){      
      p0 = 0.0; 
      for(k = 0; k < nchar; k++){
	q = W[i + k*nchar] * W[j+ k*nchar] * elambdat[k];
	p0 += q;
      }
      f[i+j*nchar]=p0;
    }

  for(i=0; i<nchar; i++)
    for(j=0; j<i; j++) f[j+i*nchar] = f[i+j*nchar]/fr[j];
  
  for(i=0; i<nchar; i++)
    for(j=0; j<=i; j++) f[i+j*nchar] /= fr[i];
}
void Mmake(int nchar, int ntaxa, double *utreec, int nrate, double *rate, 
	   double *wt, double *fr, double *W, double *Lam, double *M, double *Mp){
  int nchar2,nb,ir,join,k,r,kup,i,j;
  double tr;
  nchar2 = nchar*nchar;
  nb = 2*ntaxa-3;
  for(ir = 0; ir < nrate; ir++)
    for(join = 0; join < ntaxa-1; join++){
      kup = (join < ntaxa-2)?2:1;
      for(k = 0; k < kup; k++){
	r = (int) utreec[k+join*4];
	tr = rate[ir]*utreec[k+2+join*4];
	Mijdp(nchar,tr,fr,W,Lam,&M[r*nchar2+ir*nchar2*nb],
	      &Mp[r*nchar2+ir*nchar2*nb]); 
	for(i = 0; i < nchar; i++)
	  for(j = 0; j < nchar; j++) 
	    Mp[i+j*nchar+r*nchar2+ir*nchar2*nb] *= rate[ir];
/* 	printf("\n"); */
/* 	  for(i = 0; i < nchar; i++){ */
/* 	    for(j = 0; j < nchar; j++) printf("%.5e ", M[i+j*nchar+ir*nchar2*nb]); */
/* 	    printf("\n"); */
/* 	  } */
      }
    }
}
int read_tree(FILE *fp, char *tstring)
{
  /* assumes 
   *   '[' and ']' delimit comments only
   *    '][' won't be used to end a comment and start a new one
   *   white space can be removed */
  int ltstring;
  char c;

  ltstring = 0;
  while((c = getc(fp)) != ';' && c!=EOF){ 
    if (c != ' ' && c != '\t' && c != '\r' && c != '\n'){ /* no white space */
      if (c == '[') /* start of comment; ignored */
	while((c = getc(fp)) != ']') ;
      else
	tstring[ltstring++]=c;
    }
  } 
  if(c==EOF){return(-1);}
  tstring[ltstring++]=';';
  tstring[ltstring++]='\0';
  return(ltstring);
}
double get_dlist(char *tstr, int ss, int de, int *ssn, int *ls, int *le)
{
  /* IN:
   * tstr - tree
   * ss - char position to start search
   * de - end position of dlist to be searched
   * OUT:
   * ssn - char position to start new search
   * ls - start pos of label; (ls-1) gives end of dlist found
   * le - end pos of label; 
   * RETURN: edge length (0.1 if not present) */

  char c,els[100]; /* note the limit on edge-length string */
  int lb=0,rb=0,lrb,i,j=0;
  double el=0.1;

  /* printf("%s\n",tstr); */
  /* printf("%i %i\n",ss,de); */
  /* for(i=ss; i<de; i++) putchar(tstr[i]); printf("\n"); */

  i=ss;
  lrb=ss-1;
  while(i <= de-1){
    if ((c=tstr[i]) == '(') lb++;
    if (c == ')'){rb++; lrb=i;}
    if (c == ',' && lb==rb) break;
    i++;
  }
  *ssn=i+1;
  *ls=lrb+1;
  i=lrb+1; 
  while((c=tstr[i]) != ':' && c != ','&& c != ')') i++;
  *le=i-1;
  if(c == ':'){
    while((c=tstr[++i]) != ',' && c != ')') els[j++]=c;
    els[j]='\0';
    el=atof(els);
  }
  return(el);
}
void underscore2blank(char *s){
  while(*s++ != '\0')
    if(*s=='_') *s=' ';
}
char **treecs(int ntaxa, char *tstr, char names[][11], double *utreec, 
	      int has_names){
  /* IN:
   * tstr - Newick tree w/o comments, whitespace and ; (see read_tree)
   * has_names = 1 if names are available
   * IN/OUT:
   * names - if has_names, used to determine ordering 0,...,ntaxa-1 of 
   *         terminal labels
   *         o/w gives names of the taxa labeled 0,...,ntaxa-1 in utreec
   * OUT:
   * utreec 
   * RETURN:
   * ilabels - internal nodel labels; (ntaxa-2); might be uninitialized
   *    not allocated as a contiguous block
   *
   * COMMENTS:
   * Based on observation that a root subtree is either
   *    (a) a leaf and has no `(' or ')' 
   * or (b) a dlist + label and el, and has equal numbers of `(' and ')' 
   *
   * violations of Newick standard
   * - quoted labels not allowed
   * - '[' and ']' only used to delimit comments
   * - limits of 10 non-null chars on leaf names
   * - underscores are not converted to blanks
   */

  int *ds; /* starting char positions of dlists */
  int *de; /* ending char positions of dlists */
  int cd=0,td=1; /* current and total dlists */
  int cj; /* current join */
  int cil; /* current internal label */
  int ctl=0; /* current terminal label */
  int *ul; /* utreec labels */
  int *ls,*le,*is_leaf,k,i,ss,K,j,ssn,ir,il;
  char **ilabels,tname[11];
  double *el;

  int tdo;

  el = (double *) malloc((size_t) ntaxa*sizeof(double));
  ds = (int *) malloc((size_t) 2*ntaxa*sizeof(int));
  de = (int *) malloc((size_t) 2*ntaxa*sizeof(int));
  ls = (int *) malloc((size_t) ntaxa*sizeof(int));
  le = (int *) malloc((size_t) ntaxa*sizeof(int));
  is_leaf = (int *) malloc((size_t) ntaxa*sizeof(int));
  ul = (int *) malloc((size_t) ntaxa*sizeof(int));
  ilabels = (char **)malloc((size_t) (ntaxa-2)*sizeof(char *));


  /* if(has_names) */
  /*   for(i = 0; i < ntaxa; i++) underscore2blank(names[i]); */

  for(i = 0; i < ntaxa; i++) el[i]=0;

  /* the initial dlist is the entire tree */
  *ds=0; *de=0; *is_leaf=0;
  while(tstr[*de] != '\0')(*de)++;
  while(tstr[*de] != ')') (*de)--; /* root label and edge ignored */

  cj=ntaxa-2;
  cil=2*ntaxa-3;
  while (cj >= 0){/* Main loop */

    /* obtain all dlists within the current dlist and update dlists */
    /* dlists indicated by ds[i], de[i], upper and lower char pos in tstr */
    ss=ds[cd]+1;
    K=0;
    while(ss <= de[cd]-1){
      el[K]=get_dlist(tstr,ss,de[cd],&ssn,&ls[K],&le[K]);
      is_leaf[K]=(ls[K] == ss)?1:0;
      if (!is_leaf[K]){
	ds[td]=ss;
	de[td]=ls[K]-1;
	td++;
      }
      ss=ssn;
      K++;
    }

    /* attach utreec labels, ul, to subtrees */
    for(i = 0; i < K-2; i++){ /* need to create labels for bifurcations */
      ilabels[cil-ntaxa]=(char *)malloc(sizeof(char));
      ilabels[cil-ntaxa][0]='\0';
      cil--;
    }
    for(i=0; i < K; i++){
      if (!is_leaf[i]){ 
	ilabels[cil-ntaxa]=(char *)malloc((le[i]-ls[i]+2)*sizeof(char));
	memcpy(ilabels[cil-ntaxa],&tstr[ls[i]],(le[i]-ls[i]+1)*sizeof(char));
	ilabels[cil-ntaxa][le[i]+1-ls[i]]='\0';
	/* underscore2blank(ilabels[cil-ntaxa]); */
	ul[i]=cil;
	cil--;
      }
      if (is_leaf[i] && !has_names){
	memcpy(names[ctl],&tstr[ls[i]],(le[i]-ls[i]+1)*sizeof(char));
	for(k=le[i]+1-ls[i]; k < 10; k++) names[ctl][k]=' ';
	names[ctl][10]='\0';
	/* underscore2blank(names[ctl]); */
	ul[i]=ctl;
	ctl++;
      }
      if (is_leaf[i] && has_names){
	memcpy(tname,&tstr[ls[i]],(le[i]-ls[i]+1)*sizeof(char));
	for(k=le[i]+1-ls[i]; k < 10; k++) tname[k]=' ';
	tname[10]='\0';
	/* underscore2blank(tname); */
	for(j = 0; j < ntaxa; j++){
	  if (strncmp(tname,names[j],11)==0) break;
	}
	if(j >= ntaxa){
	  printf("treecs: unable to find %s among taxa\n",tname); exit(0);}
	ul[i]=j;
      } 
    }/* end of attach utreec labels ... */

    /* for(k=cd; k<td; k++){ */
    /*   for(j = ds[k]; j <= de[k]; j++) putchar(tstr[j]); printf("\n"); */
    /* } */
    
    /* printf("current subtree %i\n",ntaxa+cj); */
    /* for(j = ds[cd]; j <= de[cd]; j++) putchar(tstr[j]); printf("\n"); */
    /* printf("number of subtrees: %i\n",K); */
    /* labels for subtrees in ul. Last label corresponds to last dlist.
     * Next one to the second last but this will not be second last ul
     * unless it was not a taxa ...*/
    /* tdo=td-1;  */
    /* for(i=K-1; i >= 0 ; i--){ */
    /*   printf("%i %f ",ul[i],el[i]); */
    /*   if(!is_leaf[i]){ */
    /* 	for(j = ds[tdo]; j <= de[tdo]; j++) putchar(tstr[j]); */
    /* 	tdo--; */
    /*   } */
    /*   printf("\n"); */
    /* } */

    /* adjust labels of existing internal edges for multifurcation */
    if(K>2)
      for(i=cj+1; i<ntaxa-1; i++)
	for(k=0; k<2; k++) 
	  if(utreec[k+i*4]>ntaxa && utreec[k+i*4]<ntaxa+cj) utreec[k+i*4] -= K-2;

    /* add root subtrees to utreec */
    for(i = 0; i < K-2; i++){
      utreec[cj*4]=ul[i]; utreec[1+cj*4]=cj-1+ntaxa; 
      utreec[2+cj*4]=el[i]; utreec[3+cj*4]=0.0;
      cj--;
    }
    if (ul[K-2] < ul[K-1]){
      ir=K-2;il=K-1;
    }
    else{
      ir=K-1;il=K-2;
    }
    utreec[cj*4]=ul[ir]; utreec[2+cj*4]=el[ir];
    utreec[1+cj*4]=ul[il]; utreec[3+cj*4]=el[il];
    cj--;
    cd++;
    
    /* for(i=cj+1; i<ntaxa-1; i++) */
    /*   printf("%i %i %i %f %f\n",i+ntaxa,(int) utreec[i*4],(int) utreec[1+i*4], */
    /* 	     utreec[2+i*4],utreec[3+i*4]); */
    /* printf("\n"); */

  } /* end of Main loop */
  /* pr_utreec(stdout,ntaxa,utreec); */
  /* for(i = 0; i < ntaxa; i++) */
  /*   printf("%s\n",names[i]); */
  
  free(is_leaf); 
  free(ul); 
  free(le); 
  free(ls); 
  free(de); 
  free(ds); 
  free(el); 
  return(ilabels);
}
void treecsnl(int ntaxa, char *tstr, char names[][11], double *utreec, 
	      int has_names){
  char **ilabels; 
  int j;
  ilabels=treecs(ntaxa, tstr, names, utreec, has_names);
  for(j=0; j<ntaxa-2; j++)free(ilabels[j]);
  free(ilabels);
}
char **read_treecs(FILE *treefile, int ntaxa, char names[][11], double *utreec, 
		   int has_names){
  /* IN:
   * treefile - currently reading a new tree
   * IN/OUT:
   * names - if has_names, used to determine ordering 0,...,ntaxa-1 of 
   *         terminal labels
   *         o/w gives names of the taxa labeled 0,...,ntaxa-1 in utreec
   * OUT:
   * utreec 
   * RETURN:
   * ilabels - internal nodel labels; (ntaxa-2); each might only have '\0'
   *    not allocated as a contiguous block
   *
   * COMMENTS:
   * Based on observation that a root subtree is either
   *    (a) a leaf and has no `(' or ')' *
   * or (b) a dlist + label and el, and has equal numbers of `(' and ')' 
   *
   * violations of Newick standard
   * - quoted labels not allowed
   * - '[' and ']' only used to delimit comments
   * - limits of 10 non-null chars on leaf names 
   * - underscores are not converted to blanks
   */
  char *tstring;
  char **ilabels;
  tstring = (char *) malloc((size_t) 100000*sizeof(char));
  read_tree(treefile,tstring);
  /* printf("%s\n",tstring); */
  ilabels=treecs(ntaxa,tstring,names,utreec,has_names);
  free(tstring);
  return(ilabels);
}
void read_treecsnl(FILE *treefile, int ntaxa, char names[][11], double *utreec, 
		   int has_names){
  char *tstring;
  /* char tstring[10000]; */
  tstring = (char *) malloc(100000*sizeof(char));
  read_tree(treefile,tstring);
  /* printf("%s\n",tstring); */
  treecsnl(ntaxa,tstring,names,utreec,has_names);
  free(tstring);
}
int ntrees_treefile(FILE *treefile){
  int ntrees;
  char c;
  ntrees=0;
  while((c=getc(treefile)) != EOF)
    if(c==';') ntrees++;
  rewind(treefile);
  return(ntrees);
}
int sp_char(char c){
  return((c == '(' || c == ')' || c == ',' || c == ':' || c == ';'));
}
void ch_name(int lstring, char *tstring, int i){ 
  /* change name given by the first lstring chars of tstring to integer label i */
  char label[11];
  int tstl, llen;
  
  /* shorten the string and get rid of the name */
  tstl = 0;
  while(tstring[tstl+lstring] != '\0'){
    tstring[tstl] = tstring[tstl+lstring];
    tstl++;
  }
  tstring[tstl] = '\0';
  
  sprintf(label, "%i", i); /* i to char label */
  llen = strlen(label);   /* number of digits */
  
  /* make room for integer label; more efficient: combine this and shorten above */
  while(tstl >= 0){
    tstring[tstl+llen] = tstring[tstl];
    tstl--;
  }
  /* place the integer label where name used to be */
  for(tstl = 0; tstl < llen; tstl++)
    tstring[tstl] = label[tstl];
}
int has_bl(char *tstring)
{
  while(*tstring != ';')
    if(*tstring == ':') /* signifies start of bl */
      return(1);
    else
      tstring++;
  return(0); /* must not be bl if we got this far */
}
void ch_labels_nbl(int ntaxa, char *tstring, char names[][11]){
  /* change all names in tstring to integer labels (no branch lengths)*/
  char taxname[11];
  int ltstring, i;

  while(*tstring != ';'){
    while(*tstring == ')' || *tstring == '(' || *tstring == ','){
      tstring++;
    }
    if(*tstring == ';') break;  /* end of tstring, no more names */

    ltstring = 0; /* beginning of new name */
    while(*tstring != ')' && *tstring != '(' && *tstring != ','){
      taxname[ltstring] = *tstring;
      tstring++;
      ltstring++;
    }

    for(i=ltstring; i < 10; i++) taxname[i] = ' '; /* pad w/ blanks */
    taxname[10] = '\0'; /* name is in taxname */
    for(i = 0; i < ltstring; i++) tstring--; /* go to start of name */
    
    for(i = 0; i < ntaxa; i++){            /* check which taxa it is */
      if (strncmp(taxname, names[i], 10) == 0){ /* i is label for taxa */
	ch_name(ltstring, tstring, i);	  
	break;
      }
    }
    /* go to end of name */
    while(*tstring != '(' && *tstring != ')' && *tstring != ',') tstring++;
  }
}
void ch_labels(int ntaxa, char *tstring, char names[][11]){
  /* change all names in tstring to integer labels */
  char taxname[11];
  int lstring, i;
  
  while(*tstring != ';'){
    while(*tstring != ':' && *tstring != ';'){ /* name always before : */
      tstring++;
    }
    if (*tstring == ';') break;  /* end of tstring, no more names */
    tstring--; lstring = 0;      /* end of a possible name */
    while(!sp_char(*tstring)){   /* find the beginning */
      tstring--; lstring++;
    }
    if (*tstring == '(' || *tstring == ','){ /* condition for a name */
      tstring++;
      strncpy(taxname, tstring, lstring);    /* name is in taxname */
      for(i=lstring; i < 10; i++) taxname[i] = ' '; /* pad w/ blanks */
      taxname[10] = '\0';

/*       for(i = 0; i < lstring; i++) putchar(taxname[i]); printf("\n");  */
      for(i = 0; i < ntaxa; i++){            /* check which taxa it is */
/*  	printf("%s\n", names[i]);  */
	if (strncmp(taxname, names[i], 10) == 0){ /* i is label for taxa */
	  ch_name(lstring, tstring, i);	  
	  break;
	}
      }
/*       printf("%s\n\n", tstring); */ 
    }
    while(*tstring != ':') tstring++; /* get back to : */
    tstring++; /* process the rest of string */
  }
}
int is_wsp(char c){
  if(c==' ' || c=='\t' || c=='\n' || c=='\t') return(1);
  return(0);
}
void rmblntree(char *tstring, int rwsp){
  /* I: wsp=1 => remove whitespace */
  int s=0,e=0;
  while(tstring[e]!=';'){
    if(tstring[e]==':')
      while(tstring[e]!=',' && tstring[e]!=')') e++;
    if(!rwsp || (tstring[e]!=' ' && tstring[e]!='\t'&& tstring[e]!='\r' && 
		 tstring[e]!='\n'))
      tstring[s++]=tstring[e];
    e++;
  }
  tstring[s++]=';';
  tstring[s]='\0';
}
```
