## Supplementary material for "Modeling site-and-branch-heterogeneity with GFmix": alpha_est_mix_rtf.h

```
#include <stdio.h>
#include <stdlib.h>
#include <math.h>
#include <string.h>
#ifndef PI
   #define PI 3.14159265358979323846
#endif
enum {
  alanine = 0, arginine, asparagine, aspartic, cysteine, 
  glutamine, glutamic, glycine, histidine, isoleucine,
  leucine, lysine, methionine, phenylalanine, proline,
  serine, threonine, tryptophan, tyrosine, valine
};
#define InverseCDFGamma(prob,alpha,beta) InverseCDFChi2(prob,2.0*(alpha))/(2.0*(beta))
#define SIGN(a,b) ((b) >= 0.0 ? fabs(a) : -fabs(a))
void Mmake_rt_mix(int nchar, int ntaxa, double *utreec,
		  int nclass, double *fr, double *S,
		  int nrate, double *rated, 
		  double *M, double *Mp, int deriv);
double fmix_garp_lnld(int nchar, int ntaxa, int npatt, int *seq, int *count,
		      double *utreec,
		      int nclass, double *fr, double *wtc,
		      double *S, int nrate, double alpha, double *lp,
		      int deriv, double *liks);
void fmix_garp_param2x(int ntaxa, int nclass, double *utreec, double *wtc,
		       double alpha, double *x);
void fmix_garp_x2param(int ntaxa, int nclass, double *x, double *utreec,
		       double *wtc, double *alpha);
FILE *fopene(char *filename, char *rw, char *err);
char get_param(int *k, char **arg, char *value);
char ctl_paraml(FILE *fp, char *opt, char *value, int *is_option);
int pmodel2smodel(int model, char *aaRmat, int nchar);
int std_paramf(FILE *ctlfile, FILE **treefile, FILE **utreecfile, FILE **seqfile,
	   FILE **Qfile, FILE **ratefile, char *aaRmat, 
	   int *model, int *nchar, int *nsite, int *ntaxa, int *nrate, 
	   int *ntrees, double *alpha, double *kappa);
void rmgap(int numsp, int *nsite, char *seqc);
void letter(int numsp, int nsite, char *seq);
int l2i(char c);
char i2l(int i);
char i2lp(int a);
int l2ip(char c);
int rseqf(FILE *fp, char *seq);
char ignore_whitespace_seqfile(FILE *infile);
int rinterleave_block(FILE *infile, int numsp, int nsite, char name[][11], 
		       char *seq, int *initb);
void rinterleavef_nolim(FILE *infile, const int *numsp, const int *nsite, 
			char name[][11], char *seq);
void rinterleave(int *numsp, int *nsite, char name[][11], char *seq);
void rinterleavef(FILE *infile, int *numsp, int *nsite, char name[][11], 
		  char *seq);
int char_freq(int nchar, int numsp, int nsite, int *seq, double *freq);
int char_freq_count(int nchar, int numsp, int endsite, int *seq, int *count, 
		     double *freq);
int patt_freq(int numsp, int nsite, int *seq, int *count);
int patt_freq_locs(int numsp, int nsite, int *seq, int *count, int *locs);
void pr_utreec(FILE *fp, int ntaxa, double *utreec);
char *read_seqc(FILE *fp, int *ntaxa, int *nsite, char (**name)[11]);
int *read_seq(FILE *fp, int nchar, int rm_gap, int *ntaxa, int *nsite, 
	      char (**name)[11]);
int DiscreteGamma (double freqK[], double rK[], double alpha, double beta, int K, int dgammamean);
int DiscreteGammad(int K, double alpha, double freqK[], double rK[]);
int DiscreteGammada(int K, double alpha, double freqK[], double rK[], double h);
void error2 (char * message);
long factorial (int n);
double LnGamma (double x);
double InverseCDFNormal (double prob);
double InverseCDFChi2 (double prob, double v);
double IncompleteGamma (double x, double alpha, double ln_gamma_alpha);
double trigamma(double x);
double digamma(double x);
int digami(double x, double p, double *d);
void makeprotfreqs(double *fr, double *W, double *Lam);
void Mijp(double t, double *fr, double *W, double *Lam, double *f);
void Mijpd(double t, double *fr, double *W, double *Lam, double *M, double *Mp, 
	   double *Mpp);
void WLamfr2exchange(int nchar, double *W, double *Lam, double *fr, double *R);
void chQ_freq(int nbin, double *W, double *Lam, double *fr, double *fro);
double int_elamrt(double t, double lam, double alpha, int deriv, double *de);
double pijJCg(int x, int y, double t, int nchar);
double pijJCg_gen(int x, int y, double t, int nchar, double *fr, double *W, 
		  double *Lam);
double ppijJCg(int x, int y, double t, int nchar);
double ppijJCg_gen(int x, int y, double t, int nchar, double *fr,
		   double *W, double *Lam);
double pp2ijJCg(int x, int y, double t, int nchar);
void pJCm(double t, int nchar, double *fr, int deriv, 
	    double *M, double *Mp, double *Mpp);
void pJCm_gen(double t, int nchar, double *fr, double *W, double *Lam, 
		int deriv, double *M, double *Mp, double *Mpp);
void pJCma(double t, int nchar, double *fr, int deriv, double alpha,
	   double *M, double *Mp, double *Mpp);
void pJCma_gen(double t, int nchar, double *fr, double *W, double *Lam, 
	       int deriv, double alpha, double *M, double *Mp, double *Mpp);
double muF81f(int nchar, double *fr);
double pijF81(int i, int j, double t, int nchar, double mu, double *fr);
double pijF81_gen(int i, int j, double t, int nchar, double *fr, double *mu,
		  double *Lam);
double ppijF81(int i, int j, double t, int nchar, double mu, double *fr);
double ppijF81_gen(int i, int j, double t, int nchar, double *fr, double *mu,
		   double *Lam);
void pF81m(double t, int nchar, double *fr, double mu, int deriv, 
	   double *M, double *Mp, double *Mpp);
void pF81m_gen(double t, int nchar, double *fr, double *mu, double *Lam, 
		 int deriv, double *M, double *Mp, double *Mpp);
void pF81ma(double t, int nchar, double *fr, double mu, int deriv, double alpha,
	   double *M, double *Mp, double *Mpp);
void pF81ma_gen(double t, int nchar, double *fr, double *mu, double *Lam, 
		int deriv, double alpha, double *M, double *Mp, double *Mpp);
int is_transitionf(int i, int j);
double muF84f(double K, double *fr);
double pijF84(int i, int j, double t, double mu, double K, double *fr);
double pijF84_gen(int i, int j, double t, int nchar, double *fr, double *mu,
		  double *K);
double ppijF84(int i, int j, double t, double mu, double K, double *fr);
double ppijF84_gen(int i, int j, double t, int nchar, double *fr, double *mu,
		  double *K);
double p2ijF84(int i, int j, double t, double mu, double K, double *fr);
double p2ijF84_gen(int i, int j, double t, int nchar, double *fr, double *mu,
		  double *K);
void pF84m(double t, double mu, double K, double *fr, int deriv, 
	   double *M, double *Mp, double *Mpp);
void pF84m_gen(double t, int nchar, double *fr, double *mu, double *K, 
	       int deriv, double *M, double *Mp, double *Mpp);
void pF84ma(double t, double mu, double K, double *fr, int deriv, double alpha,
	    double *M, double *Mp, double *Mpp);
void pF84ma_gen(double t, int nchar, double *fr, double *mu, double *K, 
		int deriv, double alpha, double *M, double *Mp, double *Mpp);
double tt2K(double R, double *fr);
double K2tt(double K, double *fr);
double muHKYf(double kappa, double *fr);
double pijHKY(int i, int j, double t, double mu, double kappa, double *fr);
double pijHKY_gen(int i, int j, double t, int nchar, double *fr, double *mu,
		  double *kappa);
double ppijHKY(int i, int j, double t, double mu, double kappa, double *fr);
double ppijHKY_gen(int i, int j, double t, int nchar, double *fr, double *mu,
		   double *kappa);
double p2ijHKY(int i, int j, double t, double mu, double kappa, double *fr);
double p2ijHKY_gen(int i, int j, double t, int nchar, double *fr, double *mu,
		   double *kappa);
void pHKYm(double t, int nchar, double *fr, double mu, double kappa,
	   int deriv, double *M, double *Mp, double *Mpp);
void pHKYm_gen(double t, int nchar, double *fr, double *mu, double *kappa, 
	       int deriv, double *M, double *Mp, double *Mpp);
void pHKYma(double t, int nchar, double *fr, double mu, double kappa,
	    int deriv, double alpha, double *M, double *Mp, double *Mpp);
void pHKYma_gen(double t, int nchar, double *fr, double *mu, double *kappa, 
	       int deriv, double alpha, double *M, double *Mp, double *Mpp);
double tt2kappa(double R, double *fr);
double kappa2tt(double kappa,  double *fr);
void rmnegpv(int m, double *pv, int incp);
void rmnegtrans(int m, double *P);
void Q2frWLam(int nbin, double *Q, double *fr, double *W, double *Lam);
void GetWLam(int nchar,double *S, double *fr, double *Q, 
	     double *W, double *Lam);
double pijGTR(int i, int j, double t, int nchar, double *fr, double *W, 
	      double *Lam);
double ppijGTR(int i, int j, double t, int nchar, double *fr, double *W, 
	       double *Lam);
double p2ijGTR(int i, int j, double t, int nchar, double *fr, double *W, 
	       double *Lam);
void pGTRm(double t, int nchar, double *fr, double *W, double *Lam, 
	   int deriv, double *M, double *Mp, double *Mpp);
void pGTRma(double t, int nchar, double *fr, double *W, double *Lam, 
	    int deriv, double alpha, double *M, double *Mp, double *Mpp);
void LG_exchangeability(double *S);
void JTT_exchangeability(double *S);
void WAG_exchangeability(double *S);
int subst_model_setp(int smodel,
		     double (*(*pij))(int i, int j, double t, int nchar,
				     double *fr, double *W, double *Lam),
		     double (*(*ppij))(int i, int j, double t, int nchar,
				       double *fr, double *W, double *Lam));
int subst_model_setpm(int smodel,
		      void (*(*pm))(double t, int nchar, double *fr, 
				    double *W, double *Lam, int deriv, 
				    double *M, double *Mp, 
				    double *Mpp));
int subst_model_setpma(int smodel,
		       void (*(*pm))(double t, int nchar, double *fr, 
				     double *W, double *Lam, int deriv, 
				     double alpha,
				     double *M, double *Mp, double *Mpp));
int subst_model_setparam(int nchar, int smodel, double *fr, double *Qk,
			 double **W, double **Lam);
int subst_model_set(int nchar, int smodel, double *fr, double *Qk, 
		    double (*(*pij))(int i, int j, double t, int nchar, 
				     double *fr, double *W, double *Lam),
		    double (*(*ppij))(int i, int j, double t, int nchar, 
				      double *fr, double *W, double *Lam),
		    double **W, double **Lam);
int subst_model_setm(int nchar, int smodel, double *fr, double *Qk, 
		     void (*(*pm))(double t, int nchar, double *fr, 
				   double *W, double *Lam, int deriv, 
				   double *M, double *Mp, 
				   double *Mpp),
		     double **W, double **Lam);
void trans_gen(int nchar, int ntaxa, double *utreec, double *fr, double *W, 
	       double *Lam, 		      
	       double pij(int i, int j, double t, int nchar, 
			  double *fr, double *W, double *Lam),
	       double ppij(int i, int j, double t, int nchar, 
			     double *fr, double *W, double *Lam),
	       double *M, double *Mp);
void trans_gen_rate(int nchar, int ntaxa, double *utreec, 
		    int nrate, double *rate, double *wt,
		    double *fr, double *W, double *Lam, 		      
		    double pij(int i, int j, double t, int nchar, 
			       double *fr, double *W, double *Lam),
		    double ppij(int i, int j, double t, int nchar, 
				double *fr, double *W, double *Lam),
		    double *M, double *Mp);
void trans_gen_rates(int nchar, int ntaxa, double *utreec, 
		    int nrate, double *rate, double *wt,
		    double *fr, double *W, double *Lam, 		      
		    double pij(int i, int j, double t, int nchar, 
			       double *fr, double *W, double *Lam),
		    double ppij(int i, int j, double t, int nchar, 
				double *fr, double *W, double *Lam),
		     double *M, double *Mp);
void transm_rate(int smodel, int nchar, int ntaxa, double *utreec, 
		 int nrate, double *rate, double alpha,
		 double *fr, double *W, double *Lam, int deriv, 
		 double *M, double *Mp, double *Mpp);
int eigenRealSym(double A[], int n, double Root[], double Offdiag[]);
void EigenSort(double d[], double U[], int n);
void HouseholderRealSym(double a[], int n, double d[], double e[]);
int EigenTridagQLImplicit(double d[], double e[], int n, double z[]);
double under_adj(int n, double *f);
void bprob_edge(int nchar, int ntaxa, int j, int r, double *M, double *fb,
		  double lfb, double *b, double *lb);
void backward_probs(int nchar, int ntaxa, double *utreec, double *fr,
		    double *M, double *fb, double *lfb, double *f, double *lf,
		    double *b, double *lb);
double fprob_edge(int nchar, int ntaxa, int r, int *s, double *M, double *f,
		  double lf, double *L);
void forward_probs(int nchar, int ntaxa, int *s, double *utreec, double *M,
		   double *f, double *lf, double *fb, double *lfb);
double Mfn_tedge_site(int nchar, int s, double *b, double lb, double *Mf,
		      double *lMf);
double Mfn_iedge_site(int nchar, double *b, double lb,
		     double *f, double lf, double *Mf, double *lMf);
void lik_deriv_calc(int nchar, int ntaxa, int *s, double *fr, double *utreec,
		    double *M, double *Mp, 
		    double *f, double *lf, double *fb, double *lfb,
		    double *lp, double *llp);
double lik_deriv_site_1rate(int nchar, int ntaxa, int *s, double *fr,
			    double *utreec, double *M, double *Mp,
			    double *lp, double *llp, double *llik, int deriv);
double lik_deriv_site_rt(int nchar, int ntaxa, int *s, double *fr,
			 double *utreec, int nrate, double *rate, double *wt, 
			 double *M, double *Mp, double *lliks,
			 double *lp, double *llp, int deriv, int rt);
void Mijdp(int nchar, double t, double *fr, double *W, double *Lam,
		double *f,double *fd);
void Mij_nod(int nchar, double t, double *fr, double *W, double *Lam,
	     double *f);
void Mmake(int nchar, int ntaxa, double *utreec, int nrate, double *rate, 
	   double *wt, double *fr, double *W, double *Lam, double *M, double *Mp);
int read_tree(FILE *fp, char *tstring);
double get_dlist(char *tstr, int ss, int de, int *ssn, int *ls, int *le);
void underscore2blank(char *s);
char **treecs(int ntaxa, char *tstr, char names[][11], double *utreec, 
	      int has_names);
void treecsnl(int ntaxa, char *tstr, char names[][11], double *utreec, 
	      int has_names);
char **read_treecs(FILE *treefile, int ntaxa, char names[][11], double *utreec, 
		   int has_names);
void read_treecsnl(FILE *treefile, int ntaxa, char names[][11], double *utreec, 
		   int has_names);
int ntrees_treefile(FILE *treefile);
int sp_char(char c);
void ch_name(int lstring, char *tstring, int i);
int has_bl(char *tstring);
void ch_labels_nbl(int ntaxa, char *tstring, char names[][11]);
void ch_labels(int ntaxa, char *tstring, char names[][11]);
int is_wsp(char c);
void rmblntree(char *tstring, int rwsp);
#define NCHAR 20
```
