## Supplementary material for "Modeling site-and-branch-heterogeneity with GFmix": Makefile

```
.SUFFIXES: .c .f .o .a  .def .exp .dll .exe 

CFLAGS=-O3
CC=gcc

SOURCES=$(wildcard -f *.c *.f)
OBJS=$(foreach i,$(SOURCES),$(basename $i).o)

all: treecns rert alpha_est_mix_rt 

treecns: treecns.o treecnsf.o
	$(CC) $(CFLAGS) -o treecns treecns.o treecnsf.o -lm

rert: rert.o rertf.o
	$(CC) $(CFLAGS) -o rert rert.o rertf.o -lm

alpha_est_mix_rt: alpha_est_mix_rt.o alpha_est_mix_rtf.o 
	$(CC) $(CFLAGS) -o alpha_est_mix_rt alpha_est_mix_rt.o \
	alpha_est_mix_rtf.o -lm

.c.o:
	$(CC)  $(CFLAGS) $($*-CFLAGS) -c $< -o $@
```
