## Supplementary material for "Modeling site-and-branch-heterogeneity with GFmix": rert.c

```
#include "rertf.h"
int main(int argc, char **argv)
{
  int ntaxa,r,i,fsi;
  double *utreec,*nutreec,tmp;
  ntaxa=atoi(argv[--argc]);
  r=atoi(argv[--argc]);
  utreec=(double *) malloc((size_t) 4*(ntaxa-1)*sizeof(double));
  nutreec=(double *) malloc((size_t) 4*(ntaxa-1)*sizeof(double));
  for(i = 0; i < 4*(ntaxa-1); i++) fsi=scanf("%lf",&utreec[i]);
  /* for(i = 0; i < ntaxa-1; i++) printf("%i %i %i\n", i, (int) utreec[i*4], (int) utreec[1+i*4]); */
  rert(r,ntaxa,utreec,nutreec);
  /* for(i = 0; i < ntaxa-1; i++) printf("%i %i %i\n", i, (int) nutreec[i*4], (int) nutreec[1+i*4]); */
  relab_int(ntaxa,nutreec,utreec);
  for(i=0; i<ntaxa-1; i++) /* r < l */
    if(nutreec[i*4]>nutreec[1+i*4]){
      r=nutreec[1+i*4]; tmp=nutreec[3+i*4];
      nutreec[1+i*4]=nutreec[i*4]; nutreec[3+i*4]=nutreec[2+i*4];
      nutreec[2+i*4]=tmp; nutreec[i*4]=r;
    }
  /* for(i = 0; i < 4*(ntaxa-1); i++) printf("%f ", utreec[i]); */
  for(i = 0; i < ntaxa-1; i++) printf("%i %i %f %f\n", (int) nutreec[i*4], (int) nutreec[1+i*4], nutreec[2+i*4], nutreec[3+i*4]);
  return(0);
}
```
