## Supplementary material for "Modeling site-and-branch-heterogeneity with GFmix": rertf.c

```
#include "rertf.h"
void parent_branch(int r, int ntaxa, double *utreec, int *p, int *l, 
		   double *bl, double *br)
{
  int i,k,done=0;
  i=r-ntaxa+1; /* only need to check above r */
  if (i < 0) i=0;
  while(!done){
    for(k = 0; k < 2; k++)
      if(((int) utreec[k+i*4])==r){
	*p=i+ntaxa;
	*br=utreec[k+2+i*4];
	*l=utreec[1-k+i*4];
	*bl=utreec[1-k+2+i*4];
	done=1;
      }
    i++;
  }
}
void relab_int(int ntaxa, double *utreec, double *nutreec)
{
  /* assumes r < l */
  int l,la,j,k,r;
 
  for(l = 0; l < 4; l++) nutreec[l+(ntaxa-2)*4] = utreec[l+(ntaxa-2)*4];
  la=2*ntaxa-3;
  /* printf("%i %i\n", (int) nutreec[(ntaxa-2)*4], (int) nutreec[1+(ntaxa-2)*4]); */
  
  for(j=(ntaxa-2); j >= 0; j--)
    for(k = 1; k >= 0; k--){
      r=nutreec[k+j*4];
      if (r >= ntaxa){
	for(l = 0; l < 4; l++) 
	  nutreec[l+(la-ntaxa)*4]=utreec[l+(r-ntaxa)*4];
	nutreec[k+j*4]=la;
	/* printf("j: %i %i\n",(int) nutreec[(ntaxa-2)*4],(int) nutreec[1+(ntaxa-2)*4]); */
	la--;
      }}
}
void rert(int r, int ntaxa, double *utreec, double *nutreec)
{
  /* IN:
   * r - label of branch tree is to be rooted at
   * ntaxa, utreec
   * OUT:
   * nutreec
   *
   * see software notes 2009-06-26
   */
  int p,l,k,kup,j,c,nc;
  double br,bl;
  
  /* If parent node of r is the root, no re-rooting is required. */
  parent_branch(r,ntaxa,utreec,&p,&l,&bl,&br);
  if (p == 2*ntaxa-2){
    for(j = 0; j < 4*(ntaxa-1); j++) nutreec[j]=utreec[j];
    return;
  }

  /* The last row of nutreec will be r 2*ntaxa-3; edge length split in half */
  c=ntaxa-2; 
  nutreec[c*4]=r; nutreec[1+c*4]=2*ntaxa-3;
  nutreec[2+c*4]=br/2; nutreec[3+c*4]=br/2;
  /* printf("%i %i %i\n", c, (int) nutreec[c*4], (int) nutreec[1+c*4]); */

  /* As long as the parent of r is not the root, create new joins in 
   * nutreec that join l with p where p is the parent of l r. 
   * This effectively reverses the orientation of p */
  while(p != 2*ntaxa-2){
    c--; 
    /* The relabeled p will be ntaxa+c-1 rather than l. 
     * New backward branchs start labeling from the root and decrease. */
    nutreec[c*4]=l; nutreec[1+c*4]=ntaxa+c-1;
    nutreec[2+c*4]=bl;
    
    r=p;
    parent_branch(r,ntaxa,utreec,&p,&l,&bl,&br);
    nutreec[3+c*4]=br;
    /* printf("%i %i %i\n", r, p, l); */
    /* printf("%i %i %i\n", c, (int) nutreec[c*4], (int) nutreec[1+c*4]); */
  }
  
  /* finish processing the root join */
  nutreec[1+c*4]=l; nutreec[3+c*4] += bl;
  /* printf("%i %i %i\n\n",  c, (int) nutreec[c*4], (int) nutreec[1+c*4]); */
  nc=c;
  c--;

  /* Start from the root of nutreec and relabel all joins so that they
   * don't conflict with the newly created ones. */
  j=ntaxa-2;
  while(j >= 0){
    kup=(j>nc)?1:2; /* kup set so that newly created joins are not changed */
    for(k = 0; k < kup; k++){
      r=nutreec[k+j*4];
      if (r >= ntaxa){
	/* printf("c j %i %i\n",c,j); */
	for(l = 0; l < 4; l++) 
	  nutreec[l+c*4]=utreec[l+(r-ntaxa)*4]; 
	nutreec[k+j*4]=c+ntaxa;
	c--;
      }
    }
    j--;
  }
}
```
