## Supplementary material for "Modeling site-and-branch-heterogeneity with GFmix": rertf.h

```
#include <stdio.h>
#include <stdlib.h>
#include <math.h>
#include <string.h>
void parent_branch(int r, int ntaxa, double *utreec, int *p, int *l, 
		   double *bl, double *br);
void relab_int(int ntaxa, double *utreec, double *nutreec);
void rert(int r, int ntaxa, double *utreec, double *nutreec);
```
