## Supplementary material for "Modeling site-and-branch-heterogeneity with GFmix": treecns.c

```
#include "treecnsf.h"
int main(int argc, char **argv)
{
  FILE *treefile,*utreecfile;
  char (*names)[11],**ilabels;
  int ntaxa,has_names,i;
  double *utreec;
  
  ntaxa=atoi(argv[--argc]);
  utreecfile=fopen(argv[--argc],"w");
  treefile=fopene(argv[--argc],"r","treefile not available");
  if (ntaxa > 0){
    names=(char (*)[11]) malloc((size_t) ntaxa*sizeof(*names));
    has_names=0;
  }
  else{
    ntaxa=names_infile(stdin,&names,0);
    has_names=1;
  }
  utreec=(double *)malloc(4*ntaxa*sizeof(double));
  ilabels=read_treecs(treefile,ntaxa,names,utreec,has_names);
  pr_utreec(utreecfile,ntaxa,utreec);
  for(i = 0; i < ntaxa; i++) printf("\"%s\"\n",names[i]);
  for(i = 0; i < ntaxa-2; i++) printf("\"%s\"\n",ilabels[i]);
  return(0);
}
```
