## Supplementary material for "Modeling site-and-branch-heterogeneity with GFmix": CommonFunctions.R

```
# Utility function.
strsplit1 <- function(x, split = "") {
  
  # Split input string by split, extract top element from list.
  s <- strsplit(x, split = split)[[1]]
  
  # Return character vector of substrings.
  return(s)
}

# Utility function.
generateAlphabet <- function() {
  alphabet <- c("A", "R", "N", "D", "C", "Q", "E", "G", "H",
                "I", "L", "K", "M", "F", "P", "S", "T", "W", "Y", "V")
  return(alphabet)
}

# Function for tabulating amino acid counts.
countAA <- function(df) {
  
  alphabet <- generateAlphabet()
  
  counts <- lapply(df$Alignment, function(x)
    table(factor(strsplit1(x, ""), levels = alphabet)))
  counts <- do.call(rbind.data.frame, counts)
  rownames(counts) <- df$IDs
  colnames(counts) <- alphabet
  
  return(counts)
}

# Function for reading in multiple sequence alignment.
readAln <- function(infile, format = "phylip-interleaved") {
  
  # Read in file.
  lines <- readLines(infile)
  
  # Parsing multiple sequence alignment in sequential PHYLIP format.
  if (any(format == "phylip", format == "phylip-sequential")) {
    # Extract number of sequences and alignment length.
    dimensions <- as.numeric(strsplit1(lines[c(1)], " ")[-c(1)])
    
    # Split ID and sequence fields in each line by whitespace,
    # bind each split pair as a row in a data.frame.
    df <- data.frame(do.call(rbind, strsplit(lines[-c(1)], "\\s+")))
  }
  
  # Parsing multiple sequence alignment in interleaved PHYLIP format.
  else if (format == "phylip-interleaved") {
    
    # Extract number of sequences and alignment length.
    dimensions <- as.numeric(strsplit1(lines[c(1)], " ")[-c(1)])
    
    # Extract ID and sequence lines while removing dimensions in header, and
    # filter to remove empty lines (lines with length = 0).
    lines <- Filter(length, strsplit(lines[-c(1)], "\\s+"))
    
    # Extract IDs from lines by extracting the first element of the first n
    # lines, for n = number of sequences.
    id <- sapply(lines[seq_len(dimensions[1])], "[[", 1)
    
    # Extract sequence from lines by taking every ith line out of n lines,
    # from i..n+i to n..n+n where i is the ith sequence out of n sequences,
    # removing sequence IDs and pasting all sublines together into a single
    # aligned sequence.
    alignment <- lapply(seq_len(dimensions[1]), function(i) {
      paste0(unlist(lines[seq(i, length(lines), by = dimensions[1])])[-c(1)],
             collapse = "")
    })
    
    # Bind ID and alignment columns together in a data.frame.
    df <- data.frame(cbind(id, alignment))
  }
  
  # Parsing multiple sequence alignment in FASTA format.
  else if (format == "fasta") {
    # Extract sequence header for each sequence by matching ">" in lines,
    # then removing ">" from the match.
    id <- gsub(">", "", lines[grepl(">", lines)])
    
    # Identify which consecutive lines do not start with ">", i.e. must be
    # alignment data (this holds for single- and multi-line FASTA files).
    matches <- which(!grepl(">", lines))
    matches <- split(matches, cumsum(c(1, diff(matches) > 1)))
    
    # For each set of consecutive sequence lines, extract them from lines and
    # paste into a single alignment.
    alignment <- lapply(matches, function(i) paste0(lines[i], collapse = ""))
    
    # Bind ID and alignment columns together in a data.frame.
    df <- data.frame(cbind(id, alignment))
  }
  
  # Add names to dataframe.
  colnames(df) <- c("IDs", "Alignment")
  
  return(df)
}

writeAln <- function(alignment, outfile, format = "phylip") {
  
  ntaxa <- nrow(alignment)
  nsites <- nchar(alignment$Alignment)[1]
  
  if (any(format == "phylip", format == "phylip-sequential")) {
    dimensions <- paste0(" ", ntaxa, " ", nsites)
    pad <- lapply(10 - nchar(alignment$IDs),
                  function(x) paste0(rep(" ", x), collapse = ""))
    alignment <- paste0(alignment$IDs, pad, alignment$Alignment)
    lines <- c(dimensions, alignment)
    
    suffix <- ".phy"
  }
  
  else if (format == "fasta") {
    lines <- paste0(">", alignment$IDs, "\n", alignment$Alignment, collapse = "\n")
    
    suffix <- ".fasta"
  }
  
  outfile <- paste0(outfile, suffix)
  
  cat(lines, sep = "\n", file = outfile)
  return(outfile)
}

# Function for running binomial test.
binomialBinning <- function(dendrogram = NULL, group_1 = NULL, counts, critical = 1.96) {
  
  alphabet <- generateAlphabet()
  
  if (is.null(group_1)) {
    clusters <- cutree(dendrogram, 2)
    clusters <- lapply(seq(unique(clusters)), function(i) names(clusters[clusters == i]))
    
    group_1 <- unlist(clusters[which.max(lengths(clusters))])
    group_2 <- unlist(clusters[!unlist(lapply(clusters, function(x) all(x %in% group_1)))])
  }
  
  else {
    group_2 <- rownames(counts)[!rownames(counts) %in% group_1]
  }
  
  group_1_prnt <- paste("Group 1: ", paste(group_1, collapse = ", "))
  group_2_prnt <- paste("Group 2: ", paste(group_2, collapse = ", "))
  write(group_1_prnt, stdout())
  write(group_2_prnt, stdout())
  
  x1 <- colSums(counts[group_1,])
  x2 <- colSums(counts[group_2,])
  
  n1 <- sum(x1)
  n2 <- sum(x2)
  
  pdiff <- (x1 / n1) - (x2 / n2)
  phat <- (x1 + x2) / (n1 + n2)
  
  se <- sqrt(phat * (1 - phat) * (1/n1 + 1/n2))
  
  zscores <- sort(pdiff / se, decreasing = TRUE)
  
  AAs <- names(zscores)
  
  assignment <- ifelse(zscores > critical, "G-class",
                       ifelse(zscores < -critical, "F-class", "O-class"))
  
  zscores <- cbind("AA" = AAs, "Z-score" = round(zscores, 2), "Assignment" = assignment)
  
  return(zscores)
}
```
