## Supplementary material for "Modeling site-and-branch-heterogeneity with GFmix": Ed_Functions.R

```
LocateRoot <- function(utreec, taxa){
  ##IN: taxa - labels 0,1,.... corresponding to one side of the split
  if(length(taxa)==1) return(taxa)
  ntaxa <- dim(utreec)[1]+1
  
  if(0 %in% taxa){
    spl.of.int <- rep(0,ntaxa)
    spl.of.int[(taxa+1)] <- 1
  }else{
    spl.of.int <- rep(1,ntaxa)
    spl.of.int[(taxa+1)] <- 0
  }
  
  splits <- utreec2splits(utreec)
  spl.label <- -1
  for(i in 1:dim(splits)[1]){
    if(sum(abs(splits[i,]-spl.of.int))==0){
      spl.label <- ntaxa+i-1
      break
    }
  }
  if(spl.label<0) stop("split not found")
  return(spl.label)
}

utreec2splits <- function(utreec){
  # OUT:
  # splits - (ntaxa-3)x ntaxa matrix of 1 and 0; 1 for taxa 1 
  ntaxa <- dim(utreec)[1]+1
  splits <- matrix(0,ncol=ntaxa,nrow=ntaxa-3)
  
  for(split in 1:(ntaxa-3))
    for(k in 1:2){
      b <- utreec[split,k]
      if (b < ntaxa)
        splits[split,(b+1)] <- 1
      else
        splits[split,] <- splits[split,] + splits[(b-ntaxa+1),]
    }
  
  splits <- split.leading1(splits)
  
  return(splits)
}

split.leading1 <- function(spls){
  for(spl in 1:dim(spls)[1])
    if (spls[spl,1] == 0){
      idx <- spls[spl,] == 0
      spls[spl,idx] <- 1
      spls[spl,!idx] <- 0
    }
  return(spls)
}

rert <- function(utree, root, ntaxa, msa_path) {
  
  program <- "bin/rert"
  
  utreecu_filename <- paste0(msa_path, ".utreecu")
  utreec_filename <- paste0(msa_path, ".utreec")
  
  cmd <- paste(program, root, ntaxa, "<", utreecu_filename, ">", utreec_filename)
  names <- system(cmd, intern = TRUE)
  
  utreec <- readLines(utreec_filename)
  utreec <- sapply(utreec, function(line) as.numeric(strsplit1(line, " ")))
  utreec <- matrix(utreec, ncol = 4, byrow = TRUE)
  
  return(utreec)
  
  
}

find.rt <- function(ptree){
  b <- 1
  at.root <- 0
  while(!at.root){
    p <- ptree[b,3]
    pp <- ptree[p,3]
    if (pp == b)
      break
    else
      b <- p
  }
  return(list(b=b, p=p))
}
relabelptree <- function(ptree){
  # relabel branch lengths 
  # should be able to construct utreec after relabelling
  ntaxa <- (dim(ptree)[1] + 2)/2
  a <- find.rt(ptree)
  np <- NULL; lnp <- 0
  if (a$b > ntaxa){
    np <- c(a$b, np)
    lnp <- lnp + 1
  }
  if (a$p > ntaxa){
    np <- c(a$p, np)
    lnp <- lnp + 1
  }
  while(lnp > 0){
    cnp <- np[lnp]
    for(j in 1:2)
      if (ptree[cnp,j] > ntaxa){
        np <- c(ptree[cnp,j], np)
        lnp <- lnp + 1
      }
    lnp <- lnp - 1
  }
  for(i in 1:(ntaxa-2))
    ptree[,1:3] <- ifelse(ptree[,1:3] == np[i], -i, ptree[,1:3])
  ptree[,1:3] <- ifelse(ptree[,1:3] < 0, ntaxa - ptree[,1:3], ptree[,1:3])
  ptree[c((ntaxa+1):(2*ntaxa-2)),] <- ptree[np,]
  return(ptree)
}
nutreec2ptree <- function(nutreec){
  nutreec[,1:2] <- nutreec[,1:2]+1
  ntaxa <- dim(nutreec)[1]+1
  ptree <- rbind(matrix(rep(0,ntaxa*2),ncol=2),nutreec[1:(ntaxa-2),1:2])
  
  cnutreec <- c(nutreec[,1:2]) # columnwise vector
  cbl <- c(nutreec[,3:4])
  p <- rep(c((ntaxa+1):(2*ntaxa-1)),2)
  
  o <- order(cnutreec)
  bl <- cbl[o]
  p <- p[o]
  
  idx.rt <- c(1:(2*ntaxa-2))[p == 2*ntaxa-1] # get root parents correct
  p[idx.rt[1]] <- idx.rt[2]
  p[idx.rt[2]] <- idx.rt[1]
  
  ptree <- cbind(ptree,p,bl)
  return(ptree)
}
rertptree <- function(b, ptree){
  # branch lengths haven't received a lot of testing
  p <- ptree[b,3]
  pp <- ptree[p,3]
  bl <- ptree[,4]
  at.root <- 0
  if (pp == b) at.root <- 1
  if (!at.root){
    ptree[b,4] <- bl[b]/2
    bl[b] <- bl[b]/2 # root branch split in half
  }
  while(!at.root){
    if (b == ptree[p,1])
      ptree[p,1] <- pp
    else
      ptree[p,2] <- pp
    ptree[p,3] <- b
    ptree[p,4] <- bl[b]
    
    b <- p
    p <- pp
    pp <- ptree[p,3]
    if(pp == b){
      at.root <- 1
      ptree[p,4] <- bl[b] + bl[p]
    }
  }
  return(ptree)
}
ptree2nutreec <- function(ptree){
  ptree <- relabelptree(ptree)
  ntaxa <- (dim(ptree)[1] + 2)/2
  nutreec <- matrix(rep(0,(ntaxa-1)*4), ncol = 4)
  for(i in 1:(ntaxa-2)){
    r <- ptree[(ntaxa+i),1]
    l <- ptree[(ntaxa+i),2]
    nutreec[i,] <- cbind(min(r,l),max(r,l),ptree[min(r,l),4],ptree[max(r,l),4])
  }
  a <- find.rt(ptree)
  nutreec[(ntaxa-1),] <- cbind(min(a$b,a$p), max(a$b,a$p),
                               ptree[min(a$b,a$p),4],ptree[max(a$b,a$p),4])
  nutreec[,1:2] <- nutreec[,1:2] - 1
  return(nutreec)
}
rertnutreec <- function(b, utreec){
  # labeling of branches is 1,2,... not 0,1....
  ptree <- nutreec2ptree(utreec)
  ptree <- rertptree(b, ptree)
  utreec <- ptree2nutreec(ptree)
  return(utreec)
}
utreedraw <- function(utreec, names, xlim, smidge = 0.1, smidgel = 0,
                      xaxt = "s", nutreec = TRUE, blabel = F, labels){
  # IN:
  # labels - if missing, 1:(2*ntaxa-1)
  ntaxa <- dim(utreec)[1]+1
  if (missing(names)) names <- as.character(c(0:(ntaxa-1)))
  if (missing(labels)) labels <- c(0:(2*ntaxa-1))
  rht <- utreec[,3]; lht <- utreec[,4]
  r <- utreec[,1] + 1
  l <- utreec[,2] + 1
  
  if (nutreec)
    trees <- vector("list", 2*ntaxa-1)
  else
    trees <- vector("list", ntaxa)
  ntree <- ntaxa
  for(i in 1:ntaxa)
    trees[[i]] <- list(names = names[i], is.tip = TRUE,
                       vseg = NULL, xvseg = 1, tht = 0,
                       hseg = NULL, yhseg = NULL, labels = labels[i])
  for(i in 1:(ntaxa - 1)){
    # merge the two trees into a new tree object:
    #   names - nvseg vector; names of taxa at tips "NODE" or taxa name
    #   is.tip - nvseg vector; indicator about whether vertical line is a tip
    #   vseg - 2xnvseg matrix; lower and upper bounds of vertical segments
    #   xvseg - nvseg vector; giving x values for vertical segments; always starts
    #           at 1
    #   hseg - njoinx2 matrix; lower and upper bounds of horizontal segments
    #   yhseg - nhseq vector; giving y values for horizontal segments
    #   labels - labels of the branches; 1-ntaxa for taxa, -1,...,-(ntaxa-1) joins
    
    tree.r <- trees[[r[i]]]
    tree.l <- trees[[l[i]]]
    r.nvseg <- length(tree.r$xvseg); l.nvseg <- length(tree.l$xvseg)
    nt <- tree.r
    nt$names <- c(tree.r$names, tree.l$names, "NODE")
    nt$labels <- c(tree.r$labels, tree.l$labels, ntaxa+i)
    nt$is.tip <- c(tree.r$is.tip, tree.l$is.tip, FALSE)
    
    xoffset <- max(tree.r$xvseg) + 1
    nt$xvseg <- c(tree.r$xvseg[r.nvseg], tree.l$xvseg[l.nvseg] + xoffset, xoffset)
    nt$names <- c(tree.r$names[r.nvseg], tree.l$names[l.nvseg], "NODE")
    nt$labels <- c(tree.r$labels[r.nvseg], tree.l$labels[l.nvseg], labels[ntaxa+i])
    nt$is.tip <- c(tree.r$is.tip[r.nvseg], tree.l$is.tip[l.nvseg], FALSE)
    if (l.nvseg > 1){
      nt$xvseg <- c(tree.l$xvseg[1:(l.nvseg-1)] + xoffset, nt$xvseg)
      nt$names <- c(tree.l$names[1:(l.nvseg-1)], nt$names)
      nt$labels <- c(tree.l$labels[1:(l.nvseg-1)], nt$labels)
      nt$is.tip <- c(tree.l$is.tip[1:(l.nvseg-1)], nt$is.tip)
    }
    if (r.nvseg > 1){
      nt$xvseg <- c(tree.r$xvseg[1:(r.nvseg-1)], nt$xvseg)
      nt$names <- c(tree.r$names[1:(r.nvseg-1)], nt$names)
      nt$labels <- c(tree.r$labels[1:(r.nvseg-1)], nt$labels)
      nt$is.tip <- c(tree.r$is.tip[1:(r.nvseg-1)], nt$is.tip)
    }
    
    sumr <- tree.r$tht + rht[i]; suml <- tree.l$tht + lht[i]
    nt$tht <- max(sumr, suml)  
    extra.dist <- nt$tht - min(sumr, suml)
    if (sumr < suml){
      if (r.nvseg > 0){
        tree.r$vseg <- tree.r$vseg + extra.dist
        tree.r$yhseg <- tree.r$yhseg + extra.dist
      }
      tree.r$tht <- tree.r$tht + extra.dist
    }
    if (sumr >= suml){
      if (l.nvseg > 0){
        tree.l$vseg <- tree.l$vseg + extra.dist
        tree.l$yhseg <- tree.l$yhseg + extra.dist
      }
      tree.l$tht <- tree.l$tht + extra.dist
    }
    
    nt$vseg <- cbind(tree.r$vseg, tree.l$vseg, c(tree.r$tht, nt$tht),
                     c(tree.l$tht, nt$tht))
    nt$yhseg <- c(tree.r$yhseg, tree.l$yhseg, nt$tht)
    nt$hseg <- rbind(tree.r$hseg, tree.l$hseg + xoffset,
                     c(tree.r$xvseg[r.nvseg], tree.l$xvseg[l.nvseg] + xoffset))
    
    trees[[r[i]]] <- nt
    if (nutreec)
      trees[[i+ntaxa]] <- nt
    else{
      if (l[i] < ntree)
        for(j in l[i]:(ntree-1)) trees[[j]] <- trees[[(j+1)]]
    }
    ntree <- ntree -1
  }
  
  if (missing(xlim)){
    xlim <- c(-0.01, (1 + smidge)*nt$tht)
  }
  ylim <- c(0, 2*ntaxa-1)
  
  plot(xlim, ylim, xlim = xlim, ylim = ylim, type = "n", yaxt = "n",
       xaxt = xaxt, bty = "n", xlab = "", ylab = "")
  nvseg <- length(nt$xvseg) - 1
  for(i in 1:nvseg) segments(nt$tht - nt$vseg[1,i], nt$xvseg[i],
                             nt$tht - nt$vseg[2,i], nt$xvseg[i])
  
  nhseg <- length(nt$yhseg)
  for(i in 1:nhseg) segments(nt$tht - nt$yhseg[i], nt$hseg[i,1],
                             nt$tht - nt$yhseg[i], nt$hseg[i,2])
  
  for(i in 1:nvseg)
    if (nt$is.tip[i]) text(nt$tht - nt$vseg[1,i] + smidgel*nt$tht,
                           nt$xvseg[i], nt$names[i], pos =4, font = 3)
  
  if (blabel){
    for(i in 1:(nvseg-1)){
      if (!nt$is.tip[i])
        text(((nt$tht - nt$vseg[1,i]) +(nt$tht - nt$vseg[2,i]))/2 ,
             nt$xvseg[i]+1.0, as.character(nt$labels[i]))
    }
  }
  return(xlim)
}
utreedraw.flex <- function(utreec, names, xlim, ylim, branchl,
                           xaxt = "s", nutreec = TRUE, 
                           blabel = F, plot=T, labels){
  # IN:
  # utreec - nutreec=T for nutreec format input tree
  # names - will be replaced by 1:ntaxa if not present
  # xlim, ylim - plotting range. can be used to zoom in or allow addl space
  # branchl - branch list. Usually obtained as output from this routine and
  #           then modeified.
  # xaxt - will place an x-axis showing scale. xaxt = "n" => no x-axis
  # blabel = T => add branch labels
  # labels - labels for branches (eg. bootstrap support) when blabel=T. set to
  #         0:2*ntaxa-2 if missing
  
  # OUT: list with components
  # xlim, ylim - plotting range
  # branchl:
  #  x, y: x-y coordinates of names for the branches, including both taxa names
  #        and branch labels (If blabel=F, name is set to "")
  #  x1, x2, yl: (x1,yl), (x2,yl) give the coordinates for the branch that
  #              will be plotted
  #  name: the name of the branch. names are used for terminal branches.
  #        labels is used for internal branches
  #        (If blabel=F, name is set to "")
  #  lwd: line width of branch. set to par()$lwd for each branch initially
  #  lcol: color of branch
  #  col: color of branch label
  #  cex: size of branch label
  #  font: font for branch label
  #  numbers: the number for the branch. an alternative way of referencing
  #           branches.
  
  # COMMENTS:
  # see tree_examples.q for examples
  
  ntaxa <- dim(utreec)[1]+1
  if (missing(names)) names <- as.character(c(0:(ntaxa-1)))
  numbers <- c(1:(2*ntaxa-1))
  if (missing(labels)) labels <- numbers-1
  rht <- utreec[,3]; lht <- utreec[,4]
  r <- utreec[,1] + 1
  l <- utreec[,2] + 1
  
  if (nutreec)
    trees <- vector("list", 2*ntaxa-1)
  else
    trees <- vector("list", ntaxa)
  ntree <- ntaxa
  for(i in 1:ntaxa)
    trees[[i]] <- list(names = names[i], is.tip = TRUE,
                       vseg = NULL, xvseg = 1, tht = 0,
                       hseg = NULL, yhseg = NULL, labels = labels[i],
                       numbers=i)
  for(i in 1:(ntaxa - 1)){
    # merge the two trees into a new tree object:
    #   names - nvseg vector; names of taxa at tips "NODE" or taxa name
    #   is.tip - nvseg vector; indicator about whether vertical line is a tip
    #   vseg - 2xnvseg matrix; lower and upper bounds of vertical segments
    #   xvseg - nvseg vector; giving x values for vertical segments; always starts
    #           at 1
    #   hseg - njoinx2 matrix; lower and upper bounds of horizontal segments
    #   yhseg - nhseq vector; giving y values for horizontal segments
    #   labels - labels of the branches; 1-ntaxa for taxa, -1,...,-(ntaxa-1) joins
    
    tree.r <- trees[[r[i]]]
    tree.l <- trees[[l[i]]]
    r.nvseg <- length(tree.r$xvseg); l.nvseg <- length(tree.l$xvseg)
    nt <- tree.r
    
    xoffset <- max(tree.r$xvseg) + 1
    nt$xvseg <- c(tree.r$xvseg[r.nvseg], tree.l$xvseg[l.nvseg] + xoffset, xoffset)
    nt$names <- c(tree.r$names[r.nvseg], tree.l$names[l.nvseg], "NODE")
    nt$labels <- c(tree.r$labels[r.nvseg], tree.l$labels[l.nvseg], labels[ntaxa+i])
    nt$numbers <- c(tree.r$numbers[r.nvseg], tree.l$numbers[l.nvseg], numbers[ntaxa+i])
    nt$is.tip <- c(tree.r$is.tip[r.nvseg], tree.l$is.tip[l.nvseg], FALSE)
    if (l.nvseg > 1){
      nt$xvseg <- c(tree.l$xvseg[1:(l.nvseg-1)] + xoffset, nt$xvseg)
      nt$names <- c(tree.l$names[1:(l.nvseg-1)], nt$names)
      nt$labels <- c(tree.l$labels[1:(l.nvseg-1)], nt$labels)
      nt$numbers <- c(tree.l$numbers[1:(l.nvseg-1)], nt$numbers)
      nt$is.tip <- c(tree.l$is.tip[1:(l.nvseg-1)], nt$is.tip)
    }
    if (r.nvseg > 1){
      nt$xvseg <- c(tree.r$xvseg[1:(r.nvseg-1)], nt$xvseg)
      nt$names <- c(tree.r$names[1:(r.nvseg-1)], nt$names)
      nt$labels <- c(tree.r$labels[1:(r.nvseg-1)], nt$labels)
      nt$numbers <- c(tree.r$numbers[1:(r.nvseg-1)], nt$numbers)
      nt$is.tip <- c(tree.r$is.tip[1:(r.nvseg-1)], nt$is.tip)
    }
    
    sumr <- tree.r$tht + rht[i]; suml <- tree.l$tht + lht[i]
    nt$tht <- max(sumr, suml)  
    extra.dist <- nt$tht - min(sumr, suml)
    if (sumr < suml){
      if (r.nvseg > 0){
        tree.r$vseg <- tree.r$vseg + extra.dist
        tree.r$yhseg <- tree.r$yhseg + extra.dist
      }
      tree.r$tht <- tree.r$tht + extra.dist
    }
    if (sumr >= suml){
      if (l.nvseg > 0){
        tree.l$vseg <- tree.l$vseg + extra.dist
        tree.l$yhseg <- tree.l$yhseg + extra.dist
      }
      tree.l$tht <- tree.l$tht + extra.dist
    }
    
    nt$vseg <- cbind(tree.r$vseg, tree.l$vseg, c(tree.r$tht, nt$tht),
                     c(tree.l$tht, nt$tht))
    nt$yhseg <- c(tree.r$yhseg, tree.l$yhseg, nt$tht)
    nt$hseg <- rbind(tree.r$hseg, tree.l$hseg + xoffset,
                     c(tree.r$xvseg[r.nvseg], tree.l$xvseg[l.nvseg] + xoffset))
    
    trees[[r[i]]] <- nt
    if (nutreec)
      trees[[i+ntaxa]] <- nt
    else{
      if (l[i] < ntree)
        for(j in l[i]:(ntree-1)) trees[[j]] <- trees[[(j+1)]]
    }
    ntree <- ntree -1
  }
  
  smidge <- 0.1 
  if (missing(xlim)){
    xlim <- c(-0.01, (1 + smidge)*nt$tht)
  }
  if (missing(ylim)){
    ylim <- c(0, 2*ntaxa-1)
  }
  
  if (plot)
    plot(xlim, ylim, xlim = xlim, ylim = ylim, type = "n", yaxt = "n",
         xaxt = xaxt, bty = "n", xlab = "", ylab = "")
  nvseg <- length(nt$xvseg) - 1
  
  nhseg <- length(nt$yhseg)
  if (plot)
    for(i in 1:nhseg) segments(nt$tht - nt$yhseg[i], nt$hseg[i,1],
                               nt$tht - nt$yhseg[i], nt$hseg[i,2])
  
  # Create a list with the x-y coordinates of names for the branches
  # and (x1,yl), (x2,yl) coordinates for the segments as well as
  # lcol, lwd (line colour and width), col and cex (colour and cex label) 
  if (missing(branchl)){
    branchl <- list(x=NULL, y=NULL, x1=NULL, x2=NULL, yl=NULL, name=NULL,
                    lcol=NULL, lwd=NULL, col=NULL, cex=NULL, font=NULL,
                    numbers=nt$numbers)
    for(i in 1:nvseg){
      branchl$x1 <- c(branchl$x1, nt$tht - nt$vseg[1,i])
      branchl$x2 <- c(branchl$x2, nt$tht - nt$vseg[2,i])
      branchl$yl <- c(branchl$yl, nt$xvseg[i])
      if (nt$is.tip[i]){
        branchl$x <- c(branchl$x,nt$tht - nt$vseg[1,i])
        branchl$y <- c(branchl$y, nt$xvseg[i])
        branchl$name <- c(branchl$name, nt$names[i])
        branchl$font <- c(branchl$font, 3)
      }
      if (!nt$is.tip[i]){
        branchl$x <- c(branchl$x,
                       0.25*(nt$tht - nt$vseg[1,i]) +
                         0.75*(nt$tht - nt$vseg[2,i]))
        branchl$y <- c(branchl$y, nt$xvseg[i]+0.8)
        if (blabel)
          branchl$name <- c(branchl$name, as.character(nt$labels[i]))
        else
          branchl$name <- c(branchl$name, "")
        branchl$font <- c(branchl$font, par()$font)
      }
    }
    branchl$cex <- rep(par()$cex, length(branchl$x))
    branchl$lcol <- rep(par()$col, length(branchl$x))
    branchl$col <- rep(par()$col, length(branchl$x))
    branchl$lwd <- rep(par()$lwd, length(branchl$x))
  }
  
  if(plot){
    text(branchl$x, branchl$y, branchl$name, pos=4, font=branchl$font,
         col=branchl$col, cex=branchl$cex)
    segments(branchl$x1, branchl$yl, branchl$x2, branchl$yl, lwd=branchl$lwd,
             col=branchl$lcol)
  }
  
  return(list(xlim=xlim, ylim=ylim, branchl=branchl))
}
#= convert utreec to matrix of splits
utreec2splits <- function(utreec){
  # OUT:
  # splits - (ntaxa-3)x ntaxa matrix of 1 and 0; 1 for taxa 1 
  ntaxa <- dim(utreec)[1]+1
  splits <- matrix(0,ncol=ntaxa,nrow=ntaxa-3)
  
  for(split in 1:(ntaxa-3))
    for(k in 1:2){
      b <- utreec[split,k]
      if (b < ntaxa)
        splits[split,(b+1)] <- 1
      else
        splits[split,] <- splits[split,] + splits[(b-ntaxa+1),]
    }
  
  splits <- split.leading1(splits)
  
  return(splits)
}

utreec2el <- function(utreec){
  edge.label <- c(utreec[,1:2])+1
  el <- c(utreec[,3:4])
  el[edge.label] <- el
  return(el)
}

utreec2splitsel <- function(utreec){
  splits <- utreec2splits(utreec)
  el <- utreec2el(utreec)
  return(list(splits=splits, el=el))
}
treecns_Ed <- function(tree,treefile,infile="",ntaxa,clean=TRUE,prcmd=FALSE){
  
  ##IN: tree (string) or treefile and (for names) infile or ntaxa
  ##RETURN: list with utreec and names
  if((missing(tree) && missing(treefile)) ||
     (!missing(tree) && !missing(treefile)))
    stop("Exactly one of tree or treefile must be specified")
  if((infile=="" && missing(ntaxa)) || (infile!="" && !missing(ntaxa)))
    stop("Exactly one of ntaxa or infile must be specified")
  
  if(!missing(tree)) {cat(tree,"\n",file="tmp.tree"); treefile <- "tmp.tree"}
  ## Charley Might want to change this
  cmdline <- "bin/treecns"
  cmdline <- paste(cmdline,treefile,"tmp.utreec")
  ## cat(cmdline,"\n")
  if((infile=="" && missing(ntaxa)) || (infile!="" && !missing(ntaxa)))
    stop("Exactly one of ntaxa or infile must be specified")
  if(infile!="")
    cmdline <- paste(cmdline,"-1 < ",infile," > tmp.out")
  if(!missing(ntaxa))
    cmdline <- paste(cmdline,ntaxa," > tmp.out")
  if(prcmd) cat(cmdline,"\n")
  system(cmdline)
  utreec <- matrix(scan("tmp.utreec",quiet=T),ncol=4,byrow=T)
  names <- scan("tmp.out",quiet=T,what=character())
  if(clean){
    file.remove("tmp.utreec","tmp.out")
    if(!missing(tree)) file.remove("tmp.tree")
  }
  return(list(utreec=utreec,names=names))
}

## Routine to produce inbv file
## 

##= Replace edge-lengths in inbv.file with edge-lengths from utreec, gradient
##  and Hessian with gradient and Hessian from paramfile. Does necessary
##  reordering of entries
## IN: inbv.file - Mock inbv.file for mcmctree. The Newick tree is the only
##       thing read in. Edge-lengths are assumed to give the order of the edge
##       in the line after the Newick tree. So there is an implicit assumption
##       that Andrew's routine has been run with numbers 0,1,.... gving the
##       ordering of edge-lengths. For instance (...(rabbit:5, ...)
##       implies the 6th entry in the list of edge lengths is the edge length
##       for the terminal `rabbit' edge-length
##     seqfile - PHYLIP format sequence file. The names in seqfile should match
##       the names in the Newick string in inbv.file
##     mundi.paramfile - paramfile output by mundi with -x option
## OUT: new.inbv.file 
FormatInBv <- function(intree, ref_tree, seqfile, mundi.paramfile, new.inbv.file){
  
  inbv.utreec <- treecns_Ed(tree=intree,infile=seqfile)$utreec
  ntaxa <- dim(inbv.utreec)[1]+1
  inbv.splits.el <- utreec2splitsel(inbv.utreec)
  
  x <- scan(mundi.paramfile,quiet=TRUE)[-c(1:2)]
  utreec <- matrix(x[1:(4*(ntaxa-1))],ncol=4,byrow=TRUE)
  utreec.splits.el <- utreec2splitsel(utreec)
  
  ## first ntaxa edges are terminal. inbv edge-lengths give order in inbv file
  req.perm <- inbv.splits.el$el[1:ntaxa]
  
  ## order in inbv file of internal edges.
  for(j in 1:dim(utreec.splits.el$splits)[1])
    for(i in 1:dim(inbv.splits.el$splits)[1]) # determine matching split
      if(sum(abs(inbv.splits.el$splits[i,] -
                 utreec.splits.el$splits[j,]))<1.0e-6){
        req.perm <- c(req.perm, inbv.splits.el$el[(i+ntaxa)])
        break
      }
  req.perm <- req.perm + 1
  
  el <- rep(-1,length(req.perm))
  el[req.perm] <- utreec.splits.el$el[1:length(req.perm)]
  
  hess <- x[-length(x)] # last entry is measure of goodness of hess approx
  hess <- hess[c((length(hess)-(2*ntaxa-3)*(2*ntaxa-3)-(2*ntaxa-3)+1):length(hess))]
  grad <- hess[1:(2*ntaxa-3)]
  hess <- matrix(hess[-c(1:(2*ntaxa-3))],ncol=2*ntaxa-3)
  
  grad[req.perm] <- grad
  hess[req.perm,req.perm] <- hess
  
  ## output to new.inbv
  cat(ntaxa,"\n\n",file=new.inbv.file)
  cat(ref_tree,"\n\n",file=new.inbv.file,append=TRUE)
  ##  cat(sprintf("%.16e ",el),"\n\n",file=new.inbv.file,append=TRUE)
  cat(sprintf("%.6g ",el),"\n\n",file=new.inbv.file,append=TRUE)
  cat(sprintf("%.16e ",grad),"\n\n",file=new.inbv.file,append=TRUE)
  cat("Hessian\n\n",file=new.inbv.file,append=TRUE)
  write(sprintf("%10.2f",hess),ncol=2*ntaxa-3,file=new.inbv.file,append=TRUE)
}
```
