## Supplementary material for "Modeling site-and-branch-heterogeneity with GFmix": GFmixFunctions.R

```
# Taken from Ed.
LocateRoot <- function(utreec, taxa){
  ##IN: taxa - labels 0,1,.... corresponding to one side of the split
  if(length(taxa)==1) return(taxa)
  ntaxa <- dim(utreec)[1]+1
  
  if(0 %in% taxa){
    spl.of.int <- rep(0,ntaxa)
    spl.of.int[(taxa+1)] <- 1
  }else{
    spl.of.int <- rep(1,ntaxa)
    spl.of.int[(taxa+1)] <- 0
  }
  
  splits <- utreec2splits(utreec)
  spl.label <- -1
  for(i in 1:dim(splits)[1]){
    if(sum(abs(splits[i,]-spl.of.int))==0){
      spl.label <- ntaxa+i-1
      break
    }
  }
  if(spl.label<0) stop("split not found")
  return(spl.label)
}

utreec2splits <- function(utreec){
  # OUT:
  # splits - (ntaxa-3)x ntaxa matrix of 1 and 0; 1 for taxa 1 
  ntaxa <- dim(utreec)[1]+1
  splits <- matrix(0,ncol=ntaxa,nrow=ntaxa-3)
  
  for(split in 1:(ntaxa-3))
    for(k in 1:2){
      b <- utreec[split,k]
      if (b < ntaxa)
        splits[split,(b+1)] <- 1
      else
        splits[split,] <- splits[split,] + splits[(b-ntaxa+1),]
    }
  
  splits <- split.leading1(splits)
  
  return(splits)
}

split.leading1 <- function(spls){
  for(spl in 1:dim(spls)[1])
    if (spls[spl,1] == 0){
      idx <- spls[spl,] == 0
      spls[spl,idx] <- 1
      spls[spl,!idx] <- 0
    }
  return(spls)
}

treeCNS <- function(msa_path, tree) {
  
  program <- "bin/treecns"
  
  utreecu_filename <- paste0(msa_path, ".utreecu")
  
  cmd <- paste(program, tree, utreecu_filename, "-1 <", msa_path)
  names <- system(cmd, intern = TRUE)
  
  utreecu <- readLines(utreecu_filename)
  utreecu <- sapply(utreecu, function(line) as.numeric(strsplit1(line, " ")))
  utreecu <- matrix(utreecu, ncol = 4, byrow = TRUE)
  
  return(utreecu)
}

rert <- function(utree, root, ntaxa, msa_path) {
  
  program <- "bin/rert"
  
  utreecu_filename <- paste0(msa_path, ".utreecu")
  utreec_filename <- paste0(msa_path, ".utreec")
  
  cmd <- paste(program, root, ntaxa, "<", utreecu_filename, ">", utreec_filename)
  names <- system(cmd, intern = TRUE)
  
  utreec <- readLines(utreec_filename)
  utreec <- sapply(utreec, function(line) as.numeric(strsplit1(line, " ")))
  utreec <- matrix(utreec, ncol = 4, byrow = TRUE)
  
  return(utreec)
  
  
}

parseIQTREE <- function(iqtree) {
  
  lines <- readLines(iqtree)
  
  # Extract tree likelihood from line containing likelihood information.
  lnl <- lines[grepl("Log-likelihood of the tree:", lines)]
  lnl <- strsplit1(lnl, ": ")[2]
  lnl <- as.numeric(strsplit1(lnl, " ")[1])
  
  # Extract alpha value from line containing alpha information.
  alpha <- lines[grepl("Gamma shape alpha:", lines)]
  alpha <- as.numeric(strsplit1(alpha, " ")[4])
  
  weights <- lines[grepl("^[ ]{1,3}\\d{1,3}[ ]{2}\\w", lines)]
  weights <- lapply(strsplit(weights, "\\s+"), function(y) y[5])
  weights <- unlist(weights)
  weights <- as.numeric(weights)
  
  # If +F, extract frequencies for +F class from IQTREE and reorder
  # weights so that the +F class weights appear at the end of the
  # weights vector.
  est_freqs <- lines[grepl("  pi", lines)]
  if (length(est_freqs) == 20) {
    plusF <- lapply(strsplit(est_freqs, "\\s+"), function(y) y[4])
    plusF <- unlist(plusF)
    plusF <- as.numeric(plusF)
    weights <- weights[c(2:length(weights), 1)]
  }
  else {
    plusF <- as.numeric(NULL)
  }
  
  parsed <- data.frame("Likelihood" = lnl,
                "Alpha" = alpha,
                "plusF" = I(list(plusF)),
                "Weights" = I(list(weights)))
  
  return(parsed)
}

convertIDs <- function(msa_path, treefile_path, format) {
  
  msa <- readAln(msa_path, format = format)
  treefile <- readLines(treefile_path)
  
  
  new_ids <- seq(0, nrow(msa) - 1)
  
  for (i in seq(new_ids)) {
    treefile <- gsub(msa$IDs[i], new_ids[i], treefile)
  }
  
  msa$IDs <- new_ids
  
  con_msa_path <- paste0(msa_path, "_INT")
  con_treefile_path <- paste0(treefile_path, "_INT")
  
  writeAln(msa, con_msa_path, format)
  write(treefile, con_treefile_path)
  
  prnt <- paste("Converted IDs in input alignment and tree to 0-index integers.",
                "File paths:", con_msa_path, "and", con_treefile_path, ".")
  write(prnt, stdout())
}

GFmix <- function(msa_path, mixture, tree, iqtree, rootfile,
                  g_class, f_class, plusF = FALSE, optimizer = FALSE,
                  afo = NULL, params = NULL) {
  
  if (optimizer) {
    program <- "bin/gfmix_for_opt"
  } else if (!is.null(afo)) {
    program <- "bin/gfmix_for_afo"
  } else {
    program <- "bin/gfmix"
  }
  
  g_class <- paste0(g_class, collapse = "")
  f_class <- paste0(f_class, collapse = "")
  
  cmd <- paste(program, "-s", msa_path, "-t", tree, "-i", iqtree,
               "-f", mixture, "-r", rootfile, "-gclass", g_class,
               "-fclass", f_class)

  if(!is.null(afo)) {
    afo <- paste0(afo, collapse = ",")
    cmd <- paste(cmd, "-afo", afo)
  }
  
  if (!plusF) {
    cmd <- paste(cmd, "-d")
  }
  if (!is.null(params)) {
    cmd <- paste(cmd, "-x", params)
  }

  lnl <- system(cmd, intern = TRUE)
  lnl <- as.numeric(lnl)
  
  return(lnl)
  
}

AnFreqOpt <- function(msa_path, mixture, utreec_path, alpha, w_filename,
                      b_filename, g_class, f_class, ss = TRUE) {
  
  program <- "bin/an_freq_opt_fast"
  
  mval <- 1.0e-8
  lflag <- 1e-02
  mflag <- 1e+02
  tol <- 1e+7
  
  alphabet <- generateAlphabet()
  o_class <- alphabet[!alphabet %in% c(g_class, f_class)]
  d_vector <- rep(2, 20)
  d_vector[alphabet %in% g_class] <- 0
  d_vector[alphabet %in% f_class] <- 1
  d_filename <- paste0(msa_path, "_", paste0(g_class, collapse = ""), "_",
                       paste0(o_class, collapse = ""),
                       "_", paste0(f_class, collapse = ""), ".dfile")
  write(d_vector, d_filename, sep = "\n")
  
  cmd <- paste(program, "-i", msa_path, "-u", utreec_path, "-w", w_filename,
               "-f", mixture, "-a", alpha, "-l", lflag,
               "-m", mflag, "-b", b_filename, "-d", d_filename, "-e", tol)

  afo <- system(cmd, intern = TRUE)
  afo <- as.numeric(afo)
  
  file.remove(d_filename)
  
  if (ss) {
    afo <- afo[1]
  } else {
    afo <- afo[c(-1)]
  }
  return(afo)
}


NewGFmix <- function(msa_path, mixture, utreec_path, alpha, w_filename,
                     g_class, f_class, mode = 4, params = NULL, multis = NULL) {
  
  program <- "bin/gfmix_lnl_fast"
  
  iprint <- 1
  
  nclass <- length(readLines(mixture))
  
  alphabet <- generateAlphabet()
  d_vector <- rep(2, 20)
  d_vector[alphabet %in% g_class] <- 0
  d_vector[alphabet %in% f_class] <- 1
  d_filename <- paste0(msa_path, "_", paste0(g_class, collapse = ""),
                       "_", paste0(f_class, collapse = ""), ".dfile")
  write(d_vector, d_filename, sep = "\n")
  
  cmd <- paste(program, "-s", msa_path, "-c", nclass, "-f", mixture,
               "-u", utreec_path, "-w ", w_filename, "-a", alpha,
               "-g", d_filename, "-o", mode, "-p", iprint)
  
  if (!is.null(params)) {
    cmd <- paste(cmd, "-x", params)
  }
  
  if (!is.null(multis)) {
    cmd <- paste(cmd, "-m", multis)
  }

  cmd <- paste(cmd, "2>&1 | tee /dev/stderr")
  
  lnl <- system(cmd, intern = TRUE)
  
  info <- paste0(tail(lnl, 15), collapse = ";")
  lnl <- as.numeric(tail(lnl, 1))
  
  return(list("lnl" = lnl, "info" = info))
  
}


AnFreq <- function(utreec, wtfile, fmatfile, alpha, bfile) {
  
  program <- "bin/an_freq"
  
  cmd <- paste(program, "-u", utreec, "-w", wtfile, "-f",
              fmatfile, "-a", alpha, "-b", bfile)
  
  af <- as.numeric(system(cmd, intern = TRUE))
  af <- matrix(af, ncol = 20, byrow = TRUE)
  af <- af / 100
  
  return(af)
}
```
