## Supplementary material for "Modeling site-and-branch-heterogeneity with GFmix": IdentificationFunctions.R

```
generateSwaps <- function(g_class, o_class, f_class) {
  
  gswap <- seq(g_class)
  fswap <- seq(f_class)
  oswap <- seq(o_class)
  
  gto <- lapply(gswap, function(i) {
    no <- c(g_class[i], o_class)
    ng <- g_class[-c(i)]
    nf <- f_class
    
    if (any(is.null(ng), is.null(nf))) {
      swap <- NULL
    }
    else {
      swap <- list("G" = ng, "O" = no, "F" = nf)
    }
    
    return(swap)
    
  })
  gto <- do.call(rbind, gto)
  
  otg <- lapply(oswap, function(i) {
    ng <- c(g_class, o_class[i])
    no <- o_class[-c(i)]
    nf <- f_class
    
    if (any(is.null(ng), is.null(nf))) {
      swap <- NULL
    }
    else {
      swap <- list("G" = ng, "O" = no, "F" = nf)
    }
    
    return(swap)
    
  })
  otg <- do.call(rbind, otg)
  
  otf <- lapply(oswap, function(i) {
    ng <- g_class
    nf <- c(o_class[i], f_class)
    no <- o_class[-c(i)]
    
    if (any(is.null(ng), is.null(nf))) {
      swap <- NULL
    }
    else {
      swap <- list("G" = ng, "O" = no, "F" = nf)
    }
    
    return(swap)
    
  })
  otf <- do.call(rbind, otf)
  
  fto <- lapply(fswap, function(i) {
    ng <- g_class
    no <- c(o_class, f_class[i])
    nf <- f_class[-c(i)]
    
    if (any(is.null(ng), is.null(nf))) {
      swap <- NULL
    }
    else {
      swap <- list("G" = ng, "O" = no, "F" = nf)
    }
    
    return(swap)
  })
  fto <- do.call(rbind, fto)
  
  starting <- list("G" = g_class, "O" = o_class, "F" = f_class)
  starting <- rbind(starting)
  
  swaps <- rbind.data.frame("Starting" = starting, gto, otg, otf, fto)
  
  return(swaps)
  
}

optimizeByChi <- function(msa, binomial_test, garpfymink = FALSE, limit = NULL,
                          track_swaps = FALSE, threads = 1) {
  
  counts <- countAA(msa)
  
  if (garpfymink) {
    g_class <- c("G", "A", "R", "P")
    f_class <- c("F", "Y", "M", "I", "N", "K")
    o_class <- generateAlphabet()[!generateAlphabet() %in% c(g_class, f_class)]
  }
  else{
    g_class <- binomial_test$AA[binomial_test$Assignment == "G-class"]
    o_class <- binomial_test$AA[binomial_test$Assignment == "O-class"]
    f_class <- binomial_test$AA[binomial_test$Assignment == "F-class"]
  }
  
  
  if (is.null(limit)) {
    limit <- Inf
  }
  
  optimize <- TRUE
  itr <- 1
  opt_time <- Sys.time()
  
  itr_table <- data.frame()
  
  while(optimize) {
    
    itr_time <- Sys.time()
    
    swaps <- generateSwaps(g_class, o_class, f_class)
    
    nswaps <- seq(nrow(swaps))
    
    opt_prnt <- paste0("Iteration ", itr, ": Assessing ",
                       length(nswaps), " swaps from starting point ",
                       paste0(g_class, collapse = ""), "/",
                       paste0(o_class, collapse = ""), "/",
                       paste0(f_class, collapse = ""), ".")
    
    write(opt_prnt, stdout())
    
    swaps$Criterion <- mclapply(nswaps, function(i) {
      
      swap <- swaps[i,]
      
      g_counts <- counts[unlist(swap$G), drop = FALSE]
      f_counts <- counts[unlist(swap$F), drop = FALSE]
      
      g_counts <- rowSums(g_counts)
      f_counts <- rowSums(f_counts)
      
      swap_time <- Sys.time()
      chi2 <- chisq.test(cbind(g_counts, f_counts))
      chi2 <- as.numeric(chi2$statistic)
      swap_time <- as.numeric(difftime(Sys.time(), swap_time ,units = "secs"))
      
      return(list("Chi2" = chi2, "Swap" = swap_time))
    }, mc.cores = threads)
    
    swap_times <- unlist(lapply(swaps$Criterion, function(x) x$Swap))
    swaps$Criterion <- unlist(lapply(swaps$Criterion, function(x) x$Chi2))
    
    best <- swaps[which.max(swaps$Criterion),]
    starting <- swaps["Starting",]
    
    if (unlist(best$Criterion) > unlist(starting$Criterion)) {
      
      g_class <- unlist(best$G)
      o_class <- unlist(best$O)
      f_class <- unlist(best$F)
      
      itr_time <- as.numeric(difftime(Sys.time(), itr_time ,units = "secs"))
      curr_time <- as.numeric(difftime(Sys.time(), opt_time ,units = "secs"))
      
      if (itr == limit) {
        
        opt_time <- curr_time
        
        opt_prnt <- paste0("Iteration ", itr, ": G/O/F at iteration limit: ",
                           paste0(g_class, collapse = ""), "/",
                           paste0(o_class, collapse = ""), "/",
                           paste0(f_class, collapse = ""), ". ",
                           "Iteration time: ", round(itr_time, 3), ", ",
                           "Final optimization time: ", round(opt_time, 3), ".")
        
        itr_row <- cbind.data.frame(best, "Optimizer" = "Chi2",
                                    "Iterations" = itr, "Iteration time" = itr_time,
                                    "Optimization time" = opt_time)
        
        if (track_swaps) {
          itr_row <- cbind.data.frame(itr_row, "Swap times" = I(list(swap_times)))
        }
        
        itr_table <- rbind(itr_table, itr_row)
        
        write(opt_prnt, stdout())
        optimize <- FALSE
        
      } else {
        
        opt_prnt <- paste0("Iteration ", itr, ": Found new starting point ",
                           paste0(g_class, collapse = ""), "/",
                           paste0(o_class, collapse = ""), "/",
                           paste0(f_class, collapse = ""), ". ",
                           "Iteration time (s): ", round(itr_time, 3), ", ",
                           "Current optimization time (s): ", round(curr_time, 3), ".")
        
        itr_row <- cbind.data.frame(best, "Optimizer" = "Chi2",
                                    "Iterations" = itr, "Iteration time" = itr_time,
                                    "Optimization time" = curr_time)
        
        if (track_swaps) {
          itr_row <- cbind.data.frame(itr_row, "Swap times" = I(list(swap_times)))
        }
        
        itr_table <- rbind(itr_table, itr_row)
        
        itr <- itr + 1
        write(opt_prnt, stdout())
      }
      
      
    } else {
      g_class <- unlist(starting$G)
      o_class <- unlist(starting$O)
      f_class <- unlist(starting$F)
      
      itr_time <- as.numeric(difftime(Sys.time(), itr_time ,units = "secs"))
      opt_time <- as.numeric(difftime(Sys.time(), opt_time ,units = "secs"))
      
      opt_prnt <- paste0("Iteration ", itr, ": G/O/F at optimization limit: ",
                         paste0(g_class, collapse = ""), "/",
                         paste0(o_class, collapse = ""), "/",
                         paste0(f_class, collapse = ""), ". ",
                         "Iteration time: ", round(itr_time, 3), ", ",
                         "Final optimization time: ", round(opt_time, 3), ".")
      
      itr_row <- cbind.data.frame(starting, "Optimizer" = "Chi2",
                                  "Iterations" = itr, "Iteration time" = itr_time,
                                  "Optimization time" = opt_time)
      
      if (track_swaps) {
        itr_row <- cbind.data.frame(itr_row, "Swap times" = I(list(swap_times)))
      }
      
      itr_table <- rbind(itr_table, itr_row)
      
      write(opt_prnt, stdout())
      optimize <- FALSE
    }
    
  }
  
  rownames(itr_table) <- seq(nrow(itr_table))
  return(itr_table)
}

optimizeBySSF <- function(msa, msa_path, binomial_test, garpfymink = FALSE, limit = NULL,
                         tree = NULL, iqtree = NULL, rootfile = NULL,
                         mixture = NULL,
                         track_swaps = FALSE, threads = 1) {
  
  ntaxa <- nrow(msa)
  
  iqtree <- parseIQTREE(iqtree)
  alpha <- iqtree$Alpha
  w_vector <- unlist(iqtree$Weights)
  w_filename <- paste0(msa_path, ".wfile")
  write(w_vector, w_filename, sep = "\n")
  
  b_vector <- rep(1, (2 * ntaxa) - 1)
  b_filename <- paste0(msa_path, ".bfile")
  write(b_vector, b_filename, sep = "\n")
  
  utree <- treeCNS(msa_path, tree)
  
  root <- as.numeric(strsplit1(readLines(rootfile)), " ")
  root <- LocateRoot(utree, root)
  
  utreec <- rert(utree, root, ntaxa, msa_path)
  utreec_path <- paste0(msa_path, ".utreec")
  
  if (garpfymink) {
    g_class <- c("G", "A", "R", "P")
    f_class <- c("F", "Y", "M", "I", "N", "K")
    o_class <- generateAlphabet()[!generateAlphabet() %in% c(g_class, f_class)]
  }
  else{
    g_class <- binomial_test$AA[binomial_test$Assignment == "G-class"]
    o_class <- binomial_test$AA[binomial_test$Assignment == "O-class"]
    f_class <- binomial_test$AA[binomial_test$Assignment == "F-class"]
  }
  
  
  if (is.null(limit)) {
    limit <- Inf
  }
  
  optimize <- TRUE
  itr <- 1
  opt_time <- Sys.time()
  
  itr_table <- data.frame()
  
  while(optimize) {
    
    itr_time <- Sys.time()
    
    swaps <- generateSwaps(g_class, o_class, f_class)
    
    nswaps <- seq(nrow(swaps))
    
    opt_prnt <- paste0("Iteration ", itr, ": Assessing ",
                       length(nswaps), " swaps from starting point ",
                       paste0(g_class, collapse = ""), "/",
                       paste0(o_class, collapse = ""), "/",
                       paste0(f_class, collapse = ""), ".")
    
    write(opt_prnt, stdout())
    
    swaps$Criterion <- mclapply(nswaps, function(i) {
      
      swap <- swaps[i,]
      g_class <- unlist(swap$G)
      f_class <- unlist(swap$F)
      
      swap_time <- Sys.time()
      
      afo <- AnFreqOpt(msa_path, mixture, utreec_path, alpha,
                       w_filename, b_filename, g_class, f_class)
      
      swap_time <- as.numeric(difftime(Sys.time(), swap_time ,units = "secs"))
      
      return(list("SS" = afo, "Swap" = swap_time))
    }, mc.cores = threads)
    
    swap_times <- unlist(lapply(swaps$Criterion, function(x) x$Swap))
    swaps$Criterion <- unlist(lapply(swaps$Criterion, function(x) x$SS))
    
    best <- swaps[which.min(swaps$Criterion),]
    starting <- swaps["Starting",]
    
    if (unlist(best$Criterion) < unlist(starting$Criterion)) {
      
      g_class <- unlist(best$G)
      o_class <- unlist(best$O)
      f_class <- unlist(best$F)
      
      itr_time <- as.numeric(difftime(Sys.time(), itr_time ,units = "secs"))
      curr_time <- as.numeric(difftime(Sys.time(), opt_time ,units = "secs"))
      
      if (itr == limit) {
        
        opt_time <- curr_time
        
        opt_prnt <- paste0("Iteration ", itr, ": G/O/F at iteration limit: ",
                           paste0(g_class, collapse = ""), "/",
                           paste0(o_class, collapse = ""), "/",
                           paste0(f_class, collapse = ""), ". ",
                           "Iteration time: ", round(itr_time, 3), ", ",
                           "Final optimization time: ", round(opt_time, 3), ".")
        
        itr_row <- cbind.data.frame(best, "Optimizer" = "SSF",
                                    "Iterations" = itr, "Iteration time" = itr_time,
                                    "Optimization time" = curr_time)
        
        if (track_swaps) {
          itr_row <- cbind.data.frame(itr_row, "Swap times" = I(list(swap_times)))
        }
        
        itr_table <- rbind(itr_table, itr_row)
        
        write(opt_prnt, stdout())
        optimize <- FALSE
        
        
      } else {
        
        opt_prnt <- paste0("Iteration ", itr, ": Found new starting point ",
                           paste0(g_class, collapse = ""), "/",
                           paste0(o_class, collapse = ""), "/",
                           paste0(f_class, collapse = ""), ". ",
                           "Iteration time (s): ", round(itr_time, 3), ", ",
                           "Current optimization time (s): ", round(curr_time, 3), ".")
        
        itr_row <- cbind.data.frame(best, "Optimizer" = "SSF",
                                    "Iterations" = itr, "Iteration time" = itr_time,
                                    "Optimization time" = curr_time)
        
        if (track_swaps) {
          itr_row <- cbind.data.frame(itr_row, "Swap times" = I(list(swap_times)))
        }
        
        itr_table <- rbind(itr_table, itr_row)
        
        itr <- itr + 1
        write(opt_prnt, stdout())
      }
      
      
    } else {
      g_class <- unlist(starting$G)
      o_class <- unlist(starting$O)
      f_class <- unlist(starting$F)
      
      itr_time <- as.numeric(difftime(Sys.time(), itr_time ,units = "secs"))
      opt_time <- as.numeric(difftime(Sys.time(), opt_time ,units = "secs"))
      
      opt_prnt <- paste0("Iteration ", itr, ": G/O/F at optimization limit: ",
                         paste0(g_class, collapse = ""), "/",
                         paste0(o_class, collapse = ""), "/",
                         paste0(f_class, collapse = ""), ". ",
                         "Iteration time: ", round(itr_time, 3), ", ",
                         "Final optimization time: ", round(opt_time, 3), ".")
      
      itr_row <- cbind.data.frame(starting, "Optimizer" = "SSF",
                                  "Iterations" = itr, "Iteration time" = itr_time,
                                  "Optimization time" = opt_time)
      
      if (track_swaps) {
        itr_row <- cbind.data.frame(itr_row, "Swap times" = I(list(swap_times)))
      }
      
      itr_table <- rbind(itr_table, itr_row)
      
      write(opt_prnt, stdout())
      optimize <- FALSE
    }
    
  }
  
  dfiles <- Sys.glob(paste0(msa_path, "*.dfile"))
  file.remove(dfiles)
  
  rownames(itr_table) <- seq(nrow(itr_table))
  return(itr_table)
}

optimizeByOGF <- function(msa_path, binomial_test, garpfymink = FALSE, limit = NULL,
                            tree = NULL, iqtree = NULL, rootfile = NULL,
                            mixture = NULL, track_swaps = FALSE, threads = 1) {
  
  if (garpfymink) {
    g_class <- c("G", "A", "R", "P")
    f_class <- c("F", "Y", "M", "I", "N", "K")
    o_class <- generateAlphabet()[!generateAlphabet() %in% c(g_class, f_class)]
  }
  else{
    g_class <- binomial_test$AA[binomial_test$Assignment == "G-class"]
    o_class <- binomial_test$AA[binomial_test$Assignment == "O-class"]
    f_class <- binomial_test$AA[binomial_test$Assignment == "F-class"]
  }
  
  if (is.null(limit)) {
    limit <- Inf
  }
  
  optimize <- TRUE
  itr <- 1
  opt_time <- Sys.time()
  
  itr_table <- data.frame()
  
  while(optimize) {
    
    itr_time <- Sys.time()
    
    swaps <- generateSwaps(g_class, o_class, f_class)
    
    nswaps <- seq(nrow(swaps))
    
    opt_prnt <- paste0("Iteration ", itr, ": Assessing ",
                       length(nswaps), " swaps from starting point ",
                       paste0(g_class, collapse = ""), "/",
                       paste0(o_class, collapse = ""), "/",
                       paste0(f_class, collapse = ""), ".")
    
    write(opt_prnt, stdout())
    
    swaps$Criterion <- mclapply(nswaps, function(i) {
      
      swap <- swaps[i,]
      g_class <- unlist(swap$G)
      f_class <- unlist(swap$F)
      
      swap_time <- Sys.time()
      
      lnl <- GFmix(msa_path, mixture, tree, iqtree, rootfile, g_class, f_class,
                   plusF = FALSE, optimizer = TRUE)
      
      swap_time <- as.numeric(difftime(Sys.time(), swap_time ,units = "secs"))
      
      return(list("lnL" = lnl, "Swap" = swap_time))
    }, mc.cores = threads)
    
    swap_times <- unlist(lapply(swaps$Criterion, function(x) x$Swap))
    swaps$Criterion <- unlist(lapply(swaps$Criterion, function(x) x$lnL))
    
    best <- swaps[which.max(swaps$Criterion),]
    starting <- swaps["Starting",]
    
    if (unlist(best$Criterion) > unlist(starting$Criterion)) {
      
      g_class <- unlist(best$G)
      o_class <- unlist(best$O)
      f_class <- unlist(best$F)
      
      itr_time <- as.numeric(difftime(Sys.time(), itr_time ,units = "secs"))
      curr_time <- as.numeric(difftime(Sys.time(), opt_time ,units = "secs"))
      
      if (itr == limit) {
        
        opt_time <- curr_time
        
        opt_prnt <- paste0("Iteration ", itr, ": G/O/F at iteration limit: ",
                           paste0(g_class, collapse = ""), "/",
                           paste0(o_class, collapse = ""), "/",
                           paste0(f_class, collapse = ""), ". ",
                           "Iteration time: ", round(itr_time, 3), ", ",
                           "Final optimization time: ", round(opt_time, 3), ".")
        
        itr_row <- cbind.data.frame(best, "Optimizer" = "OGF",
                                    "Iterations" = itr, "Iteration time" = itr_time,
                                    "Optimization time" = curr_time)
        
        if (track_swaps) {
          itr_row <- cbind.data.frame(itr_row, "Swap times" = I(list(swap_times)))
        }
        
        itr_table <- rbind(itr_table, itr_row)
        
        write(opt_prnt, stdout())
        optimize <- FALSE
        
      } else {
        
        opt_prnt <- paste0("Iteration ", itr, ": Found new starting point ",
                           paste0(g_class, collapse = ""), "/",
                           paste0(o_class, collapse = ""), "/",
                           paste0(f_class, collapse = ""), ". ",
                           "Iteration time (s): ", round(itr_time, 3), ", ",
                           "Current optimization time (s): ", round(curr_time, 3), ".")
        
        itr_row <- cbind.data.frame(best, "Optimizer" = "OGF",
                                    "Iterations" = itr, "Iteration time" = itr_time,
                                    "Optimization time" = curr_time)
        
        if (track_swaps) {
          itr_row <- cbind.data.frame(itr_row, "Swap times" = I(list(swap_times)))
        }
        
        itr_table <- rbind(itr_table, itr_row)
        
        itr <- itr + 1
        write(opt_prnt, stdout())
      }
      
      
    } else {
      g_class <- unlist(starting$G)
      o_class <- unlist(starting$O)
      f_class <- unlist(starting$F)
      
      itr_time <- as.numeric(difftime(Sys.time(), itr_time ,units = "secs"))
      opt_time <- as.numeric(difftime(Sys.time(), opt_time ,units = "secs"))
      
      opt_prnt <- paste0("Iteration ", itr, ": G/O/F at optimization limit: ",
                         paste0(g_class, collapse = ""), "/",
                         paste0(o_class, collapse = ""), "/",
                         paste0(f_class, collapse = ""), ". ",
                         "Iteration time: ", round(itr_time, 3), ", ",
                         "Final optimization time: ", round(opt_time, 3), ".")
      
      itr_row <- cbind.data.frame(starting, "Optimizer" = "OGF",
                                  "Iterations" = itr, "Iteration time" = itr_time,
                                  "Optimization time" = opt_time)
      
      if (track_swaps) {
        itr_row <- cbind.data.frame(itr_row, "Swap times" = I(list(swap_times)))
      }
      
      itr_table <- rbind(itr_table, itr_row)
      
      write(opt_prnt, stdout())
      optimize <- FALSE
    }
    
  }
  
  rownames(itr_table) <- seq(nrow(itr_table))
  return(itr_table)
  
}

formatOptTable <- function(optimization) {
  
  optimization$G <- lapply(optimization$G, function(i) paste(i, collapse = ""))
  optimization$O <- lapply(optimization$O, function(i) paste(i, collapse = ""))
  optimization$F <- lapply(optimization$F, function(i) paste(i, collapse = ""))
  
  optimization$`Swap times` <- lapply(optimization$`Swap times`, function(i) paste(i, collapse = ","))
  optimization <- apply(optimization, 2, as.character)
  
  return(optimization)
  
}
```
