## Supplementary material for "Modeling site-and-branch-heterogeneity with GFmix": LikelihoodFunctions.R

```
runGFmix <- function(msa_path, mixture, tree, iqtree, rootfile,
                     g_class, f_class, plusF, params) {
  
  
  lnl <- GFmix(msa_path, mixture, tree, iqtree, rootfile, g_class, f_class,
               plusF, optimizer = FALSE, afo = NULL, params = params)
  
  return(lnl)
}

runAnFreqOpt <- function(msa_path, mixture, tree, iqtree_path, rootfile,
                         g_class, f_class, plusF, params) {
  
  msa <- readAln(msa_path, format = format)
  ntaxa <- nrow(msa)
  
  iqtree <- parseIQTREE(iqtree_path)
  alpha <- iqtree$Alpha
  w_vector <- unlist(iqtree$Weights)
  w_filename <- paste0(msa_path, ".wfile")
  write(w_vector, w_filename, sep = "\n")
  
  b_vector <- rep(1, (2 * ntaxa) - 1)
  b_filename <- paste0(msa_path, ".bfile")
  write(b_vector, b_filename, sep = "\n")
  
  utree <- treeCNS(msa_path, tree)
  
  root <- as.numeric(strsplit1(readLines(rootfile), " "))
  root <- LocateRoot(utree, root)
  
  utreec <- rert(utree, root, ntaxa, msa_path)
  utreec_path <- paste0(msa_path, ".utreec")
  
  g_class <- strsplit1(g_class, split = "")
  f_class <- strsplit1(f_class, split = "")
  
  afo <- AnFreqOpt(msa_path, mixture, utreec_path, alpha,
                   w_filename, b_filename, g_class, f_class,
                   ss = FALSE)
  
  lnl <- GFmix(msa_path, mixture, tree, iqtree_path, rootfile, g_class, f_class,
               plusF, optimizer = FALSE, afo = afo, params = params)
  
  file.remove(w_filename, b_filename, utreec_path, paste0(utreec_path, "u", collapse = NULL))
  
  return(lnl)
}

runGFmixLnL <- function(msa_path, mixture, tree, iqtree_path, rootfile,
                        g_class, f_class, mode = 4, plusF, params, multis) {
  
  msa <- readAln(msa_path, format = format)
  ntaxa <- nrow(msa)
  
  iqtree <- parseIQTREE(iqtree_path)
  alpha <- iqtree$Alpha
  w_vector <- unlist(iqtree$Weights)
  w_filename <- paste0(msa_path, ".wfile")
  write(w_vector, w_filename, sep = "\n")
  
  b_vector <- rep(1, (2 * ntaxa) - 1)
  b_filename <- paste0(msa_path, ".bfile")
  write(b_vector, b_filename, sep = "\n")
  
  utree <- treeCNS(msa_path, tree)
  
  root <- as.numeric(strsplit1(readLines(rootfile), " "))
  root <- LocateRoot(utree, root)
  
  utreec <- rert(utree, root, ntaxa, msa_path)
  utreec_path <- paste0(msa_path, ".utreec")
  
  g_class <- strsplit1(g_class, split = "")
  f_class <- strsplit1(f_class, split = "")
  
  lnl <- NewGFmix(msa_path, mixture, utreec_path, alpha, w_filename,
                  g_class, f_class, mode, params, multis)
  
  return(lnl)
  
}
```
