## Supplementary material for "Modeling site-and-branch-heterogeneity with GFmix": PerformanceFunctions.R

```
generateTrueNodeFreqs <- function(mixture, ntaxa) {
  
  alpha <- 0.5
  
  if (ntaxa == 16) {

    utreec <- "sim_topologies/16_taxa.utreec"
    bfile <- "sim_topologies/16_taxa.bfile"
    
    row_names <- c("A1", "A2", "A3", "A4",
                   "B1", "B2", "B3", "B4",
                   "C1", "C2", "C3", "C4",
                   "D1", "D2", "D3", "D4",
                   "C1+C2", "C3+C4", "D1+D2", "D3+D4",
                   "B1+B2", "B3+B4", "C-stem", "D-stem",
                   "A3+A4", "B-stem", "C+D", "A1+A2",
                   "A-stem", "A+B", "Root")
    
  } else if (ntaxa == 4) {
    
    utreec <- "sim_topologies/4_taxa.utreec"
    bfile <- "sim_topologies/4_taxa.bfile"
    
    row_names <- c("A", "B", "C", "D", 
                   "C+D", "A+B", "Root")
    
  }
  
  wtfile <- "sim_topologies/C20.wtfile"
  
  af <- AnFreq(utreec, wtfile, mixture, alpha, bfile)
  colnames(af) <- generateAlphabet()
  rownames(af) <- row_names
  
  return(af)
}

parseGFmixParams <- function(params, ntaxa) {
  
  lines <- readLines(params)
  
  alpha <- as.numeric(lines[1])
  
  utreec <- lines[2:ntaxa]
  utreec <- as.matrix(read.table(text = utreec))
  colnames(utreec) <- NULL
  
  weights <- lines[ntaxa + 1]
  weights <- as.numeric(strsplit1(weights, " "))
  
  b <- as.numeric(lines[(ntaxa + 2):length(lines)])
  
  params <- list("alpha" = alpha,
                 "utreec" = utreec,
                 "weights" = weights,
                 "b" = b)
  
  return(params)  
}

parseGFmixLnLParams <- function(params, ntaxa) {
  
  lines <- readLines(params)
  
  alpha <- as.numeric(lines[1])
  
  utreec <- lines[2:ntaxa]
  utreec <- as.matrix(read.table(text = utreec))
  colnames(utreec) <- NULL

  weights <- lines[ntaxa + 1]
  weights <- as.numeric(strsplit1(weights, " "))
  
  cG <- lines[(ntaxa + 2):length(lines)]
  cG <- lapply(cG, function(line) {
    values <- as.numeric(strsplit1(line, " "))
    list(cG = values[1], cF = values[2])
  })
  cF <- sapply(cG, function(x) x$cF)
  cG <- sapply(cG, function(x) x$cG)
  
  params <- list("alpha" = alpha,
                 "utreec" = utreec,
                 "weights" = weights,
                 "cG" = cG,
                 "cF" = cF)
  
  return(params)  
}

uTreecToTable <- function(utreec, ntaxa, ids) {
  
  # Get branch length and b-value information for each node first.
  tree_table <- lapply(seq(0, 2 * ntaxa - 2), function(i) {
    
    l <- which(utreec[, 1] == i)
    r <- which(utreec[, 2] == i)
    
    # First condition should capture root node (not defined in utreec, so
    # shouldn't show up in either l or r).
    if (identical(l, r)) {
      tip <- list(0, i + 1, 0, NA)
    }
    else if (identical(r, integer(0))) {
      tip <- list(0, i + 1, utreec[l, 3], ids[i + 1])
    }
    else if (identical(l, integer(0))) {
      tip <- list(0, i + 1, utreec[r, 4], ids[i + 1])
    }
    
    return(tip)
  })
  
  # Use structure of utreec matrix to assign parent nodes.
  # Set parent of root node to 0.
  lapply(seq(ntaxa, 2 * ntaxa - 1), function(i) {
    
    line <- i - (ntaxa - 1)
    if (line <= nrow(utreec)) {
      children <- utreec[line, 1:2]
      
      lapply(children, function(j) {
        tree_table[[j + 1]][1] <<- i + 1
      })
      
      label <- paste0(lapply(children, function(j) {
        tree_table[[j + 1]][[4]]
      }), collapse = "+")
      
      tree_table[[i + 1]][[4]] <<- label
    }
    else {
      tree_table[[i]][1] <<- 0
      tree_table[[i]][4] <<- "Root"
    }
  })
  
  tree_table <- do.call(rbind.data.frame, tree_table)
  columns <- c("parent", "node", "edge.length", "label")
  colnames(tree_table) <- columns
  
  tree_table$group <- NA
  
  get_clade <- function(label) {
    if (label == "Root") {
      unique_clades <- "Root"
    }
    else {
      unique_clades <- sort(unique(strsplit(gsub("[0-9+]", "", label), "")[[1]]))
    }
    return(paste(unique_clades, collapse = "+"))
  }

  tree_table$group <- sapply(tree_table$label, get_clade)
  tree_table$group <- factor(tree_table$group, levels = c("A", "B", "C", "D", "A+B", "C+D", "Root"))

  return(tree_table)
}


# Function taken from Ed Susko.
deriveRateMatrix <- function(exchangabilities, profile) {
  Q <- exchangabilities %*% diag(profile)
  diag(Q) <- -apply(Q, 1, sum)
  mu <- -1 / sum(profile * diag(Q))
  Q <- mu*Q
  return(Q)
}

# Function taken from Ed Susko.
deriveEigens <- function(Q, fr){
  e <- eigen(diag(sqrt(fr)) %*% Q %*% diag(1/sqrt(fr)))
  W <- diag(sqrt(fr)) %*% e$vectors
  Lam <- e$values
  mu <- -1 / sum(diag(W %*% diag(Lam) %*% t(W)))
  Lam <- Lam*mu
  return(list(W=W,Lam=Lam))
}

readSubstitutionMatrix <- function(infile, format = "nexus") {
  alphabet <- generateAlphabet()
  
  lines <- readLines(infile)
  
  if (format == "nexus") {
    
    # Extract substitution probabilities.
    lines <- lines[grep("^[[:digit:]]", lines)]
    lines <- strsplit(gsub(";", "", lines), " ")
    lines <- lapply(lines, function(i) c(as.numeric(i), rep(0, 20 - length(i))))
    
    # Make symmetric matrix.
    exchangabilities <- list(rep(0, 20))
    exchangabilities <- append(exchangabilities, lines)
    exchangabilities <- do.call(rbind.data.frame, exchangabilities)
    exchangabilities <- as.matrix(exchangabilities)
    exchangabilities <- t(exchangabilities) + exchangabilities - diag(diag(exchangabilities))
    colnames(exchangabilities) <- rownames(exchangabilities) <- generateAlphabet()
    
    return(exchangabilities)
  }
}
```
