## Supplementary material for "Modeling site-and-branch-heterogeneity with GFmix": SimulationFunctions.R

```
generateTree <- function(ntaxa = 16, exclude_bl = FALSE) {
  
  if (ntaxa == 16) {
    
    tip <- stem <- 0.5
    zeroAC <- 0.001
    zeroBD <- 0.002
    internal <- 0.25
    
    newick <- paste0("((((A1:", tip, ",A2:", tip, "):", zeroAC, ",(A3:", tip, ",A4:", tip, "):", zeroAC, "):", stem,
                     ",((B1:", tip, ",B2:", tip, "):", zeroBD, ",(B3:", tip, ",B4:", tip, "):", zeroBD, "):", stem,
                     "):", internal,
                     ",(((C1:", tip, ",C2:", tip, "):", zeroAC, ",(C3:", tip, ",C4:", tip, "):", zeroAC, "):", stem,
                     ",((D1:", tip, ",D2:", tip, "):", zeroBD, ",(D3:", tip, ",D4:", tip, "):", zeroBD, "):", stem,
                     "):", internal, ");")
    
    
  }
  else if (ntaxa == 4) {
    
    # Set tip lengths to 1, internal lengths to 0.25.
    tip <- 1
    internal <- 0.25
    
    newick <- paste0("((A:", tip, ",B:", tip, "):", internal,
                     ",(C:", tip, ",D:", tip, "):", internal, ");")
  }
  
  if (exclude_bl) {
    newick <- gsub(":[0-9.\\-eE]+", "", newick)
  }
  
  return(newick)
}

generateGammaRates <- function(alpha = 0.5, k = 4) {
  
  if (k == 1) {
    rates <- 1
  }
  else {
    probs <- (1:k - 1) / k
    
    quantiles <- qgamma(probs, shape = alpha, rate = alpha)
    cdf <- c(0, pgamma(quantiles * alpha, alpha + 1), 1)
    rates <- diff(cdf) * k
  }
  
  rates <- rates[rates > 0]
  
  return(rates)
  
}

generateMixture <- function(mixture = "C20") {
  
  alphabet <- generateAlphabet()
  
  if (mixture == "C20") {
    
    classes <- scan("bin/C20.aafreq.dat", what = double(),
                    sep = " ", quiet = TRUE)
    classes <- na.omit(classes)
    classes <- as.data.frame(matrix(classes, ncol = 20, byrow = TRUE))
    
    weights <- c(0.0559910600, 0.0514824870, 0.0812922124,
                 0.0721976867, 0.0556718858, 0.0331003080,
                 0.0589501763, 0.0263756889, 0.0307584220,
                 0.0376701125, 0.0303058290, 0.0808775576,
                 0.0263349134, 0.0579101455, 0.0371248064,
                 0.0586867766, 0.0561479138, 0.0349810886,
                 0.0544937394, 0.0596471901)
    
    classes <- cbind.data.frame(classes, weights)
    rownames(classes) <- paste0(mixture, "pi", 1:20)
    colnames(classes) <- c(alphabet, "Weight")
  }
  
  return(classes)
}

modifyMixture <- function(mixture = NULL, g_class, f_class, comp_ratio) {
  
  prnt <- paste0("Modifying mixture so ", paste0(g_class, collapse = ""),
                 "/",
                 paste0(f_class, collapse = ""), " ratio = ", comp_ratio, "...")
  write(prnt, stdout())
  
  alphabet <- generateAlphabet()
  
  o_class <- alphabet[!alphabet %in% c(g_class, f_class)]
  
  classes <- mixture[c(-ncol(mixture))]
  
  if (nrow(classes) > 1) {
    sumG <- apply(classes[,alphabet %in% g_class], 1, sum)
    sumF <- apply(classes[,alphabet %in% f_class], 1, sum)
  }
  else {
    sumG <- sum(classes[,alphabet %in% g_class])
    sumF <- sum(classes[,alphabet %in% f_class])
  }
  
  sumO <- 1 - sumG - sumF
  
  cG <- cF <- 1
  mu <- rep(0, nrow(classes))
  wtc <- mixture$Weight
  b <- comp_ratio
  
  # Solve SG, SF and mu such that:
  #   1) The sum of each modified site profile = 1,
  #   2) The sum of all G site profiles divided by
  #      the sum of all F site profiles is equal to b.
  #
  # This routine is derived from the gfmix software written
  # by Ed Susko.
  for (iter in 1:10000) {
    mu <- 1 / (cG * sumG + cF * sumF + sumO)
    bo <- sum(wtc * mu * sumG) / sum(wtc * mu * sumF)
    cF <- (1 + bo) / (1 + b)
    cG <- cF * b / bo
    err <- sum(abs(1 - mu * (cG * sumG + cF * sumF + sumO)))
    err <- err + abs(b - bo * cG / cF)
    if (err < 1.0e-6) break
  }
  
  classes[,alphabet %in% g_class] <- classes[,alphabet %in% g_class] * cG * mu
  classes[,alphabet %in% f_class] <- classes[,alphabet %in% f_class] * cF * mu
  classes[,alphabet %in% o_class] <- classes[,alphabet %in% o_class] * mu
  
  mixture <- cbind.data.frame(classes, "Weight" = wtc)
  
  return(mixture)
}

resampleProfileSites <- function(mixture, nsites) {
    
  minimum <- 3
  
  weights <- mixture$Weight
  weights <- rmultinom(1, nsites, weights)
  
  check <- all(weights > minimum)
  
  while (!check) {
    weights <- rmultinom(1, nsites, weights)
    check <- all(weights > minimum)
  }
  
  mixture$Weight <- drop(weights)
  return(mixture)
  
}

resampleGammaSites <- function(nsites) {
  
  minimum <- 0
  
  weights <- c(0.25, 0.25, 0.25, 0.25)
  
  weights <- rmultinom(1, nsites, weights)
  
  check <- all(weights > minimum)
  
  while (!check) {
    weights <- rmultinom(1, nsites, weights)
    check <- all(weights > minimum)
  }
  
  return(weights)
}

addProfileToNewick <- function(newick, equal_profile, fymink_profile, prefix, ntaxa) {
  
  if (ntaxa == 16) {
    
    # Add "base" frequencies to terminal A and C branches.
    newick <- gsub("([A,C]+\\d*:\\d+(\\.\\d+)?)",
                   paste0("\\1[&model=", prefix, "{",
                          paste0(equal_profile[1:20], collapse = "/"), "}]"), newick)
    
    # Add "modified" frequencies to terminal B and branches.
    newick <- gsub("([B,D]+\\d*:\\d+(\\.\\d+)?)",
                   paste0("\\1[&model=", prefix, "{",
                          paste0(fymink_profile[1:20], collapse = "/"), "}]"), newick)
    
    # Add "base" frequencies to A1-4 and C1-4 branch.
    newick <- gsub("(:+\\d\\.\\d),\\(",
                   paste0("\\1[&model=", prefix, "{",
                          paste0(equal_profile[1:20], collapse = "/"), "}],("), newick)
    
    # Add "modified" frequencies to B1-4 and D1-4 branch.
    newick <- gsub("(:\\d.\\d)\\)",
                   paste0("\\1[&model=", prefix, "{",
                          paste0(fymink_profile[1:20], collapse = "/"), "}])"), newick)
    
    # Add correct frequencies to "tiny" branches in A, C and B, D (replacing
    # placeholder BL for the latter).
    newick <- gsub("(:0.001)", paste0("\\1[&model=", prefix, "{",
                                      paste0(equal_profile[1:20], collapse = "/"), "}]"), newick)
    newick <- gsub("(:0.002)", paste0(":0.001[&model=", prefix, "{",
                                      paste0(fymink_profile[1:20], collapse = "/"), "}]"), newick)
    
    # Add "base" frequencies to internal branches.
    newick <- gsub("(:\\d.\\d*)\\,",
                   paste0("\\1[&model=", prefix, "{",
                          paste0(equal_profile[1:20], collapse = "/"), "}],"), newick)
    newick <- gsub("(:\\d.\\d*)\\)",
                   paste0("\\1[&model=", prefix, "{",
                          paste0(equal_profile[1:20], collapse = "/"), "}])"), newick)
  }
  
  return(newick)
  
}

generateNexus <- function(prefix, equal_profile, rates, gamma_weights, psites) {
  
  profile_name <- rownames(equal_profile)
  profile_freqs <- equal_profile[1:20]
  
  category_sites <- gamma_weights
  
  if (sum(category_sites) != psites) {
    browser()
  }
  
  # Write nexus header.
  nexus <- c("#nexus\nbegin sets;\n")
  
  # Generate rate category definition lines.
  deflines <- paste0(c(paste0("\tcharset ", profile_name, "_", 1:4, " = AA, ")))
  
  positions <- paste0(c(1, cumsum(category_sites)[1:3] + 1),
                      "-", cumsum(category_sites), ";\n")
  
  deflines <- paste0(deflines, positions)
  charpar <- paste0("\tcharpartition ", profile_name, " = ")
  deflines <- append(deflines, charpar)
  nexus <- append(nexus, deflines)
  
  freqlines <- paste0(prefix, "{", paste0(profile_freqs, collapse = ","), "}:",
                      paste0( profile_name, "_", 1:4),
                      paste0("{", rates, "},"))
  
  freqlines[4] <- gsub("},", "};\n", freqlines[4])
  nexus <- append(nexus, freqlines)
  nexus <- append(nexus, "end;")
  
  return(nexus)
  
}

runAliSim <- function(tree, ntaxa, nsites, rates, alpha, equal, fymink, outdir) {
  
  prefix <- "LG+F"
  n_profiles <- nrow(equal)
  
  outdir_path <- file.path(getwd(), outdir)
  
  if (!dir.exists(outdir_path)) {
    dir.create(outdir_path, recursive = TRUE)
  }
  
  for (i in 1:n_profiles) {
    
    equal_profile <- equal[i,]
    fymink_profile <- fymink[i,]
    psites <- equal_profile$Weight
    
    prnt <- paste0(rownames(equal_profile)[1],
                   ": Generating AliSim inputs with new frequencies, ",
                   "profile weight and rate category weights...")
    write(prnt, stdout())
    
    gamma_weights <- resampleGammaSites(psites)
    
    newick <- addProfileToNewick(tree, equal_profile, fymink_profile, prefix, ntaxa)
    nexus <- generateNexus(prefix, equal_profile, rates, gamma_weights, psites)
    
    newick_outfile <- paste0(outdir_path, "/", rownames(equal_profile), ".nwk")
    nexus_outfile <- paste0(outdir_path, "/", rownames(equal_profile), ".nex")
    fasta_outfile <- paste0(outdir_path, "/", rownames(equal_profile))
    
    cat(newick, file = newick_outfile)
    cat(nexus, file = nexus_outfile)
    
    prnt <- paste0(rownames(equal_profile)[1],
                   ": Running AliSim...")
    write(prnt, stdout())
    
    cmd <- paste(c("iqtree3", "--alisim", fasta_outfile, "-t", newick_outfile,
                   "-p", nexus_outfile, "-af fasta -quiet"), collapse = " ")
    
    system(cmd)
  }
  
  prnt <- paste0("Created ", n_profiles, " alignments for each mixture profile.\n",
                 "Profile alignment sites: ", paste0(equal$Weight, collapse = ", "), 
                 ".\nSum sites: ", sum(equal$Weight),
                 ", Target sites: ", nsites, ", Number of taxa: ", ntaxa, ".")
  write(prnt, stdout())
  
  return(outdir_path)
}

concatenateAlignments <- function(indir, outfile, ntaxa = 16) {
  
  fastas <- paste0(indir, "/", "C20pi", 1:20, ".fa")
  
  concatenated <- data.frame(IDs = rep(NA, ntaxa), Alignment = rep(NA, ntaxa))
  
  for (alignment in fastas) {
    
    alignment <- readAln(alignment, "fasta")
    
    if (all(is.na(concatenated$IDs))) {
      concatenated$IDs <- alignment$IDs
    }
    if (all(is.na(concatenated$Alignment))) {
      concatenated$Alignment <- alignment$Alignment
    }
    else {
      concatenated$Alignment <- paste0(concatenated$Alignment, alignment$Alignment)
    }
  }
  
  prnt <- paste0("Concatenated profile alignments into full simulated alignment.")
  write(prnt, stdout())
  
  return(concatenated)
}

downsampleAlignment <- function(alignment) {
  
  alignment <- alignment[c(1, 5, 9, 13),]
  alignment$IDs <- c("A", "B", "C", "D")
  
  prnt <- paste0("Created downsampled alignment.")
  write(prnt, stdout())
  
  return(alignment)
}
```
