## Supplementary material for "Modeling site-and-branch-heterogeneity with GFmix": IdentifyHeterogeneity.R

```
#!/usr/bin/env Rscript
source("functions/CommonFunctions.R")
source("functions/GFmixFunctions.R")
source("functions/IdentificationFunctions.R")

library(parallel)

# Handling arguments.
args <- commandArgs()
iarg <- length(args)

seqfile <- groupfile <- iqtreefile <- treefile <- rootfile <- NULL
mixture <- frmt <- criterion <- limit <- NULL
frmt <- "phylip"
criterion <- "chi"
outfile <- stdout()
convert <- FALSE
  
while(iarg>=5){
  if(substring(args[iarg],1,1)=='-'){
    opt <- args[iarg]
    is.opt <- TRUE
  }else{
    val <- args[iarg];
    is.opt <- FALSE
  }
  if(is.opt){
    not.an.option <- TRUE
    if(opt=="-aln"){
      seqfile <- val; not.an.option <- FALSE
    }
    if(opt=="-fmt"){
      format <- val; not.an.option <- FALSE
    }
    if (opt=="-crit"){
      criterion <- val; not.an.option <- FALSE
    }
    if (opt=="-group"){
      groupfile <- val; not.an.option <- FALSE
    }
    if (opt=="-mix"){
      mixture <- val; not.an.option <- FALSE
    }
    if (opt=="-iqtree"){
      iqtreefile <- val; not.an.option <- FALSE
    }
    if (opt=="-tree"){
      treefile <- val; not.an.option <- FALSE
    }
    if (opt=="-root"){
      rootfile <- val; not.an.option <- FALSE
    }
    if (opt=="-iter"){
      limit <- as.numeric(val); not.an.option <- FALSE
    }
    if (opt=="-out"){
      outfile <- val; not.an.option <- FALSE
    }
    if (opt=="-conv"){
      convert <- TRUE; not.an.option <- FALSE
    }
  }
  iarg <- iarg-1
}

threads <- 10
garpfymink <- FALSE
track_swaps <- TRUE

if (criterion %in% c("ss", "gfmix")) {
  if (any(is.null(mixture), is.null(treefile),
          is.null(iqtreefile), is.null(rootfile))) {
    stop(paste0("Check command line: input missing for SS/lnL optimization",
         " (-mix, -iqtree, -tree or -root)."))
  }
}

# Read alignment and count amino acids.
alignment <- readAln(seqfile, format = frmt)
counts <- countAA(alignment)

if (!is.null(groupfile)) {
  group_1 <- strsplit1(readLines(groupfile), " ")
  
  binomial_test <- data.frame(binomialBinning(group_1 = group_1, counts = counts))
} else {
  chi_residuals <- chisq.test(counts)$residuals
  chi_dend <- hclust(dist(chi_residuals), method = "average")
  
  # Run binomial test and present results.
  binomial_test <- data.frame(binomialBinning(dendrogram = chi_dend, counts = counts))
}

if (convert) {
  
  convertIDs(seqfile, treefile, format)
  treefile <- paste0(treefile, "_INT")
  if (format == "phylip") {
    seqfile <- paste0(seqfile, "_INT.phy")
  }
  
}

if (criterion == "bi") {
  
  g_class <- binomial_test[binomial_test$Assignment == "G-class",]$AA
  o_class <- binomial_test[binomial_test$Assignment == "O-class",]$AA
  f_class <- binomial_test[binomial_test$Assignment == "F-class",]$AA
  
  optimization <- cbind.data.frame("G" = I(list(g_class)), "O" = I(list(o_class)),
                             "F" = I(list(f_class)), "Criterion" = 0,
                             "Optimizer" = "Binomial", "Iterations" = 0,
                             "Iteration time" = 0, "Optimization time" = 0,
                             "Swap times" = I(list(0)))
  
} else if (criterion == "chi") {
  optimization <- optimizeByChi(msa = alignment, binomial_test = binomial_test,
                                 garpfymink = garpfymink,
                                 limit = limit, track_swaps = track_swaps,
                                 threads = threads)
} else if (criterion == "ssf") {
  optimization <- optimizeBySSF(msa = alignment, msa_path = seqfile,
                                binomial_test = binomial_test,
                                garpfymink = garpfymink,
                                limit = limit, track_swaps = track_swaps,
                                tree = treefile,
                                iqtree = iqtreefile, rootfile = rootfile,
                                mixture = mixture,
                                threads = threads)
} else if (criterion == "ogf") {
  optimization <- optimizeByOGF(msa_path = seqfile,
                                   binomial_test = binomial_test,
                                   garpfymink = garpfymink,
                                   limit = limit, track_swaps = track_swaps,
                                   tree = treefile,
                                   iqtree = iqtreefile, rootfile = rootfile,
                                   mixture = mixture,
                                   threads = threads)
}

optimization <- formatOptTable(optimization)

# If output defaults to 1col instead of 1row (for binomial and sometimes chi2), transpose it.
# This is stupid and should be a temporary fix (something to do with apply?).
if (is.null(ncol(optimization))) {
  optimization <- t(optimization)
}


write.table(optimization, file = outfile, sep = "\t", quote = FALSE, row.names = FALSE)
```
