## Supplementary material for "Modeling site-and-branch-heterogeneity with GFmix": MakePlots.R

```
library(ggplot2)
library(ggtree)
library(treeio)
library(patchwork)
library(tidyr)
library(dplyr)
library(reshape2)
library(legendry)

source("functions/CommonFunctions.R")
source("functions/SimulationFunctions.R")
source("functions/GFmixFunctions.R")
source("functions/PerformanceFunctions.R")
source("functions/Ed_Functions.R")

# UTREEVIEWER functions.
uTreecToTable <- function(utreec, ids, ntaxa, b = NULL, cG = NULL, cF = NULL) {
  
  # Get branch length and parameter information for each node first.
  tree_table <- lapply(seq(0, 2 * ntaxa - 2), function(i) {
    
    l <- which(utreec[, 1] == i)
    r <- which(utreec[, 2] == i)
    
    # First condition should capture root node (not defined in utreec, so
    # shouldn't show up in either l or r).
    if (identical(l, r)) {
      tip <- list(0, i + 1, 0, NA)
    }
    else if (identical(r, integer(0))) {
      tip <- list(0, i + 1, utreec[l, 3], ids[i + 1])
    }
    else if (identical(l, integer(0))) {
      tip <- list(0, i + 1, utreec[r, 4], ids[i + 1])
    }
    
    if (!is.null(b)) {
      params <- b[i + 1]
    }
    else {
      params <- c(cG[i + 1], cF[i + 1])
    }
    
    tip <- c(tip, params)
    return(tip)
  })
  
  # Use structure of utreec matrix to assign parent nodes.
  # Set parent of root node to 0.
  lapply(seq(ntaxa, 2 * ntaxa - 1), function(i) {
    
    line <- i - (ntaxa - 1)
    if (line <= nrow(utreec)) {
      children <- utreec[line, 1:2]
      
      lapply(children, function(j) {
        tree_table[[j + 1]][1] <<- i + 1
      })
      
      label <- paste0(lapply(children, function(j) {
        tree_table[[j + 1]][[4]]
      }), collapse = "+")
      
      tree_table[[i + 1]][[4]] <<- label
    }
    else {
      tree_table[[i]][1] <<- 0
      tree_table[[i]][4] <<- "ROOT"
    }
  })
  
  tree_table <- do.call(rbind.data.frame, tree_table)
  columns <- c("parent", "node", "edge.length", "label")
  if (!is.null(b)) {
    columns <- c(columns, "b")
  }
  else if (!is.null(cG)) {
    columns <- c(columns, "cG", "cF")
  }
  
  colnames(tree_table) <- columns
  
  return(tree_table)
}

plotUtree <- function(tree_table, hex = 0.1, title = NULL) {
  
  tree <- as_tibble(tree_table)
  nc <- ncol(tree)
  tree <- tree %>% as.treedata(edge.length, label)
  
  if (nc == 5) {
    utree_plot <- ggtree(tree, aes(color = b)) + geom_tiplab() +
      scale_color_gradient(low = "navy", high = "skyblue") +
      hexpand(hex, direction = 1) + guides() + theme_tree2()
  }
  else if (nc == 6) {
    cG_utree <- ggtree(tree, aes(color = cG)) + geom_tiplab() +
      hexpand(hex, direction = 1) + guides() + theme_tree2() +
      scale_color_gradient(low = "navy", high = "skyblue")
    cF_utree <- ggtree(tree, aes(color = cF)) + geom_tiplab() +
      hexpand(hex, direction = 1) + guides() + theme_tree2() +
      scale_color_gradient(low = "darkred", high = "orange")
    utree_plot <- (cG_utree | cF_utree)
  }
  else {
    utree_plot <- ggtree(tree) + geom_tiplab() +
      hexpand(hex, direction = 1) + guides() + theme_tree2()
  }
  
  if (!is.null(title)) {
    utree_plot <- utree_plot & plot_annotation(title = title)
  }
  
  return(utree_plot)
}

plotFromParams <- function(params, type = "old", ids, hex = 0.1, title = NULL) {
  
  ids <- readLines(ids)
  ntaxa <- length(ids)
  
  if (type == "old") {
    params <- parseGFmixParams(params = params, ntaxa = ntaxa)
    
    utreec <- params$utreec
    b <- params$b
    
    tree_table <- uTreecToTable(utreec = utreec, ids = ids, ntaxa = ntaxa, b = b)
  }
  else {
    params <- parseGFmixLnLParams(params = params, ntaxa = ntaxa)
    
    utreec <- params$utreec
    cG <- params$cG
    cF <- params$cF
    
    tree_table <- uTreecToTable(utreec = utreec, ids = ids, ntaxa = ntaxa, cG = cG, cF = cF)
  }
  
  utreec_plots <- plotUtree(tree_table = tree_table, hex = hex, title = title)
  
  return(utreec_plots)
}

UTREEViewer <- function(params, type = "old", ids, hex, title = "UTreeplot") {
  
  utreeplot <- plotFromParams(params, type, ids, hex, title)
  
  outfile <- paste0(title, ".pdf", collapse = "")
  
  ggsave(outfile, utreeplot, device = "pdf", height = 10, width = 20)
  
  msg <- paste("Plot saved to", outfile)
  print(msg)
}

# Quick function to generate GARP/FYMINK ratios based on AnFreq given a param file.
getRatio <- function(params) {
  
  write.table(params$utreec, "utreec", row.names = FALSE, col.names = FALSE)
  write(format(params$weights, digits = 17, scientific = TRUE), ncol = 1, file = "weights")
  write(params$b, "bfile", sep = "\n")
  
  af <- AnFreq("utreec", "weights", "bin/C20.aafreq.dat", params$alpha, "bfile")
  colnames(af) <- generateAlphabet()
  
  params$ratio <- rowSums(af[,c("G", "A", "R", "P")]) /
    rowSums(af[,c("F", "Y", "M", "I", "N", "K")])
  
  file.remove("bfile", "utreec", "weights")
  
  return(params)
  
}

# Trees figure.
simulatingTrees <- function() {
  tree_16 <- generateTree(ntaxa = 16)
  tree_4 <- generateTree(ntaxa = 4)
  
  tree_16 <- gsub(":0.002", ":0.001", tree_16)
  
  t1 <- read.tree(text = tree_16)
  t2 <- read.tree(text = tree_4)
  
  t1 <- groupClade(as_tibble(t1), c(22, 29), group_name = "High")
  t1$High <- as.numeric(t1$High)
  t1$High <- ifelse(t1$High > 1, "π(GARP) < π(FYMINK)", "π(GARP) = π(FYMINK)")
  t1 <- t1 %>% as.treedata(High, branch.length)
  t1@phylo$root.edge <- 0.01
  
  t2 <- groupClade(as_tibble(t2), c(2, 4), group_name = "High")
  t2$High <- as.numeric(t2$High)
  t2$High <- ifelse(t2$High > 1, "π(GARP) < π(FYMINK)", "π(GARP) = π(FYMINK)")
  t2 <- t2 %>% as.treedata(High, branch.length)
  t2@phylo$root.edge <- 0.01
  
  g1 <- ggtree(t1, aes(color = High), size= 1.25) + geom_tiplab(fontface = "bold", size = 5) +
    geom_rootedge(size = 1.25) +
    geom_text(aes(x = branch, vjust = 1.5,
                  label = ifelse(branch.length > 0.05, branch.length, "")),
              show.legend = FALSE, fontface = "bold", size = 5) +
    labs(color = "Composition") + 
    scale_color_manual(values = c("gray50", "black")) +
    theme(legend.position = "none")
    
  g2 <- ggtree(t2, aes(color = High), size = 1.25) + geom_tiplab(fontface = "bold", size = 5) +
    geom_rootedge(size = 1.25) +
    geom_text(aes(x = branch, vjust = 1.5,
                  label = ifelse(branch.length > 0.05, branch.length, "")),
              show.legend = FALSE, fontface = "bold", size = 5) +
    labs(color = "Composition") + 
    scale_color_manual(values = c("gray50", "black")) +
    theme(legend.position = "none")
  
  plot <- g1 / g2
  
  return(plot)

}

# Binning figure.
assessPatterns <- function(optimization) {

  alphabet <- c("G", "A", "R", "P",
                "D", "C", "Q", "E", "H", "L", "S", "T", "W", "V",
                "F", "Y", "M", "I", "N", "K")
  
  nopt <- length(unique(optimization$Optimizer))
  nseq <- nrow(optimization) / nopt
  
  optimization <- cbind("Replicate" = rep(1:nseq, each = nopt), optimization)
  
  rep_results <- lapply(seq(nseq), function(i) {
    
    replicate <- optimization[optimization$"Replicate" == i,]
    replicate <- replicate[,c("G", "O", "F", "Optimizer")]
    
    opt_results <- lapply(seq(nopt), function(j) {
      
      opt <- replicate[j,]
      assignment <- lapply(alphabet, function(a) {
        if (grepl(a, opt$G)) {
          0
        } else if (grepl(a, opt$F)) {
          1
        } else {
          2
        }
      })
      
      opt_result <- cbind.data.frame("Replicate" = i, "AA" = alphabet,
            "Assignment" = unlist(assignment), "Optimizer" = opt$Optimizer)
      
    })
    
    opt_results <- do.call(rbind, opt_results)
  })
  
  results <- do.call(rbind, rep_results)
  results$AA <- factor(results$AA, levels = alphabet)
  
  results <- results %>% group_by(Replicate, Optimizer) %>%
    pivot_wider(names_from = AA, values_from = Assignment)
  
  occurrences <- results %>% group_by(across(G:K), Optimizer) %>% count() %>%
    pivot_wider(names_from = Optimizer, values_from = n,
                values_fill = 0)
  
  n_occurrences <- nrow(occurrences)
  
  occurrences$Correct <- lapply(seq(n_occurrences), function(i) {
    
    row <- occurrences[i,]
    correct <- sum(row[1:4]) == 0 &
      sum(row[5:14]) == 20 & sum(row[15:20]) == 6
    
    })
  
  occurrences$Correct <- unlist(occurrences$Correct)
  occurrences <- cbind("Pattern" = seq(nrow(occurrences)), occurrences)
  
  patterns <- melt(occurrences[1:21], id.vars = "Pattern")
  colnames(patterns) <- c("Pattern", "AA", "Assigned")
  patterns$Expected <- ifelse(patterns$AA %in% c("G", "A", "R", "P"), "Expected G",
                     ifelse(patterns$AA %in% c("F", "Y", "M", "I", "N", "K"),
                            "Expected F", "Expected O"))
  patterns$Expected <- factor(patterns$Expected, levels = c("Expected G", "Expected O",
                                          "Expected F"))
  patterns$Assigned <- factor(patterns$Assigned, levels = c(0, 2, 1))
  patterns$Observed <- ifelse(patterns$Assigned == 0, "Observed G",
                             ifelse(patterns$Assigned == 1, "Observed F", "Observed O"))
  patterns$Observed <- factor(patterns$Observed, levels = c("Observed G", "Observed O",
                                                "Observed F"))
  patterns$Correct <- ifelse(substring(patterns$Expected, 10) == substring(patterns$Observed, 10),
                             TRUE, FALSE)

  patterns <- patterns %>% group_by(Pattern) %>% mutate(Falses = sum(!Correct)) %>% ungroup()
  patterns <- patterns %>% mutate(Pattern = reorder(Pattern, -Falses))
  occurrences$Pattern <- factor(occurrences$Pattern, levels = levels(patterns$Pattern))
  
  heatmap <- ggplot(patterns, aes(interaction(x = AA, Expected, sep = "&"),
                               y = Pattern, fill = Assigned)) +
    geom_tile(aes(color = ifelse(Assigned == 0, "white", "black"))) +
    scale_fill_manual(values = c("black", "white", "gray"),
                      labels = c("Observed G", "Observed O", "Observed F"),
                      guide = guide_legend(override.aes = list(color = "black"))) +
    scale_color_identity() +
    geom_text(aes(label = ifelse(Pattern == which(occurrences$Correct),  "\u2713", "")),
              color = "green", fontface = "bold", size = 5) +
    theme_classic() + coord_equal() +
    guides(x = guide_axis_nested(key = "&")) +
    labs(x = "Amino acid", y = "Assigned G/O/F across methods",
         fill = "Assigment") +
    theme(axis.text.y = element_blank(), axis.ticks.y = element_blank(),
          legend.position = "bottom")
    
  counts <- melt(occurrences, id.vars = "Pattern",
                 measure.vars = unique(optimization$Optimizer))
  
  counts$variable <- factor(counts$variable,
                            levels = c("Binomial", "Chi2", "SSF", "OGF"))
  
  nplot <- ggplot(counts, aes(y = Pattern, x = variable, fill = value)) +
    geom_tile(color = "black") + geom_text(aes(label = ifelse(value > 0, value, NA),
                                               color = ifelse(value > nseq / 2, "white", "black"))) +
    scale_color_identity() +
    scale_fill_viridis_c(direction = -1,
                         guide = guide_legend(override.aes = list(color = "black"))) +
    facet_wrap(~variable, nrow = 1, scales = "free_x") + theme_classic() +
    theme(axis.text = element_blank(), axis.line = element_blank(), axis.title.y = element_blank(),
          axis.ticks = element_blank(), legend.position = "bottom") +
    labs(x = paste0("Counts observed across ",
                    nseq, " replicates per method", collapse = ""),
                                               fill = "Count")
  
  plot <- (heatmap | nplot) + plot_layout(guides = "collect") & theme(legend.position = "bottom")
  return(plot)
}

# Timing figures.
assessOpts <- function(optimization) {
  
  optimization <- optimization[optimization$Optimizer != "Binomial",]
  nopt <- length(unique(optimization$Optimizer))
  nseq <- nrow(optimization) / nopt
  optimization <- cbind("Replicate" = rep(1:nseq, each = nopt), optimization)
  optimization$Optimizer <- factor(optimization$Optimizer, levels = c("Chi2", "SSF", "OGF"))
 
  
  iterations <- melt(optimization, id.vars = c("Replicate", "Optimizer"), measure.vars = "Iterations")
  itr_plot <- ggplot(iterations, aes(y = Optimizer, x = value)) +
    geom_point(shape = 1,
               stat = "summary", fun = "mean", size = 3,
               position = position_dodge(width = 0.9)) +
    stat_summary(
      aes(group = Optimizer),
      fun.data = function(x) {
        mean_val <- mean(x)
        sd_val <- sd(x)
        ymin <- mean_val - sd_val
        ymax <- mean_val + sd_val
        return(data.frame(y = mean_val, ymin = ymin, ymax = ymax))
      },
      geom = "errorbarh",
      position = "dodge"
    ) + theme_classic() + labs(x = "Iterations until G/O/F optimized per 100 replicates.")
  
  fulltime <- melt(optimization, id.vars = c("Replicate", "Optimizer"), measure.vars = "Optimization time")
  ft_plot <- ggplot(fulltime, aes(y = Optimizer, x = value)) + 
    geom_point(shape = 0,
               stat = "summary", fun = "mean", size = 3,
               position = position_dodge(width = 0.9)) +
    stat_summary(
      aes(group = Optimizer),
      fun.data = function(x) {
        mean_val <- mean(x)
        sd_val <- sd(x)
        ymin <- mean_val - sd_val
        ymax <- mean_val + sd_val
        return(data.frame(y = mean_val, ymin = ymin, ymax = ymax))
      },
      geom = "errorbarh",
      position = "dodge"
    ) + theme_classic() + labs(x = "Total optimization time (in seconds) per 100 replicates.")
  
  ittime <- melt(optimization, id.vars = c("Replicate", "Optimizer"), measure.vars = "Iteration time")
  it_plot <- ggplot(ittime, aes(y = Optimizer, x = value)) + 
    geom_point(shape = 0,
               stat = "summary", fun = "mean", size = 3,
               position = position_dodge(width = 0.9)) +
    stat_summary(
      aes(group = Optimizer),
      fun.data = function(x) {
        mean_val <- mean(x)
        sd_val <- sd(x)
        ymin <- mean_val - sd_val
        ymax <- mean_val + sd_val
        return(data.frame(y = mean_val, ymin = ymin, ymax = ymax))
      },
      geom = "errorbarh",
      position = "dodge"
    ) + theme_classic() + labs(x = "Time for final optimization iteration (in seconds) per 100 replicates.")
  
  
  swaps <- lapply(optimization$`Swap times`, function(i) as.numeric(strsplit1(i, ",")))
  
  swaptimes <- lapply(seq(nrow(optimization)), function(i) {
    
    rep <- optimization[i,]$Replicate
    opt <- optimization[i,]$Optimizer
    swap <- swaps[[i]]
    
    times <- cbind.data.frame("Replicate" = rep, "Optimizer" = opt, "Swaptime" = swap)
  })
  swaptimes <- do.call(rbind, swaptimes)
  
  swapplot <- ggplot(swaptimes, aes(y = Optimizer, x = Swaptime)) +
    geom_point(shape = 0, stat = "summary", fun = "mean", size = 3,
               position = position_dodge(width = 0.9)) +
    stat_summary(
      aes(group = Optimizer),
      fun.data = function(x) {
        mean_val <- mean(x)
        sd_val <- sd(x)
        ymin <- mean_val - sd_val
        ymax <- mean_val + sd_val
        return(data.frame(y = mean_val, ymin = ymin, ymax = ymax))
      },
      geom = "errorbarh",
      position = "dodge"
    ) + theme_classic() + labs(x = "Swap times for final iteration (in seconds) per 100 replicates.")
    
  
  plots <- (itr_plot) / (ft_plot) / (it_plot) / (swapplot)
  
  return(plots)
  
}

# BL vs. AA freq figure.
blenVsFreq <- function() {
  
  bl_vector <- seq(0, 10, length = 1000)
  overall_freqs <- matrix(0, length(bl_vector), 20)
  lg <- readSubstitutionMatrix("bin/LG.nex")
  
  mixture <- generateMixture(mixture = "C20")
  g_class <- c("G", "A", "R", "P")
  f_class <- c("F", "Y", "M", "I", "N", "K")
  
  starting_profiles <- modifyMixture(mixture, g_class, f_class, 1)
  shifted_profiles <- modifyMixture(mixture, g_class, f_class, 0.1)
  
  for (i in seq(nrow(mixture))) {

    starting_profile <- unlist(starting_profiles[i,][1:20])
    shifted_profile <- unlist(shifted_profiles[i,][1:20])
    
    shifted_q <- deriveRateMatrix(lg, shifted_profile)
    colnames(shifted_q) <- generateAlphabet()
    
    eigens <- deriveEigens(shifted_q, shifted_profile)
    v <- t(starting_profile) %*% eigens$W
    
    time_freq <- matrix(0, length(bl_vector), 20)
    colnames(time_freq) <- generateAlphabet()
    
    for(itv in 1:length(bl_vector)){
      P <- eigens$W %*% diag(exp(eigens$Lam * bl_vector[itv])) %*% t(eigens$W) / shifted_profile
      time_freq[itv,] <- t(starting_profile) %*% P
    }
    
    overall_freqs <- overall_freqs + (time_freq * starting_profiles$Weight[i])
    
  }
  
  melted_time <- melt(overall_freqs)
  names(melted_time) <- c("Time", "AA", "Frequency")
  melted_time$Time <- rep(bl_vector, 20)
  melted_time$Class <- ifelse(melted_time$AA %in% g_class, "G-class",
                              ifelse(melted_time$AA %in% f_class, "F-class",
                              "O-class"))
  melted_time$Class <- factor(melted_time$Class, levels = c("G-class", "O-class", "F-class"))
  melted_time$Label <- ifelse(melted_time$Time == 0, as.character(melted_time$AA),
                              ifelse(melted_time$Time == 10, as.character(melted_time$AA),
                                     NA))
  
  
  frequency_plot <- ggplot(melted_time, aes(x = Time, y = Frequency, group = AA, label = Label)) +
    geom_line(aes(size = ifelse(AA %in% c("L", "V", "S"), 1, 0.5))) +
    geom_label() + facet_wrap(~Class) + theme_classic() +
    labs(y = "Amino acid frequency under default GFmix model", x = "Branch length") +
    theme(legend.position = "bottom") + scale_size_identity()
  
  return(frequency_plot)
}

# Function for plotting the empirical trees.
empiricalTrees <- function() {
  
  apart <- read.tree("Nucleomorphs/16_NM_Apart.tree")
  together <- read.tree("Nucleomorphs/16_NM_Together.tree")
  
  ids <- readLines("Nucleomorphs/Old_Sequences_IDs.txt")
  
  red <- ids[1:9]
  green <- ids[c(11,12,13,15,16)]
  
  apart$tip.label <- ids[c(15,16,13,14,11,12,2,1,3,4,5,8,6,7,9,10)]
  together$tip.label <- ids[c(15,16,13,11,12,2,1,3,4,5,8,6,7,9,14,10)]
  
  gapart <- ggtree(apart, size = 1.5) +
    geom_tiplab(aes(color = ifelse(label %in% red, "red3", 
                                   ifelse(label %in% green, "royalblue", "black"))), size = 4, fontface = "bold") +
    hexpand(0.75) + scale_color_identity(guide = "none") +
    geom_rootpoint(size = 4)
  gtogether <- ggtree(together, size = 1.5) +
    geom_tiplab(aes(color = ifelse(label %in% red, "red3",
                                   ifelse(label %in% green, "royalblue", "black"))), size = 4, fontface = "bold") +
    hexpand(0.75) + scale_color_identity(guide = "none") +
    geom_rootpoint(size = 4)
  
  eplot <- (gapart | gtogether)
  return(eplot)
}

# Function for comparing node frequencies across different GFmix implementations.
compareNodeFreqs <- function(ntaxa = 16) {
  
  if (ntaxa == 16) {
    ids <- c("A1", "A2", "A3", "A4",
                   "B1", "B2", "B3", "B4",
                   "C1", "C2", "C3", "C4",
                   "D1", "D2", "D3", "D4",
                   "C1+C2", "C3+C4", "D1+D2", "D3+D4",
                   "B1+B2", "B3+B4", "C-stem", "D-stem",
                   "A3+A4", "B-stem", "C+D", "A1+A2",
                   "A-stem", "A+B", "Root")
    
    ogf <- Sys.glob("Simulations/*_performance/OGF.params")
    ogf <- ogf[!grepl("downsampled", ogf)]
    ssf <- Sys.glob("Simulations/*_performance/SSF.params")
    ssf <- ssf[!grepl("downsampled", ssf)]
    gfp <- Sys.glob("Simulations/*_performance/GFP.params")
    gfp <- gfp[!grepl("downsampled", gfp)]
    gff <- Sys.glob("Simulations/*_performance/GFF.params")
    gff <- gff[!grepl("downsampled", gff)]
    
  }
  
  else if (ntaxa == 4) {
    ids <- c("A", "B", "C", "D", "C+D", "A+B", "Root")
    
    ogf <- Sys.glob("Simulations/*downsampled_performance/OGF.params")
    ssf <- Sys.glob("Simulations/*downsampled_performance/SSF.params")
    gfp <- Sys.glob("Simulations/*downsampled_performance/GFP.params")
    gff <- Sys.glob("Simulations/*downsampled_performance/GFF.params")
    
  }
    
  trues <- generateTrueNodeFreqs("bin/C20.aafreq.dat", ntaxa)
  trues <- rowSums(trues[,c("G", "A", "R", "P")]) /
      rowSums(trues[,c("F", "Y", "M", "I", "N", "K")])
  mixture <- generateMixture(mixture = "C20")
    
  ogf <- lapply(ogf, function(i) {
      params <- parseGFmixParams(i, ntaxa)
      
      params <- getRatio(params)
      params$diff <- params$ratio - trues
      params
    })
    
  ssf <- lapply(ssf, function(i) {
      params <- parseGFmixParams(i, ntaxa)
      
      params <- getRatio(params)
      params$diff <- params$ratio - trues
      params
    })
    
  gfp <- lapply(gfp, function(i) {
      params <- parseGFmixLnLParams(i, ntaxa)
      
      params$b <- (params$cG * sum(params$weights * mixture[,c("G", "A", "R", "P")])) /
        (params$cF * sum(params$weights * mixture[,c("F", "Y", "M", "I", "N", "K")]))
      
      params <- getRatio(params)
      params$diff <- params$ratio - trues
      params
    })
  
  gff <- lapply(gff, function(i) {
      params <- parseGFmixLnLParams(i, ntaxa)
      
      params$b <- (params$cG * sum(params$weights * mixture[,c("G", "A", "R", "P")])) /
        (params$cF * sum(params$weights * mixture[,c("F", "Y", "M", "I", "N", "K")]))
      
      params <- getRatio(params)
      params$diff <- params$ratio - trues
      params
    })
  
  rogf <- do.call(rbind, lapply(ogf, function(i) i$ratio))
  rogf <- cbind.data.frame("Replicate" = 1:nrow(rogf), "Model" = "OGF", rogf)
  
  rssf <- do.call(rbind, lapply(ssf, function(i) i$ratio))
  rssf <- cbind.data.frame("Replicate" = 1:nrow(rssf), "Model" = "SSF", rssf)
  
  rgfp <- do.call(rbind, lapply(gfp, function(i) i$ratio))
  rgfp <- cbind.data.frame("Replicate" = 1:nrow(rgfp), "Model" = "GFP", rgfp)
  
  rgff <- do.call(rbind, lapply(gff, function(i) i$ratio))
  rgff <- cbind.data.frame("Replicate" = 1:nrow(rgff), "Model" = "GFF", rgff)
    
  ogf <- do.call(rbind, lapply(ogf, function(i) i$diff))
  ogf <- cbind.data.frame("Replicate" = 1:nrow(ogf), "Model" = "OGF", ogf)
    
  ssf <- do.call(rbind, lapply(ssf, function(i) i$diff))
  ssf <- cbind.data.frame("Replicate" = 1:nrow(ssf), "Model" = "SSF", ssf)
    
  gfp <- do.call(rbind, lapply(gfp, function(i) i$diff))
  gfp <- cbind.data.frame("Replicate" = 1:nrow(gfp), "Model" = "GFP", gfp)
    
  gff <- do.call(rbind, lapply(gff, function(i) i$diff))
  gff <- cbind.data.frame("Replicate" = 1:nrow(gff), "Model" = "GFF", gff)
  
  all_models <- rbind(ogf, ssf, gfp, gff)
  
  nfs <- melt(all_models, id.vars = c("Replicate", "Model"))
  colnames(nfs)[3:4] <- c("Node", "Difference")
  
  rmses <- rbind(rogf, rssf, rgfp, rgff)
  colnames(rmses) <- colnames(all_models)
  rmses <- melt(rmses, id.vars = c("Replicate", "Model"))
  colnames(rmses)[3:4] <- c("Node", "Ratio")  
  
  if (ntaxa == 16) {
      retain <- c("A1", "B1", "C1", "D1", "A+B", "C+D", "A-stem", "B-stem", "C-stem", "D-stem", "Root")
      
      nfs <- subset(nfs, nfs$Node %in% retain)
      nfs$Node <- factor(nfs$Node,
                                  levels =  retain)
      nfs$Group <- ifelse(grepl("1", nfs$Node), "Terminal",
                          ifelse(grepl("stem", nfs$Node), "Stem", "Internal"))
      nfs$Model <- factor(nfs$Model, levels = c("OGF", "SSF", "GFP", "GFF"))
      nfs$Group <- factor(nfs$Group, levels = c("Terminal", "Stem", "Internal"))
      
      rmses <- subset(rmses, rmses$Node %in% retain)
      rmses$Node <- factor(rmses$Node,
                         levels =  retain)
      
      rmses <- rmses %>% mutate(group = ifelse(Node %in% c("A1", "C1"), "ACTips",
                                               ifelse(Node %in% c("A-stem", "C-stem"), "ACStem",
                                               ifelse(Node %in% c("B-stem", "D-stem"), "BDStem", 
                                               ifelse(Node %in% c("B1", "D1"), "BDTips",
                                                      ifelse(Node %in% c("Root"), "Root", "Int"))))))
      rmses$group <- factor(rmses$group, levels = c("ACTips", "ACStem", "BDTips", "BDStem", "Int", "Root"))
      
      rmses <- rmses %>% mutate(True = trues[as.character(Node)])
      
      rmses <- rmses %>% group_by(group, Model) %>% summarize(rmse = sqrt(mean((Ratio - True)^2)))
      rmses <- rmses %>% pivot_wider(names_from = group, values_from = rmse)
      
    }
    
  else if (ntaxa == 4) {
    
    rmses <- rmses %>% mutate(group = ifelse(Node %in% c("A", "C"), "AC",
                                         ifelse(Node %in% c("B", "D"), "BD",
                                                ifelse(Node %in% c("Root"), "Root", "Int"))))
    rmses$group <- factor(rmses$group, levels = c("AC", "BD", "Int", "Root"))
    rmses$True <- trues[rmses$Node]
    rmses <- rmses %>% group_by(group, Model) %>% summarize(rmse = sqrt(mean((Ratio - True)^2)))
    rmses <- rmses %>% pivot_wider(names_from = group, values_from = rmse)
    rmses$Model <- factor(rmses$Model, levels = c("OGF", "SSF", "GFP", "GFF"))  
    nfs$Node <- factor(nfs$Node, levels = c("A", "B", "C", "D", "A+B", "C+D", "Root"))
    nfs$Group <- ifelse(grepl("\\+", nfs$Node), "Internal",
                                  ifelse(nfs$Node == "Root", "Internal", "Terminal"))
    nfs$Model <- factor(nfs$Model, levels = c("OGF", "SSF", "GFP", "GFF"))
    nfs$Group <- factor(nfs$Group, levels = c("Terminal", "Internal"))
      
  }
    
  nplot <- ggplot(nfs, aes(y = Node, x = Difference)) +
      geom_vline(xintercept = 0, linetype = "dotted", color = "grey", size = 1) +
      geom_point(aes(shape = Model, color = Model), stat = "summary", fun = "mean", size = 3, position = position_dodge(width = 0.9)) +
      stat_summary(
        aes(group = Model),
        fun.data = function(x) {
          mean_val <- mean(x)
          sd_val <- sd(x)
          ymin <- mean_val - sd_val
          ymax <- mean_val + sd_val
          return(data.frame(y = mean_val, ymin = ymin, ymax = ymax))
        },
        geom = "errorbarh",
        position = "dodge"
      ) +
      scale_shape_manual(name = "Model", labels = c("OGF", "SSF", "GFP", "GFF"),
                         values = c(22, 23, 24, 25)) +
      scale_color_manual(name = "Model", labels = c("OGF", "SSF", "GFP", "GFF"),
                         values=c('#ff4444','#448cff', '#8644ff', '#db44ff')) + theme_classic() +
      labs(y = "Node types", x = "GARP/FYMINK ratio: Estimated - True") +
      facet_wrap(~Group, scales = "free_y", ncol = 1) + theme(legend.position = "bottom", axis.title.y = element_blank()) +
      scale_y_discrete(limits = rev)
  
    return(list("rmse" = rmses, "plot" = nplot))
}

# Function for comparing branch lengths with/without ML re-optimization.
compareBranchLengths <- function(ntaxa = 16, nreps = 100) {
  
  if (ntaxa == 16) {
    
    ids <- c("A1", "A2", "A3", "A4",
             "B1", "B2", "B3", "B4",
             "C1", "C2", "C3", "C4",
             "D1", "D2", "D3", "D4",
             "C1+C2", "C3+C4", "D1+D2", "D3+D4",
             "B1+B2", "B3+B4", "C-stem", "D-stem",
             "A3+A4", "B-stem", "C+D", "A1+A2",
             "A-stem", "A+B", "Root")
    
    uml <- Sys.glob("Simulations/*_performance/OGF.params")
    uml <- uml[!grepl("downsampled", uml)]
    gff <- Sys.glob("Simulations/*_performance/GFF.params")
    gff <- gff[!grepl("downsampled", gff)]
  }
  
  else if (ntaxa == 4) {
    
    ids <- c("A", "B", "C", "D", "C+D", "A+B", "Root")
    
    uml <- Sys.glob("Simulations/*downsampled_performance/OGF.params")
    gff <- Sys.glob("Simulations/*downsampled_performance/GFF.params")

  }
  
  uml <- lapply(uml, function(i) {
    params <- parseGFmixParams(i, ntaxa)
    uTreecToTable(params$utreec, ids, ntaxa)
  })
  
  gff <- lapply(gff, function(i) {
    params <- parseGFmixLnLParams(i, ntaxa)
    uTreecToTable(params$utreec, ids, ntaxa)
  })
  
  uml <- do.call(rbind, uml)
  uml <- cbind.data.frame("Replicate" = rep(seq(1, nreps), each = 2*ntaxa-1), "Model" = "UML", uml)
  
  gff <- do.call(rbind, gff)
  gff <- cbind.data.frame("Replicate" = rep(seq(1, nreps), each = 2*ntaxa-1), "Model" = "GFF", gff)
  
  all_models <- rbind(uml, gff)
    
  if (ntaxa == 16) {

      retain <- c("A1", "B1", "C1", "D1", "B1+B2+B3+B4+A3+A4+A1+A2", "C1+C2+C3+C4+D1+D2+D3+D4",
                  "A3+A4+A1+A2", "B1+B2+B3+B4", "C1+C2+C3+C4", "D1+D2+D3+D4")
      new_ids <- c("A1", "B1", "C1", "D1", "A+B", "C+D", "A-stem", "B-stem", "C-stem", "D-stem")
    
      all_models <- subset(all_models, all_models$label %in% retain)
      all_models$label <- as.character(all_models$label)

      for (i in seq(retain)) {
        pattern <- gsub("\\+", "\\\\+", retain[i])
        all_models$label <- gsub(pattern, new_ids[i], all_models$label)
      }
      
      rmses <- subset(all_models, all_models$label != "ROOT")
      
      rmses <- rmses %>% mutate(rgroup = ifelse(label %in% c("A1", "C1", "A-stem", "C-stem"), "AC",
                                                ifelse(label %in% c("B1", "D1", "B-stem", "D-stem"), "BD",
                                                       "Int")))
      rmses$rgroup <- factor(rmses$rgroup)
      levels(rmses$rgroup) <- c("AC", "BD", "Int")
      
      true_values <- c(`A1` = 0.5, `A-stem` = 0.5,
                       `B1` = 0.5, `B-stem` = 0.5,
                       `C1` = 0.5, `C-stem` = 0.5,
                       `D1` = 0.5, `D-stem` = 0.5,
                       `C+D` = 0.25, `A+B` = 0.25)
      rmses$true_values <- true_values[rmses$label]
      
      rmses <- rmses %>% group_by(rgroup, Model) %>% summarize(rmse = sqrt(mean((edge.length - true_values)^2)))
      rmses <- rmses %>% pivot_wider(names_from = rgroup, values_from = rmse)
      rmses$Model <- factor(rmses$Model, levels = c("UML", "GFF"))
      
      all_models$label <- factor(all_models$label, levels =  new_ids)
      all_models$edge.length <- ifelse(grepl("\\+", all_models$label),
                                        all_models$edge.length - 0.25,
                                        all_models$edge.length - 0.5)
      all_models$Group <- ifelse(grepl("\\+", all_models$label), "Internal",
                                  ifelse(grepl("stem", all_models$label), "Stem", "Terminal"))
      all_models$Model <- factor(all_models$Model, levels = c("UML", "GFF"))
      all_models$Group <- factor(all_models$Group, levels = c("Terminal", "Stem", "Internal"))
  }
    
  else if (ntaxa == 4) {
    
    rmses <- subset(all_models, all_models$label != "ROOT")
    
    rmses <- rmses %>% mutate(rgroup = ifelse(label %in% c("A", "C"), "AC", ifelse(label %in% c("B", "D"), "BD", "Int")))
    rmses$rgroup <- factor(rmses$rgroup)
    levels(rmses$rgroup) <- c("AC", "BD", "Int")
    
    true_values <- c(A = 1.0, B = 1.0, C = 1.0, D = 1.0, `C+D` = 0.25, `A+B` = 0.25)
    rmses$true_values <- true_values[rmses$label]
    
    rmses <- rmses %>% group_by(rgroup, Model) %>% summarize(rmse = sqrt(mean((edge.length - true_values)^2)))
    rmses <- rmses %>% pivot_wider(names_from = rgroup, values_from = rmse)
    rmses$Model <- factor(rmses$Model, levels = c("UML", "GFF"))
    
    all_models <- subset(all_models, all_models$label != "ROOT")
    
    all_models$edge.length <- ifelse(grepl("\\+", all_models$label),
                                     all_models$edge.length - 0.25,
                                     all_models$edge.length - 1)
    
    all_models$Group <- ifelse(grepl("\\+", all_models$label), "Internal", "Terminal")
    all_models$Model <- factor(all_models$Model, levels = c("UML", "GFF"))
    all_models$label <- factor(all_models$label, levels = c("A", "B", "C", "D",
                                                                "A+B", "C+D"))
    all_models$Group <- factor(all_models$Group, levels = c("Terminal", "Internal"))
  }
  
  bplot <- ggplot(all_models, aes(y = label, x = edge.length)) +
      geom_vline(xintercept = 0, linetype = "dotted", color = "grey", size = 1) +
      geom_point(aes(shape = Model, color = Model), stat = "summary", fun = "mean", size = 3, position = position_dodge(width = 0.9)) +
      stat_summary(
        aes(group = Model),
        fun.data = function(x) {
          mean_val <- mean(x)
          sd_val <- sd(x)
          ymin <- mean_val - sd_val
          ymax <- mean_val + sd_val
          return(data.frame(y = mean_val, ymin = ymin, ymax = ymax))
        },
        geom = "errorbarh",
        position = "dodge"
      ) +
      scale_shape_manual(name = "Model", labels = c("UML", "GFF"),
                         values = c(21, 25)) +
      scale_color_manual(name = "Model", labels = c("UML", "GFF"),
                         values=c('#44ff65', '#db44ff')) +
      theme_classic() +
      labs(x = "Branch Length: Estimated - True",
           y = "Branch") +
      scale_y_discrete(limits = rev) +
      facet_wrap(~Group, ncol = 1, scales = "free_y") + theme(legend.position = "bottom")
  
  return(list("rmse" = rmses, "plot" = bplot))
}

###############################################################################
# Note! This script will generate figures like in the accompanying manuscript.#
# Run this in RStudio!                                                        #
###############################################################################

# Figure 1: Draw simulating trees.
fig_1 <- simulatingTrees()

# Figures 2 and 3: Draw comparison of 16/4-taxa branch length and node frequency plots.
bl_16 <- compareBranchLengths(ntaxa = 16)
bl_4 <- compareBranchLengths(ntaxa = 4)
nf_16 <- compareNodeFreqs(ntaxa = 16)
nf_4 <- compareNodeFreqs(ntaxa = 4)

fig_2 <- bl_16$plot | nf_16$plot
fig_3 <- bl_4$plot | nf_4$plot

# Generate tables for RMSEs of branch lengths and node frequencies.
tab_2 <- bl_16$rmse
tab_3 <- bl_4$rmse
tab_4 <- nf_16$rmse
tab_5 <- nf_4$rmse

# Figures 5 and 6: Assess patterns of G/O/F assignment across different ID methods.
optimization_16 <- read.table("OptSummary.txt", sep = "\t", header = TRUE)
optimization_4 <- read.table("OptDownSummary.txt", sep = "\t", header = TRUE)
colnames(optimization_16) <- colnames(optimization_4) <- c("G", "O", "F", "Criterion", "Optimizer",
                                                           "Iterations",
                                                           "Iteration time", "Optimization time",
                                                           "Swap times")
fig_5 <- assessPatterns(optimization_16)
fig_6 <- assessPatterns(optimization_4)

# Figure 9: Draw tested topologies for empirical datasets.
fig_7 <- empiricalTrees()

# Figure 10: Draw G/O/F frequencies as a function of branch length under GFmix.
fig_8 <- blenVsFreq()

# Figures S1 and S2: optimization times and iterations.
fig_s1 <- assessOpts(optimization_16)
fig_s2 <- assessOpts(optimization_4)

# Add code for tree visualizations if necessary.

ggsave("Figure1_SimulatingTrees.pdf", fig_1, width = 210, height = 297, units = "mm", dpi = 300)
ggsave("Figure2_PerformancePlot.pdf", fig_2, width = 210, height = 297, units = "mm", dpi = 300)
ggsave("Figure3_4TaxaPerformancePlot.pdf", fig_3, width = 210, height = 297, units = "mm", dpi = 300)
ggsave("Figure5_16TaxonBinning.pdf", fig_5, width = 210, height = 297, units = "mm", dpi = 300, device = cairo_pdf)
ggsave("Figure6_4TaxonBinning.pdf", fig_6, width = 210, height = 297, units = "mm", dpi = 300, device = cairo_pdf)
ggsave("Figure7_NucleomorphTrees.pdf", fig_7, height = 210, width = 297, units = "mm", dpi = 300)
ggsave("Figure8_FrequencyPlot.pdf", fig_8, width = 210, height = 297, units = "mm", dpi = 300)

ggsave("FigureS1_16TaxonTimings.pdf", fig_s1, width = 210, height = 297, units = "mm", dpi = 300)
ggsave("FigureS2_4TaxonTimings.pdf", fig_s2, width = 210, height = 297, units = "mm", dpi = 300)
```
