## Supplementary material for "Modeling site-and-branch-heterogeneity with GFmix": QuickBinomial.R

```
#!/usr/bin/env Rscript
source("CommonFunctions.R")

# Handling arguments.
args <- commandArgs()
iarg <- length(args)
seqfile <- format <- outfile <- ""
while(iarg>=3){
  if(substring(args[iarg],1,1)=='-'){
    opt <- args[iarg]
    is.opt <- TRUE
  }else{
    val <- args[iarg];
    is.opt <- FALSE
  }
  if(is.opt){
    not.an.option <- TRUE
    if(opt=="-aln"){
      seqfile <- val; not.an.option <- FALSE
    }
    if(opt=="-fmt"){
      frmt <- val; not.an.option <- FALSE
    }
    if(opt=="-out"){
      outfile <- val; not.an.option <- FALSE
    }
  }
  iarg <- iarg-1
}

# Read alignment and count amino acids.
alignment <- readAln(seqfile, format = frmt)
counts <- countAA(alignment)

# Run chi2 test and make hclust tree based on residuals.
chi_residuals <- chisq.test(counts)$residuals
chi_dend <- hclust(dist(chi_residuals), method = "average")

# Run binomial test and present results.
binomial_test <- binomialBinning(chi_dend, counts)
write.table(binomial_test, file = outfile, sep = "\t", quote = FALSE, row.names = FALSE)
```
