## Supplementary material for "Modeling site-and-branch-heterogeneity with GFmix": README

# Scripts for McCarthy, Susko and Roger (2025).

## Introduction

This folder contains R scripts used in several analyses involving the GFmix model for McCarthy, Susko and Roger (2025). There are a number of R scripts in the main folder, of which the following in **bold** are executable:

-   **SimulateHeteroData.R**: simulate a 16-taxon multiple sequence alignment under a site-and-branch heterogeneous LG+C20+G{0.5}+GFmix{GARP/FYMINK} mixture model, with the option to also generate a 4-taxon down-sampling of the same alignment.

-   **QuickBinomial.R**: perform a quick binomial test on a multiple sequence alignment to determine amino acid heterogeneity to be modelled via GFmix.

-   **IdentifyHeterogeneity.R**: perform a more precise optimization procedure to determine amino acid heterogeneity to be modelled via GFmix.

-   **RunGFmix.R**: run one of four implementations of the GFmix model tested in the manuscript, providing an input alignment, tree, model and preferred amino acid compositional heterogeneity (default: GARP/FYMINK).

To replicate the analysis on simulated data in McCarthy, Susko and Roger (2025), I've provided a number of shell scripts in the "**ShellScripts**" folder which will automate most of the tasks involved. This is done so sequentially, so expect some of the scripts involving more computationally-intensive parts (e.g. likelihood calculations) to take some time. These scripts should be run from within the main folder. Scripts are listed in the order that their tasks occur in the manuscript:

-   **Simulate100Alignments.sh**: this will generate 100 simulated LG+C20+G+{GARP/FYMINK} alignments using SimulateHeteroData.R. Output alignments will be in the folder Simulations, output AliSim data for simulated alignment 1 will be in the folder Simulations/001, et cetera.

-   **RunIQTree.sh**: this will run IQ-Tree 3 on all simulated and downsampled alignments, fitting a LG+C20+G model. This is a required step for the next two parts. Output for simulated alignment 1 will be in the folder Simulations/001_iqtree/, output for downsampled alignment 1 will be in the folder Simulations/001_downsampled_iqtree, et cetera.

-   **RunModels.sh and RunModelsDownsampled.sh**: these will run each implementation of GFmix separately on all simulated and downsampled alignments. Output for simulated alignment 1 will be in the folders Simulations and Simulations/001_performance/, output for downsampled alignment 1 will be in the folders Simulations and Simulations/001_downsampled_performance/, et cetera.

-   **OptimizeGF.sh and OptimizeGFDownsampled.sh**: these will run each G-class / F-class identification and optimization routine separately on all simulated and downsampled alignments. Output for simulated alignment 1 will be in the folders Simulations and Simulations/001_performance/, output for downsampled alignment 1 will be in the folders Simulations and Simulations/001_downsampled_performance/, et cetera.

-   **SummarizeOptims.sh**: this will summarize the results of each optimization for all simulated and downsampled alignments into TSV files. Output for all simulated alignments will be OptSummary.txt and output for all downsampled alignments will be OptDownSummary.txt.

Finally, this a plotting script - **MakePlots.R** -that will produce calculations and plots similar to those in the manuscript using much of the output generated by the above shell scripts. I recommend interacting with this script via RStudio or some other IDE and not on the command line.

More information on the main R scripts themselves follows below.

## SimulateHeteroData.R

### Usage

```         
$ SimulateHeteroData.R -aln <alignment_name> -fmt <phylip | fasta> [-downsample] [-trees]
```

### Command-line options

**-aln (default: test)**: Filename for the output alignment(s). Can be provided with or without a filename extension. Will also be the name of an output folder containing all input files for AliSim and log files generated in the background. Supports writing output to new folders - e.g. "simulations/test" will also create a folder called "simulations" if the latter doesn't exist.

**-fmt (default: phylip)**: Format of output alignment(s). Either sequential PHYLIP or FASTA permitted.

**-downsample**: If provided on the command line, script will also generate downsampled 4-taxon alignment of the original 16-taxon alignment in the same format.

**-trees**: If provided, output simulating tree (and downsampled tree if "-downsample" is provided) without branch lengths for use as user fixed tree for IQTREE (-te option).

### Output

1)  A 16-taxon 10,000-site alignment in FASTA or PHYLIP format simulated under a LG+C20+G{0.5}+GFmix{GARP/FYMINK} model.

2)  If option "-downsample" is specified, a 4-taxon 10,000-site alignment downsampled from 1).

3)  A folder of the same name as 1) containing all input files for AliSim and output log files from AliSim. If "-trees" is provided, this folder with also contain the simulating tree (and downsampled tree if "-downsampled" is provided) in Newick format.

### Details

This script simulates a 16-taxon alignment under a site-and-branch heterogeneous LG+C20+G{0.5}+GFmix{GARP/FYMINK} mixture model, with the option to also generate a 4-taxon down-sampling of the same alignment. Simulation is performed using the AliSim component of the IQTREE software package.

As simulating branch-heterogeneity is not currently a native function of AliSim, the branch-heterogeneity component of the simulation is performed using a custom procedure. This procedure is detailed in the **Supplementary Information** of McCarthy, Susko and Roger (2025).

This script is used to generate 16-taxon simulated alignments to assess 1) the performance of different implementations of the GFmix model and 2) the performance of different G/F class identification methods in McCarthy, Susko and Roger (2025). The down-sampled 4-taxon alignments are used to perform similar assessments under more sparse taxon sampling conditions.

## QuickBinomial.R

### Usage

```         
$ QuickBinomial.R -aln <alignment_file> -fmt <phylip | phylip-interleaved | fasta> -out <output_file>
```

### Command-line options

**-aln (required)**: Path to multiple sequence alignment file.

**-fmt (required)**: Format of MSA file - sequential or relaxed PHYLIP (default), interleaved PHYLIP or FASTA.

**-out**: Path to output file, print output to screen if not provided.

### Output

The output of this script is a tab-separated table containing three columns as in the simplified example below:

```         
AA  Z-score Assignment
A   10.29   G-class
D   1.73    O-class
N   -6.4    F-class
```

where **AA** is an amino acid, ***Z*****-score** is the result of the binomial test for that amino acid, and **Assignment** indicates whether that amino acid is assigned to the **G-**, **O-** or **F-class** for GFmix.

### Details

This script performs a simple binomial test of two proportions for a multiple sequence alignment to determine which amino acids are compositionally heterogeneous across sequences for GFmix (Baker et al 2024, Williamson et al 2025). The counts of all 20 amino acids are tabulated for the input MSA file. A Pearson's χ^2^ test is then performed on these counts, and the residuals of this test are extracted. These residuals are converted into a distance matrix and used to hierarchically cluster sequences using UPGMA. The resulting dendrogram is cut in two, to provide the two groups of sequences needed for the binomial test.

The binomial test of two proportions measures the difference in the number of amino acids across the two groups of sequences as a *Z*-score. Amino acids with *Z* \> 1.96 are assigned as "**G-class**" amino acids for GFmix. Amino acids with *Z* \< -1.96 are assigned as "**F-class**" amino acids for GFmix. The **G-** and **F-classes** are the classes of amino acids expected to be compositionally heterogeneous across sequences. All remaining amino acids are assigned as "**O-class**" amino acids for GFmix, and are expected to be compositionally homogeneous across sequences.

## IdentifyHeterogeneity.R

### Usage

```         
$ IdentifyHeterogeneity.R -aln <alignment_file> -fmt <phylip | phylip-interleaved | fasta> -crit <bi | chi | ssf | ogf> -out <output_file> [-group <group_file> -iqtree <iqtree_file> -tree <tree_file> -root <root_file> -mix <mixture_file>]
```

### Command-line options

The following options apply to all criteria:

**-aln (required)**: Path to multiple sequence alignment file.

**-fmt (required)**: Format of MSA file - sequential or relaxed PHYLIP (default), interleaved PHYLIP or FASTA.

**-crit (default = chi)**: Criterion used to identify heterogeneous G/F classes. bi = Binomial test, chi = χ² test, ssf = sum-of-squares calculation from SSF implementation of GFmix, ogf = likelihood calculation from OGF implementation of GFmix. If bi is selected, the binomial test will proceed as in **QuickBinomial.R** above unless a groupfile is specified via -group. All other criteria will first perform a binomial test to get a starting G/F, and then perform an optimization of G/F class identification such that G/F returns a minimum (ssf) or maximum (chi, ogf) criterion value.

**-group (optional):** Path to file listing all taxa in 0-index numerical taxon IDs order to be assigned to group 1 for the binomial test. All other taxa in the alignment are assigned to group 2. The test will then proceed as outline in **QuickBinomial.R** above.

**-out**: Path to output file, print output to screen if not provided.

For tree/model-dependent criteria SSF and OGF, the following options are required:

**-iqtree:** Path to IQTREE report file, used to extract mixture weights and alpha from initial branch-homogeneous model fitting.

**-tree:** Path to IQTREE tree file in Newick format.

**-root:** Path to file specifying all 0-index numerical taxon IDs on one side of the root split of the tree.

**-mix:** Path to mixture frequency file defined in PAML format. See e.g. /bin/C10.aafreq.dat and similar files for examples.

### Output

The output of this script is a tab-separated table containing nine columns as in the example below:

```         
G	O	F	Criterion	Optimizer	Iterations	Iteration time	Optimization time	Swap times
INKSFYC	LW	THQDMPGERVA	-297403.457148472	GFmix	1	20.5378649234772	20.5381350517273	6.69813680648804,7.07171702384949,6.13255596160889,6.89132213592529,6.14467668533325,7.21630525588989,7.22756910324097,6.21991729736328,6.12342095375061,7.5722804069519,7.48150897026062,6.68500685691833,6.30993390083313,7.1081178188324,6.22749519348145,6.92600727081299,6.91627287864685,6.21400308609009,6.30487322807312,6.70887923240662,6.33306813240051,5.87276887893677
INKFYC	SLW	THQDMPGERVA	-297304.861787549	GFmix	2	20.2848722934723	40.8244557380676	6.12418437004089,6.02716970443726,7.67252635955811,6.2017548084259,6.25790357589722,7.46998977661133,7.21854448318481,6.35003447532654,7.22700953483582,7.18159770965576,6.07644891738892,6.88717651367188,6.00872111320496,6.13095831871033,6.70149421691895,6.65720152854919,6.71095514297485,6.04676532745361,6.84607219696045,7.07174921035767,5.90395402908325,6.1761839389801,6.59189343452454
INKFYC	SLWD	THQMPGERVA	-297282.982375572	GFmix	3	20.5320293903351	61.3579752445221	6.47557616233826,7.09872388839722,7.14146637916565,7.24875950813293,6.0775625705719,7.11838126182556,6.19135355949402,6.33416557312012,6.1560595035553,7.79260945320129,6.31776738166809,7.06242704391479,5.99902677536011,6.79112243652344,7.17432618141174,7.04719638824463,6.07590007781982,6.27450609207153,6.37050318717957,6.73439121246338,6.42640733718872,6.36274027824402,5.88187694549561,5.69265222549438
INKFYC	SLWDE	THQMPGRVA	-297276.69162113	GFmix	4	20.3952906131744	81.754460811615	6.04374837875366,7.05961346626282,7.38548278808594,7.17769002914429,7.29365706443787,6.13704299926758,6.17062592506409,7.14532494544983,6.16820573806763,6.10510230064392,6.04489731788635,6.00628995895386,6.63558959960938,6.07484149932861,6.78895688056946,6.74252796173096,7.01699233055115,6.12856340408325,7.04669499397278,6.23496556282043,6.46155619621277,5.88197803497314,6.36025285720825,5.79745721817017,6.30330538749695
INKFYC	SLWDE	THQMPGRVA	-297276.69162113	GFmix	5	20.8720452785492	102.627875566483	7.0819718837738,6.12668251991272,6.08473467826843,6.30716323852539,6.73491859436035,7.09119820594788,6.09684181213379,6.89634132385254,7.12700629234314,7.22654986381531,7.6830906867981,6.22978925704956,7.17844367027283,6.18623828887939,6.95319533348083,6.15571475028992,6.19138741493225,6.15588641166687,7.05212783813477,6.83798933029175,6.10110259056091,7.00507116317749,5.7416775226593,5.94215822219849,6.5214831829071,6.53870368003845
```

where **G, O** and **F** are the optimum G/O/F classes at a given step in the optimization procedure, **Criterion** is the criterion value, **Optimizer** is the criterion being optimized, **Iterations** is the current iteration of the optimization procedure, **Iteration time** is the time in seconds for that iteration in the optimization procedure, **Optimization time** is the time in seconds to date for the optimization procedure and **Swap times** are the list of times in seconds for assessment of each candidate G/O/F class for a given iteration in the optimization procedure. The final line in this file is the optimized G/O/F class for the input tree/model under a given optimization criterion.

### Details

This script is used to determine the most compositionally-heterogeneous G/F classes for a given alignment (with optional information from a corresponding tree and protein evolution model) for modeling with GFmix. In McCarthy, Susko and Roger (2025), this script was used to assess the performance of different identification criteria on 16- and 4-taxon simulated data where the simulating G/F = GARP/FYMINK and the OGF-based identification method was applied to a real algal nucleomorph dataset.

The binomial criterion operates as in the binomial test of two proportions implemented in **QuickBinomial.R**. The χ² test criterion uses the binomial criterion as a starting point, and then seeks to assign G/O/F classes for an alignment that maximize χ² through an optimization procedure detailed in the main text of McCarthy, Susko and Roger (2025). Both criteria only require an alignment as input, and optionally a group file which manually assigns the groups used in the binomial test of two proportions.

The SSF and OGF-based criteria use the SSF and OGF implementations of the GFmix model, and thus require additional tree and model information. Both criteria use the same binomial-based starting point and optimization procedure as the χ² test criterion. See the main text of McCarthy, Susko and Roger (2025) for evaluation and discussion of the performance of each criterion.

## RunGFmix.R

### Usage

```         
$ RunGFmix.R -aln <alignment_file> -fmt <phylip | phylip-interleaved | fasta> -type <ogf | ssf | gfp | gff> -mix <mixture_file> -iqtree <iqtree_file> -tree <tree_file> -root <root_file> [-out <out_file> -conv -params <params_file> -gclass <GARP> -fclass <FYMINK>]
```

### Command-line options

**-aln (required)**: Path to multiple sequence alignment file.

**-fmt (required)**: Format of MSA file - sequential or relaxed PHYLIP (default), interleaved PHYLIP or FASTA.

**-type:** Implementation of GFmix to use. OGF = original implementation in which branch-composition parameters are estimated using a hierarchical procedure, SSF = implementation in which branch-composition parameters are optimized to reduce the sum-of-squares difference between branch and model frequencies, GFP = maximum-likelihood implementation in which branch-composition parameters are optimized, GFF = maximum-likelihood implementation in which all tree and model parameters are optimized.

**-iqtree:** Path to IQTREE report file, used to extract mixture weights and alpha from initial branch-homogeneous model fitting.

**-tree:** Path to IQTREE tree file in Newick format.

**-root:** Path to file specifying all 0-index numerical taxon IDs on one side of the root split of the tree.

**-mix:** Path to mixture frequency file defined in PAML format. See e.g. /bin/C10.aafreq.dat and similar files for examples.

**-out**: Path to output file, print output to screen if not provided.

**-params:** Path to params file, which outputs likelihood, alpha, tree parameters and mixture weights to file.

**-conv:** convert sequence IDs to 0th-index sequential numerical IDs.

**-gclass:** G-class for GFmix modeling (default = GARP).

**-fclass:** F-class for GFmix modeling (default = FYMINK).

### Output

The output of this script is a tab-separated table containing ten columns as in the example below:

```         
Alignment	Mixture	+F	Tree	Method	G	F	Likelihood	Runtime	NewGFmixInfo
Nucleomorphs/Downsampled.phy	bin/C20.aafreq.dat	FALSE	Nucleomorphs/Downsampled_NM_Apart_LG+C20+G/Downsampled_NM_Apart_LG+C20+G.treefile	OGF	THQMPGRVA	FYCINK	-297276.69162113	5.65325570106506	NA
```

where **Alignment** is the input alignment file, **Mixture** is the input mixture file, **+F** indicates whether the observed frequency class (+F) was included in the base model, **Tree** is the input tree file, **Method** is the implementation of GFmix used, **G** and **F** are the G and F classes modeled GFmix, **Likelihood** is the likelihood of the tree given the model and alignment, **Runtime** is the time taken to evaulate the likelihood and **NewGFmixInfo** is additional information taken from the output of the GFP or GFF implementations of GFmix which use maximum-likelihood parameter optimization.

### Details

This script applies one of the four GFmix model implementations - OGF, SSF, GFP or GFP - to a given alignment, tree and branch-homogeneous protein evolution model. See McCarthy, Susko and Roger (2025) for details on these implementations and the GFmix model more generally. Each implementation requires input from IQTREE to extract information (i.e. alpha, mixture weights) for the base branch-homogeneous model, as well as information on how to root the tree (as IQTREE by default outputs unrooted trees). By default, **RunGFmix.R** runs GFmix with G/F = GARP/FYMINK but any valid G/F may be provided - including G/F classes determined by **IdentifyHeterogeneity.R** above.

As each implementation of the GFmix model expects taxon IDs to be 0th-index numerical IDs, it is usually necessary (and advised) to convert sequence IDs to 0th-index sequential IDs via the -conv option. The -params option may be used to output a file containing the tree (in custom UTREE format), tree and model parameters and final likelihood under a given model. It is advised to use this option if the user wants to compare estimated parameters under different GFmix implementations.

When RunGFmix.R is called with either the GFP or GFF implementation of GFmix, the output of the L-BFGS-B optimization algorithm will be printed to screen. This may be redirected to a specific file for logging purposes using `2> <logging_file>` on the command-line. See e.g. **RunModels.sh** for an example of such a redirect.

## Important notes

These scripts have been tested primarily using MSAs in sequential, relaxed or interleaved PHYLIP format. FASTA-formatted MSAs should also be compatible, but have not been as extensively tested.

## Installation

These scripts have been written and tested on remote and desktop Linux environments, and should also work on macOS. Usage on Windows has not been tested and is not advisable. All scripts must be located in the same directory. If the scripts are not already executable on your operating system, please use `chmod +x` on each of the executable scripts to enable run permission.

These scripts depend on the `parallel` library, which is included in most R installations. **MakePlots.R** has additional dependencies which can usually be installed for the user if the script is opened through RStudio.

## References

McCarthy, Susko and Roger (2025). Modeling site-and-branch-heterogeneity with GFmix. Submitted.

Williamson, K et al (2025). A robustly rooted tree of eukaryotes reveals their excavate ancestry. *Nature* **640**, 974–981. <https://doi.org/10.1038/s41586-025-08709-5>

Baker, BA et al (2024). Expanded phylogeny of extremely halophilic archaea shows multiple independent adaptations to hypersaline environments. *Nat Microbiol* **9**, 964-975. <https://doi.org/10.1038/s41564-024-01647-4>

Munoz-Gomez, S et al (2022). A site-and-branch-heterogeneous model on an expanded dataset favor mitochondria as sister to known Alphaproteobacteria. *Nat Ecol Evol* **6**, 253-262. <https://doi.org/10.1038/s41559-021-01638-2>
