## Supplementary material for "Modeling site-and-branch-heterogeneity with GFmix": RunGFmix.R

```
#!/usr/bin/env Rscript
source("functions/CommonFunctions.R")
source("functions/GFmixFunctions.R")
source("functions/LikelihoodFunctions.R")

# Handling arguments.
args <- commandArgs()
iarg <- length(args)

seqfile <- iqtreefile <- treefile <- rootfile <- NULL
mixture <- NULL
frmt <- "phylip"
program <- "gfmix"
g_class <- "GARP"
f_class <- "FYMINK"
outfile <- stdout()
params <- NULL
plusF <- FALSE
convert <- FALSE
multis <- NULL

while(iarg>=5){
  if(substring(args[iarg],1,1)=='-'){
    opt <- args[iarg]
    is.opt <- TRUE
  }else{
    val <- args[iarg];
    is.opt <- FALSE
  }
  if(is.opt){
    not.an.option <- TRUE
    if(opt=="-aln"){
      seqfile <- val; not.an.option <- FALSE
    }
    if(opt=="-fmt"){
      format <- val; not.an.option <- FALSE
    }
    if (opt=="-type"){
      program <- val; not.an.option <- FALSE
    }
    if (opt=="-mix"){
      mixture <- val; not.an.option <- FALSE
    }
    if (opt=="-iqtree"){
      iqtreefile <- val; not.an.option <- FALSE
    }
    if (opt=="-tree"){
      treefile <- val; not.an.option <- FALSE
    }
    if (opt=="-root"){
      rootfile <- val; not.an.option <- FALSE
    }
    if (opt=="-out"){
      outfile <- val; not.an.option <- FALSE
    }
    if (opt=="-conv"){
      convert <- TRUE; not.an.option <- FALSE
    }
    if (opt=="-params"){
      params <- val; not.an.option <- FALSE
    }
    if (opt=="-gclass"){
      g_class <- val; not.an.option <- FALSE
    }
    if (opt=="-fclass"){
      f_class <- val; not.an.option <- FALSE
    }
    if (opt=="-multis"){
      multis <- val; not.an.option <- FALSE
    }
  }
  iarg <- iarg-1
}

if (convert) {
  
  convertIDs(seqfile, treefile, format)
  treefile <- paste0(treefile, "_INT")
  if (format == "phylip") {
    seqfile <- paste0(seqfile, "_INT.phy")
  }
  
}

if (program == "ogf") {
  
  run_time <- Sys.time()
  lnl <- runGFmix(seqfile, mixture, treefile, iqtreefile, rootfile, g_class,
                    f_class, plusF, params = params)
  run_time <- as.numeric(difftime(Sys.time(), run_time ,units = "secs"))
  
  result <- cbind.data.frame("Alignment" = seqfile,
                  "Mixture" = mixture,
                  "+F" = plusF,
                  "Tree" = treefile,
                  "Method" = "OGF",
                  "G" = g_class,
                  "F" = f_class,
                  "Likelihood" = lnl,
                  "Runtime" = run_time,
                  "NewGFmixInfo" = NA)
  
} else if (program == "ssf") {
  
  run_time <- Sys.time()
  lnl <- runAnFreqOpt(seqfile, mixture, treefile, iqtreefile, rootfile, g_class,
                               f_class, plusF, params)
  run_time <- as.numeric(difftime(Sys.time(), run_time ,units = "secs"))
  
  result <- cbind.data.frame("Alignment" = seqfile,
                  "Mixture" = mixture,
                  "+F" = plusF,
                  "Tree" = treefile,
                  "Method" = "SSF",
                  "G" = g_class,
                  "F" = f_class,
                  "Likelihood" = lnl,
                  "Runtime" = run_time,
                  "NewGFmixInfo" = NA)
  
} else if (program == "gfp") {
  
  run_time <- Sys.time()
  
  lnl <- runGFmixLnL(seqfile, mixture, treefile, iqtreefile, rootfile, g_class,
                      f_class, mode = 4, plusF, params, multis)
  run_time <- as.numeric(difftime(Sys.time(), run_time ,units = "secs"))
  
  result <- cbind.data.frame("Alignment" = seqfile,
                             "Mixture" = mixture,
                             "+F" = plusF,
                             "Tree" = treefile,
                             "Method" = "GFP",
                             "G" = g_class,
                             "F" = f_class,
                             "Likelihood" = lnl$lnl,
                             "Runtime" = run_time,
                             "NewGFmixInfo" = lnl$info)
  
} else if (program == "gff") {
  
  run_time <- Sys.time()
  lnl <- runGFmixLnL(seqfile, mixture, treefile, iqtreefile, rootfile, g_class,
                     f_class, mode = 6, plusF, params, multis)
  run_time <- as.numeric(difftime(Sys.time(), run_time ,units = "secs"))
  
  result <- cbind.data.frame("Alignment" = seqfile,
                             "Mixture" = mixture,
                             "+F" = plusF,
                             "Tree" = treefile,
                             "Method" = "GFF",
                             "G" = g_class,
                             "F" = f_class,
                             "Likelihood" = lnl$lnl,
                             "Runtime" = run_time,
                             "NewGFmixInfo" = lnl$info)
  
}

write.table(result, file = outfile, sep = "\t", quote = FALSE, row.names = FALSE)
```
