## Supplementary material for "Modeling site-and-branch-heterogeneity with GFmix": NucelomorphGFmix.sh

```
# 16-taxon dataset Apart

  ./RunGFmix.R -aln Nucleomorphs/Downsampled.phy -fmt phylip -type ogf -mix bin/C20.aafreq.dat \
  -iqtree Nucleomorphs/Downsampled_NM_Apart_LG+C20+G/Downsampled_NM_Apart_LG+C20+G.iqtree \
  -tree Nucleomorphs/Downsampled_NM_Apart_LG+C20+G/Downsampled_NM_Apart_LG+C20+G.treefile \
  -root Nucleomorphs/16_NM_Apart.rootfile \
  -gclass THQMPGRVA -fclass FYCINK \
  -params Nucleomorphs/Downsampled_NM_Apart_LG+C20+G+GFmix/OGF_optimized.params \
  -out Nucleomorphs/Downsampled_NM_Apart_LG+C20+G+GFmix/OGF_optimized.txt
  
  ./RunGFmix.R -aln Nucleomorphs/Downsampled.phy -fmt phylip -type ssf -mix bin/C20.aafreq.dat \
  -iqtree Nucleomorphs/Downsampled_NM_Apart_LG+C20+G/Downsampled_NM_Apart_LG+C20+G.iqtree \
  -tree Nucleomorphs/Downsampled_NM_Apart_LG+C20+G/Downsampled_NM_Apart_LG+C20+G.treefile \
  -root Nucleomorphs/16_NM_Apart.rootfile \
  -gclass THQMPGRVA -fclass FYCINK \
  -params Nucleomorphs/Downsampled_NM_Apart_LG+C20+G+GFmix/SSF_optimized.params \
  -out Nucleomorphs/Downsampled_NM_Apart_LG+C20+G+GFmix/SSF_optimized.txt
  
  ./RunGFmix.R -aln Nucleomorphs/Downsampled.phy -fmt phylip -type gfp -mix bin/C20.aafreq.dat \
  -iqtree Nucleomorphs/Downsampled_NM_Apart_LG+C20+G/Downsampled_NM_Apart_LG+C20+G.iqtree \
  -tree Nucleomorphs/Downsampled_NM_Apart_LG+C20+G/Downsampled_NM_Apart_LG+C20+G.treefile \
  -root Nucleomorphs/16_NM_Apart.rootfile \
  -gclass THQMPGRVA -fclass FYCINK \
  -params Nucleomorphs/Downsampled_NM_Apart_LG+C20+G+GFmix/GFP_optimized.params \
  -out Nucleomorphs/Downsampled_NM_Apart_LG+C20+G+GFmix/GFP_optimized.txt 2> Nucleomorphs/Downsampled_NM_Apart_LG+C20+G+GFmix/GFP_optimized.iterations;
  
  ./RunGFmix.R -aln Nucleomorphs/Downsampled.phy -fmt phylip -type gff -mix bin/C20.aafreq.dat \
  -iqtree Nucleomorphs/Downsampled_NM_Apart_LG+C20+G/Downsampled_NM_Apart_LG+C20+G.iqtree \
  -tree Nucleomorphs/Downsampled_NM_Apart_LG+C20+G/Downsampled_NM_Apart_LG+C20+G.treefile \
  -root Nucleomorphs/16_NM_Apart.rootfile \
  -gclass THQMPGRVA -fclass FYCINK \
  -params Nucleomorphs/Downsampled_NM_Apart_LG+C20+G+GFmix/GFF_optimized.params \
  -out Nucleomorphs/Downsampled_NM_Apart_LG+C20+G+GFmix/GFF_optimized.txt 2> Nucleomorphs/Downsampled_NM_Apart_LG+C20+G+GFmix/GFF_optimized.iterations;
  
```
