## Supplementary material for "Modeling site-and-branch-heterogeneity with GFmix": OptimizeGF.sh

```
for i in $(seq -w 1 100);
do
  
  mkdir Simulations/"$i"_optimization;

    ./IdentifyHeterogeneity.R -aln Simulations/$i.phy -fmt phylip \
   -crit bi -group sim_topologies/16_taxa.groupfile \
   -out Simulations/"$i"_optimization/bi_optimization.txt;

   ./IdentifyHeterogeneity.R -aln Simulations/$i.phy -fmt phylip \
    -crit chi -group sim_topologies/16_taxa.groupfile \
    -out Simulations/"$i"_optimization/chi_optimization.txt;

    ./IdentifyHeterogeneity.R -aln Simulations/$i.phy -fmt phylip \
     -crit ssf -group sim_topologies/16_taxa.groupfile \
     -iqtree Simulations/"$i"_iqtree/"$i"_iqtree.iqtree \
     -tree Simulations/"$i"_iqtree/"$i"_iqtree.treefile \
     -root sim_topologies/16_taxa.rootfile \
     -mix bin/C20.aafreq.dat \
     -out Simulations/"$i"_optimization/ssf_optimization.txt;

   ./IdentifyHeterogeneity.R -aln Simulations/$i.phy -fmt phylip \
    -crit ogf -group sim_topologies/16_taxa.groupfile \
    -iqtree Simulations/"$i"_iqtree/"$i"_iqtree.iqtree \
    -tree Simulations/"$i"_iqtree/"$i"_iqtree.treefile \
    -root sim_topologies/16_taxa.rootfile \
    -mix bin/C20.aafreq.dat \
    -out Simulations/"$i"_optimization/gfmix_optimization.txt;

done
```
