## Supplementary material for "Modeling site-and-branch-heterogeneity with GFmix": OptimizeGFDownsampled.sh

```
for i in $(seq -w 1 100);
do

  mkdir Simulations/"$i"_downsampled_optimization;

   ./IdentifyHeterogeneity.R -aln Simulations/"$i"_downsampled.phy -fmt phylip \
    -crit bi -group sim_topologies/4_taxa.groupfile \
    -out Simulations/"$i"_downsampled_optimization/bi_optimization.txt;

   ./IdentifyHeterogeneity.R -aln Simulations/"$i"_downsampled.phy -fmt phylip \
    -crit chi -group sim_topologies/4_taxa.groupfile \
    -out Simulations/"$i"_downsampled_optimization/chi_optimization.txt;

   ./IdentifyHeterogeneity.R -aln Simulations/"$i"_downsampled.phy -fmt phylip \
    -crit ssf -group sim_topologies/4_taxa.groupfile \
    -iqtree Simulations/"$i"_downsampled_iqtree/"$i"_downsampled_iqtree.iqtree \
    -tree Simulations/"$i"_downsampled_iqtree/"$i"_downsampled_iqtree.treefile \
    -root sim_topologies/4_taxa.rootfile \
    -mix bin/C20.aafreq.dat \
    -out Simulations/"$i"_downsampled_optimization/ssf_optimization.txt;

   ./IdentifyHeterogeneity.R -aln Simulations/"$i"_downsampled.phy -fmt phylip \
    -crit ogf -group sim_topologies/4_taxa.groupfile \
    -iqtree Simulations/"$i"_downsampled_iqtree/"$i"_downsampled_iqtree.iqtree \
    -tree Simulations/"$i"_downsampled_iqtree/"$i"_downsampled_iqtree.treefile \
    -root sim_topologies/4_taxa.rootfile \
    -mix bin/C20.aafreq.dat \
    -out Simulations/"$i"_downsampled_optimization/ogf_optimization.txt;
    
done
```
