## Supplementary material for "Modeling site-and-branch-heterogeneity with GFmix": RunIQTree.sh

```
for i in $(seq -w 1 100);
do
  mkdir Simulations/"$i"_iqtree;
  iqtree3 -s Simulations/$i.phy -m LG+C20+G -te sim_topologies/16_taxa.tree \
   -nt 30 --prefix Simulations/"$i"_iqtree/"$i"_iqtree;
  
  mkdir Simulations/"$i"_downsampled_iqtree;
  iqtree3 -s Simulations/"$i"_downsampled.phy -m LG+C20+G \
   -te sim_topologies/4_taxa.tree -nt 30 \
   --prefix Simulations/"$i"_downsampled_iqtree/"$i"_downsampled_iqtree;
done
```
