## Supplementary material for "Modeling site-and-branch-heterogeneity with GFmix": RunModels.sh

```
for i in $(seq -w 1 100);
do

  mkdir Simulations/"$i"_performance;

  ./RunGFmix.R -aln Simulations/"$i".phy -fmt phylip -type ogf -mix bin/C20.aafreq.dat \
  -iqtree Simulations/"$i"_iqtree/"$i"_iqtree.iqtree \
  -tree Simulations/"$i"_iqtree/"$i"_iqtree.treefile \
  -root sim_topologies/16_taxa.rootfile \
  -params Simulations/"$i"_performance/OGF.params \
  -out Simulations/"$i"_performance/OGF.txt 

  ./RunGFmix.R -aln Simulations/"$i".phy -fmt phylip -type ssf -mix bin/C20.aafreq.dat \
  -iqtree Simulations/"$i"_iqtree/"$i"_iqtree.iqtree \
  -tree Simulations/"$i"_iqtree/"$i"_iqtree.treefile \
  -root sim_topologies/16_taxa.rootfile \
  -params Simulations/"$i"_performance/SSF.params \
  -out Simulations/"$i"_performance/SSF.txt 

  ./RunGFmix.R -aln Simulations/"$i".phy -fmt phylip -type gfp -mix bin/C20.aafreq.dat \
  -iqtree Simulations/"$i"_iqtree/"$i"_iqtree.iqtree \
  -tree Simulations/"$i"_iqtree/"$i"_iqtree.treefile \
  -root sim_topologies/16_taxa.rootfile \
  -params Simulations/"$i"_performance/GFP.params \
  -out Simulations/"$i"_performance/GFP.txt 2> Simulations/"$i"_performance/GFP.iterations;

  ./RunGFmix.R -aln Simulations/"$i".phy -fmt phylip -type gff -mix bin/C20.aafreq.dat \
  -iqtree Simulations/"$i"_iqtree/"$i"_iqtree.iqtree \
  -tree Simulations/"$i"_iqtree/"$i"_iqtree.treefile \
  -root sim_topologies/16_taxa.rootfile \
  -params Simulations/"$i"_performance/GFF.params \
  -out Simulations/"$i"_performance/GFF.txt 2> Simulations/"$i"_performance/GFF.iterations;
  
done
```
