## Supplementary material for "Modeling site-and-branch-heterogeneity with GFmix": SummarizeOptims.sh

```
for i in $(seq -w 1 100); do tail -n 1 Simulations/"$i"_optimization/*_optimization.txt | grep -v "=" | grep -Ev "^$"; done > OptSummary.txt

sed -i "1s/^/G	O	F	Criterion	Optimizer	Iterations	Iteration time	Optimization time	Swap times\n/" OptSummary.txt

for i in $(seq -w 1 100); do tail -n 1 Simulations/"$i"_downsampled_optimization/*_optimization.txt | grep -v "=" | grep -Ev "^$"; done > OptDownSummary.txt

sed -i "1s/^/G	O	F	Criterion	Optimizer	Iterations	Iteration time	Optimization time	Swap times\n/" OptDownSummary.txt
```
