## Supplementary material for "Modeling site-and-branch-heterogeneity with GFmix": SimulateHeteroData.R

```
#!/usr/bin/env Rscript
source("functions/CommonFunctions.R")
source("functions/SimulationFunctions.R")

# Handling arguments.
args <- commandArgs()
iarg <- length(args)

ntaxa <- 16
nsites <- 10000
alpha <- 0.5
k <- 4
g_class <- strsplit1("GARP")
f_class <- strsplit1("FYMINK")
downsample <- FALSE
format <- "phylip"
aln_name <- "test"
trees <- FALSE

while(iarg>=4){
  if(substring(args[iarg],1,1)=='-'){
    opt <- args[iarg]
    is.opt <- TRUE
  }else{
    val <- args[iarg];
    is.opt <- FALSE
  }
  if(is.opt){
    not.an.option <- TRUE
    if(opt=="-aln"){
      aln_name <- val; not.an.option <- FALSE
    }
    if(opt=="-fmt"){
      format <- val; not.an.option <- FALSE
    }
    if(opt=="-downsample"){
      downsample <- TRUE; not.an.option <- FALSE
    }
    if(opt=="-trees"){
      trees <- TRUE; not.an.option <- FALSE
    }
  }
  iarg <- iarg-1
}

tree <- generateTree(ntaxa = ntaxa)
rates <- generateGammaRates(alpha = alpha, k = k)
mixture <- generateMixture(mixture = "C20")

# Resample mixture weights.
mixture <- resampleProfileSites(mixture, nsites)

# Generate models with class frequencies where GARP/FYMINK = 1 and 0.1.
equal <- modifyMixture(mixture = mixture, g_class = g_class,
                         f_class = f_class, comp_ratio = 1)
fymink <- modifyMixture(mixture = mixture, g_class = g_class,
                        f_class = f_class, comp_ratio = 0.1)

# Generate newick and nexus files and run AliSim for each mixture profile.
alisim_dir <- runAliSim(tree, ntaxa, nsites, rates,
                        alpha, equal, fymink, aln_name)

# Concatenate profile alignments into one single alignment.
simulated_aln <- concatenateAlignments(alisim_dir, aln_name)
output <- writeAln(simulated_aln, aln_name, format)

prnt <- paste0("Written concatenated alignment to file: ./",
               output, ". AliSim I/O files are in the folder: ./", aln_name, ".")
write(prnt, stdout())

if (trees) {
  tree <- generateTree(ntaxa = 16, exclude_bl = TRUE)
  write(tree, file = paste0(aln_name, "/16_taxa.tree", collapse = ""))
  
  prnt <- paste0("Written simulating tree topology ",
                 "(w/o branch length) to", aln_name ,"/16_taxa.tree.")
  write(prnt, stdout())
  
}

if (downsample) {
  daln_name <- paste0(aln_name, "_downsampled", collapse = "")
  downsampled_aln <- downsampleAlignment(simulated_aln)
  output <- writeAln(downsampled_aln, daln_name, format)
  
  prnt <- paste0("Written downsampled alignment to file: ./",
                 output, ".")
  write(prnt, stdout())
  
  if (trees) {
    tree <- generateTree(ntaxa = 4, exclude_bl = TRUE)
    write(tree, file = paste0(aln_name, "/4_taxa.tree", collapse = ""))
    
    prnt <- paste0("Written downsampled tree topology ",
                   "(w/o branch length) to", aln_name ,"/4_taxa.tree.")
    write(prnt, stdout())
  }

}
```
